# Hypoxia-conditioned HNSCC cell line secretomes drive phenotypic, functional, and transcriptional reprogramming of human neutrophils

**DOI:** 10.64898/2026.07.31.741959

**Authors:** Jana Pereckova, Filip Zavadil Kokas, Simona Voznicova, Tamara Kolarova, Roman Hrstka, Ondrej Vasicek, Tomas Perecko

## Abstract

Neutrophils display marked functional plasticity in cancer; however, it remains poorly understood how soluble factors derived from hypoxic and irradiated head and neck squamous cell carcinoma (HNSCC) cells reprogram neutrophil phenotype and function.

Here, we employed a well characterized and controlled *in vitro* model to examine how tumor-conditioned media (TCM) from HNSCC cell lines cultured under ambient (21% O_2_) or hypoxic (1% O_2_) conditions, with or without 6 Gy gamma irradiation, modulate human neutrophil phenotype, and functional and transcriptional responses. Initial analyses were performed using TCM from three different HNSCC cell lines, whereas subsequent mechanistic characterization focused on FaDu-derived TCM.

We show that TCM prolongs neutrophil survival in a cell line–dependent manner. Among the tested cell lines, hypoxia-conditioned FaDu-derived TCM promoted immunomodulatory neutrophil state characterized by enhanced survival, selective priming of ROS production, and elevated TRAIL-R3/TRAIL-R2 ratio. Induction of classical activation markers (CD11b, CD62L) was not evident. Transcriptomic analysis revealed minimal effects of normoxic TCM. Hypoxia-conditioned TCM induced a pronounced transcriptional program enriched in hypoxia- and stress-associated pathways. In contrast, irradiation of tumor cells had a limited additional impact on neutrophil reprogramming.

Together, these findings indicate hypoxia-conditioned tumor secretomes as important drivers of neutrophil functional adaptation *in vitro*, supporting a model in which soluble factors alone are sufficient to induce a persistent, immunomodulatory neutrophil phenotype. This work provides mechanistic insight into tumor–neutrophil crosstalk, highlighting hypoxia-driven signaling as a potential therapeutic target in radioresistant HNSCC and supporting a role for neutrophil reprogramming in this context.

## Introduction

Neutrophils are among the most abundant leukocytes in blood and are increasingly recognized as important regulators of cancer biology, rather than merely short-lived immune effectors (reviewed in [1]). Tumor associated neutrophils (TANs) were initially conceptualized within a dichotomous framework as either anti-tumoral (N1) or pro-tumoral (N2) populations [2]. However, current evidence indicates that TANs exhibit considerable functional, phenotypic, and transcriptional plasticity, and are better understood as a continuum of states shaped by the tumor microenvironment (TME) (reviewed in [3]). Depending on the local TME, neutrophils may contribute to anti-tumor responses through direct cytotoxicity – e.g. production of neutrophil extracellular traps (NETs) [4] or reactive oxygen species (ROS) [5]. However, neutrophils can contribute to pro-tumoral effects by immune suppression [5–7], enhancing angiogenesis (reviewed in [8]), and supporting metastatic spread [9], indicating double-edged sword of neutrophil defense mechanisms (e.g., ROS, NETs – [9, 10]). This makes TANs context-dependent immune populations in solid tumors, including head and neck squamous cell carcinoma (HNSCC).

HNSCC is in the top ten most common cancers worldwide [11]. Based on human papillomavirus (HPV) status, HNSCC can be classified as HPV-negative tumors, generally associated with tobacco or alcohol abuse, or HPV-positive tumors, primarily linked to HPV-16 infection [12]. HPV oncoproteins E6 and E7 inhibit the functions of p53 and the retinoblastoma tumor suppressor protein, respectively [13].

From a therapeutic perspective, more than 50% of patients with HNSCC have an indication for radiotherapy [14]. Hypoxia is a common feature of HNSCC tumors and contributes to poor prognosis and reduced sensitivity to radiotherapy by limiting oxygen-dependent ROS formation and enhancing DNA repair mechanisms (reviewed in [12]),[15]. Hypoxic conditions stabilize the hypoxia-inducible factor 1-alpha (HIF-1α) transcription factor, a key regulator of cellular adaptive responses to low oxygen levels (reviewed in [16]). Hypoxia advances tumor development, progression, and metastasis (reviewed in [17]). HPV-positive HNSCC cells generally exhibit increased radiosensitivity compared to HPV-negative cells; however, under hypoxic conditions, they show comparable relative radioresistance [18].

Hypoxia also affects immune components of the TME, particularly neutrophils. Low oxygen conditions can prolong neutrophil lifespan [19]. In a glioblastoma study, infiltration of neutrophils was increased in the group with high HIF-1α expression [20]. In a mouse model, neutrophils within hypoxic tumors persist longer and promote angiogenesis [21], supporting the concept that tumor-derived signals and oxygen conditions within TME drive neutrophil reprogramming. Glucose transporter type 1, a marker of hypoxia, was strongly expressed in TANs and its deletion increased tumor sensitivity to radiotherapy [22]. These features make hypoxic TME a relevant determinant of immune regulation in cancer.

However, despite growing interest in tumor neutrophils, the combined impact of hypoxia and radiotherapy-related tumor conditioning on neutrophils in HNSCC remains unclear. In our previous work, we characterized the effects of hypoxia and gamma irradiation on the secretome of HNSCC cell lines [23]; here, we extend these findings to investigate how tumor-conditioned medium (TCM) derived from HNSCC cells cultured under ambient or hypoxic conditions, with or without gamma irradiation, modulate human neutrophil phenotype (programmed death-ligand 1 (PD-L1), CD170 (Siglec-5), tumor necrosis factor– related apoptosis-inducing ligand receptors (TRAIL-Rs)), function (apoptosis, ROS, migration), and transcriptional activity. For TCM generation, we used three different HNSCC cell lines: FaDu (primary origin), Detroit-562 (metastatic origin), and 2A3, an isogenic derivative of FaDu stably expressing HPV-16 E6/E7 oncogenes.

HNSCC-derived TCM prolonged neutrophil survival in a cell line- and oxygen-dependent manner, and hypoxic conditioning selectively altered neutrophil ROS priming, and the expression of selected immunoregulatory surface markers. RNA sequencing of neutrophils exposed to FaDu-derived TCM showed that normoxic TCM induced only limited transcriptional changes, whereas short-term exposure to hypoxia-conditioned TCM triggered a broader transcriptional response. Overall, our study provides a framework linking soluble tumor-derived signals to neutrophil phenotypic and transcriptional adaptation.

## Material and Methods

### HNSCC cell culture and tumor-conditioned media (TCM)

TCM were generated as previously described [23]. Briefly, FaDu (primary origin), Detroit-562 (metastatic origin) and 2A3 (FaDu cells stably expressing HPV-16 E6 and E7 oncogenes) cells were cultured under normoxic (21% O_2_, 5% CO₂) or hypoxic (1% O_2_, 5% CO_2_, 94% N_2_; InvivO_2_ 400, Baker Ruskinn, UK) conditions. Cells (2×10^5^) were plated in 6-well plates in 1.5 ml of RPMI medium supplemented with 10% low-endotoxin fetal bovine serum (LE-FBS). The next day, cells were irradiated with 0 or 6 Gy of gamma rays (dose rate 1 Gy/min) using ⁶⁰Co gamma-ray source (Chisostat, Czech Republic). Hypoxic samples were placed in airtight pouches during irradiation. After irradiation, the medium was replaced with fresh oxygen-matched RPMI with 10% LE-FBS, and cells were incubated under their respective oxygen conditions for 48 hours. TCM was aspirated, centrifuged at 500×g for 10 minutes at 4°C to remove cell debris, and transferred to new tubes. Samples were processed under the corresponding oxygen conditions, aliquoted, and stored at −20°C until use. Most TCM batches had been previously characterized in our secretome study [23], while additional batches were generated using the same protocol for experiments performed subsequently.

### Neutrophil isolation and culture with TCM

Neutrophils were isolated as previously [24]. Buffy coats were obtained from healthy donors at the Department of Transfusion and Tissue Medicine, University Hospital Brno, under agreement approved by the Ethics Committee of UH Brno and in accordance with the Declaration of Helsinki principles. Cells were resuspended in RPMI with 10% LE-FBS. Neutrophils were adjusted to 3.5 × 10⁶ cells/ml and incubated 1:1 with TCM (final volume 200 μl) at 37°C in a humidified CO₂ incubator under ambient conditions for the indicated times before functional or phenotypic analyses, using tubes sealed with Parafilm. Control cells were cultured in RPMI with LE-FBS only.

### Neutrophil viability

Neutrophils were incubated with TCM for 24 and 72 hours at 37°C. Following incubation, samples were centrifuged (250×g, room temperature, 7 minutes). The cell pellet was resuspended in 150 µl Annexin V binding buffer containing 1 µl Annexin V–FITC (Exbio, Czech Republic) and incubated for 20 minutes at room temperature in the dark. Samples were then washed, and the pellet was resuspended in 150 µl Annexin V binding buffer and kept on ice. Immediately before acquisition, 2 µl propidium iodide (PI; Sigma-Aldrich, Germany) was added, samples were mixed, and analyzed on SONY SP6800 spectral flow cytometer (Sony Biotechnology, USA).

### Correlation of IL-8 concentration in TCM with neutrophil viability

IL-8 concentrations in TCM were measured previously using the Human IL-8/CXCL8 DuoSet ELISA kit (#DY208, Bio-Techne, USA) and published in our earlier work [23]. Previously obtained IL-8 values corresponding to the same TCM samples used in the present study were used exclusively for correlation analyses with neutrophil viability.

### Neutrophil ROS production

Neutrophils were incubated for 24 hours with and without TCM. *Fusobacterium nucleatum* (1 neutrophil:100 bacteria) was added 30 minutes before the end of incubation as the activator of ROS production. DHR123 (100 µM) was added to all samples for 10 minutes at 37°C in the dark. Cells were then washed and resuspended in 150 µl PBS containing 0.5% bovine serum albumin (BSA, Sigma-Aldrich, Germany). Samples were transported on ice and analyzed on SONY SP6800 spectral flow cytometer (Sony Biotechnology, USA).

### Assessment of neutrophil activation

Neutrophils were incubated with TCM for 2 hours at 37°C. After incubation, cells were washed and resuspended in 150 µl PBS containing 0.5% BSA. The following antibodies were added (all from Sony Biotechnology, USA): anti-CD11b (3 µl; 2107020) and anti-CD62L (3 µl; 2124050). Samples were incubated for 20 minutes at 4°C, protected from light. Cells were then washed and resuspended in 150 µl PBS + 0.5% BSA, kept on ice, and analyzed on BD FACSVerse flow cytometer (BD Biosciences, USA). Control samples consisted of neutrophils incubated in RPMI with 10% LE-FBS; for positive controls, *Fusobacterium nucleatum* suspension was added at a ratio of 1 neutrophil:100 bacteria.

### Neutrophil phenotyping

Phenotypic changes in neutrophils were analyzed after 48-hour incubation with TCM at 37°C. Following incubation, cells were resuspended in 150 µl PBS with 0.5% BSA. The following antibodies from Sony Biotechnology (USA) were added: anti-PD-L1 (2 µl; 2248570), anti-CD170 (2 µl; 2360030), anti-TRAIL-R2 (5 µl; 2137040), and anti-TRAIL-R3 (5 µl; 2135030). Samples were incubated for 20 minutes at 4°C, protected from light. After incubation, samples were washed and the cells resuspended in 150 µl PBS + 0.5% BSA. Samples were kept on ice before analysis. Data were acquired on BD FACSVerse flow cytometer (BD Biosciences, USA), and analyzed in FlowJo v10.10.1 (BD Biosciences, USA). Control neutrophils were incubated in RPMI with 10% LE-FBS, and interferon-γ (IFN-γ, 10 ng/ml) served as a positive control.

### Neutrophil migration

Neutrophil motility was monitored using time-lapse microscopy. Cells were seeded into µ-slides (Ibidi, Germany) with channels coated with 50 µl fibrinogen (100 µg/ml in RPMI) and incubated for 2 hours at 37°C (Parafilm sealed). Channels were then washed with PBS and prefilled with 50 µl RPMI before neutrophils application. Neutrophil nuclei were stained with Hoechst 33342 (4 ng/ml) for 30 minutes at 37°C in the dark. A 30 µl suspension of neutrophils (1 × 10⁶ cells/ml) was added to each channel. The slide was placed into a temperature-controlled microscope chamber (37°C) for 30 minutes to allow cells to settle. Imaging positions and parameters were set (2-minute interval, 91 frames). Subsequently, 10 µl TCM was added to one side of the channel; recombinant IL-8 (2 ng/ml) served as a positive control and RPMI as a negative control. Cell movement was tracked over 60 minutes at 37°C. Migration trajectories were quantified using the *TrackMate* plugin in ImageJ.

### RNA isolation, sequencing, and differential gene expression analysis in neutrophils

Neutrophils (15×10^6^/sample) were incubated with TCM diluted 1:1 in fresh RPMI medium with 10% LE-FBS (final volume 500 µl per sample). After a 4-hour incubation in a humidified CO_2_ incubator under ambient conditions, neutrophils were washed with PBS and lysed in 200 µl RNA later (R0901, Merck, USA). Total RNA was extracted from cells by TRI-Reagent (Sigma-Aldrich, USA). Only RNA samples with an RNA integrity number (RIN) ≥7 determined by a Bioanalyzer (RNA 6000 Nano Kit, Agilent Technologies, USA) passed to library preparation. The TruSeq Stranded Total RNA LT Sample Prep Kit (Illumina, USA) was used to convert 0.5 mg of total RNA into a library of template molecules. The library was validated using Bioanalyzer (DNA 1000 Kit, Agilent. USA) and quantified according to the manufacturer’s instructions by qPCR (KAPA Library Quantification Kit Illumina platforms, Kapa Biosystems, USA) on Quant studio 5 Real-Time PCR System (Thermo Fisher Scientific, USA). Sequencing was performed on NextSeq 500 (Illumina, USA).

Low-quality reads were removed from the raw sequencing data, adaptor sequences clipped, and low-quality leading or trailing regions (below phred score 18) were trimmed using Trimmomatic v0.33 and BBDuk2 (both from the Joint Genome Institute, USA). RNA sequencing data obtained from three biological replicates per group were aligned to the hg38 [25] reference genome using the HISAT [26] aligner. Reads mapped to gene regions were subsequently quantified with FeatureCounts [27], and differential expression analysis was performed using DESeq2 [28]. Genes were considered differentially expressed if they had an adjusted p-value (padj) <0.05. Genes which were evaluated as differentially expressed were visualized via InteractiVenn [29] and iDEP [30]. WEB-based GEne SeT AnaLysis Toolkit 2024 (WebGestalt 2024) [31], using Over-Representation Analysis based on Fisher’s exact test, was employed to analyze biological processes associated with differentially expressed genes (DEGs) including KEGG [32, 33].

### External validation of the hypoxia score in TCGA-HNSCC

Validation of the hypoxia score was performed using publicly available TCGA-HNSCC data obtained from cBioPortal [34, 35]. The principal source of expression and clinical data was the Head and Neck Squamous Cell Carcinoma (TCGA GDC, 2025) study (hnsc_tcga_gdc), containing hg38-harmonized data derived from the NCI Genomic Data Commons [36]. The published Head and Neck Squamous Cell Carcinoma (TCGA, Nature 2015) dataset (hnsc_tcga_pub), which comprised genomic characterization of 279 HNSCC tumors, was used only to supplement HPV-status and anatomical-site annotations [37]. Because the two datasets represent overlapping TCGA-HNSCC cases, the published dataset was not considered an independent validation cohort.

Only primary tumor samples were included, and one sample per patient was retained, preferentially selecting the sample with the largest sequencing library. RNA-seq counts were TMM-normalized and transformed to logCPM values using edgeR [38]. Entrez Gene identifiers were mapped to gene symbols using AnnotationDbi and org.Hs.eg.db within the Bioconductor framework [39]. The hypoxia score was calculated as the difference between the mean gene-wise z-score of genes upregulated and downregulated in the experimentally derived hypoxia signature.

For Kaplan–Meier analysis, patients were stratified into high- and low-score groups using the median hypoxia score. The association between the continuous standardized score and overall survival was evaluated using a multivariable Cox proportional-hazards model adjusted for age, sex, and pathological T and N stages. Associations with pathological N+ status and advanced T stage (T3/T4) were assessed using logistic regression adjusted for available HPV-status and anatomical-site variables. Score distributions between pathological-stage groups were compared using the Wilcoxon rank-sum test. Analyses were conducted in R using the survival [40], dplyr, edgeR [38], AnnotationDbi, org.Hs.eg.db [39], ggplot2 [41], and patchwork [42] packages.

### Statistics

Data are presented as arithmetic means ± Standard Deviation (SD), unless otherwise stated. Statistical analysis was performed using one-way ANOVA followed by appropriate post hoc tests. Comparisons between two groups were performed using Student’s t-test. A p-value of <0.05 was considered statistically significant. Statistical analyses were conducted using GraphPad Prism v10.4 (GraphPad Software, USA, www.graphpad.com). Statistical methods and the number of independent experiments are provided in the respective figure legends.

## Results

### TCM prolongs neutrophil survival in a cell line–specific manner modulated by hypoxia

TCM derived from HNSCC cell lines increased neutrophil viability compared with RPMI controls at both analyzed time points (Figure 1A–C). At 24 hours, hypoxia-conditioned FaDu-derived TCM induced a significant pro-survival effect in neutrophils (Figure 1A). Further, hypoxia-conditioned FaDu-derived TCM prolonged neutrophil lifespan compared with normoxic TCM, although this effect was observed only in the 0 Gy samples (Figure 1A). TCM derived from 2A3 (Figure 1B) and Detroit-562 (Figure 1C) cells increased neutrophil viability either non-significantly or significantly, respectively, irrespective of oxygen or radiation conditions.

**Figure 1.**
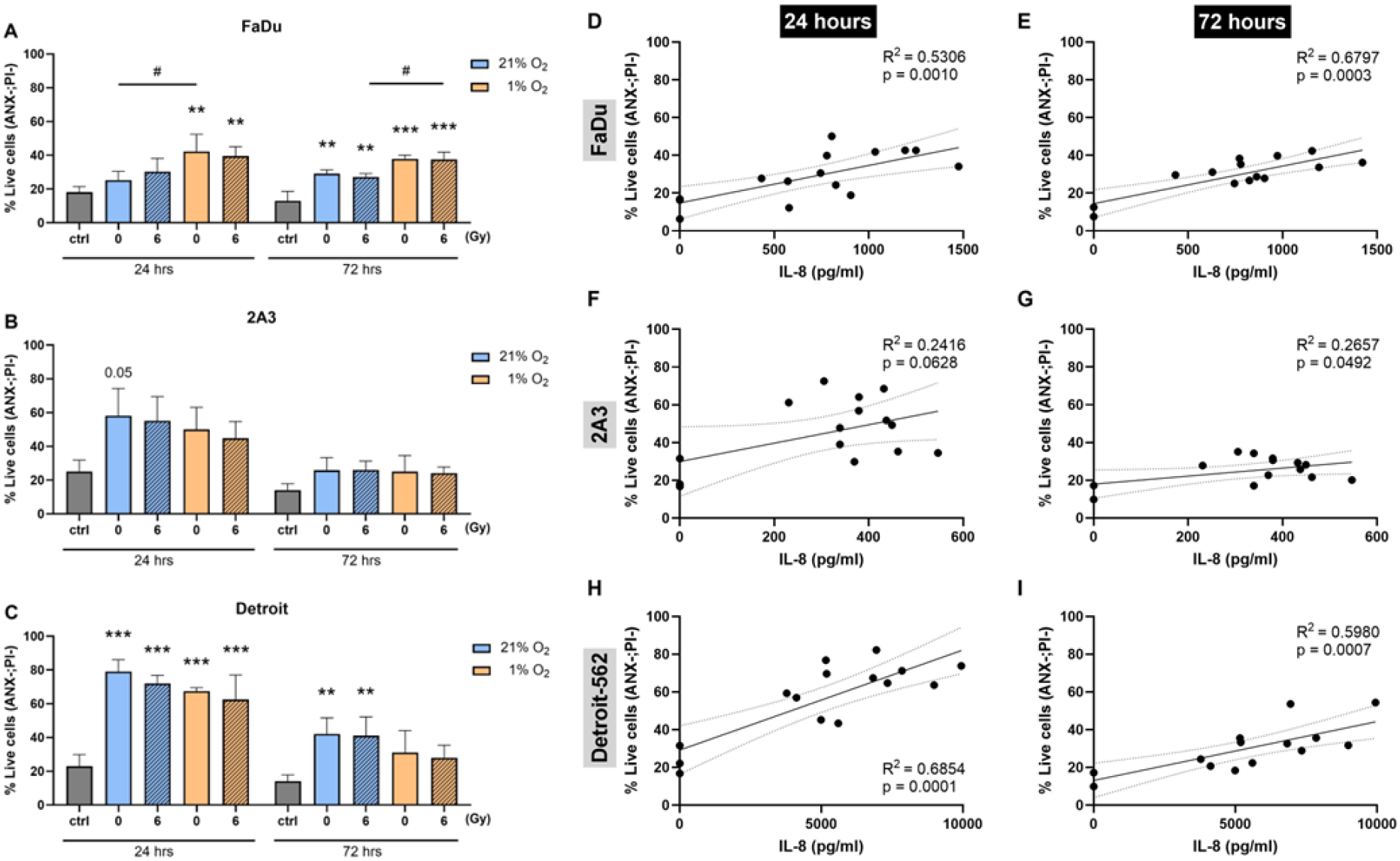
Tumor-conditioned media (TCM) from HNSCC cell lines differentially modulate human neutrophil viability in a cell line- and oxygen-dependent manner. Neutrophil viability was assessed after 24 and 72 hours of incubation with TCM generated 48 hours after irradiation (0 or 6 Gy) from FaDu (A), 2A3 (B), and Detroit-562 (C) cells processed under normoxic (21% O_2_) or hypoxic (1% O_2_) conditions. Viability was quantified by flow cytometry using Annexin V/propidium iodide (PI) staining and is reported as the percentage of double-negative (Annexin V⁻/PI⁻) neutrophils. Data are presented as mean ± SD. Statistical analysis was performed separately for each time point using one-way ANOVA with Tukey’s multiple comparisons test versus RPMI control (**p < 0.01; ***p < 0.001) or between compared oxygen conditions (#p < 0.05); n = 4. (D–I) IL-8 concentrations (pg/ml) measured in the same TCM samples were correlated with neutrophil viability using simple linear regression. Solid lines indicate the best-fit linear regression, with dashed lines showing the 95% confidence interval. Coefficients of determination (R²) and corresponding p-values are shown in each panel. Each data point represents one individual TCM sample. Data were obtained from three independent experiments with five distinct TCM samples per experiment (total n = 14–16 per cell line and time point).

At 72 hours, neutrophil viability remained significantly elevated following incubation with FaDu- or Detroit-562-derived TCM (Figure 1A,C). Hypoxia-conditioned FaDu-derived TCM further prolonged neutrophil lifespan compared with normoxic TCM, although this effect was observed only in the 6 Gy samples (Figure 1A). In contrast, normoxic Detroit-562-derived TCM increased neutrophil viability more strongly than hypoxic TCM (Figure 1C).

The viability of neutrophils incubated with 2A3- and Detroit-562-derived TCM for 72 hours (Figure 1B,C) decreased relative to the 24-hour time point, indicating that TCM delays—but does not fully prevent— neutrophil apoptosis during prolonged exposure in a cell line-dependent manner.

The neutrophil-supportive activity of TCM was preserved following tumor cell irradiation, as media generated from irradiated and non-irradiated cells induced comparable neutrophil viability (Figure 1A–C). In contrast, oxygen availability emerged as a critical determinant of TCM activity, with pronounced cell line–specific differences.

As shown in Figure 1D–I, neutrophil viability positively correlated with IL-8 concentrations in TCM derived from FaDu and Detroit-562 cells. This association was observed at both 24 and 72 hours of incubation (Figure 1D,E,H,I). In contrast, no significant correlation between IL-8 concentration and neutrophil viability was detected for TCM derived from 2A3 cells at the 24-hour time point (Figure 1F), with only borderline (p=0.0492) significance observed at 72 hours (Figure 1F). These data support a cell line–dependent relationship between tumor-derived IL-8 and neutrophil viability.

### Oxygen-dependent, radiation-independent priming of neutrophil ROS production by TCM

Because TCM alone did not induce spontaneous ROS production in neutrophils (data not shown), we assessed whether TCM primes neutrophils for ROS production in response to a commensal oral bacterium. Neutrophil ROS production in response to *F. nucleatum* stimulation was modulated by TCM in a cell line- and oxygen condition-dependent manner. As shown in Figure 2A, TCM derived from hypoxic FaDu cells significantly increased ROS production following *F. nucleatum* stimulation compared with control. Moreover, TCM from irradiated FaDu cells under hypoxic conditions caused significantly higher ROS production relative to irradiated FaDu cells cultured under ambient air (Figure 2A).

**Figure 2.**
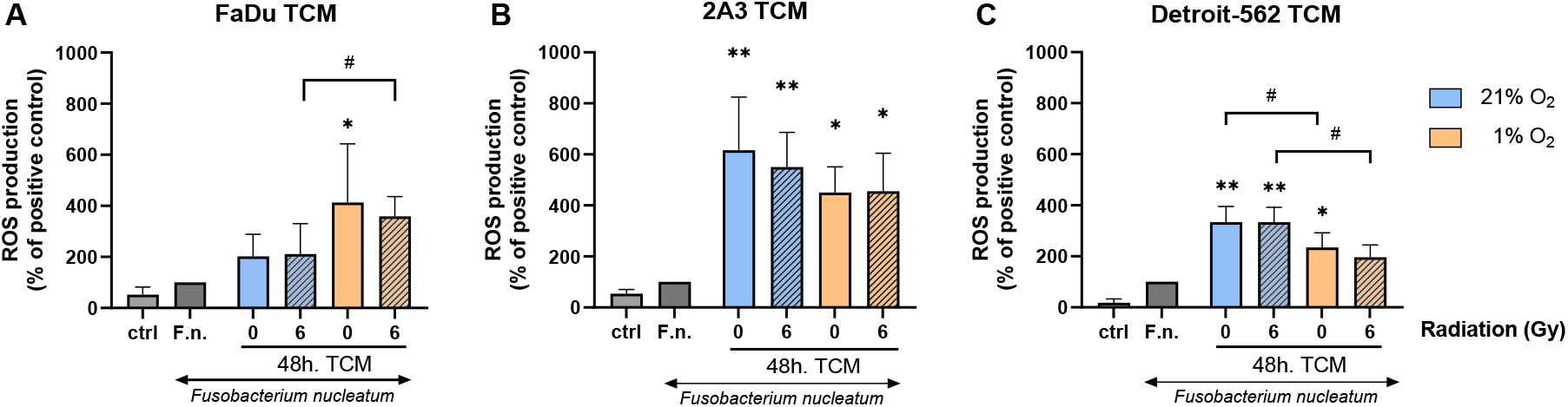
Tumor-conditioned media (TCM) from HNSCC cell lines differentially prime neutrophil ROS production. Reactive oxygen species (ROS) production was quantified by flow cytometry using DHR123. Neutrophils were incubated for 24 hours with TCM derived from (A) FaDu, (B) 2A3, or (C) Detroit-562 cells 48 hours after irradiation under 21 or 1% oxygen, and ROS production was measured for 30 minutes following stimulation with *F. nucleatum*. Results are expressed as a percentage of the stimulated control. Data are shown as mean ± SD. Statistical analysis was performed using one-way ANOVA with Dunnett’s multiple tests compared to the stimulated control (*p < 0.05; **p < 0.01) and unpaired t-tests for pairwise comparisons between indicated conditions (#p < 0.05); n = 3–4.

In contrast, TCM derived from 2A3 (Figure 2B) and Detroit-562 (Figure 2C) cells cultured under ambient oxygen conditions enhanced stimulated ROS production in neutrophils. For 2A3, relatively comparable increases in ROS generation were observed across all four TCM conditions, resulting in the highest level of ROS induction among the tested HNSCC cell lines (Figure 2B). For Detroit-562, TCM generated under ambient oxygen conditions induced significantly higher ROS production compared with hypoxia TCM (Figure 2C).

Notably, TCM derived from irradiated tumor cells did not differ from non-irradiated TCM in their ability to prime neutrophil ROS production (Figure 2A–C). Overall, these data suggest that tumor-derived factors regulate neutrophil ROS production in a cell line– and oxygen-dependent manner, with minimal contribution from irradiation.

Because FaDu-derived TCM displayed the most consistent and reproducible oxygen-dependent effects on neutrophil viability and ROS production, this cell line was selected for a more detailed evaluation, including phenotypic, migratory, and transcriptomic analyses.

### Hypoxia-conditioned FaDu-derived TCM promotes an immunomodulatory neutrophil phenotype

We next examined whether exposure to FaDu-derived TCM alters the immunoregulatory phenotype of neutrophils. Notably, classical neutrophil activation markers, such as CD11b and CD62L, showed no significant changes in surface expression, respectively, in response to short-term incubation with FaDu-derived TCM, regardless of the tested conditions (Supplementary file 1, Figure S1A, B).

We therefore analyzed the surface expression of the immunoregulatory molecules PD-L1, CD170, and TRAIL-R2 and -R3. As shown in Figure 3A, PD-L1 expression—which contributes to immune suppression— was not increased in neutrophils incubated for 48 hours with TCM derived under different conditioning of FaDu cells. TCM generated from hypoxic FaDu cells showed a tendency to increase neutrophil expression of CD170, independent of irradiation status (Figure 3B).

**Figure 3.**
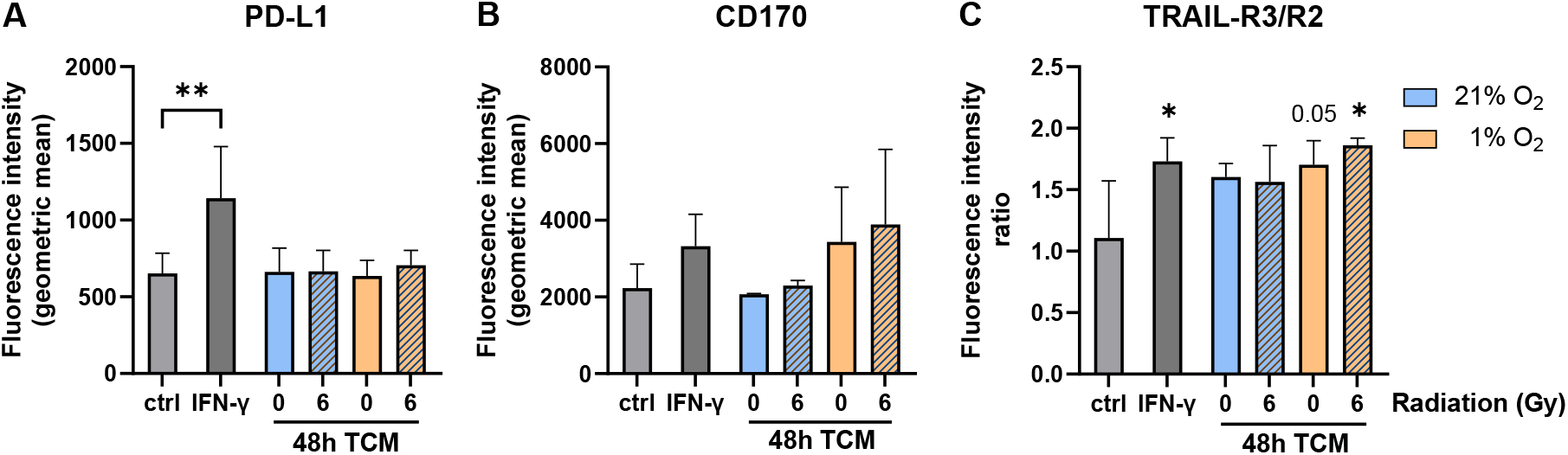
FaDu-derived tumor-conditioned media (TCM) modulate immunoregulatory surface markers on neutrophils. Surface expression of (A) PD-L1, (B) CD170, and (C) the TRAIL-R3/TRAIL-R2 ratio was quantified by flow cytometry after 48 hours of incubation with FaDu-derived TCM generated 48 hours after irradiation under 21% or 1% oxygen. Interferon-γ (IFN-γ) served as a positive control. Results are expressed as the geometric mean fluorescence intensity or its ratio ± SD. Statistical analysis was performed using one-way ANOVA followed by Dunnett’s multiple comparisons test versus the control (*p < 0.05, **p < 0.01); n = 3–4.

In contrast, expression of TRAIL-R2, a functional death receptor, showed a slight decrease following exposure to FaDu-derived TCM, except in the case of non-irradiated hypoxic TCM (Supplementary file 1, Figure S1C), whereas the decoy receptor TRAIL-R3 tended to increase regardless of the TCM conditions (Supplementary file 1, Figure S1D). Accordingly, the calculated TRAIL-R3/TRAIL-R2 ratio was significantly increased in neutrophils incubated with TCM derived from irradiated hypoxic FaDu cells and showed a near-significant increase in response to hypoxic FaDu TCM alone (Figure 3C).

Collectively, these data indicate that hypoxia-conditioned FaDu-derived TCM may promote modulation of selected neutrophil immunoregulatory surface markers.

### Migration of neutrophils in response to FaDu-derived TCM

To further characterize neutrophil functional responses, we evaluated neutrophil migration in response to FaDu-derived TCM. Neutrophils migrated comparable distances (Figure 4A) and exhibited similar migration speeds (Figure 4B) in response to the different TCM conditions, with values overall comparable to those induced by IL-8, which was included as a positive control as a well-established neutrophil chemoattractant [43]. TCM generated from non-irradiated FaDu cells cultured under hypoxic conditions induced a significantly greater migration distance compared with TCM derived from non-irradiated FaDu cells cultured under ambient oxygen conditions (Figure 4A).

**Figure 4.**
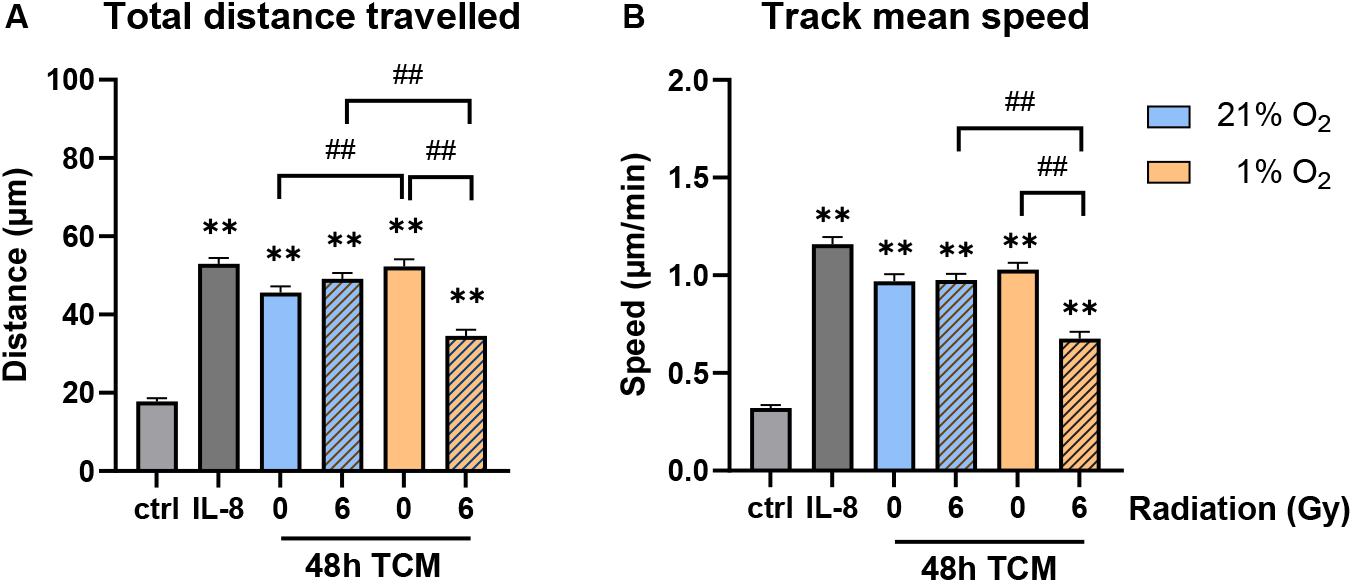
FaDu-derived tumor-conditioned media (TCM) modulate neutrophil migration. Migration parameters shown include (A) total distance traveled and (B) track mean speed. Migration was stimulated with 48-hour FaDu-derived TCM or IL-8 (2 ng/ml) used as a positive control. Data are presented as mean ± SEM. Statistical analysis was performed using one-way ANOVA by Dunnett’s multiple comparisons test versus the control (**p < 0.01) and unpaired t-tests for between-condition comparisons (##p < 0.01); n = 4-5.

In contrast, TCM derived from irradiated FaDu cells under hypoxic conditions induced significantly reduced migration distances and lower migration speeds compared with hypoxic TCM from non-irradiated FaDu cells, as well as compared with the corresponding TCM generated under ambient oxygen conditions (Figure 4A,B). Collectively, these data indicate that hypoxia-conditioned tumor-derived factors modulate neutrophil migratory behavior in a radiation-dependent manner.

### Limited normoxic response versus hypoxia-driven transcriptomic remodeling in neutrophils cultured with FaDu-derived TCM

Finally, we analyzed how FaDu-derived TCM alters the neutrophil transcriptome. Transcriptomic data used in this study are in Supplementary file 2. Neutrophils incubated with TCM derived from FaDu cells cultured under ambient oxygen conditions, either with or without irradiation, exhibited very limited transcriptional changes (Figure 5A, C), consistent with the low basal transcriptional activity of unstimulated neutrophils [44].

**Figure 5.**
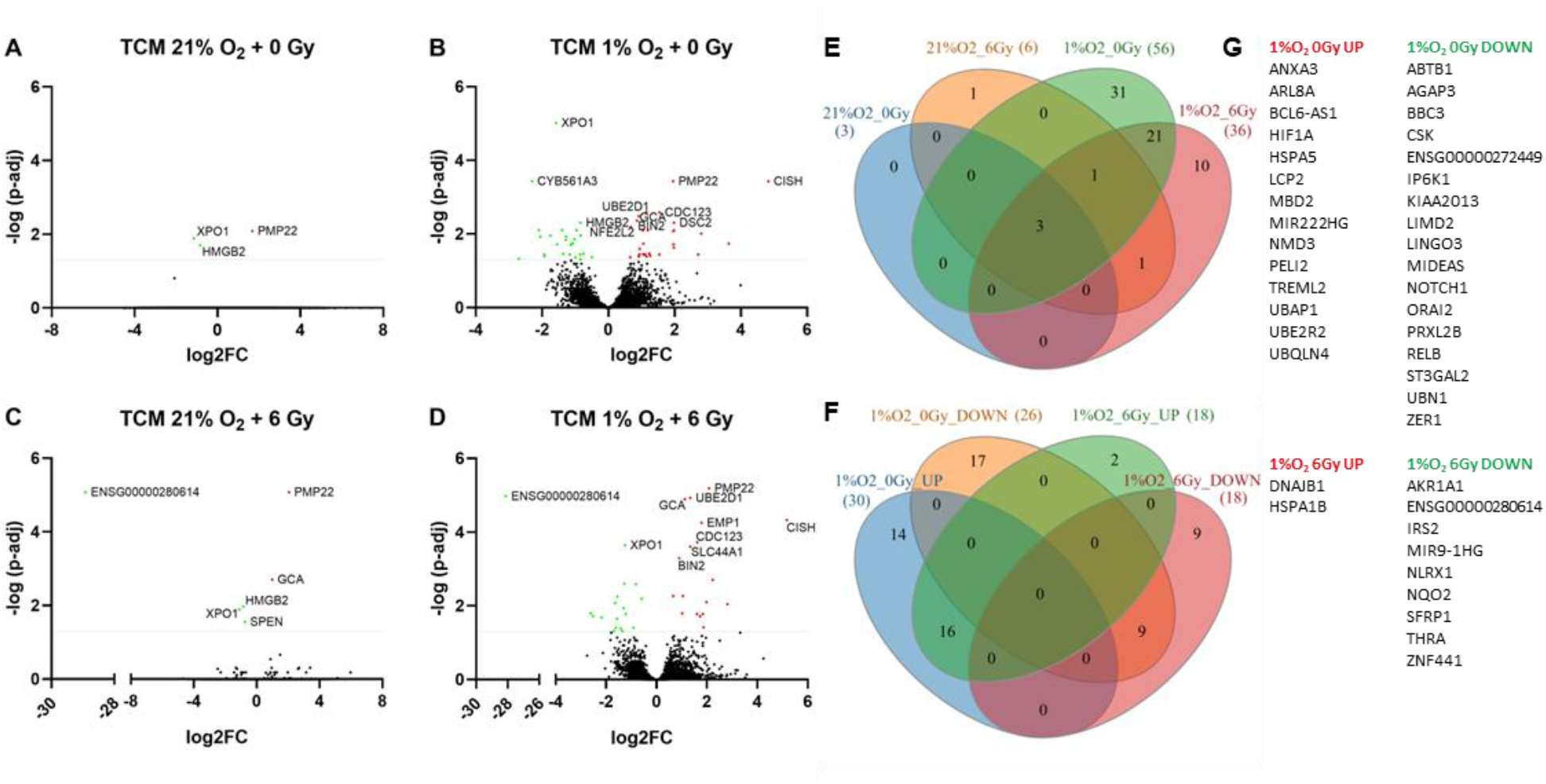
Differentially expressed genes (DEGs) in neutrophils incubated with FaDu-derived tumor-conditioned media (TCM). Neutrophils isolated from healthy donors were incubated for 4 hours with TCM generated from FaDu cells irradiated with 0 or 6 Gy under ambient (21% O_2_) or hypoxic (1% O_2_) conditions. TCM was harvested 48 hours after irradiation. TCM from three independent experiments was incubated with neutrophils from three independent donors. The three biological replicates per condition were analyzed jointly using DESeq2, which modeled the replicate-level raw read counts without prior averaging. Statistical significance was determined from the group-level comparison, and individual significance in all three replicates was not required. (A–D) DEGs in neutrophils incubated with FaDu-derived TCM under the indicated conditions, shown as volcano plots. Data are presented as log_2_ fold change versus the negative decimal logarithm of the adjusted p-value. The ten most significantly altered genes are annotated. Thresholds applied: adjusted p-value <0.05. (E) Venn diagram illustrating significantly regulated DEGs in response to radiation, oxygen tension, and their combinations. (F) Venn diagram showing significantly regulated DEGs in response to irradiation under hypoxic conditions, stratified by up- or downregulation. (G) List of DEGs differentially expressed exclusively in response to hypoxic TCM either without irradiation or following irradiation, and not overlapping with DEGs common to both conditions (DEGs in neutrophils in response to hypoxic TCM, independent of irradiation, are shown in Supplementary file 1, Table T1).

Nevertheless, mRNA levels of *XPO1* (nuclear export) and *HMGB2* (chromatin remodeling) were downregulated, whereas *PMP22* (integral membrane protein) was upregulated, with comparable fold changes and p-values across all four conditions. *GCA* (calcium-binding protein) showed a similar trend but was not detected as DEG in the normoxic non-irradiated condition. Collectively, these genes may reflect a generalized, low-level transcriptional response of neutrophils to TCM (Figure 5A–D).

In contrast, TCM derived from hypoxic FaDu cells markedly increased the number of DEGs. We identified 56 DEGs in neutrophils exposed to hypoxic TCM and 36 DEGs in response to TCM from hypoxic and irradiated FaDu cells, with 25 DEGs shared between both conditions (Figure 5E). The overlapping DEGs displayed highly similar fold-change magnitudes and directionality, with only minor differences observed for *CYB561A3, EMP1*, and *ZC3H12C*. These findings suggest that neutrophil responses to hypoxic TCM from non-irradiated and irradiated FaDu cells are predominantly qualitative rather than quantitative at the transcriptional level. Based on this observation, we focused subsequent analyses on DEGs uniquely induced by hypoxic TCM alone and by hypoxic, irradiated FaDu-derived TCM (Figure 5F). DEGs in neutrophils in response to hypoxic TCM, independent of irradiation, are shown in Supplementary file 1, Table T1.

Exposure of neutrophils to non-irradiated hypoxic FaDu-derived TCM resulted in the upregulation of 14 genes associated with hypoxia response and cellular stress (*HIF1A, HSPA5, MBD2, UBQLN4, NMD3*) or myeloid cell activation (*LCP2, PELI2, TREML2, ANXA3, ARL8A*) (Figure 5G, upper left panel). Thus, even short-term exposure to diluted hypoxic TCM activated HIF-1α pathway (Supplementary file 1, Figure S2). In contrast, incubation with irradiated hypoxia-conditioned FaDu media induced only two upregulated genes, *DNAJB1* and *HSPA1B*, both linked to generic cellular stress responses, indicating the absence of substantial additional transcriptional reprogramming compared with non-irradiated hypoxic TCM (Figure 5G, lower left panel).

Hypoxic TCM also suppressed the expression of genes involved in apoptotic and growth-associated pathways (*BBC3, ABTB1, ZER1*) (Figure 5G, upper right panel; and Supplementary file 1, Figure S3) and regulators of immune priming rather than direct activation (*CSK, RELB, NOTCH1, ORAI2*) (Figure 5G, upper right panel), consistent with the observed CD11b and CD62L surface expression profiles (Supplementary file 1, Figure S1A,B). Downregulated genes specific to irradiated hypoxic TCM were primarily related to metabolic and redox regulation (*AKR1A1, NQO2, IRS2*) and developmental signaling pathways (*SFRP1, THRA*), rather than immune-specific programs (Figure 5G, lower right panel). This further supports the absence of a distinct neutrophil activation signature in response to irradiated FaDu-derived TCM generated under hypoxic conditions.

### Translational validation of the neutrophil-derived hypoxia score in TCGA-HNSCC

To explore the clinical relevance of the experimentally derived neutrophil hypoxia-response signature, we evaluated its association with clinical outcome in publicly available TCGA-HNSCC datasets. Importantly, these datasets comprise predominantly bulk RNA-seq profiles of primary HNSCC tumors rather than neutrophil-specific transcriptomes. Consequently, this analysis should be interpreted as a translational validation of the neutrophil-derived gene signature at the tumor level, reflecting its representation within the overall tumor transcriptome and microenvironment rather than direct validation in patient neutrophils.

Median-based stratification into high- and low-hypoxia-score groups did not reveal a significant difference in overall survival by Kaplan–Meier analysis (HR = 1.15, 95% CI 0.88–1.50; log-rank p = 0.301; Figure 6A). However, when analyzed as a continuous standardized variable, the hypoxia score was independently associated with overall survival in a multivariable Cox model adjusted for age, sex, and pathological T and N stages, corresponding to an 18% increase in the hazard of death per 1-SD increase in the score (HR = 1.18, 95% CI 1.01–1.38; p = 0.039; Figure 6A,B).

**Figure 6.**
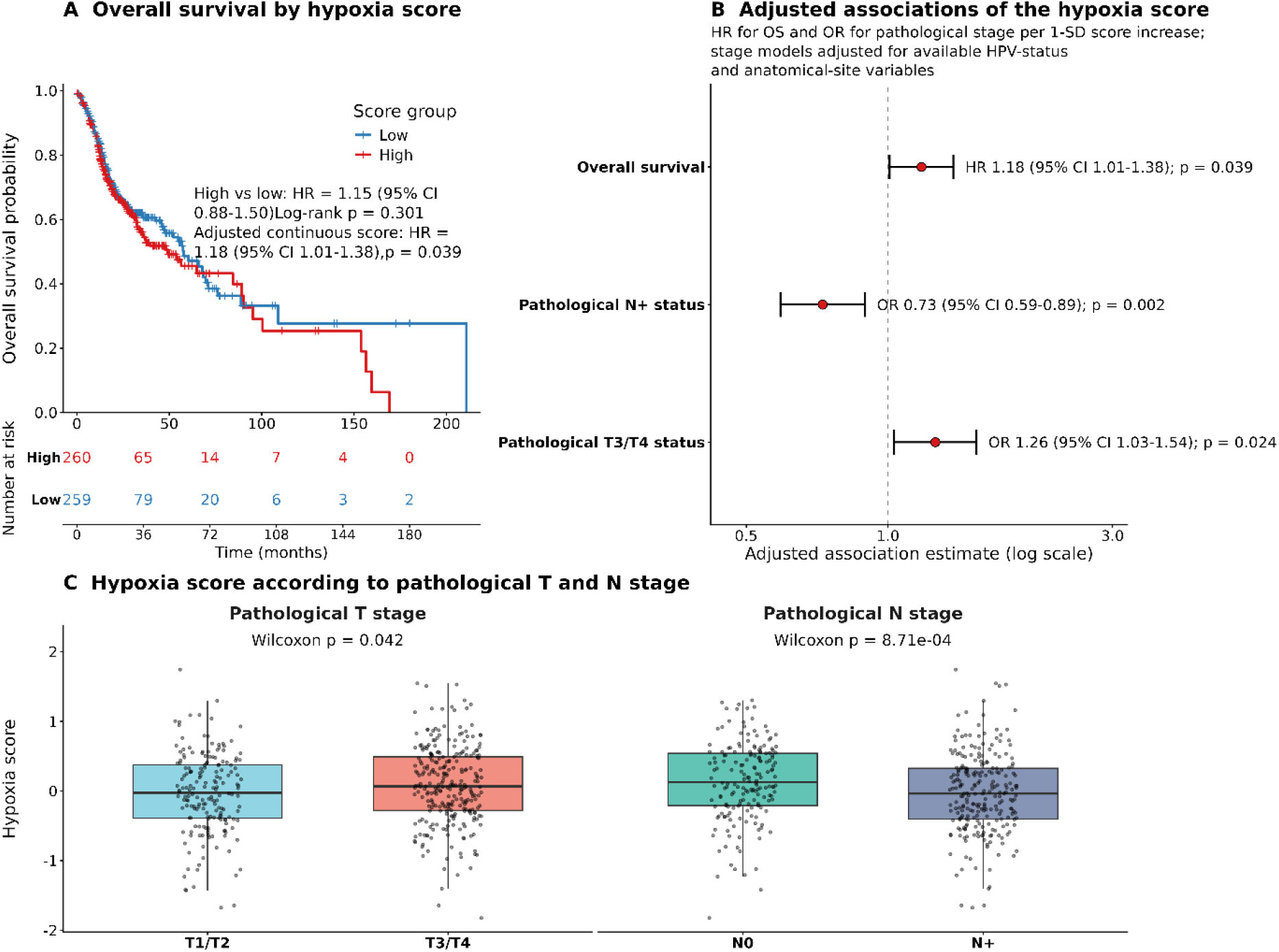
Clinical validation of the experimentally derived hypoxia score in TCGA-HNSCC. (A) Kaplan-Meier analysis of overall survival after median dichotomization of the hypoxia score. The number-at-risk table is shown below the survival curves. The panel also reports the continuous-score multivariable Cox model using the available clinical covariates specified in the analysis: HR per 1-SD increase = 1.178 (95% CI 1.008-1.377), p = 0.039. (B) Combined forest plot of the hypoxia score association with overall survival and pathological stage. The overall-survival estimate is a hazard ratio from the multivariable Cox model; pathological N+ and T3/T4 estimates are odds ratios from logistic models adjusted for available HPV-status and anatomical-site variables. (C) Distribution of the hypoxia score in pathological T1/T2 versus T3/T4 tumors and N0 versus N+ tumors. P-values were calculated using two-sided Wilcoxon rank-sum tests.

After adjustment for available HPV status and primary anatomical site, a higher hypoxia score was associated with lower odds of pathological nodal positivity (OR = 0.73, 95% CI 0.59–0.89; p = 0.002), but with higher odds of advanced pathological T stage (T3/T4; OR = 1.26, 95% CI 1.03–1.54; p = 0.024; Figure 6B). Consistently, hypoxia-score distributions differed between both T-stage and N-stage groups in unadjusted comparisons (Figure 6C). These apparently divergent associations may indicate that the signature is more closely related to local tumor growth and hypoxic features of the primary tumor rather than to lymphatic dissemination; however, this interpretation remains exploratory.

In contrast, the hypoxia-plus-irradiation score was not independently associated with overall survival. Both scores were strongly correlated, indicating that they capture substantially overlapping transcriptional programs (Supplementary file 1, Figure S4A,B).

## Discussion

Neutrophils are frequently associated with reduced overall survival across multiple cancer types; however, their prognostic impact is strongly cancer-type dependent [45]. Importantly, distinct subsets of TANs have been linked to adverse clinical outcomes [46], underscoring the need for improved stratification of neutrophil functional states and the local TME like hypoxia.

Hypoxia directly delays constitutive neutrophil apoptosis through activation of the oxygen-sensing transcription factor HIF-1α [19]. Neutrophil-specific deletion of HIF-1α selectively reduces the viability of tumor-infiltrating neutrophils [47]. Consistent with this mechanism, neutrophils preferentially accumulate in hypoxic tumor regions [48], where hypoxia promotes the persistence of pro-tumorigenic neutrophil phenotypes [46].

In HNSCC, IL-8 emerges as a hypoxia-associated prognostic biomarker [49] and hypoxia directly induces IL-8 expression in cancer cells under 1% O₂ conditions [50]. Clinically, HPV-negative HNSCC has been associated with higher IL-8 production [51]. Beyond its chemotactic function [43], IL-8 can delay spontaneous and TNF-α-mediated neutrophil apoptosis *in vitro* [52]. In our study, neutrophil viability correlated with IL-8 concentrations in FaDu- and Detroit-562-derived TCM. However, survival effects were also observed under ambient oxygen conditions, consistent with prior evidence [43]. This indicates that IL-8 and hypoxia are contributory rather than exclusive determinants of neutrophil persistence.

Together, these data suggest that oxygen tension within the TME and cancer cell type may shape IL-8– dependent neutrophil recruitment and persistence. Importantly, the effects observed in our model do not reflect a direct influence of hypoxia on neutrophils, as neutrophils were cultured under ambient conditions throughout. Instead, our data isolate the impact of soluble factors derived from hypoxia-conditioned tumor cells. Such priming may represent an early step in neutrophil reprogramming, extending the influence of tumor hypoxia beyond the local tumor niche.

ROS play a dual role in neutrophil–cancer interactions: ROS can contribute to direct tumor cell killing [47], while ROS-linked effector mechanisms can concurrently suppress cell–mediated anti-tumor immunity [53]. Increased oxidative burst was described in laryngeal carcinoma [54], while decreased ROS were reported in broader HNSCC cohorts [55], highlighting context-dependent effects. Notably, exposure of healthy donor neutrophils to TCM induces low-level spontaneous ROS production [55]. In our model, TCM alone did not elicit spontaneous ROS production. Instead, TCM primed neutrophils for stimulus-induced ROS production in a cell line- and oxygen-dependent manner. This may reflect a primed neutrophil state following exposure to tumor secretomes. Nevertheless, tumor hypoxia may play an important role in modulation of ROS production. In mouse, scRNA-seq identified enrichment of ROS-associated gene signatures in specific TAN subsets, whereas these signatures were not enriched in hypoxia/glycolysis-high TANs, suggesting that hypoxia-driven reprogramming defines a TAN state distinct from ROS-associated pro-inflammatory programs [56]. Together, these findings support the concept that tumor-secreted factors and hypoxia can shape neutrophil ROS readiness.

Neutrophils isolated from newly diagnosed, untreated HNSCC patients exhibit a CD16^high^CD62L^dim^ and CD18^high^/CD11b^high^ phenotype, indicative of chronic phenotypic reprogramming rather than acute activation [57]. Prolonged (72-hour) exposure of neutrophils to hypoxic TCM has been shown to increase the proportion of CD62L^dim^ neutrophils *in vitro* [58]. However, we did not observe changes in CD11b and CD62L following short-term incubation with TCM, regardless of oxygen or radiation conditions. This lack of response likely reflects the absence of acute activation [59], and may fundamentally differ from the sustained phenotypic alterations observed in cancer.

Consistent with previous findings, FaDu-derived TCM induced neutrophil migration *in vitro* [43]. In our model, TCM promoted neutrophil migration irrespective of oxygen levels or irradiation status of cultured FaDu cells; however, migratory capacity was significantly reduced in response to TCM derived from hypoxic, irradiated FaDu cells. In the study of Millrud *et al.* neutrophils demonstrate enhanced migratory capacity, and a higher proportion of CD16^high^CD62L^dim^ neutrophil subset correlates with improved patient survival [57]. In contrast, advanced tumor stage in HNSCC, characterized by extensive neutrophil infiltration, has been associated with poor prognosis [43], further supporting the notion that tissue-specific environmental cues drive neutrophil phenotypic adaptation.

Taken together, our results suggest that hypoxic tumor-derived soluble factors primarily support neutrophil persistence, whereas acquisition of an activated phenotype requires additional microenvironmental cues. These findings underscore the importance of prolonged or direct exposure to microenvironmental signals in shaping neutrophil activation. However, further studies are needed to identify the molecular and cellular signals governing these processes.

Next, we focused on selected surface molecules important in neutrophil-cancer interactions. We did not observe increased PD-L1 expression in neutrophils incubated with FaDu-derived TCM. Elevated PD-L1 expression in tumor cells has been associated with poor prognosis across multiple cancer types [60, 61] and increased PD-L1 expression has also been reported in neutrophils isolated from tumors and lymph nodes of patients with advanced-stage HNSCC [6]. PD-L1–positive neutrophils are known to exert immunosuppressive effects [6, 62]. Our findings therefore suggest that hypoxia-conditioned TCM alone was not sufficient to induce PD-L1 expression in neutrophils.

Similarly, CD170 expression on neutrophils has been linked to immunoregulatory and immunosuppressive phenotypes [7]. CD170 (Siglec-5 in humans, functionally related to Siglec-F in mice) plays an important role in tumor–neutrophil interactions. Engagement of tumor-associated sialic acids with Siglec-5 on neutrophils inhibits tumor cell killing [63], and Siglec-F^high^ neutrophils in tumor-bearing mice display tumor-promoting gene signatures [64]. In our model, TCM derived from hypoxic FaDu induced a modest increase in CD170 expression on neutrophils, independent of irradiation of tumor cells. Our findings suggest that tumor-derived soluble factors may contribute to neutrophil reprogramming by enhancing inhibitory receptor expression, even in the absence of direct hypoxic exposure. While increased Siglec-5 expression has been reported in other immune cell types under hypoxic conditions [65, 66], its regulation in neutrophils in response to hypoxia-conditioned tumor secretomes and/or irradiation-associated tumor signaling remains poorly defined and warrants further investigation.

In mouse, decoy TRAIL-R1^+^ neutrophils preferentially localize to hypoxic tumor regions and show pro-tumoral characteristics [21]. In humans, neutrophils predominantly express TRAIL-R2 (a functional death receptor) and TRAIL-R3 (a decoy receptor), on their surface [67]. TRAIL receptors in neutrophils may function not primarily as death receptors but rather as modulators of survival and inflammatory signaling [68]. Neutrophils typically express low levels of TRAIL-R2 and high levels of TRAIL-R3, with TRAIL-R3 being further regulated by TNF-α (negatively) and IFN-γ (positively) [69]. Thus, the balance between functional and decoy receptors is skewed toward the non-functional receptor in neutrophils [70]. The findings of Ng *et al.* indicate that hypoxia alone is not sufficient to induce decoy TRAIL-R1 expression in murine neutrophils, whereas tumor-derived signals are required for TRAIL-R1-associated neutrophil reprogramming [21]. In agreement with this, TCM induced a trend toward increased expression of the decoy receptor TRAIL-R3 and decreased expression of the functional receptor TRAIL-R2, resulting in an increased TRAIL-R3/TRAIL-R2 ratio in neutrophils exposed to hypoxic FaDu-derived TCM.

However, the role of TRAIL-R2 and -R3 in tumor-associated neutrophils *in vivo* remains to be defined. In contrast to neutrophils, targeting TRAIL-R2 in patients with HNSCC selectively reduced myeloid-derived suppressor cells, while neutrophil numbers remained unchanged [71]. Importantly, downregulation of decoy TRAIL-Rs in neutrophils from healthy donors renders these cells sensitive to TRAIL-R2 agonistic antibody [71]. Despite the limited data on TRAIL-Rs function in neutrophils, the increased ratio of the decoy TRAIL-R3 observed in our model may contribute to sustained neutrophil persistence in the hypoxic tumor niche.

Transcriptomic profiling further supported this interpretation. Even short-term exposure to hypoxia-conditioned TCM induced a transcriptional response characterized by hypoxia- and stress-associated genes. Notably, *HIF1A* and *NFE2L2*, which are components of transcriptional programs shared with tumor-associated neutrophils, were upregulated, reflecting activation of hypoxia signaling and oxidative stress pathways with links to angiogenic regulation [72]. Interestingly, neutrophils also exhibited strong induction of the negative regulator of cytokine signaling *CISH*, suggesting activation of feedback mechanisms that limit excessive inflammatory responses despite cellular activation. In parallel, the increased expression of the growth factor genes *TGFA* and *HBEGF*, along with reduced expression of the pro-apoptotic gene *BBC3*, indicate enhanced tissue-remodeling capacity and prolonged cell survival rather than a classical acute inflammatory phenotype. Collectively, these findings suggest that soluble factors released by hypoxic HNSCC cells are sufficient to induce a persistent adaptive transcriptional program in neutrophils, characterized by attenuation of inflammatory signaling while preserving functions associated with tissue remodeling and tumor promoting [73, 74].

This study has several limitations. First, our findings are based on an *in vitro* reductionist model using healthy donor neutrophils exposed to TCM rather than direct contact with tumor, stromal, and vascular components of the native microenvironment. Thus, our system captures soluble tumor-derived signals, but not the full cellular and spatial complexity of tumors. Nevertheless, this simplified but well-defined approach allows us to directly examine how tumor-derived soluble factors influence neutrophil responses, providing mechanistic insights that would be difficult to isolate in more complex systems. Second, hypoxia was modeled using a single oxygen concentration (1% O2) during tumor cell conditioning, which does not reflect the dynamic oxygen gradients present *in vivo* and over different time points. Despite this simplification, the use of a defined hypoxic condition allows for reproducible assessment of biologically relevant hypoxia-driven effects and establishes a clear baseline for future studies incorporating gradient models. Third, neutrophils were exposed to hypoxic TCM for a short-term period under standard ambient culture conditions rather than continuous matched hypoxia, so our data primarily address hypoxia-driven changes in the tumor secretome rather than the combined direct and indirect effects of hypoxia on neutrophils. However, this model is complementary to recent studies [21] and this design isolates the contribution of hypoxia-conditioned tumor secretome, thereby offering valuable insight into indirect hypoxia-mediated modulation of neutrophil function. Despite these limitations, our results provide a framework for future mechanistic studies on mediators responsible for the observed phenotypic changes in tumor-associated neutrophils.

## Conclusion

Here we showed that hypoxia-conditioned TCM drive neutrophil reprogramming, promoting prolonged survival and selective priming of ROS production in cell line-dependent manner in HNSCC. Further, hypoxic FaDu-derived TCM increased TRAIL-R3/TRAIL-R2 ratio and induced significant transcriptional changes in neutrophils. On the other hand, normoxic TCM had only minimal effects on the neutrophil transcriptome. Further, gamma irradiation of tumor cells had only limited additional effects on neutrophil transcriptional activity (Scheme 1). Although our study is limited by its reductionist *in vitro* design using healthy donor neutrophils and soluble factors only, this mechanistic model enabled isolation of hypoxia-conditioned tumor secretome effects. Translationally, these findings provide a rationale for further exploring hypoxia-driven tumor–neutrophil crosstalk and associated neutrophil states in hypoxic HNSCC, particularly in the context of radiotherapy.

**Scheme 1.**
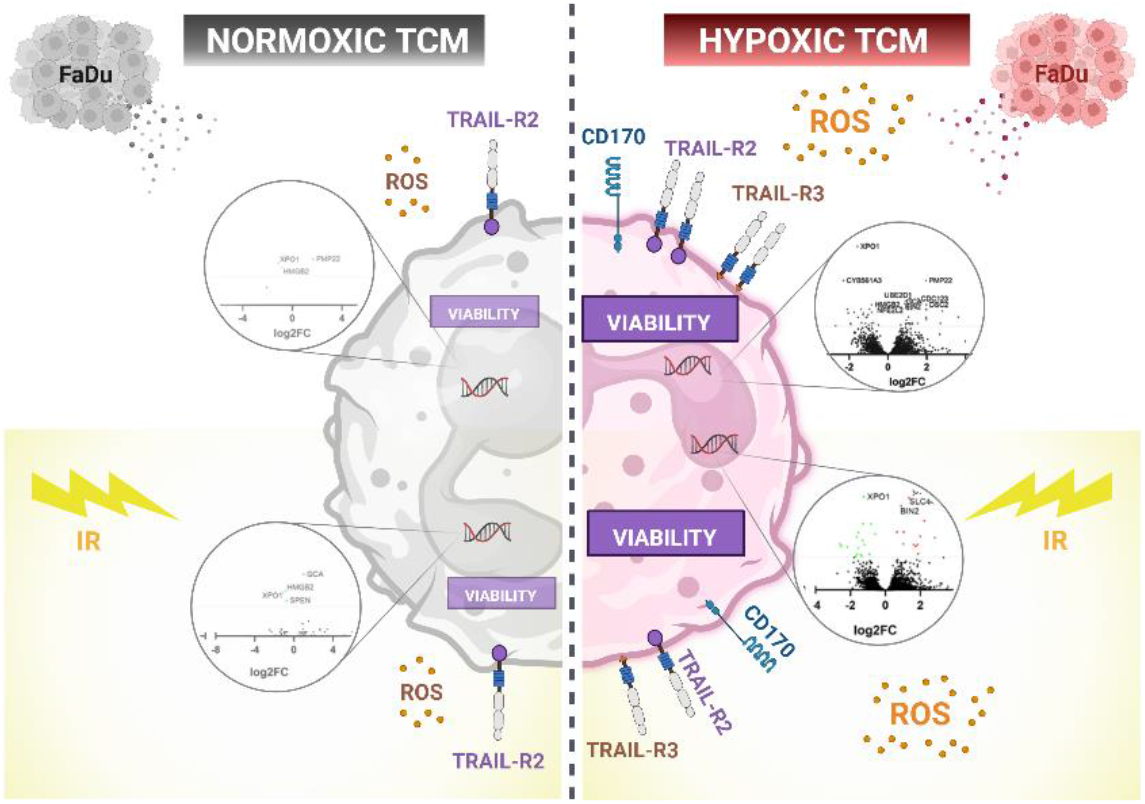
Proposed model of hypoxia-driven neutrophil reprogramming by HNSCC-derived conditioned media. Hypoxia-conditioned tumor secretomes promote neutrophil survival, selective ROS priming, phenotypic changes characterized by increased TRAIL-R3/TRAIL-R2 ratio and increased transcriptional activity, whereas gamma irradiation exerts only limited additional effects. Created in BioRender. Pereckova, J. (2026) https://BioRender.com/1ufep1l.

## Acknowledgements

This publication is based upon work from the COST Action Converting molecular profiles of myeloid cells into biomarkers for inflammation and cancer (Mye-InfoBank), CA20117 (R.H., T.P. members), supported by COST (European Cooperation in Science and Technology). Scheme 1 was created with BioRender (Pereckova, J. (2026) https://BioRender.com/1ufep1l).

## Author Contributions

Conceptualization: J.P., R.H., O.V., T.P.; Designed and performed experiments: J.P., F.Z.K., S.V., O.V., T.K., R.H., T.P.; Analyzed transcriptomic data: F.Z.K.; Writing - original draft preparation: J.P., F.Z.K., R.H., O.V., T.P.; Writing – review and editing: J.P., F.Z.K., S.V., O.V., T.K., R.H., T.P.; Funding acquisition: F.Z.K., R.H., T.P. All authors approved the final manuscript.

## Use of Generative AI

A large language model (Microsoft Copilot, GPT-5 chat model) was used for language editing and stylistic refinement. The authors confirm the originality and accuracy of the content and take full responsibility for it.

## Data Availability Statement

The NGS datasets generated and analyzed in this study will be made publicly available upon acceptance in the NCBI public database on the URL: http://www.ncbi.nlm.nih.gov/bioproject/1477777.

Annotated source datasets supporting the results of this study and underlying the figures, supplementary materials, and statistical analyses presented in this study will be publicly available upon acceptance in the public repository on the URL: https://doi.org/10.57680/asep.0651384.

## Supplementary Information

Supplementary file 1: SI-Figures (PDF format)

Contains Figures S1–S4 showing expression of surface molecules CD11b and CD62L, KEGG pathway maps of HIF-1α and apoptosis signaling pathways, and overall survival by hypoxia + irradiation score and correlation between hypoxia versus hypoxia + irradiation score. The SI also contains table T1 with DEGs common for neutrophils cultured with TCM derived from non-irradiated and irradiated FaDu under hypoxia.

Supplementary file 2: Results_LFC_Pval_DESeq2 (XLSX format)

Contains differentially expressed genes analyses used in this study.

---

**SI Figure S1.**
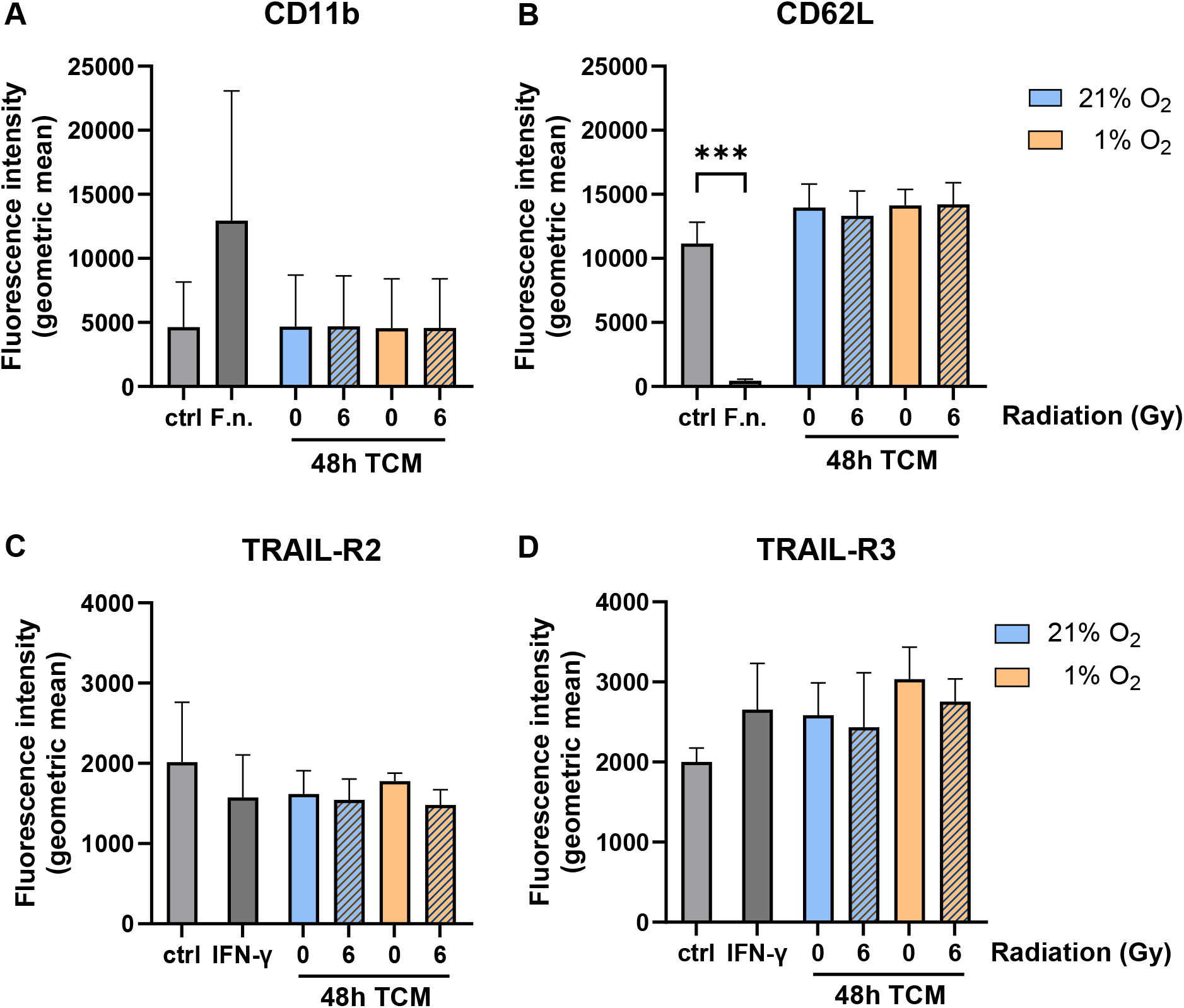
Effects of FaDu-derived tumor-conditioned media (TCM) on expression of neutrophil activation markers and TRAIL-receptors. Surface expression of (A) CD11b and (B) CD62L on neutrophils was quantified by flow cytometry after 2 hours of incubation with FaDu-derived TCM generated 48 hours after irradiation under normoxic (21% O₂) or hypoxic (1% O₂) conditions. *Fusobacterium nucleatum* (F.n.) served as a positive control. Results are presented as geometric mean fluorescence intensity ± SD. Surface expression of (C) TRAIL-R2 and (D) TRAIL-R3 on neutrophils was quantified by flow cytometry after 48 hours of incubation with FaDu-derived TCM generated 48 hours after irradiation under 21 or 1% oxygen. Interferon gamma (IFN-γ) served as a positive control. Statistical analysis was performed using one-way ANOVA with Dunnett’s multiple comparisons test versus the control (***p < 0.001); n = 3.

**SI Table T1.**
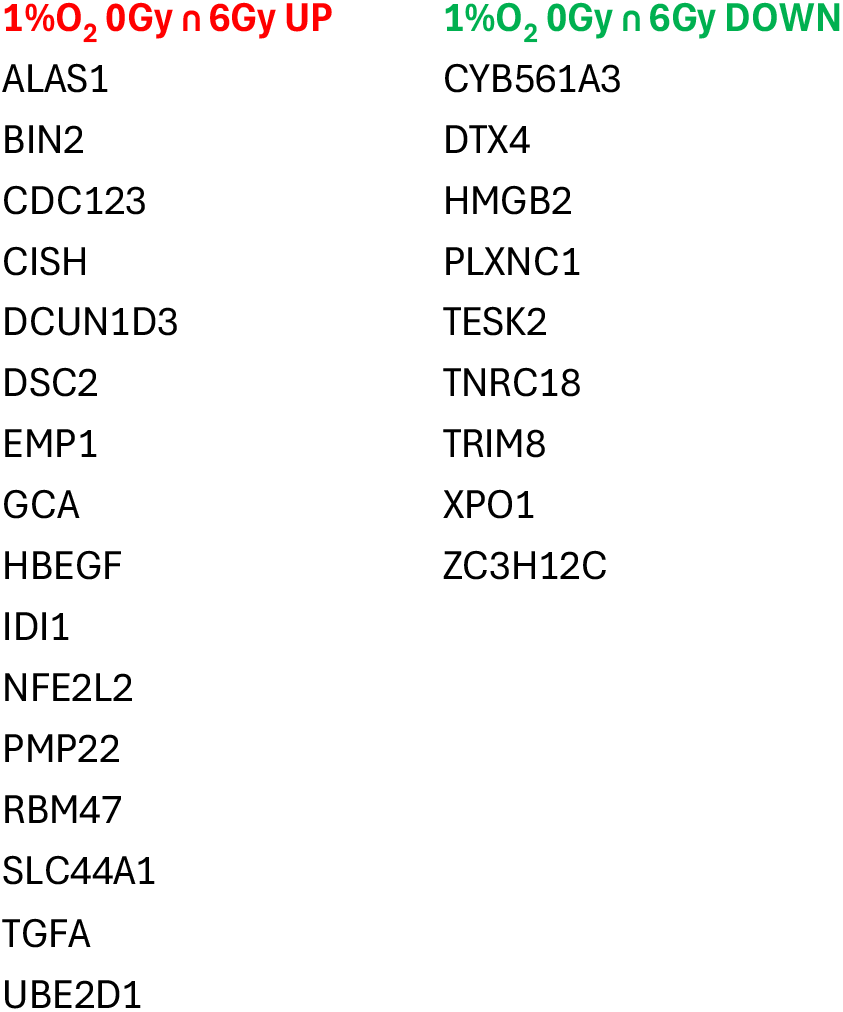
Differentially expressed genes (DEGs) in neutrophils in response to hypoxic tumor-conditioned medium (TCM), independent of irradiation. Neutrophils were incubated for 4 hours with TCM generated 48 hours after irradiation (0 or 6 Gy) from FaDu cells cultured under normoxic (21% O₂) or hypoxic (1% O₂) conditions. The table lists genes that were identified as significantly differentially expressed under hypoxic conditions in both irradiation settings (0 Gy ∩ 6 Gy), including both up- and downregulated genes. The analysis is from three independent biological replicates (n=3) and met the inclusion criterion of an adjusted p-value (padj) < 0.05.

**SI Figure S2.**
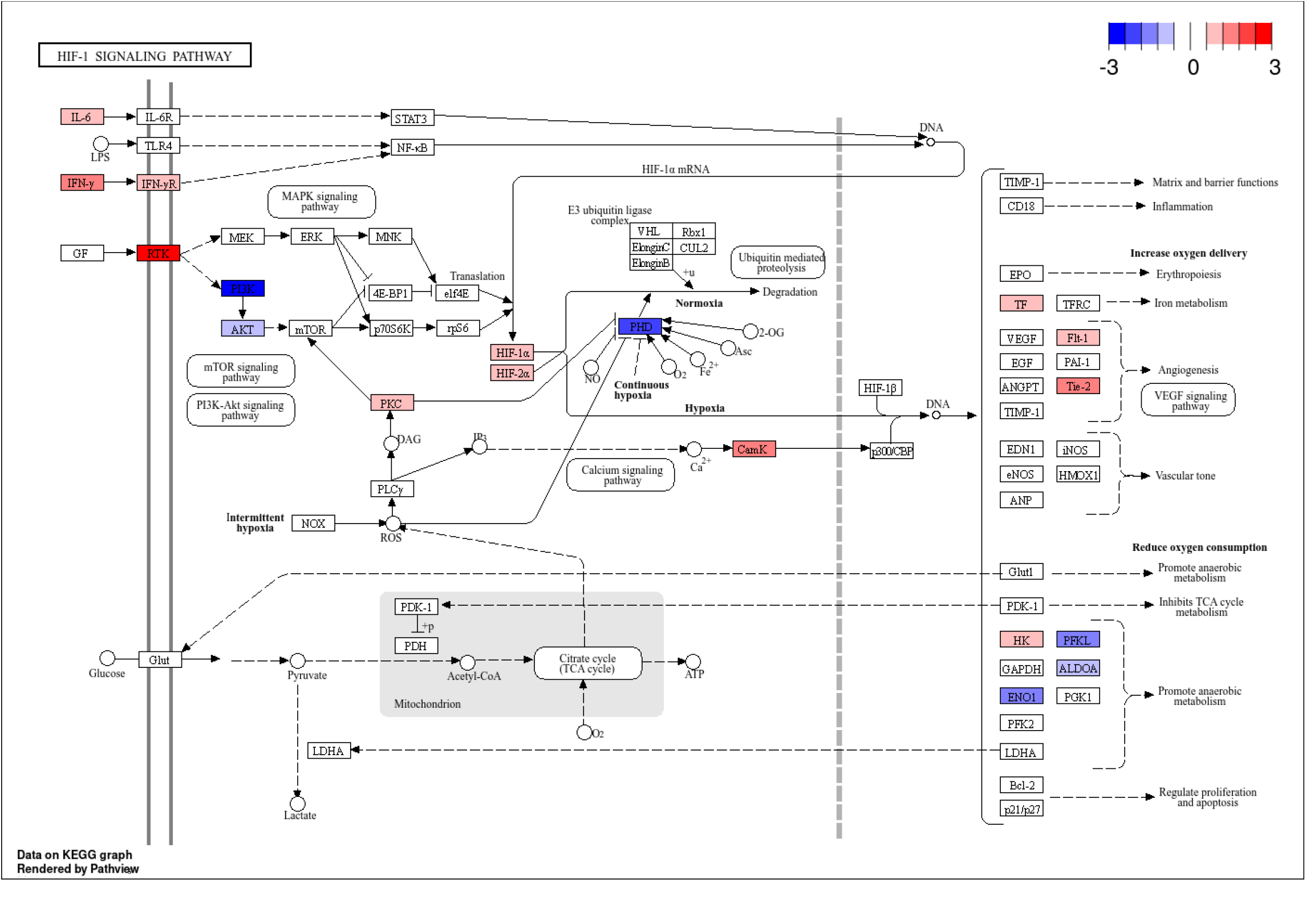
HIF-1α signaling pathway. Differentially expressed genes from neutrophils incubated with hypoxic FaDu-derived TCM. Genes are color-coded according to log2 fold change: red indicates upregulation, blue indicates downregulation, and white denotes no significant change. Threshold applied: adjusted p-value < 0.05. The map was generated using the KEGG database (DOI: 10.1093/nar/gkac963) and visualized with iDep (DOI: 10.1186/s12859-018-2486-6).

**SI Figure S3.**
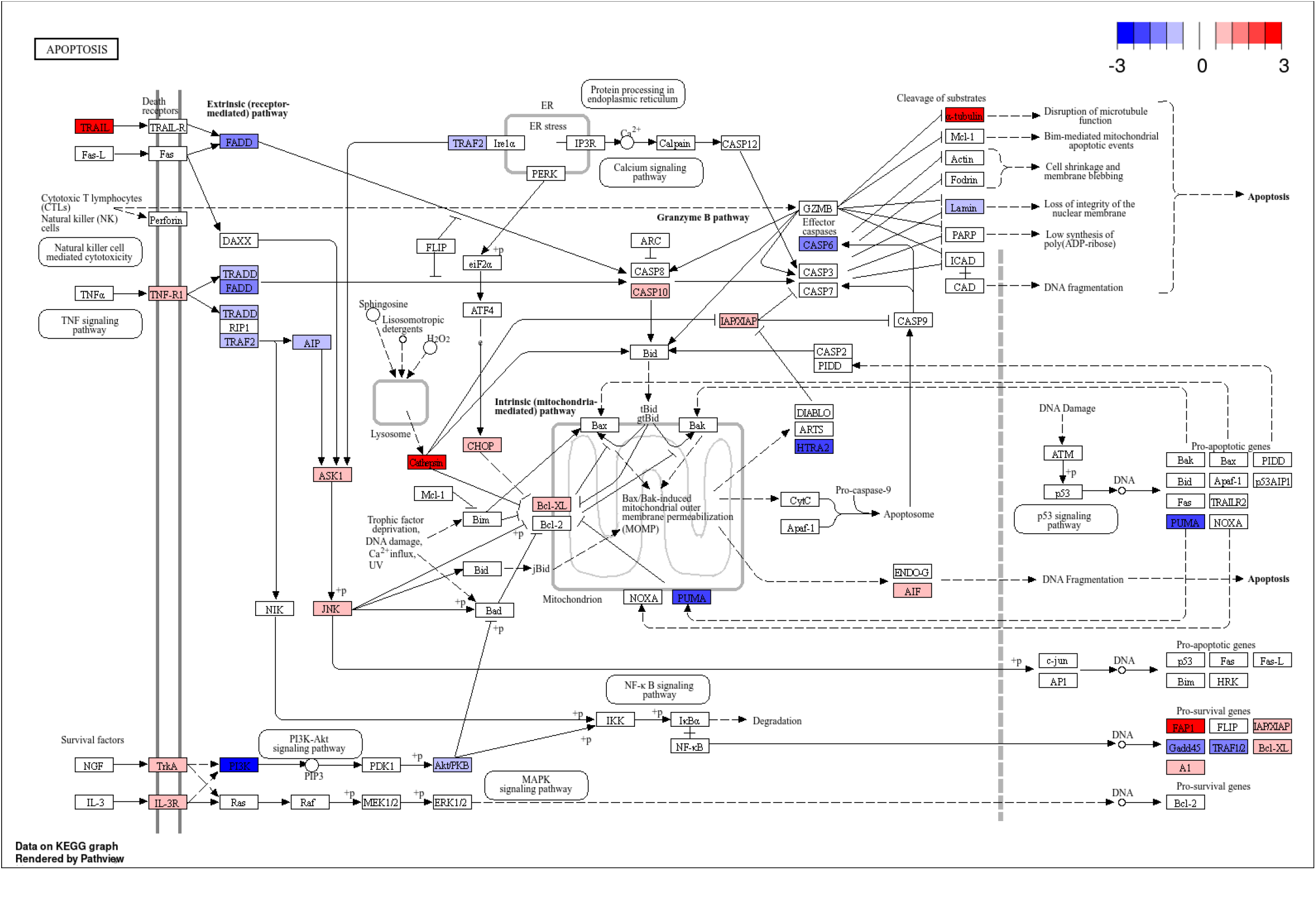
Apoptosis signaling pathway. Differentially expressed genes from neutrophils incubated with hypoxic FaDu-derived TCM. Genes are color-coded according to log2 fold change: red indicates upregulation, blue indicates downregulation, and white denotes no significant change. Threshold applied: adjusted p-value < 0.05. The map was generated using the KEGG database (DOI: 10.1093/nar/gkac963) and visualized with iDep (DOI: 10.1186/s12859-018-2486-6).

**SI Figure S4.**
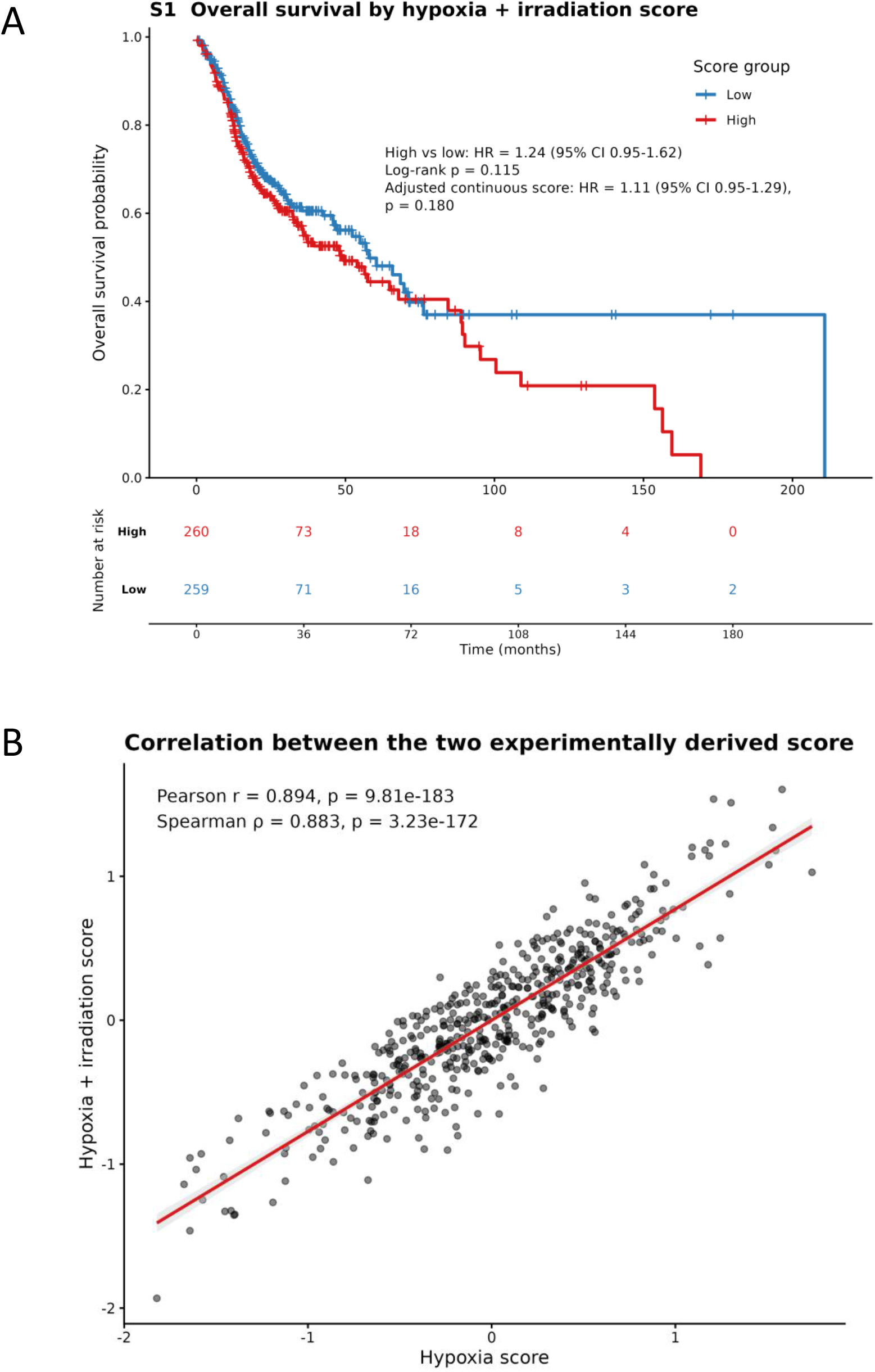
Overall survival according to the hypoxia-plus-irradiation score and correlation between the hypoxia and hypoxia-plus-irradiation scores in the TCGA-HNSCC cohort. (A) Patients were stratified into low- and high-score groups using the median hypoxia-plus-irradiation score. Overall survival was evaluated using Kaplan–Meier analysis, and differences between groups were assessed using the log-rank test. The plot shows the unadjusted hazard ratio for the high- versus low-score group and the hazard ratio for the continuous standardized score derived from a multivariable Cox proportional-hazards model adjusted for age, sex, and pathological T and N stages. The number of patients at risk is shown below the survival curves. (B) Correlation between the hypoxia and hypoxia-plus-irradiation scores in the TCGA-HNSCC cohort. Each point represents one tumor sample. The line shows the fitted linear regression with the corresponding 95% confidence interval. Pearson’s correlation coefficient, Spearman’s rank correlation coefficient, and their associated p-values are displayed in the plot.

