## Supplementary File 2 for "Hypoxia-conditioned HNSCC cell line secretomes drive phenotypic, functional, and transcriptional reprogramming of human neutrophils"

| symbol | ensembl_ID | baseMean |
|---|---|---|
| ENSG00000112096 | ENSG00000112096 | 86853,13305 |
| SRGN | ENSG00000122862 | 99556,40465 |
| CXCL8 | ENSG00000169429 | 145579,2926 |
| B2M ENSG00000166710 | ENSG00000166710 | 90819,62332 |
| NAMPT | ENSG00000105835 | 91084,10181 |
| GLUL | ENSG00000135821 | 32550,50411 |
| IFIT2 | ENSG00000119922 | 19466,33976 |
| ACTB | ENSG00000075624 | 31822,72882 |
| SAT1 | ENSG00000130066 | 44771,38141 |
| MXD1 | ENSG00000059728 | 39064,25135 |
| IVNS1ABP | ENSG00000116679 | 41240,10086 |
| MALAT1 | ENSG00000251562 | 31091,10775 |
| MT-RNR2 | ENSG00000210082 | 31582,29955 |
| LAPTM5 | ENSG00000162511 | 29872,10573 |
| LCP1 | ENSG00000136167 | 27910,42074 |
| HLA-B ENSG00000234745 | ENSG00000234745 | 21834,37167 |
| TMSB4X | ENSG00000205542 | 22780,76147 |
| AQP9 | ENSG00000103569 | 28209,86581 |
| FTL | ENSG00000087086 | 19863,32751 |
| LITAF | ENSG00000189067 | 22657,65197 |
| S100A9 | ENSG00000163220 | 21084,80119 |
| IFIT3 | ENSG00000119917 | 11723,27918 |
| TXNIP | ENSG00000265972 | 16982,74548 |
| MYH9 | ENSG00000100345 | 17317,72845 |
| CXCR4 | ENSG00000121966 | 18104,62877 |
| CSF3R | ENSG00000119535 | 20216,20409 |
| FTH1 | ENSG00000167996 | 14210,65667 |
| TNFRSF1B | ENSG00000028137 | 12725,57212 |
| H3-3B | ENSG00000132475 | 16894,65662 |
| HIF1A | ENSG00000100644 | 23299,24054 |
| HLA-E ENSG00000204592 | ENSG00000204592 | 15202,31836 |
| SDCBP | ENSG00000137575 | 16547,2953 |
| HLA-C ENSG00000204525 | ENSG00000204525 | 14324,06883 |
| ITM2B | ENSG00000136156 | 17257,86758 |
| NFKBIA | ENSG00000100906 | 13948,38311 |
| PTPRC ENSG00000081237 | ENSG00000081237 | 17475,44817 |
| TNIP1 | ENSG00000145901 | 11320,68478 |
| TNFAIP2 | ENSG00000185215 | 10689,17886 |
| EEF1A1 | ENSG00000156508 | 10488,14368 |
| MCL1 | ENSG00000143384 | 16589,36039 |
| EHD1 | ENSG00000110047 | 12191,70589 |
| ACSL1 | ENSG00000151726 | 14586,89755 |
| IFIT1 | ENSG00000185745 | 7744,134194 |
| TNFAIP3 | ENSG00000118503 | 12164,43383 |
| MARCKS | ENSG00000277443 | 13087,56178 |
| C5AR1 | ENSG00000197405 | 12774,54242 |
| IL1RN | ENSG00000136689 | 13474,66351 |
| RSAD2 | ENSG00000134321 | 6037,338821 |
| DENND5A | ENSG00000184014 | 10871,08935 |
| H3-3A | ENSG00000163041 | 12027,02469 |
| PLEK | ENSG00000115956 | 14622,48306 |
| LCP2 | ENSG00000043462 | 15762,2166 |
| AZIN1 | ENSG00000155096 | 9575,0399 |
| FCGR3B | ENSG00000162747 | 10447,48276 |

| | | |
|---|---|---|
| PSAP | ENSG00000197746 | 8435,899205 |
| S100A8 | ENSG00000143546 | 10905,78293 |
| FPR1 | ENSG00000171051 | 9740,908417 |
| ITGAX | ENSG00000140678 | 13964,80632 |
| TAGLN2 | ENSG00000158710 | 9065,625076 |
| SLC25A37 | ENSG00000147454 | 10930,46756 |
| ARRDC3 | ENSG00000113369 | 9776,635017 |
| TREM1 | ENSG00000124731 | 11782,57038 |
| PLAUR | ENSG00000011422 | 11878,26099 |
| SLC2A3 | ENSG00000059804 | 12709,89085 |
| HLA-A ENSG00000206503 | ENSG00000206503 | 7465,688149 |
| LYN | ENSG00000254087 | 10588,90614 |
| RASSF5 | ENSG00000266094 | 9468,871511 |
| ALOX5AP | ENSG00000132965 | 11304,05377 |
| FLNA | ENSG00000196924 | 9812,997403 |
| YWHAZ | ENSG00000164924 | 10676,4368 |
| SERPINA1 ENSG00000197249 | ENSG00000197249 | 9053,068529 |
| IFITM2 | ENSG00000185201 | 9638,013937 |
| ECE1 | ENSG00000117298 | 7451,50622 |
| CTSS | ENSG00000163131 | 10151,3566 |
| VASP | ENSG00000125753 | 7933,70491 |
| ANPEP | ENSG00000166825 | 9537,559802 |
| DDX3X | ENSG00000215301 | 9891,193595 |
| BCL2A1 | ENSG00000140379 | 11306,09745 |
| SLC43A2 ENSG00000167703 | ENSG00000167703 | 6392,341964 |
| NCF2 | ENSG00000116701 | 7996,003934 |
| SNN | ENSG00000184602 | 6698,410858 |
| SMCHD1 | ENSG00000101596 | 10239,90516 |
| TMEM154 | ENSG00000170006 | 7000,396595 |
| SERPINB1 | ENSG00000021355 | 9942,25354 |
| BTG2 | ENSG00000159388 | 7172,864912 |
| ADAM8 | ENSG00000151651 | 6499,466241 |
| NINJ1 | ENSG00000131669 | 7208,558802 |
| KDM6B | ENSG00000132510 | 7994,730419 |
| PABPC1 | ENSG00000070756 | 7350,355081 |
| LPCAT1 ENSG00000153395 | ENSG00000153395 | 7822,243783 |
| CD53 | ENSG00000143119 | 8642,423317 |
| PLAU | ENSG00000122861 | 8068,017861 |
| RASSF2 | ENSG00000101265 | 6362,80959 |
| TPT1 | ENSG00000133112 | 8248,851518 |
| FOSL2 | ENSG00000075426 | 7676,639881 |
| ZFP36L1 | ENSG00000185650 | 7480,472595 |
| MX1 | ENSG00000157601 | 5126,48891 |
| G0S2 | ENSG00000123689 | 9895,519426 |
| NFKBIZ | ENSG00000144802 | 8417,542967 |
| PIK3R5 | ENSG00000141506 | 7366,687586 |
| CD55 | ENSG00000196352 | 8677,571814 |
| ADAR | ENSG00000160710 | 6961,8037 |
| GPCPD1 | ENSG00000125772 | 8816,92911 |
| FGR | ENSG00000000938 | 6242,371811 |
| IFNGR2 ENSG00000159128 | ENSG00000159128 | 6449,233461 |
| TPM4 | ENSG00000167460 | 8905,579992 |
| SKIL | ENSG00000136603 | 8480,361428 |
| ACTG1 | ENSG00000184009 | 6396,45447 |
| SEC14L1 | ENSG00000129657 | 7408,427646 |

| | | |
|---|---|---|
| PDE4B | ENSG00000184588 | 6973,645851 |
| SORL1 | ENSG00000137642 | 6031,985104 |
| RNF19B | ENSG00000116514 | 5971,127437 |
| PTAFR | ENSG00000169403 | 5204,13791 |
| CD44 | ENSG00000026508 | 5774,996585 |
| NFKB2 | ENSG00000077150 | 5147,127165 |
| TNFAIP6 | ENSG00000123610 | 5596,275465 |
| THBS1 | ENSG00000137801 | 5084,460937 |
| GNG2 | ENSG00000186469 | 8122,213612 |
| MYO1F | ENSG00000142347 | 5125,229751 |
| RANBP2 | ENSG00000153201 | 6578,469738 |
| VNN2 | ENSG00000112303 | 5998,496091 |
| NEAT1 | ENSG00000245532 | 7923,403003 |
| BASP1 | ENSG00000176788 | 6432,408238 |
| APLP2 | ENSG00000084234 | 7297,90531 |
| PLXNC1 | ENSG00000136040 | 4638,923341 |
| SLC11A1 | ENSG00000018280 | 7447,719222 |
| MSN | ENSG00000147065 | 5365,690531 |
| SELL | ENSG00000188404 | 7313,137761 |
| PNRC1 | ENSG00000146278 | 6833,999198 |
| RESF1 | ENSG00000174718 | 4851,590001 |
| N4BP1 | ENSG00000102921 | 7010,824108 |
| TLR4 | ENSG00000136869 | 6448,108219 |
| RNF213 | ENSG00000173821 | 4158,99399 |
| PREX1 | ENSG00000124126 | 5410,022986 |
| LIMK2 | ENSG00000182541 | 5436,479125 |
| ASAH1 | ENSG00000104763 | 5886,186925 |
| UBC | ENSG00000150991 | 4946,849136 |
| FMNL1 | ENSG00000184922 | 5861,604146 |
| GBP2 | ENSG00000162645 | 6126,931609 |
| IQGAP1 | ENSG00000140575 | 6270,02815 |
| ADGRG3 | ENSG00000182885 | 6088,519868 |
| PELI1 | ENSG00000197329 | 7189,210843 |
| PPIF | ENSG00000108179 | 5473,293131 |
| IFNGR1 | ENSG00000027697 | 7398,426909 |
| CMTM6 | ENSG00000091317 | 5607,539012 |
| MT-RNR1 | ENSG00000211459 | 5677,762544 |
| CYRIB | ENSG00000153310 | 6725,106657 |
| SLC16A3 | ENSG00000141526 | 4664,825831 |
| CXCL16 | ENSG00000161921 | 4617,962996 |
| COTL1 | ENSG00000103187 | 5157,120369 |
| ATP6V1B2 | ENSG00000147416 | 4779,958808 |
| PFKFB3 | ENSG00000170525 | 6902,357143 |
| FLOT1 ENSG00000137312 | ENSG00000137312 | 5831,778057 |
| CDC42SE1 | ENSG00000197622 | 5193,729886 |
| MX2 | ENSG00000183486 | 6104,652825 |
| DAZAP2 | ENSG00000183283 | 5334,366813 |
| DDX17 | ENSG00000100201 | 5727,02129 |
| ZFP36L2 | ENSG00000152518 | 3990,801294 |
| MAP2K3 | ENSG00000034152 | 4387,349342 |
| TLE3 | ENSG00000140332 | 5234,087906 |
| STAT1 | ENSG00000115415 | 5275,631915 |
| SSH2 | ENSG00000141298 | 4294,167758 |
| RILPL2 | ENSG00000150977 | 4740,695893 |
| RHOA | ENSG00000067560 | 5281,490462 |

| | | |
|---|---|---|
| NUP98 | ENSG00000110713 | 5161,69903 |
| TMEM123 | ENSG00000152558 | 4622,433786 |
| EFHD2 | ENSG00000142634 | 3750,61646 |
| RAPGEF1 | ENSG00000107263 | 4362,058247 |
| NAMPTP1 | ENSG00000229644 | 5154,813157 |
| S100A11 | ENSG00000163191 | 5025,038553 |
| STK40 | ENSG00000196182 | 4058,127476 |
| ZC3H12A | ENSG00000163874 | 3844,480779 |
| PARP14 | ENSG00000173193 | 3681,622608 |
| DOCK5 | ENSG00000147459 | 5595,569789 |
| SQSTM1 ENSG00000161011 | ENSG00000161011 | 3830,11087 |
| HCK | ENSG00000101336 | 4074,805178 |
| SLC6A6 | ENSG00000131389 | 4673,285877 |
| HNRNPC | ENSG00000092199 | 4542,771685 |
| STAT3 | ENSG00000168610 | 4792,683196 |
| RNF149 | ENSG00000163162 | 5574,65057 |
| SPI1 | ENSG00000066336 | 3901,497553 |
| RGS2 | ENSG00000116741 | 4516,365101 |
| BTG1 | ENSG00000133639 | 4187,724512 |
| PIM2 | ENSG00000102096 | 3800,161686 |
| RAB8B | ENSG00000166128 | 5307,517457 |
| MARCKSL1 | ENSG00000175130 | 3709,235289 |
| RAB21 | ENSG00000080371 | 4543,349138 |
| MYD88 | ENSG00000172936 | 4144,524457 |
| GK | ENSG00000198814 | 5783,794707 |
| ZDHHC18 | ENSG00000204160 | 3904,627684 |
| CHD2 | ENSG00000173575 | 5435,605564 |
| BACH1 | ENSG00000156273 | 4729,202797 |
| NFKB1 | ENSG00000109320 | 4074,222644 |
| VSIR | ENSG00000107738 | 3243,200175 |
| BOD1L1 | ENSG00000038219 | 5015,724276 |
| PGK1 | ENSG00000102144 | 4399,14345 |
| SERPINB9 | ENSG00000170542 | 5566,342731 |
| GNA13 | ENSG00000120063 | 5171,402856 |
| ACTR3 | ENSG00000115091 | 4823,656554 |
| CFLAR | ENSG00000003402 | 4040,94014 |
| IL1B | ENSG00000125538 | 4511,799041 |
| LSP1 | ENSG00000130592 | 4128,28367 |
| MT-CO1 | ENSG00000198804 | 3719,103726 |
| SPAG9 | ENSG00000008294 | 4541,505209 |
| C15orf39 | ENSG00000167173 | 3667,962574 |
| GBP5 | ENSG00000154451 | 4910,736355 |
| HCLS1 | ENSG00000180353 | 4115,335007 |
| CAPZA1 | ENSG00000116489 | 4815,71078 |
| SAMD9 | ENSG00000205413 | 3174,898656 |
| GNAI2 | ENSG00000114353 | 3606,871094 |
| PICALM | ENSG00000073921 | 5442,237299 |
| PLEKHO2 | ENSG00000241839 | 3942,666959 |
| CSF2RB | ENSG00000100368 | 4123,381895 |
| C3AR1 | ENSG00000171860 | 3582,398202 |
| LRRFIP1 | ENSG00000124831 | 3989,658489 |
| YPEL5 | ENSG00000119801 | 5134,005738 |
| CPD | ENSG00000108582 | 5103,067378 |
| ARAP1 | ENSG00000186635 | 3761,579221 |
| NBN | ENSG00000104320 | 4765,797996 |

| | | |
|---|---|---|
| BAZ1A | ENSG00000198604 | 4125,804239 |
| PPP1R15A | ENSG00000087074 | 4664,48617 |
| GNB1 | ENSG00000078369 | 3951,179569 |
| ATP2B1 | ENSG00000070961 | 4616,710995 |
| CHST15 | ENSG00000182022 | 4483,380608 |
| PLEC | ENSG00000178209 | 3713,110821 |
| ATG2A | ENSG00000110046 | 4012,251968 |
| ATP13A3 | ENSG00000133657 | 5948,951115 |
| PPP1R18 ENSG00000146112 | ENSG00000146112 | 3466,751185 |
| SAMSN1 | ENSG00000155307 | 6044,334189 |
| DUSP1 | ENSG00000120129 | 4568,349245 |
| RAC2 | ENSG00000128340 | 3836,568252 |
| THEMIS2 | ENSG00000130775 | 3861,900718 |
| SERP1 | ENSG00000120742 | 3740,040042 |
| UBE2D3 | ENSG00000109332 | 4240,005262 |
| HERC5 | ENSG00000138646 | 2492,955806 |
| CCNL1 | ENSG00000163660 | 4887,997844 |
| CLIC1 ENSG00000213719 | ENSG00000213719 | 3854,181361 |
| WDR1 | ENSG00000071127 | 3369,904375 |
| ACTR2 | ENSG00000138071 | 4175,647326 |
| WIPF1 | ENSG00000115935 | 3861,79929 |
| NIBAN1 | ENSG00000135842 | 3534,177128 |
| OASL | ENSG00000135114 | 2407,093991 |
| SF3B1 | ENSG00000115524 | 4129,282121 |
| IRF1 | ENSG00000125347 | 3016,851086 |
| CD82 | ENSG00000085117 | 2627,206897 |
| KCTD20 | ENSG00000112078 | 3301,607015 |
| DDX5 | ENSG00000108654 | 4557,364162 |
| QKI | ENSG00000112531 | 4068,892539 |
| TLR2 | ENSG00000137462 | 4239,112727 |
| GRB2 | ENSG00000177885 | 3122,350945 |
| GAPDH | ENSG00000111640 | 3535,683084 |
| PFN1 | ENSG00000108518 | 3036,009935 |
| NEDD9 | ENSG00000111859 | 3727,69798 |
| DENND3 | ENSG00000105339 | 4405,829725 |
| CEBPB | ENSG00000172216 | 4201,14543 |
| MOB3A | ENSG00000172081 | 3120,985503 |
| SERINC1 | ENSG00000111897 | 3772,739828 |
| SIRPA | ENSG00000198053 | 2809,710018 |
| LRRK2 | ENSG00000188906 | 3251,522267 |
| SARAF | ENSG00000133872 | 3554,59143 |
| CDC42 | ENSG00000070831 | 3618,696049 |
| PRKCD | ENSG00000163932 | 3037,538711 |
| NABP1 | ENSG00000173559 | 4823,479112 |
| TMBIM6 | ENSG00000139644 | 3429,803812 |
| TRIM22 | ENSG00000132274 | 2723,860418 |
| MAP3K2 | ENSG00000169967 | 4063,008412 |
| PPP1CB | ENSG00000213639 | 4382,619022 |
| EIF1 | ENSG00000173812 | 3046,572262 |
| GRINA | ENSG00000178719 | 3578,582807 |
| JUNB | ENSG00000171223 | 2682,032446 |
| ITGB2 | ENSG00000160255 | 2731,418117 |
| AMPD2 | ENSG00000116337 | 3671,728512 |
| DDX60L | ENSG00000181381 | 3522,113109 |
| OXSR1 | ENSG00000172939 | 3077,414396 |

| | | |
|---|---|---|
| TAGAP | ENSG00000164691 | 3823,258383 |
| BHLHE40 | ENSG00000134107 | 4599,72721 |
| IL10RA | ENSG00000110324 | 2528,277873 |
| CHD1 | ENSG00000153922 | 4568,898197 |
| MEFV | ENSG00000103313 | 3207,99402 |
| PTP4A2 | ENSG00000184007 | 3259,434381 |
| FFAR2 | ENSG00000126262 | 2497,456636 |
| SLA | ENSG00000155926 | 4131,994625 |
| ZMIZ1 | ENSG00000108175 | 2624,872824 |
| TLN1 | ENSG00000137076 | 3637,780612 |
| OLR1 | ENSG00000173391 | 4472,193595 |
| CDC42EP3 | ENSG00000163171 | 4069,224643 |
| STK4 | ENSG00000101109 | 3174,682155 |
| NADK | ENSG00000008130 | 3334,765007 |
| BID | ENSG00000015475 | 3116,00135 |
| TNFRSF1A | ENSG00000067182 | 3949,446755 |
| PTPRE | ENSG00000132334 | 3808,904673 |
| B4GALT5 | ENSG00000158470 | 2837,558284 |
| ICAM1 | ENSG00000090339 | 3395,193324 |
| ZNFX1 | ENSG00000124201 | 3393,177765 |
| ARPC2 | ENSG00000163466 | 3882,627794 |
| PRKAR1A | ENSG00000108946 | 3309,570602 |
| ARRB2 | ENSG00000141480 | 3328,110278 |
| ISG15 | ENSG00000187608 | 1719,567369 |
| C9orf72 | ENSG00000147894 | 3976,817945 |
| SERINC3 | ENSG00000132824 | 3600,833604 |
| ZNF267 | ENSG00000185947 | 3954,226355 |
| EIF4G2 | ENSG00000110321 | 3659,02495 |
| STAT5B | ENSG00000173757 | 2910,093491 |
| CSRNP1 | ENSG00000144655 | 3428,587959 |
| ARHGDIB | ENSG00000111348 | 3161,839887 |
| DSE | ENSG00000111817 | 2866,047112 |
| PAK1 | ENSG00000149269 | 3080,889787 |
| IFITM3 | ENSG00000142089 | 2210,220828 |
| ABHD2 | ENSG00000140526 | 2592,150784 |
| GNAI3 | ENSG00000065135 | 3577,337822 |
| TSC22D3 | ENSG00000157514 | 2273,298667 |
| CD46 | ENSG00000117335 | 3672,311236 |
| OAS3 | ENSG00000111331 | 1621,191553 |
| CHMP4B | ENSG00000101421 | 3111,470176 |
| CNN2 | ENSG00000064666 | 2935,002583 |
| NUMB | ENSG00000133961 | 2955,950385 |
| IRAK2 | ENSG00000134070 | 3091,125086 |
| GABARAPL1 | ENSG00000139112 | 3261,84374 |
| R3HDM4 | ENSG00000198858 | 2511,840222 |
| ARPC5 | ENSG00000162704 | 3437,022478 |
| FCGR2A | ENSG00000143226 | 3195,680626 |
| FPR2 | ENSG00000171049 | 3052,704419 |
| CYLD | ENSG00000083799 | 3063,795103 |
| PPP1R15B | ENSG00000158615 | 3717,639149 |
| SUSD6 | ENSG00000100647 | 2856,434538 |
| NOTCH1 | ENSG00000148400 | 2021,483143 |
| LUCAT1 | ENSG00000248323 | 3215,417787 |
| TMEM127 | ENSG00000135956 | 2425,65324 |
| JMJD1C | ENSG00000171988 | 3843,984294 |

| | | |
|---|---|---|
| GTPBP1 | ENSG00000100226 | 3593,258523 |
| MT-ND4 | ENSG00000198886 | 2894,220564 |
| CDC42SE2 | ENSG00000158985 | 2841,52384 |
| IER5 | ENSG00000162783 | 2255,030575 |
| BIRC3 | ENSG00000023445 | 2461,852331 |
| TPM3 | ENSG00000143549 | 3417,834127 |
| PTPN12 | ENSG00000127947 | 3372,214131 |
| MAP3K8 | ENSG00000107968 | 3604,355049 |
| STK10 | ENSG00000072786 | 3003,536023 |
| TRIB1 | ENSG00000173334 | 3545,964228 |
| TYROBP | ENSG00000011600 | 2592,665332 |
| MAP4K4 | ENSG00000071054 | 2344,297368 |
| VIM | ENSG00000026025 | 2789,227067 |
| VEGFA | ENSG00000112715 | 4068,543523 |
| FAM107B | ENSG00000065809 | 2663,000876 |
| CCNG2 | ENSG00000138764 | 2126,219285 |
| FCER1G | ENSG00000158869 | 3362,834213 |
| ABCA1 | ENSG00000165029 | 3981,775187 |
| MYO9B | ENSG00000099331 | 2865,144985 |
| NOTCH2 | ENSG00000134250 | 2277,931872 |
| PSEN1 | ENSG00000080815 | 2489,749457 |
| HNRNPK | ENSG00000165119 | 2658,513998 |
| PLK3 | ENSG00000173846 | 3171,554928 |
| GBP1 | ENSG00000117228 | 2451,491199 |
| PILRA | ENSG00000085514 | 2145,655046 |
| CYFIP2 | ENSG00000055163 | 2776,426856 |
| HLA-DRA ENSG00000204287 | ENSG00000204287 | 2045,578505 |
| RAB31 | ENSG00000168461 | 2337,300282 |
| NFKBID | ENSG00000167604 | 2377,721426 |
| ZFP36 | ENSG00000128016 | 3139,635734 |
| AKNA | ENSG00000106948 | 1681,477112 |
| TMEM71 | ENSG00000165071 | 3374,763915 |
| CCL4L2 ENSG00000276070 | ENSG00000276070 | 1476,460027 |
| DOK3 | ENSG00000146094 | 2704,859981 |
| SP3 | ENSG00000172845 | 2809,486915 |
| INSIG1 | ENSG00000186480 | 2232,184619 |
| ERGIC1 | ENSG00000113719 | 2477,280501 |
| CD74 | ENSG00000019582 | 2228,636988 |
| CD93 | ENSG00000125810 | 2224,791024 |
| DGAT2 | ENSG00000062282 | 2820,572827 |
| HSP90AA1 | ENSG00000080824 | 2283,20471 |
| TOM1 | ENSG00000100284 | 2516,661686 |
| ELOVL5 | ENSG00000012660 | 2549,126403 |
| PIK3AP1 | ENSG00000155629 | 2819,610567 |
| IFIT5 | ENSG00000152778 | 1914,449348 |
| FCAR ENSG00000186431 | ENSG00000186431 | 2766,366931 |
| UBE2B | ENSG00000119048 | 2902,897399 |
| IL6R | ENSG00000160712 | 2041,161435 |
| PECAM1 | ENSG00000261371 | 2500,478815 |
| CAMKK2 | ENSG00000110931 | 2102,541498 |
| ANP32A | ENSG00000140350 | 2278,585863 |
| RHOG | ENSG00000177105 | 2138,341825 |
| APOBEC3A ENSG00000128383 | ENSG00000128383 | 2415,44527 |
| MNDA | ENSG00000163563 | 2693,346906 |
| DDX6 | ENSG00000110367 | 2530,104102 |

| | | |
|---|---|---|
| PIM3 | ENSG00000198355 | 2284,671051 |
| LAMP2 | ENSG00000005893 | 3044,971771 |
| XPO6 | ENSG00000169180 | 3188,838443 |
| PDLIM7 | ENSG00000196923 | 2642,756134 |
| PACSIN2 | ENSG00000100266 | 1955,68835 |
| MT-CO3 | ENSG00000198938 | 2482,765254 |
| SAMD9L | ENSG00000177409 | 2177,489867 |
| KMT2E | ENSG00000005483 | 2691,365962 |
| LYZ | ENSG00000090382 | 2178,115994 |
| ARF1 | ENSG00000143761 | 2374,262969 |
| DBNL | ENSG00000136279 | 2513,575437 |
| IQSEC1 | ENSG00000144711 | 2544,615514 |
| RIGI | ENSG00000107201 | 2166,628741 |
| WTAP | ENSG00000146457 | 2658,353284 |
| CCNI | ENSG00000118816 | 2719,431125 |
| KLF6 | ENSG00000067082 | 2642,798533 |
| IRF2BP2 | ENSG00000168264 | 2447,253736 |
| ISG20 | ENSG00000172183 | 2299,834602 |
| GADD45B | ENSG00000099860 | 2367,918516 |
| CNBP | ENSG00000169714 | 2517,831129 |
| CKAP4 | ENSG00000136026 | 2107,579822 |
| JOSD1 | ENSG00000100221 | 2348,521209 |
| REL | ENSG00000162924 | 2628,946909 |
| ITGA5 | ENSG00000161638 | 2748,314749 |
| BCL3 | ENSG00000069399 | 2568,014076 |
| ATP6V1A | ENSG00000114573 | 1932,998824 |
| USP15 | ENSG00000135655 | 3284,365809 |
| NCOA4 | ENSG00000266412 | 2285,483776 |
| ARHGAP26 | ENSG00000145819 | 2503,258601 |
| ADIPOR1 | ENSG00000159346 | 3036,546351 |
| TOR1B | ENSG00000136816 | 1897,39438 |
| TMBIM1 | ENSG00000135926 | 1864,755031 |
| OGA | ENSG00000198408 | 2136,235355 |
| HNRNPU | ENSG00000153187 | 2445,691426 |
| UBR4 | ENSG00000127481 | 2364,039076 |
| IFRD1 | ENSG00000006652 | 3378,004647 |
| NFAM1 | ENSG00000235568 | 1695,291981 |
| SBNO2 ENSG00000064932 | ENSG00000064932 | 2074,556218 |
| HELZ2 | ENSG00000130589 | 1996,662077 |
| RIT1 | ENSG00000143622 | 3108,016287 |
| ELF1 | ENSG00000120690 | 2795,310409 |
| RIPOR2 | ENSG00000111913 | 1699,150973 |
| RICTOR | ENSG00000164327 | 2399,677488 |
| RNF144B | ENSG00000137393 | 1728,020989 |
| SASH3 | ENSG00000122122 | 2064,101094 |
| OAZ1 | ENSG00000104904 | 2199,327043 |
| NBEAL2 | ENSG00000160796 | 2581,672949 |
| MBOAT7 ENSG00000125505 | ENSG00000125505 | 2220,866969 |
| IFIH1 | ENSG00000115267 | 2143,412232 |
| EIF2AK2 | ENSG00000055332 | 1858,213562 |
| USP10 | ENSG00000103194 | 1638,482215 |
| RPL3 | ENSG00000100316 | 1475,378608 |
| SP100 | ENSG00000067066 | 2525,387627 |
| MYL12B | ENSG00000118680 | 2409,849043 |
| EHBP1L1 | ENSG00000173442 | 1716,468301 |

| | | |
|---|---|---|
| TAP1 ENSG00000168394 | ENSG00000168394 | 1874,271171 |
| C15orf48 | ENSG00000166920 | 1929,466425 |
| PARP9 | ENSG00000138496 | 2049,885097 |
| GRAMD1A | ENSG00000089351 | 1821,610971 |
| CERK | ENSG00000100422 | 2139,791512 |
| SLC12A6 | ENSG00000140199 | 2403,042146 |
| IL1R2 | ENSG00000115590 | 2249,195951 |
| JAK1 | ENSG00000162434 | 2383,440288 |
| SLC15A4 | ENSG00000139370 | 1593,494586 |
| MT-CO2 | ENSG00000198712 | 1947,802415 |
| ITPRIP | ENSG00000148841 | 2030,477663 |
| HNRNPA2B1 | ENSG00000122566 | 2979,241525 |
| PAK2 | ENSG00000180370 | 2260,588279 |
| ARHGAP30 | ENSG00000186517 | 1802,645273 |
| HECA | ENSG00000112406 | 2366,17336 |
| ATP2A3 | ENSG00000074370 | 2425,390886 |
| KLF10 | ENSG00000155090 | 2283,529711 |
| STX11 | ENSG00000135604 | 2211,335915 |
| DOCK8 | ENSG00000107099 | 2313,399853 |
| MPEG1 | ENSG00000197629 | 1644,384989 |
| RALB | ENSG00000144118 | 2194,849349 |
| HIVEP2 | ENSG00000010818 | 2126,185326 |
| SRRM2 | ENSG00000167978 | 2411,017095 |
| PTPN6 | ENSG00000111679 | 1882,099609 |
| IL4R | ENSG00000077238 | 2597,279579 |
| STAT6 | ENSG00000166888 | 1855,958936 |
| RAP1B | ENSG00000127314 | 1875,902419 |
| CD83 | ENSG00000112149 | 1693,701624 |
| PRRC2C | ENSG00000117523 | 2246,778996 |
| TAB2 | ENSG00000055208 | 2351,788329 |
| GRK2 | ENSG00000173020 | 1719,682635 |
| CCR1 | ENSG00000163823 | 2068,970001 |
| BZW1 | ENSG00000082153 | 2206,009679 |
| PLSCR1 | ENSG00000188313 | 1952,227273 |
| IGF2R | ENSG00000197081 | 1833,743868 |
| GPBP1 | ENSG00000062194 | 2357,163895 |
| BIRC2 | ENSG00000110330 | 2582,189632 |
| PJA2 | ENSG00000198961 | 2160,313529 |
| CSNK1G2 | ENSG00000133275 | 1633,467503 |
| STK17B | ENSG00000081320 | 2063,836401 |
| MED13L | ENSG00000123066 | 2234,556786 |
| B4GALT1 | ENSG00000086062 | 2075,198911 |
| RASSF3 | ENSG00000153179 | 1838,541921 |
| IRAK3 | ENSG00000090376 | 1681,064343 |
| VOPP1 | ENSG00000154978 | 1594,466429 |
| MIR23AHG | ENSG00000267519 | 2228,277593 |
| ARL5B | ENSG00000165997 | 2692,539442 |
| TPD52L2 | ENSG00000101150 | 2093,364565 |
| CYTIP | ENSG00000115165 | 2155,888265 |
| RPL41 | ENSG00000229117 | 1666,285198 |
| SYAP1 | ENSG00000169895 | 2358,019485 |
| TMCC3 | ENSG00000057704 | 2378,671575 |
| RPS6 | ENSG00000137154 | 1666,852046 |
| SCARF1 ENSG00000074660 | ENSG00000074660 | 2063,146725 |
| TACC1 | ENSG00000147526 | 1703,346026 |

| | | |
|---|---|---|
| ADGRE5 | ENSG00000123146 | 2061,912486 |
| LIMS1 | ENSG00000169756 | 2101,140269 |
| PTTG1IP | ENSG00000183255 | 1360,167074 |
| LAT2 | ENSG00000086730 | 1771,544062 |
| DNM2 | ENSG00000079805 | 2340,003972 |
| ZYX ENSG00000159840 | ENSG00000159840 | 2007,151654 |
| ARRDC2 | ENSG00000105643 | 1234,282219 |
| CREM | ENSG00000095794 | 2798,30708 |
| EEF2 | ENSG00000167658 | 1265,707176 |
| RPS9 ENSG00000170889 | ENSG00000170889 | 1609,113833 |
| CLK1 | ENSG00000013441 | 2431,4081 |
| FOS | ENSG00000170345 | 2181,511721 |
| ETF1 | ENSG00000120705 | 2368,891892 |
| SLC15A3 | ENSG00000110446 | 1775,87254 |
| MOB1A | ENSG00000114978 | 2091,578153 |
| TGFB1 | ENSG00000105329 | 1622,602777 |
| TNFSF13B | ENSG00000102524 | 1815,624335 |
| PCNX1 | ENSG00000100731 | 2116,083259 |
| TFE3 | ENSG00000068323 | 1782,60503 |
| MAP7D1 | ENSG00000116871 | 1736,643665 |
| ARID4B | ENSG00000054267 | 2265,498011 |
| TALDO1 | ENSG00000177156 | 1639,216527 |
| MAP1LC3B | ENSG00000140941 | 2586,83099 |
| CXCR2 | ENSG00000180871 | 1742,584553 |
| RPS6KA1 ENSG00000117676 | ENSG00000117676 | 1526,501667 |
| FCGR3A | ENSG00000203747 | 1722,430816 |
| MIDN | ENSG00000167470 | 2073,617022 |
| ARHGAP45 | ENSG00000180448 | 1634,603186 |
| IFI16 | ENSG00000163565 | 1840,905346 |
| GAB2 | ENSG00000033327 | 2029,849162 |
| PITPNA | ENSG00000174238 | 1930,947557 |
| MMP25 | ENSG00000008516 | 1709,040435 |
| TGM2 | ENSG00000198959 | 1600,60628 |
| HSH2D | ENSG00000196684 | 1768,830469 |
| MKNK2 | ENSG00000099875 | 1558,816244 |
| DIAPH1 | ENSG00000131504 | 1909,184934 |
| CSDE1 | ENSG00000009307 | 2092,897225 |
| S100P | ENSG00000163993 | 1381,968396 |
| CLEC4E | ENSG00000166523 | 1600,103047 |
| CDKN1A | ENSG00000124762 | 1831,841677 |
| TIMP1 | ENSG00000102265 | 2146,697799 |
| PPP1R3B ENSG00000173281 | ENSG00000173281 | 2237,802042 |
| SP110 | ENSG00000135899 | 1891,253798 |
| TDP2 | ENSG00000111802 | 2081,838296 |
| ARPC3 | ENSG00000111229 | 2154,857201 |
| ANKRD13A | ENSG00000076513 | 1691,872285 |
| NCF1 | ENSG00000158517 | 1478,512257 |
| TCIRG1 | ENSG00000110719 | 1769,825436 |
| PHACTR1 | ENSG00000112137 | 2248,802047 |
| CD58 | ENSG00000116815 | 2141,041681 |
| LRP10 | ENSG00000197324 | 1630,243996 |
| PHF20L1 | ENSG00000129292 | 2837,664761 |
| ETS2 | ENSG00000157557 | 3067,789793 |
| FYB1 | ENSG00000082074 | 2121,520078 |
| SLC7A5 | ENSG00000103257 | 1838,83786 |

| | | |
|---|---|---|
| AMPD3 | ENSG00000133805 | 2001,114784 |
| TAPBP ENSG00000231925 | ENSG00000231925 | 1747,488764 |
| LMNB1 | ENSG00000113368 | 2277,555185 |
| OSM | ENSG00000099985 | 2727,17949 |
| CYBB | ENSG00000165168 | 1516,098897 |
| CYBA | ENSG00000051523 | 1567,720235 |
| HIP1 | ENSG00000127946 | 1762,685552 |
| RBM39 | ENSG00000131051 | 2169,043255 |
| LYST | ENSG00000143669 | 2161,933568 |
| MAPRE1 | ENSG00000101367 | 1790,787174 |
| AGO2 | ENSG00000123908 | 1933,896329 |
| GNAS | ENSG00000087460 | 1809,954931 |
| FNBP1 | ENSG00000187239 | 1549,230056 |
| MAPKAPK2 | ENSG00000162889 | 1825,432692 |
| MYL6 | ENSG00000092841 | 1876,103125 |
| STAT2 | ENSG00000170581 | 1479,53481 |
| IL1R1 | ENSG00000115594 | 2146,402808 |
| USP9X | ENSG00000124486 | 1922,498634 |
| QSOX1 | ENSG00000116260 | 1812,931544 |
| FERMT3 | ENSG00000149781 | 1607,292713 |
| ADM | ENSG00000148926 | 1904,995536 |
| ATG16L2 | ENSG00000168010 | 1654,195532 |
| AFTPH | ENSG00000119844 | 2069,202074 |
| MAPK13 | ENSG00000156711 | 2037,880054 |
| SHKBP1 | ENSG00000160410 | 1601,490605 |
| LASP1 | ENSG00000002834 | 1392,035311 |
| ZCCHC2 | ENSG00000141664 | 1248,458021 |
| ZEB2 | ENSG00000169554 | 1974,994002 |
| IFNAR1 | ENSG00000142166 | 1790,482887 |
| RPL4 | ENSG00000174444 | 1324,40837 |
| B3GNT5 | ENSG00000176597 | 2663,154182 |
| RPL13A | ENSG00000142541 | 1195,372061 |
| VAV1 | ENSG00000141968 | 1787,855874 |
| IER3 ENSG00000137331 | ENSG00000137331 | 1881,659117 |
| LRRC25 | ENSG00000175489 | 1309,394464 |
| S100A6 | ENSG00000197956 | 1454,958724 |
| TGFBR2 | ENSG00000163513 | 1660,705073 |
| PTGES | ENSG00000148344 | 1579,78592 |
| SNX10 | ENSG00000086300 | 2237,952898 |
| MGAM ENSG00000257335 | ENSG00000257335 | 1701,286907 |
| ELL2 | ENSG00000118985 | 2351,446229 |
| SEMA4D | ENSG00000187764 | 2062,69235 |
| TRIM38 | ENSG00000112343 | 1694,395809 |
| SIGLEC14 | ENSG00000254415 | 1428,355374 |
| MYADM | ENSG00000179820 | 2124,782976 |
| PTPRJ | ENSG00000149177 | 1348,769775 |
| ACTN1 | ENSG00000072110 | 2377,217688 |
| TUT7 | ENSG00000083223 | 2001,559864 |
| ENO1 | ENSG00000074800 | 1410,005569 |
| VMP1 | ENSG00000062716 | 2082,126958 |
| APOL6 | ENSG00000221963 | 1224,087405 |
| WDR26 | ENSG00000162923 | 2049,064703 |
| MT-CYB | ENSG00000198727 | 1659,279953 |
| STX3 | ENSG00000166900 | 1814,480911 |
| PTMA | ENSG00000187514 | 1391,757245 |

| | | |
|---|---|---|
| CYP4F3 | ENSG00000186529 | 1709,783264 |
| ARHGEF2 | ENSG00000116584 | 1971,413215 |
| KPNB1 | ENSG00000108424 | 1618,648939 |
| CFL1 | ENSG00000172757 | 1768,743376 |
| APOBR | ENSG00000184730 | 1226,310605 |
| CORO1A | ENSG00000102879 | 1344,055371 |
| SH3BP2 | ENSG00000087266 | 1577,360618 |
| ELF4 | ENSG00000102034 | 1719,379191 |
| RPL10 | ENSG00000147403 | 1167,444834 |
| RUBCNL | ENSG00000102445 | 2211,213823 |
| RARA | ENSG00000131759 | 2213,892389 |
| RBPJ | ENSG00000168214 | 2221,45489 |
| BCL6 | ENSG00000113916 | 2545,57629 |
| ORAI2 | ENSG00000160991 | 1097,423913 |
| H2AC6 | ENSG00000180573 | 1304,115452 |
| BRD2 ENSG00000204256 | ENSG00000204256 | 1475,504155 |
| ADGRE2 | ENSG00000127507 | 1736,987292 |
| TNFSF10 | ENSG00000121858 | 963,7352427 |
| CLTC | ENSG00000141367 | 1660,346737 |
| DNAJA1 | ENSG00000086061 | 1614,036815 |
| SOCS3 | ENSG00000184557 | 2335,305116 |
| GPSM3 ENSG00000213654 | ENSG00000213654 | 1466,502743 |
| SNX18 | ENSG00000178996 | 2068,618669 |
| STX6 | ENSG00000135823 | 1732,659253 |
| CMPK2 | ENSG00000134326 | 875,9030565 |
| RAC1 | ENSG00000136238 | 1632,98226 |
| DHX34 | ENSG00000134815 | 1725,579113 |
| MT-ND1 | ENSG00000198888 | 1895,612516 |
| SLC16A6 | ENSG00000108932 | 1238,261927 |
| ANKRD12 | ENSG00000101745 | 1556,156736 |
| ZSWIM6 | ENSG00000130449 | 2115,960841 |
| SHOC2 | ENSG00000108061 | 1739,917701 |
| RXRA | ENSG00000186350 | 1220,164948 |
| ANTXR2 | ENSG00000163297 | 1546,12573 |
| CAP1 | ENSG00000131236 | 1834,536353 |
| PURB | ENSG00000146676 | 1830,365778 |
| KIAA0513 | ENSG00000135709 | 1137,423249 |
| GBP4 | ENSG00000162654 | 1179,046536 |
| RRM2B | ENSG00000048392 | 1510,70147 |
| JARID2 | ENSG00000008083 | 1592,537981 |
| RPS27 | ENSG00000177954 | 1235,986902 |
| IFI6 | ENSG00000126709 | 925,7265679 |
| OAS2 | ENSG00000111335 | 922,9637919 |
| NIN | ENSG00000100503 | 1845,905451 |
| MPP1 | ENSG00000130830 | 2142,797636 |
| CLIC4 | ENSG00000169504 | 1114,272073 |
| TRIP12 | ENSG00000153827 | 1695,831153 |
| PIM1 | ENSG00000137193 | 1859,133624 |
| RAB7A | ENSG00000075785 | 1656,076149 |
| WASHC4 | ENSG00000136051 | 1782,580929 |
| FNDC3B | ENSG00000075420 | 1570,47494 |
| ELL | ENSG00000105656 | 1665,117496 |
| PDCD4 | ENSG00000150593 | 1786,948899 |
| ZFAND5 | ENSG00000107372 | 1661,156248 |
| CCL4 ENSG00000275302 | ENSG00000275302 | 969,1073457 |

| | | |
|---|---|---|
| WBP2 | ENSG00000132471 | 1598,059547 |
| FGD3 | ENSG00000127084 | 1366,942316 |
| CYTH4 | ENSG00000100055 | 1709,682023 |
| HSPA5 | ENSG00000044574 | 2020,944016 |
| MME | ENSG00000196549 | 2086,474831 |
| SIPA1L1 | ENSG00000197555 | 1610,22917 |
| VPS35 | ENSG00000069329 | 1709,667895 |
| EVI2B | ENSG00000185862 | 1720,733888 |
| HMGA1 | ENSG00000137309 | 1748,108021 |
| ERN1 | ENSG00000178607 | 1418,879878 |
| VDR | ENSG00000111424 | 1687,158136 |
| PRKDC | ENSG00000253729 | 1881,351413 |
| PTGS2 | ENSG00000073756 | 1670,430397 |
| IGSF6 ENSG00000140749 | ENSG00000140749 | 1576,019386 |
| RYBP ENSG00000163602 | ENSG00000163602 | 1612,688923 |
| RPS4X | ENSG00000198034 | 1137,858754 |
| DTX3L | ENSG00000163840 | 1142,672559 |
| YTHDF3 | ENSG00000185728 | 1571,832779 |
| CAPZA2 | ENSG00000198898 | 1765,067697 |
| AP1G1 | ENSG00000166747 | 1772,013531 |
| PPP4R1 | ENSG00000154845 | 1526,043878 |
| PTK2B | ENSG00000120899 | 1432,810105 |
| ALOX5 ENSG00000012779 | ENSG00000012779 | 1623,576274 |
| TANK | ENSG00000136560 | 1804,140869 |
| TYMP | ENSG00000025708 | 1276,184864 |
| CREBBP | ENSG00000005339 | 1321,518526 |
| ZNF217 | ENSG00000171940 | 1220,414775 |
| RNF13 | ENSG00000082996 | 1845,593117 |
| SON | ENSG00000159140 | 1425,149747 |
| NR3C1 | ENSG00000113580 | 1510,395227 |
| ATP6V0B | ENSG00000117410 | 1370,29291 |
| RAD21 | ENSG00000164754 | 1492,655129 |
| LILRB3 ENSG00000204577 | ENSG00000204577 | 1525,18054 |
| STAT5A | ENSG00000126561 | 948,735723 |
| FBRS | ENSG00000156860 | 1213,744759 |
| STK17A | ENSG00000164543 | 1514,106121 |
| RAP2C | ENSG00000123728 | 1664,955528 |
| KRAS | ENSG00000133703 | 1667,026858 |
| TET2 | ENSG00000168769 | 2042,662632 |
| RAF1 | ENSG00000132155 | 1342,827593 |
| SEC62 | ENSG00000008952 | 1786,396857 |
| GNS | ENSG00000135677 | 1727,714329 |
| NFKBIE | ENSG00000146232 | 1376,960588 |
| STAG2 | ENSG00000101972 | 1707,891503 |
| PPP2R5C | ENSG00000078304 | 1483,679575 |
| RPS11 | ENSG00000142534 | 1171,824192 |
| MT-ND2 | ENSG00000198763 | 1574,147584 |
| DMXL2 | ENSG00000104093 | 1201,347712 |
| CD14 | ENSG00000170458 | 1961,433436 |
| GCC2 | ENSG00000135968 | 1739,498057 |
| LBR | ENSG00000143815 | 1800,47716 |
| ERBIN | ENSG00000112851 | 1285,334867 |
| OSBPL8 | ENSG00000091039 | 1632,997712 |
| CLEC7A | ENSG00000172243 | 2158,705446 |
| IL1RAP | ENSG00000196083 | 2008,636359 |

| | | |
|---|---|---|
| GPR65 | ENSG00000140030 | 1650,146363 |
| CYTH1 | ENSG00000108669 | 1550,060596 |
| CD69 | ENSG00000110848 | 1315,793027 |
| ABTB1 | ENSG00000114626 | 920,0734065 |
| UBE2H | ENSG00000186591 | 1406,884671 |
| SIPA1 | ENSG00000213445 | 1378,540478 |
| HIVEP1 | ENSG00000095951 | 1489,11197 |
| WAC | ENSG00000095787 | 1655,43671 |
| EIF2S3 | ENSG00000130741 | 1304,924586 |
| FOXO3 | ENSG00000118689 | 1561,304116 |
| GNB2 | ENSG00000172354 | 1281,454011 |
| CAPZB | ENSG00000077549 | 1463,860283 |
| RTN4 | ENSG00000115310 | 1592,286354 |
| MYL12A | ENSG00000101608 | 1760,899759 |
| SKI | ENSG00000157933 | 1200,605489 |
| PGGHG | ENSG00000142102 | 1466,23203 |
| MFN2 | ENSG00000116688 | 1651,871451 |
| KCNJ15 | ENSG00000157551 | 1859,625843 |
| TYK2 | ENSG00000105397 | 1404,198378 |
| TM9SF3 | ENSG00000077147 | 1592,620988 |
| RELA | ENSG00000173039 | 1389,932415 |
| USP32 | ENSG00000170832 | 1997,440229 |
| PI3 | ENSG00000124102 | 681,6965139 |
| RAPGEF2 | ENSG00000109756 | 1627,918428 |
| RAP1A | ENSG00000116473 | 1549,038217 |
| DUSP2 | ENSG00000158050 | 1061,341037 |
| ZBTB18 | ENSG00000179456 | 1253,242872 |
| ARID5A | ENSG00000196843 | 1465,511405 |
| LDHA | ENSG00000134333 | 1123,356416 |
| SNRK | ENSG00000163788 | 1449,56436 |
| SETX | ENSG00000107290 | 1344,02396 |
| PHF12 | ENSG00000109118 | 1310,774372 |
| MAP3K11 | ENSG00000173327 | 1078,308683 |
| KAT6A | ENSG00000083168 | 1446,379427 |
| ATP1A1 | ENSG00000163399 | 1363,221246 |
| ADPGK | ENSG00000159322 | 1629,289851 |
| MSL1 | ENSG00000188895 | 1296,934646 |
| LRG1 | ENSG00000171236 | 1258,034314 |
| GALC | ENSG00000054983 | 1388,551151 |
| TECPR2 | ENSG00000196663 | 951,7379003 |
| FRMD4B | ENSG00000114541 | 2019,178162 |
| S100A4 | ENSG00000196154 | 1205,234013 |
| MAPK1 | ENSG00000100030 | 1081,621295 |
| CHSY1 | ENSG00000131873 | 1791,063682 |
| GHITM | ENSG00000165678 | 1237,823898 |
| SUN2 | ENSG00000100242 | 1244,802545 |
| PRR5L | ENSG00000135362 | 1003,663155 |
| NFE2L2 | ENSG00000116044 | 1737,368867 |
| FAM8A1 | ENSG00000137414 | 1337,864709 |
| NFIL3 | ENSG00000165030 | 1783,651024 |
| ADGRE3 | ENSG00000131355 | 1575,235224 |
| IFI44 | ENSG00000137965 | 830,046882 |
| SUPT6H | ENSG00000109111 | 1623,959506 |
| EIF5 | ENSG00000100664 | 1552,260355 |
| SCAF11 | ENSG00000139218 | 1468,743992 |

| | | |
|---|---|---|
| MBP | ENSG00000197971 | 1488,385755 |
| FAS | ENSG00000026103 | 1175,792927 |
| INPP5D ENSG00000168918 | ENSG00000168918 | 1546,547027 |
| ATP6V1C1 | ENSG00000155097 | 1437,668142 |
| RAB5A | ENSG00000144566 | 1419,672245 |
| DDIT4 | ENSG00000168209 | 1190,296879 |
| CHP1 | ENSG00000187446 | 1333,049737 |
| DNAJC3 | ENSG00000102580 | 1068,198182 |
| MT-ND5 | ENSG00000198786 | 1291,766987 |
| SLC45A4 | ENSG00000022567 | 1357,321077 |
| BNIP3L | ENSG00000104765 | 1328,311202 |
| RNF166 | ENSG00000158717 | 1365,640542 |
| DDX21 | ENSG00000165732 | 1760,90488 |
| PTBP3 | ENSG00000119314 | 1506,297245 |
| GMFG | ENSG00000130755 | 1326,342215 |
| NECAP1 | ENSG00000089818 | 1320,878701 |
| DNAJA2 | ENSG00000069345 | 1423,238939 |
| PHC2 | ENSG00000134686 | 1290,084105 |
| PPT1 | ENSG00000131238 | 1323,527403 |
| TAF7 | ENSG00000178913 | 1397,154412 |
| C5AR2 | ENSG00000134830 | 975,4736251 |
| RPS3 | ENSG00000149273 | 856,4420112 |
| KMT2C | ENSG00000055609 | 1383,527229 |
| NMI | ENSG00000123609 | 1318,338741 |
| CSNK1D | ENSG00000141551 | 1354,257427 |
| THAP9-AS1 | ENSG00000251022 | 1856,430342 |
| HNRNPA3 | ENSG00000170144 | 1705,283593 |
| RTN3 | ENSG00000133318 | 1454,543983 |
| PER1 | ENSG00000179094 | 1177,315402 |
| IST1 | ENSG00000182149 | 1326,497155 |
| TNFRSF10B | ENSG00000120889 | 891,3988632 |
| YPEL3 | ENSG00000090238 | 1164,590934 |
| CHASERR | ENSG00000272888 | 1601,324309 |
| GRK6 | ENSG00000198055 | 976,6992627 |
| ASAP1 | ENSG00000153317 | 1526,392412 |
| PTEN ENSG00000171862 | ENSG00000171862 | 1537,313891 |
| SPATA13 ENSG00000182957 | ENSG00000182957 | 1582,103696 |
| ARF4 | ENSG00000168374 | 1503,836182 |
| EML4 | ENSG00000143924 | 1615,82934 |
| CSGALNACT2 | ENSG00000169826 | 1558,062503 |
| TKT | ENSG00000163931 | 1055,806024 |
| LAMB3 | ENSG00000196878 | 1494,977858 |
| GRN | ENSG00000030582 | 1336,902939 |
| SGK1 | ENSG00000118515 | 1187,039585 |
| PLEKHB2 | ENSG00000115762 | 1706,371602 |
| GABARAPL2 | ENSG00000034713 | 1196,74143 |
| DNAJB6 | ENSG00000105993 | 1314,670563 |
| OSER1 | ENSG00000132823 | 1236,762356 |
| MED13 | ENSG00000108510 | 1272,171928 |
| TFDP1 | ENSG00000198176 | 1530,837335 |
| RTF2 | ENSG00000022277 | 1369,897399 |
| NHSL2 | ENSG00000204131 | 965,9542798 |
| CPEB4 | ENSG00000113742 | 1479,220701 |
| BRPF3 | ENSG00000096070 | 1008,989726 |
| HIPK1 | ENSG00000163349 | 1252,588713 |

| | | |
|---|---|---|
| FUS | ENSG00000089280 | 1679,628507 |
| HCP5 ENSG00000206337 | ENSG00000206337 | 1056,976739 |
| HLA-F ENSG00000204642 | ENSG00000204642 | 948,2159092 |
| CARD8 | ENSG00000105483 | 1338,357087 |
| UNC119 | ENSG00000109103 | 977,0941559 |
| PADI2 | ENSG00000117115 | 1617,526072 |
| HMGB2 | ENSG00000164104 | 838,5873151 |
| AIF1 ENSG00000204472 | ENSG00000204472 | 1407,435868 |
| PPM1A | ENSG00000100614 | 1283,898172 |
| SIGLEC10 | ENSG00000142512 | 771,6762842 |
| TXNRD1 | ENSG00000198431 | 1293,46394 |
| FNIP1 | ENSG00000217128 | 1407,498569 |
| RNF141 | ENSG00000110315 | 1481,190382 |
| RPL21 | ENSG00000122026 | 1097,78101 |
| MARF1 ENSG00000166783 | ENSG00000166783 | 1128,642769 |
| DAPP1 | ENSG00000070190 | 1152,404608 |
| TGOLN2 | ENSG00000152291 | 1081,023117 |
| RFFL | ENSG00000092871 | 998,6191711 |
| RPL13 | ENSG00000167526 | 750,2374512 |
| PPP1R9B | ENSG00000108819 | 727,7916092 |
| PPM1F | ENSG00000100034 | 942,4862925 |
| RPL19 | ENSG00000108298 | 926,1319047 |
| RPS3A | ENSG00000145425 | 941,220277 |
| AATK | ENSG00000181409 | 1200,069552 |
| HK2 | ENSG00000159399 | 874,2952369 |
| DYSF | ENSG00000135636 | 1267,827736 |
| SEC16A | ENSG00000148396 | 1538,77977 |
| CEACAM3 | ENSG00000170956 | 1138,613563 |
| MT-ATP6 | ENSG00000198899 | 1191,944022 |
| C6orf62 | ENSG00000112308 | 1287,112757 |
| MAN2A2 | ENSG00000196547 | 1170,5121 |
| ID2 | ENSG00000115738 | 750,9057116 |
| TRANK1 | ENSG00000168016 | 978,4916672 |
| ATP11B | ENSG00000058063 | 1603,915939 |
| PAFAH1B1 | ENSG00000007168 | 1225,068574 |
| SH3BGRL3 | ENSG00000142669 | 1012,247762 |
| CANX ENSG00000127022 | ENSG00000127022 | 1001,729692 |
| DNAJC5 | ENSG00000101152 | 998,5174004 |
| PXN | ENSG00000089159 | 1182,380327 |
| PCBP1 | ENSG00000169564 | 976,7996844 |
| BRD4 | ENSG00000141867 | 1175,4693 |
| RALGDS | ENSG00000160271 | 990,2652951 |
| CASC3 | ENSG00000108349 | 1077,986548 |
| WSB1 | ENSG00000109046 | 1368,345538 |
| MORF4L1 | ENSG00000185787 | 1172,591928 |
| NIPBL | ENSG00000164190 | 1324,211566 |
| TLR1 | ENSG00000174125 | 1174,298335 |
| MIDEAS | ENSG00000156030 | 854,5165668 |
| RPL9 | ENSG00000163682 | 1064,885816 |
| KLHL21 | ENSG00000162413 | 845,6686959 |
| ATP6V0D1 | ENSG00000159720 | 1145,880557 |
| CYRIA | ENSG00000197872 | 1205,485875 |
| SRSF3 | ENSG00000112081 | 1465,721069 |
| CNOT1 | ENSG00000125107 | 1076,236075 |
| SLC3A2 | ENSG00000168003 | 876,0537058 |

| | | |
|---|---|---|
| RAB1A | ENSG00000138069 | 1265,463646 |
| DUSP6 | ENSG00000139318 | 1279,789596 |
| HMGB1 | ENSG00000189403 | 1420,664874 |
| HNRNPH2 | ENSG00000126945 | 1305,521353 |
| KIAA0232 | ENSG00000170871 | 1192,682682 |
| FBXL5 | ENSG00000118564 | 1441,631635 |
| KATNBL1 | ENSG00000134152 | 1275,38359 |
| COPA | ENSG00000122218 | 1385,460695 |
| DOCK2 | ENSG00000134516 | 1196,328187 |
| LIPN | ENSG00000204020 | 1048,16785 |
| RPL11 | ENSG00000142676 | 929,0101039 |
| TES | ENSG00000135269 | 1021,556674 |
| IFITM1 | ENSG00000185885 | 1067,763565 |
| EIF4H | ENSG00000106682 | 1017,300846 |
| CD164 | ENSG00000135535 | 1349,627173 |
| MTMR6 | ENSG00000139505 | 1247,40791 |
| HBP1 ENSG00000105856 | ENSG00000105856 | 1013,84226 |
| RPS27A | ENSG00000143947 | 866,1143729 |
| TP53INP2 | ENSG00000078804 | 893,6235003 |
| YTHDC1 ENSG00000083896 | ENSG00000083896 | 1289,063946 |
| GOLGA7 | ENSG00000147533 | 1007,045505 |
| PHF20 | ENSG00000025293 | 1151,447779 |
| UBA52 | ENSG00000221983 | 1001,832816 |
| FHOD1 | ENSG00000135723 | 1102,4703 |
| RELB | ENSG00000104856 | 814,2716364 |
| CCDC88B | ENSG00000168071 | 1059,910334 |
| AREL1 | ENSG00000119682 | 1062,148823 |
| SF3A1 | ENSG00000099995 | 1066,704538 |
| TOP1 | ENSG00000198900 | 1274,131695 |
| SIN3A | ENSG00000169375 | 925,8023919 |
| LST1 ENSG00000204482 | ENSG00000204482 | 1076,713049 |
| SETD5 | ENSG00000168137 | 1073,651005 |
| ARHGAP25 | ENSG00000163219 | 916,1132406 |
| WAS | ENSG00000015285 | 997,8083246 |
| IFI44L | ENSG00000137959 | 661,0336988 |
| MEF2D | ENSG00000116604 | 853,8529996 |
| GCA | ENSG00000115271 | 1603,034133 |
| SMG1 | ENSG00000157106 | 1076,440293 |
| MID1IP1 | ENSG00000165175 | 716,8386149 |
| KLF3 | ENSG00000109787 | 930,0273301 |
| PATL1 | ENSG00000166889 | 1290,354676 |
| HNRNPA1 | ENSG00000135486 | 889,4831168 |
| CXCR1 | ENSG00000163464 | 854,9440077 |
| KDM5A | ENSG00000073614 | 1038,250651 |
| KBTBD2 | ENSG00000170852 | 1023,135789 |
| KDM3A | ENSG00000115548 | 1032,462091 |
| EP300 | ENSG00000100393 | 925,0888532 |
| DEDD2 | ENSG00000160570 | 787,408383 |
| WARS1 | ENSG00000140105 | 1192,227897 |
| CREB1 | ENSG00000118260 | 1070,777271 |
| IRS2 | ENSG00000185950 | 691,9908029 |
| MARCHF6 | ENSG00000145495 | 1297,288104 |
| NAGK | ENSG00000124357 | 1091,603782 |
| MLLT6 ENSG00000275023 | ENSG00000275023 | 1064,500384 |
| RAB11FIP1 | ENSG00000156675 | 867,9495185 |

| | | |
|---|---|---|
| RPLP1 | ENSG00000137818 | 738,0764544 |
| ZHX2 | ENSG00000178764 | 850,5928891 |
| RPS2 | ENSG00000140988 | 734,2538187 |
| CSK | ENSG00000103653 | 728,9404997 |
| MLF2 | ENSG00000089693 | 881,8213214 |
| SIK3 | ENSG00000160584 | 1145,891411 |
| MTF1 | ENSG00000188786 | 919,5135756 |
| PLCG2 | ENSG00000197943 | 1237,984989 |
| SRSF5 | ENSG00000100650 | 1247,572732 |
| TRAF3 | ENSG00000131323 | 868,8258555 |
| NBR1 | ENSG00000188554 | 898,7257917 |
| PSD4 | ENSG00000125637 | 905,0196686 |
| LILRB2 ENSG00000131042 | ENSG00000131042 | 1111,839434 |
| UNC13D | ENSG00000092929 | 843,899648 |
| GLT1D1 | ENSG00000151948 | 826,9969875 |
| ANKRD11 | ENSG00000167522 | 1052,797341 |
| WDFY3 | ENSG00000163625 | 1127,706497 |
| RPL7 | ENSG00000147604 | 843,4278147 |
| GCH1 | ENSG00000131979 | 1005,74729 |
| IRAK1 | ENSG00000184216 | 1110,675589 |
| CASP8 | ENSG00000064012 | 1240,39595 |
| EDEM1 | ENSG00000134109 | 1109,892309 |
| PGS1 | ENSG00000087157 | 1356,933298 |
| UBXN4 | ENSG00000144224 | 1197,365886 |
| ZNF24 | ENSG00000172466 | 995,9486326 |
| IL13RA1 | ENSG00000131724 | 1078,942402 |
| RIN3 | ENSG00000100599 | 934,3596529 |
| FGL2 | ENSG00000127951 | 1271,319474 |
| SULF2 | ENSG00000196562 | 887,4482666 |
| ST3GAL1 | ENSG00000008513 | 848,3246236 |
| GMIP | ENSG00000089639 | 946,9698087 |
| WDR82 | ENSG00000164091 | 1012,252777 |
| PLAGL2 | ENSG00000126003 | 953,1935355 |
| ENSG00000241860 | ENSG00000241860 | 1309,185944 |
| ACOT9 | ENSG00000123130 | 1270,965313 |
| RPS8 | ENSG00000142937 | 724,2368301 |
| CALM2 | ENSG00000143933 | 1148,662452 |
| IL16 | ENSG00000172349 | 602,7317207 |
| SAMD4B | ENSG00000179134 | 1015,997234 |
| HAPSTR1 | ENSG00000182831 | 1190,954901 |
| RPL15 | ENSG00000174748 | 740,8938179 |
| CASP4 | ENSG00000196954 | 1302,262709 |
| YWHAB | ENSG00000166913 | 1144,654259 |
| RPL8 | ENSG00000161016 | 714,316639 |
| HMGN2 | ENSG00000198830 | 971,692354 |
| THRAP3 | ENSG00000054118 | 977,7357653 |
| ROCK1 | ENSG00000067900 | 1294,0041 |
| DYRK1A | ENSG00000157540 | 986,6908729 |
| ZNF292 | ENSG00000188994 | 856,7002501 |
| CASS4 | ENSG00000087589 | 1159,832912 |
| ARL6IP5 | ENSG00000144746 | 769,7114639 |
| SLU7 | ENSG00000164609 | 1048,870298 |
| TBK1 | ENSG00000183735 | 1204,038918 |
| SNX20 | ENSG00000167208 | 839,6543935 |
| NDRG1 | ENSG00000104419 | 1346,599926 |

| | | |
|---|---|---|
| CALCOCO2 | ENSG00000136436 | 1134,271001 |
| ELF2 | ENSG00000109381 | 929,0813763 |
| HNRNPH1 ENSG00000169045 | ENSG00000169045 | 1102,583734 |
| NRDC | ENSG00000078618 | 1286,076921 |
| NR4A3 | ENSG00000119508 | 1206,831225 |
| NUP58 | ENSG00000139496 | 983,5699564 |
| EIF4A2 | ENSG00000156976 | 1232,567481 |
| RPS18 ENSG00000231500 | ENSG00000231500 | 719,6709139 |
| MIER1 | ENSG00000198160 | 1148,442851 |
| TAX1BP1 | ENSG00000106052 | 1202,457302 |
| MBNL1 | ENSG00000152601 | 1080,209767 |
| SMAP2 | ENSG00000084070 | 1224,332216 |
| CTNNB1 | ENSG00000168036 | 1137,47407 |
| EMP3 | ENSG00000142227 | 682,1717414 |
| RAB10 | ENSG00000084733 | 821,1974808 |
| MARCHF7 | ENSG00000136536 | 1056,688541 |
| PELATON | ENSG00000224397 | 1112,716202 |
| SOS2 | ENSG00000100485 | 1133,135821 |
| GATAD2A | ENSG00000167491 | 859,8396933 |
| BSG | ENSG00000172270 | 807,3933945 |
| RPL27A | ENSG00000166441 | 728,3051113 |
| RPS16 | ENSG00000105193 | 780,0550299 |
| BRAF | ENSG00000157764 | 1222,951566 |
| HSD17B11 | ENSG00000198189 | 988,616541 |
| EIF4EBP2 | ENSG00000148730 | 1002,265536 |
| SLC9A8 | ENSG00000197818 | 1038,187624 |
| FCGRT | ENSG00000104870 | 776,2319176 |
| ANXA5 | ENSG00000164111 | 912,348076 |
| RPL5 | ENSG00000122406 | 767,348848 |
| SVIL | ENSG00000197321 | 1006,367456 |
| CXCL1 | ENSG00000163739 | 1587,210756 |
| TMEM167B | ENSG00000215717 | 1232,128026 |
| PRPF8 ENSG00000174231 | ENSG00000174231 | 921,9488154 |
| PRKCB | ENSG00000166501 | 855,6647145 |
| NAB1 | ENSG00000138386 | 845,89121 |
| CD63 | ENSG00000135404 | 1072,013133 |
| EMD | ENSG00000102119 | 955,5898715 |
| RBM23 | ENSG00000100461 | 913,9396555 |
| PLEKHA2 | ENSG00000169499 | 1002,819855 |
| G3BP2 | ENSG00000138757 | 900,3539434 |
| CORO1C | ENSG00000110880 | 929,087826 |
| YME1L1 | ENSG00000136758 | 971,5583912 |
| IER2 | ENSG00000160888 | 692,4825075 |
| PIK3CD | ENSG00000171608 | 769,6114237 |
| SYNE2 | ENSG00000054654 | 1245,694481 |
| HCAR2 | ENSG00000182782 | 1170,713131 |
| HNRNPUL1 | ENSG00000105323 | 738,0644951 |
| RLIM | ENSG00000131263 | 1226,015246 |
| LFNG | ENSG00000106003 | 601,613791 |
| TRIM25 | ENSG00000121060 | 980,8912428 |
| NUB1 | ENSG00000013374 | 832,4971744 |
| DHRS7 | ENSG00000100612 | 959,3014769 |
| GPAT4 | ENSG00000158669 | 1206,772829 |
| ZNF710 | ENSG00000140548 | 665,2051628 |
| VCP | ENSG00000165280 | 847,6180094 |

| | | |
|---|---|---|
| FOSB | ENSG00000125740 | 1084,097381 |
| PPP4R2 | ENSG00000163605 | 1229,289405 |
| RPS25 ENSG00000118181 | ENSG00000118181 | 820,9392211 |
| FAM53C | ENSG00000120709 | 868,2388754 |
| RGS14 | ENSG00000169220 | 302,1866202 |
| SP1 | ENSG00000185591 | 878,1894319 |
| PNPLA8 | ENSG00000135241 | 1006,822191 |
| NOD2 | ENSG00000167207 | 634,0697687 |
| ARL4C | ENSG00000188042 | 602,4641058 |
| SLC44A2 | ENSG00000129353 | 955,99616 |
| AGTPBP1 | ENSG00000135049 | 1099,411492 |
| ZC3HAV1 | ENSG00000105939 | 817,4827096 |
| CRTC2 | ENSG00000160741 | 949,1046991 |
| MAPK6 | ENSG00000069956 | 873,3033084 |
| KDM3B | ENSG00000120733 | 957,6716625 |
| SH2B3 | ENSG00000111252 | 835,0811446 |
| CPPED1 | ENSG00000103381 | 1055,478678 |
| P2RY8 | ENSG00000182162 | 650,1756484 |
| TRAF1 | ENSG00000056558 | 593,675189 |
| EZR | ENSG00000092820 | 860,6378502 |
| CDKN1B | ENSG00000111276 | 901,067778 |
| BAZ2A | ENSG00000076108 | 1031,004249 |
| FAM177A1 | ENSG00000151327 | 1046,993362 |
| AFF4 | ENSG00000072364 | 1029,980193 |
| TBL1XR1 | ENSG00000177565 | 1018,667672 |
| SRPK1 | ENSG00000096063 | 1066,176536 |
| DDX60 | ENSG00000137628 | 650,3498919 |
| SEC22B | ENSG00000265808 | 997,4826921 |
| PSME1 ENSG00000092010 | ENSG00000092010 | 915,1653064 |
| NCF1C | ENSG00000165178 | 672,0099909 |
| JUND | ENSG00000130522 | 660,7223722 |
| CMIP | ENSG00000153815 | 933,7495063 |
| RPS12 | ENSG00000112306 | 629,640794 |
| JAML | ENSG00000160593 | 813,2217863 |
| RB1CC1 | ENSG00000023287 | 940,9277244 |
| ACAP2 | ENSG00000114331 | 1104,128824 |
| SESN2 ENSG00000130766 | ENSG00000130766 | 941,7887345 |
| SF1 | ENSG00000168066 | 1064,706358 |
| ITGB1 | ENSG00000150093 | 866,9030066 |
| PAPOLA | ENSG00000090060 | 1081,901055 |
| IRAG2 | ENSG00000118308 | 1101,540462 |
| NDST1 | ENSG00000070614 | 761,9093573 |
| TMEM30A | ENSG00000112697 | 1012,041084 |
| RPL7A ENSG00000148303 | ENSG00000148303 | 688,3941805 |
| SQOR | ENSG00000137767 | 814,9054073 |
| NAPA | ENSG00000105402 | 1078,521622 |
| OTUD5 | ENSG00000068308 | 933,6825535 |
| ANXA11 | ENSG00000122359 | 905,7888817 |
| PPCDC | ENSG00000138621 | 810,6468591 |
| FCN1 | ENSG00000085265 | 653,2264594 |
| PLEKHO1 | ENSG00000023902 | 813,1830037 |
| PUM2 | ENSG00000055917 | 934,3959477 |
| QPCT | ENSG00000115828 | 948,183363 |
| CDA | ENSG00000158825 | 612,5106824 |
| UBR2 | ENSG00000024048 | 836,6174835 |

| | | |
|---|---|---|
| ETS1 | ENSG00000134954 | 716,8140468 |
| C16orf54 | ENSG00000185905 | 907,3997565 |
| ANXA7 | ENSG00000138279 | 746,1277273 |
| JMJD6 | ENSG00000070495 | 1083,135858 |
| NAA50 | ENSG00000121579 | 795,3119052 |
| FGD4 | ENSG00000139132 | 1161,046982 |
| WNK1 | ENSG00000060237 | 794,469594 |
| SEL1L | ENSG00000071537 | 802,518754 |
| FKBP8 | ENSG00000105701 | 694,7105337 |
| RPS14 | ENSG00000164587 | 603,7513959 |
| SEPTIN7 | ENSG00000122545 | 1031,072516 |
| RPS20 | ENSG00000008988 | 718,1133083 |
| OSBPL2 | ENSG00000130703 | 804,3058046 |
| CYB5R4 | ENSG00000065615 | 928,5340646 |
| CACUL1 | ENSG00000151893 | 982,2931524 |
| GGA1 | ENSG00000100083 | 858,7365623 |
| RPL28 | ENSG00000108107 | 703,491011 |
| SPPL2A | ENSG00000138600 | 765,2246867 |
| ITSN2 | ENSG00000198399 | 975,4956856 |
| RNF130 | ENSG00000113269 | 858,3227163 |
| TNFRSF14 ENSG00000157873 | ENSG00000157873 | 787,5355751 |
| IRF7 ENSG00000185507 | ENSG00000185507 | 642,0022066 |
| CNPY3 | ENSG00000137161 | 741,9783074 |
| ARIH1 | ENSG00000166233 | 1034,304509 |
| PPP2CA | ENSG00000113575 | 890,2072559 |
| MAEA | ENSG00000090316 | 897,9673666 |
| KCNJ2 | ENSG00000123700 | 589,3345019 |
| CEP170 ENSG00000143702 | ENSG00000143702 | 1043,92218 |
| CALM1 | ENSG00000198668 | 953,4069041 |
| ZFAND6 | ENSG00000086666 | 999,8727942 |
| SELENOT | ENSG00000198843 | 911,7077182 |
| RPS24 | ENSG00000138326 | 801,1974767 |
| SLK | ENSG00000065613 | 758,8931452 |
| USP4 | ENSG00000114316 | 927,7900137 |
| GPR108 | ENSG00000125734 | 789,99148 |
| UBXN2B | ENSG00000215114 | 1127,766665 |
| KDM2A | ENSG00000173120 | 751,6486933 |
| UBAP1 | ENSG00000165006 | 1179,409516 |
| SAR1A | ENSG00000079332 | 860,3065478 |
| ITGAM | ENSG00000169896 | 1125,907917 |
| TNFSF14 | ENSG00000125735 | 725,8431461 |
| RC3H1 | ENSG00000135870 | 859,5192066 |
| XRN1 | ENSG00000114127 | 934,2092848 |
| UBA1 | ENSG00000130985 | 746,6964967 |
| CREBRF | ENSG00000164463 | 798,3576081 |
| GPATCH2L | ENSG00000089916 | 904,6641322 |
| CRISPLD2 | ENSG00000103196 | 547,6607705 |
| RPLP0 | ENSG00000089157 | 597,686939 |
| BLTP1 | ENSG00000138688 | 1066,606692 |
| S100A12 | ENSG00000163221 | 814,3095646 |
| PBX2 ENSG00000204304 | ENSG00000204304 | 853,6560992 |
| MAST3 | ENSG00000099308 | 635,3075755 |
| ZBP1 | ENSG00000124256 | 745,9434926 |
| ANXA1 | ENSG00000135046 | 689,1044668 |
| MAML1 ENSG00000161021 | ENSG00000161021 | 623,2374089 |

| | | |
|---|---|---|
| PIK3CG | ENSG00000105851 | 953,494065 |
| NLRC5 | ENSG00000140853 | 718,9666926 |
| SUB1 | ENSG00000113387 | 1042,108385 |
| POLR2A ENSG00000181222 | ENSG00000181222 | 746,7341531 |
| RAB5B | ENSG00000111540 | 697,1247391 |
| CMTM2 | ENSG00000140932 | 836,2497991 |
| GLIPR2 | ENSG00000122694 | 835,592285 |
| CIB1 | ENSG00000185043 | 952,0647715 |
| PML | ENSG00000140464 | 697,0463034 |
| TTPAL | ENSG00000124120 | 1040,279744 |
| RPL23 | ENSG00000125691 | 719,6103773 |
| VNN3P | ENSG00000093134 | 1042,333142 |
| MTPN | ENSG00000105887 | 909,3727934 |
| RCSD1 | ENSG00000198771 | 587,8408084 |
| DNTTIP2 | ENSG00000067334 | 1231,865214 |
| SEPTIN2 | ENSG00000168385 | 922,5666586 |
| PGD | ENSG00000142657 | 945,9154306 |
| BRI3 | ENSG00000164713 | 817,9009852 |
| RMND5A | ENSG00000153561 | 654,9293418 |
| RPL31 | ENSG00000071082 | 620,665227 |
| TM9SF4 | ENSG00000101337 | 572,7001738 |
| RAB11FIP4 | ENSG00000131242 | 646,8998733 |
| PHF1 ENSG00000112511 | ENSG00000112511 | 832,1248 |
| NDEL1 | ENSG00000166579 | 1026,982936 |
| FURIN | ENSG00000140564 | 671,7503484 |
| MSRB1 | ENSG00000198736 | 652,143469 |
| FAU | ENSG00000149806 | 715,3645236 |
| CHST11 | ENSG00000171310 | 778,425283 |
| BNIP2 | ENSG00000140299 | 1055,336992 |
| RPL34 | ENSG00000109475 | 669,6226019 |
| SELENOK | ENSG00000113811 | 1011,351533 |
| ULK1 | ENSG00000177169 | 625,7137789 |
| PRKD2 | ENSG00000105287 | 839,212322 |
| SPOCK2 | ENSG00000107742 | 573,2666886 |
| OGFRL1 | ENSG00000119900 | 757,5119784 |
| SDE2 | ENSG00000143751 | 797,1465652 |
| TIPARP | ENSG00000163659 | 875,7905195 |
| LIMD2 | ENSG00000136490 | 546,2110781 |
| XBP1 | ENSG00000100219 | 732,0448444 |
| SHISA5 | ENSG00000164054 | 785,0960721 |
| LAP3 | ENSG00000002549 | 668,3844733 |
| H2BC21 | ENSG00000184678 | 548,6404121 |
| SKAP2 | ENSG00000005020 | 850,2055074 |
| GUK1 | ENSG00000143774 | 643,7681583 |
| GAPT | ENSG00000175857 | 761,1278293 |
| MMP9 | ENSG00000100985 | 599,4673845 |
| VHL | ENSG00000134086 | 795,9810811 |
| CWC25 ENSG00000273559 | ENSG00000273559 | 896,4613488 |
| CD300E | ENSG00000186407 | 638,1956186 |
| RPL32 | ENSG00000144713 | 631,7118017 |
| NLRP3 | ENSG00000162711 | 800,9745221 |
| CELF2 | ENSG00000048740 | 955,5862766 |
| ST3GAL2 | ENSG00000157350 | 490,5818433 |
| KIF5B | ENSG00000170759 | 892,4351344 |
| ANKRD13D | ENSG00000172932 | 663,5430739 |

| | | |
|---|---|---|
| ARHGAP4 | ENSG00000089820 | 669,7401559 |
| UBR5 | ENSG00000104517 | 923,6204437 |
| STK38L | ENSG00000211455 | 1058,663017 |
| ADAM10 | ENSG00000137845 | 841,2712932 |
| TMEM59 | ENSG00000116209 | 809,567988 |
| BEST1 | ENSG00000167995 | 700,6647311 |
| RNF11 | ENSG00000123091 | 934,8462693 |
| EIF4B | ENSG00000063046 | 663,9432726 |
| CHMP2A | ENSG00000130724 | 766,2995718 |
| HSP90AB1 | ENSG00000096384 | 684,9203389 |
| USP7 | ENSG00000187555 | 827,8179129 |
| TOR1AIP1 | ENSG00000143337 | 1071,344868 |
| DENND4A | ENSG00000174485 | 694,9039363 |
| KDM7A | ENSG00000006459 | 821,9639405 |
| SPEN | ENSG00000065526 | 604,5370321 |
| KLF13 ENSG00000169926 | ENSG00000169926 | 519,7240967 |
| EVI2A | ENSG00000126860 | 761,537349 |
| SRPRA | ENSG00000182934 | 736,6261128 |
| PDXK | ENSG00000160209 | 624,9994985 |
| TOP2B | ENSG00000077097 | 773,1466677 |
| CSNK1A1 | ENSG00000113712 | 833,9233834 |
| MT-ND3 | ENSG00000198840 | 754,9761074 |
| ABCG1 | ENSG00000160179 | 1040,830008 |
| GSN | ENSG00000148180 | 552,9455504 |
| NUFIP2 | ENSG00000108256 | 952,5544356 |
| PBXIP1 | ENSG00000163346 | 607,5279133 |
| NCOR2 | ENSG00000196498 | 747,3123406 |
| TMEM50A | ENSG00000183726 | 677,1656289 |
| RP2 | ENSG00000102218 | 887,0051894 |
| RHBDF2 | ENSG00000129667 | 651,263331 |
| CDV3 | ENSG00000091527 | 776,8960483 |
| OAZ2 | ENSG00000180304 | 794,3263537 |
| UPP1 | ENSG00000183696 | 1079,075318 |
| AKAP13 | ENSG00000170776 | 725,8992337 |
| MDM2 | ENSG00000135679 | 607,6108034 |
| TIMP2 | ENSG00000035862 | 716,6586427 |
| CTDSP2 | ENSG00000175215 | 588,9814669 |
| RLF | ENSG00000117000 | 851,8268247 |
| SH2D3C | ENSG00000095370 | 685,2873105 |
| ACSL4 | ENSG00000068366 | 917,2656843 |
| KHDRBS1 | ENSG00000121774 | 715,796972 |
| BAZ2B | ENSG00000123636 | 826,3322235 |
| KPNA4 | ENSG00000186432 | 869,2788028 |
| ARHGEF1 | ENSG00000076928 | 675,7468838 |
| PIAS1 | ENSG00000033800 | 898,1392244 |
| MAPK14 | ENSG00000112062 | 744,3167604 |
| EPSTI1 | ENSG00000133106 | 466,5521804 |
| PARP8 | ENSG00000151883 | 988,9804086 |
| PIK3R1 | ENSG00000145675 | 647,2129636 |
| NLRP1 | ENSG00000091592 | 783,7819461 |
| RACK1 | ENSG00000204628 | 562,7875698 |
| NFAT5 | ENSG00000102908 | 958,701142 |
| RPSA | ENSG00000168028 | 525,0578244 |
| SH3KBP1 | ENSG00000147010 | 832,2769979 |
| CHMP1B | ENSG00000255112 | 733,7598401 |

| | | |
|---|---|---|
| PDIA3 | ENSG00000167004 | 842,8151045 |
| TMEM140 | ENSG00000146859 | 653,9163467 |
| MECP2 | ENSG00000169057 | 835,7697619 |
| VPS4B | ENSG00000119541 | 640,7025204 |
| RGS19 | ENSG00000171700 | 568,4552617 |
| ETV6 | ENSG00000139083 | 857,0050501 |
| PAFAH1B2 | ENSG00000168092 | 750,708055 |
| SRSF2 | ENSG00000161547 | 933,5726456 |
| INAFM2 | ENSG00000259330 | 666,157216 |
| OSGIN2 | ENSG00000164823 | 763,5981272 |
| AP2B1 | ENSG00000006125 | 686,5709888 |
| NXF1 | ENSG00000162231 | 689,1078993 |
| AHNAK | ENSG00000124942 | 407,1127433 |
| RNF10 | ENSG00000022840 | 826,7866771 |
| MCEMP1 | ENSG00000183019 | 926,5542573 |
| METRNL ENSG00000176845 | ENSG00000176845 | 685,1393368 |
| HSPA8 | ENSG00000109971 | 611,5494931 |
| ACAA1 | ENSG00000060971 | 592,7637178 |
| XRCC5 | ENSG00000079246 | 767,3658619 |
| RCOR1 | ENSG00000089902 | 839,8753401 |
| APMAP | ENSG00000101474 | 485,1206066 |
| PPP4C | ENSG00000149923 | 672,7436509 |
| STXBP3 | ENSG00000116266 | 762,6423906 |
| STK24 | ENSG00000102572 | 872,9691954 |
| RSRP1 | ENSG00000117616 | 690,3224223 |
| PARP4 | ENSG00000102699 | 728,8681601 |
| CDK17 | ENSG00000059758 | 846,578828 |
| UBE2R2 | ENSG00000107341 | 871,7567022 |
| AKAP17A | ENSG00000197976 | 763,2355148 |
| KLF2 | ENSG00000127528 | 742,1035004 |
| GOLGB1 | ENSG00000173230 | 873,6367477 |
| OSTF1 | ENSG00000134996 | 691,8707237 |
| MSL2 | ENSG00000174579 | 567,7761586 |
| OGT | ENSG00000147162 | 761,1711885 |
| RELT | ENSG00000054967 | 841,4520884 |
| NCF4 ENSG00000100365 | ENSG00000100365 | 793,1693763 |
| HSP90B1 | ENSG00000166598 | 829,6327003 |
| GZF1 | ENSG00000125812 | 917,6004983 |
| TMF1 | ENSG00000144747 | 683,1704773 |
| HPCAL1 | ENSG00000115756 | 811,5133281 |
| RAB27A | ENSG00000069974 | 801,330174 |
| AHCTF1 | ENSG00000153207 | 628,2183932 |
| FLOT2 | ENSG00000132589 | 492,2908657 |
| PIK3IP1 | ENSG00000100100 | 571,9944113 |
| HUWE1 | ENSG00000086758 | 659,6993511 |
| CPEB2 | ENSG00000137449 | 797,2232763 |
| RPL6 | ENSG00000089009 | 551,4113144 |
| OAS1 | ENSG00000089127 | 337,6135638 |
| ABHD5 | ENSG00000011198 | 832,9768573 |
| MIR223HG | ENSG00000274536 | 907,1378146 |
| TSPAN14 | ENSG00000108219 | 626,3495381 |
| PLCB2 | ENSG00000137841 | 773,8013083 |
| NUP153 | ENSG00000124789 | 798,5842198 |
| RASGRP4 | ENSG00000171777 | 677,1177229 |
| CIR1 | ENSG00000138433 | 712,6924531 |

| | | |
|---|---|---|
| UBQLN1 | ENSG00000135018 | 789,7887556 |
| TACC3 | ENSG00000013810 | 823,5897802 |
| RPL37 | ENSG00000145592 | 605,0997428 |
| WAPL | ENSG00000062650 | 740,7608306 |
| KDM1B | ENSG00000165097 | 567,2417132 |
| ANKRD44 | ENSG00000065413 | 763,193931 |
| MAT2B | ENSG00000038274 | 603,3662085 |
| PRRC2A ENSG00000204469 | ENSG00000204469 | 591,1818474 |
| MORC3 | ENSG00000159256 | 758,9855645 |
| SAMHD1 | ENSG00000101347 | 610,648286 |
| PPP1R12A | ENSG00000058272 | 873,7542556 |
| USP34 | ENSG00000115464 | 756,2398409 |
| RPS7 | ENSG00000171863 | 549,4385203 |
| ESYT2 | ENSG00000117868 | 556,3657615 |
| OS9 | ENSG00000135506 | 746,8856634 |
| SPTY2D1 | ENSG00000179119 | 641,8014407 |
| RAB11A | ENSG00000103769 | 577,4086875 |
| RPS19 | ENSG00000105372 | 480,5442 |
| TNRC18 | ENSG00000182095 | 471,4802371 |
| POR | ENSG00000127948 | 795,835211 |
| RBM5 | ENSG00000003756 | 711,9486474 |
| NCOA2 | ENSG00000140396 | 765,8464853 |
| CNOT6L | ENSG00000138767 | 661,2744256 |
| KDM6A | ENSG00000147050 | 750,9637444 |
| CD8A | ENSG00000153563 | 330,5707451 |
| RPL12 | ENSG00000197958 | 536,6692853 |
| HDGF | ENSG00000143321 | 584,6355644 |
| SYK | ENSG00000165025 | 560,5510996 |
| SFT2D1 | ENSG00000198818 | 673,7174369 |
| USP3 | ENSG00000140455 | 613,3958262 |
| SEPTIN9 ENSG00000184640 | ENSG00000184640 | 409,2026012 |
| CARD19 | ENSG00000165233 | 782,9228134 |
| TDRD7 | ENSG00000196116 | 665,9658605 |
| TSC22D4 | ENSG00000166925 | 596,6581945 |
| RPL39 | ENSG00000198918 | 577,7179647 |
| LY6E ENSG00000160932 | ENSG00000160932 | 313,8088377 |
| AKIRIN1 | ENSG00000174574 | 684,3872829 |
| AGO4 | ENSG00000134698 | 587,1113189 |
| RNF145 | ENSG00000145860 | 725,9902606 |
| RPL27 | ENSG00000131469 | 506,6975121 |
| NUAK2 | ENSG00000163545 | 610,2251965 |
| BCOR | ENSG00000183337 | 496,6674963 |
| TSG101 | ENSG00000074319 | 738,7201547 |
| RPL30 | ENSG00000156482 | 580,0243481 |
| APBB1IP | ENSG00000077420 | 689,6764245 |
| ARPP19 | ENSG00000128989 | 631,7308358 |
| DOT1L | ENSG00000104885 | 757,3664888 |
| SEMA6B | ENSG00000167680 | 705,5004561 |
| TAOK1 | ENSG00000160551 | 662,1419644 |
| PAIP2 | ENSG00000120727 | 711,9107543 |
| BIRC6 | ENSG00000115760 | 719,0654691 |
| F11R | ENSG00000158769 | 602,7837951 |
| CD37 | ENSG00000104894 | 685,4501319 |
| NCSTN | ENSG00000162736 | 807,3437617 |
| ZBTB1 | ENSG00000126804 | 524,6024912 |

| | | |
|---|---|---|
| LAMP1 | ENSG00000185896 | 718,4222439 |
| CTSB ENSG00000164733 | ENSG00000164733 | 686,0033543 |
| TPR | ENSG00000047410 | 776,7424452 |
| BLTP3B | ENSG00000111647 | 854,6744665 |
| PARP12 | ENSG00000059378 | 510,6287804 |
| ARHGAP9 | ENSG00000123329 | 683,396409 |
| CHI3L1 | ENSG00000133048 | 460,7066937 |
| HMOX1 | ENSG00000100292 | 662,0253414 |
| ATP6V0E1 | ENSG00000113732 | 648,8741222 |
| PLPPR2 | ENSG00000105520 | 681,1693424 |
| TPP1 | ENSG00000166340 | 796,4042395 |
| KIF1B | ENSG00000054523 | 791,3071573 |
| DNAJB1 | ENSG00000132002 | 804,2373469 |
| STX4 | ENSG00000103496 | 813,966677 |
| DUSP16 ENSG00000111266 | ENSG00000111266 | 697,3409999 |
| PLEKHG3 | ENSG00000126822 | 381,2681839 |
| HPSE | ENSG00000173083 | 936,4105086 |
| PKM | ENSG00000067225 | 587,8383816 |
| NAP1L1 | ENSG00000187109 | 648,0648678 |
| CCRL2 | ENSG00000121797 | 742,964813 |
| PSME4 | ENSG00000068878 | 718,3860361 |
| GDI1 | ENSG00000203879 | 733,1439366 |
| TET3 | ENSG00000187605 | 669,4571156 |
| TP53INP1 | ENSG00000164938 | 625,4003318 |
| ABLIM1 | ENSG00000099204 | 467,1480301 |
| SNRNP200 | ENSG00000144028 | 568,149456 |
| SMARCA5 | ENSG00000153147 | 737,5291827 |
| RPL18A | ENSG00000105640 | 387,3768805 |
| SYNJ1 | ENSG00000159082 | 835,3726259 |
| RPLP2 | ENSG00000177600 | 421,2821126 |
| RBM25 | ENSG00000119707 | 925,2410909 |
| CIRBP | ENSG00000099622 | 615,1436553 |
| RUNX3 | ENSG00000020633 | 439,3581512 |
| COX4I1 | ENSG00000131143 | 561,4475788 |
| PGAP6 | ENSG00000129925 | 471,0325033 |
| KCNAB2 | ENSG00000069424 | 554,4773689 |
| CR1 | ENSG00000203710 | 758,6558571 |
| CALR | ENSG00000179218 | 510,809853 |
| ARID4A | ENSG00000032219 | 610,9344713 |
| ST8SIA4 | ENSG00000113532 | 563,9100098 |
| AP5B1 | ENSG00000254470 | 562,4228034 |
| DOCK4 | ENSG00000128512 | 714,6334452 |
| FOXN3 | ENSG00000053254 | 666,9021196 |
| ACTN4 ENSG00000130402 | ENSG00000130402 | 608,5584852 |
| GTPBP2 | ENSG00000172432 | 666,2880841 |
| PAG1 | ENSG00000076641 | 830,5785248 |
| ALPL | ENSG00000162551 | 693,5269012 |
| MDM4 | ENSG00000198625 | 660,117229 |
| EWSR1 | ENSG00000182944 | 679,7703184 |
| MIR22HG ENSG00000186594 | ENSG00000186594 | 864,0298545 |
| PHF11 | ENSG00000136147 | 703,0837421 |
| MKRN1 | ENSG00000133606 | 631,9860989 |
| MIR29B2CHG | ENSG00000203709 | 613,415263 |
| FAM91A1 | ENSG00000176853 | 751,5634965 |
| TRAF3IP3 | ENSG00000009790 | 828,6990758 |

| | | |
|---|---|---|
| CALM3 | ENSG00000160014 | 568,1248125 |
| PXK | ENSG00000168297 | 675,7735199 |
| TMEM41B | ENSG00000166471 | 818,5091135 |
| PAN3 | ENSG00000152520 | 769,5158363 |
| CCNDBP1 | ENSG00000166946 | 691,1953372 |
| ATP6AP2 | ENSG00000182220 | 733,9452139 |
| MGAT1 | ENSG00000131446 | 583,2088116 |
| ADAM17 | ENSG00000151694 | 678,2702078 |
| ARFGEF1 | ENSG00000066777 | 695,4533704 |
| PARP10 | ENSG00000178685 | 513,7356705 |
| ZDHHC5 | ENSG00000156599 | 579,7317765 |
| ZFC3H1 | ENSG00000133858 | 755,0252352 |
| IL7R | ENSG00000168685 | 487,4534968 |
| IK | ENSG00000113141 | 707,9969415 |
| PLBD1 | ENSG00000121316 | 754,0257986 |
| ACOX1 | ENSG00000161533 | 528,7714973 |
| HMGN1 | ENSG00000205581 | 704,6300053 |
| NT5C2 | ENSG00000076685 | 808,172891 |
| TXNDC11 | ENSG00000153066 | 736,7822582 |
| COPB1 | ENSG00000129083 | 664,3133476 |
| WIPF2 | ENSG00000171475 | 518,864778 |
| P4HB | ENSG00000185624 | 501,9414193 |
| MYLIP | ENSG00000007944 | 831,5662662 |
| DCUN1D3 | ENSG00000188215 | 1045,058945 |
| IQGAP2 | ENSG00000145703 | 752,8258924 |
| ARF6 | ENSG00000165527 | 530,2017238 |
| MTMR14 | ENSG00000163719 | 513,9383551 |
| MTCH1 | ENSG00000137409 | 598,0104529 |
| FMR1 | ENSG00000102081 | 657,3853211 |
| RBM47 | ENSG00000163694 | 747,139904 |
| TLR6 | ENSG00000174130 | 573,2227666 |
| STXBP2 | ENSG00000076944 | 694,2308727 |
| ENC1 | ENSG00000171617 | 252,303813 |
| MCTP2 | ENSG00000140563 | 744,6287967 |
| GPBP1L1 | ENSG00000159592 | 581,4486232 |
| HERPUD2 | ENSG00000122557 | 596,0150212 |
| SLCO3A1 | ENSG00000176463 | 606,9878553 |
| RPL24 | ENSG00000114391 | 513,9284116 |
| RAB2A | ENSG00000104388 | 637,637712 |
| VEZF1 | ENSG00000136451 | 645,757909 |
| RAP2B | ENSG00000181467 | 325,841357 |
| ZC3H18 | ENSG00000158545 | 634,8902318 |
| CPQ | ENSG00000104324 | 647,3163051 |
| TMSB10 | ENSG00000034510 | 610,4126008 |
| RHOQ | ENSG00000119729 | 534,6749065 |
| UBN1 | ENSG00000118900 | 462,9377539 |
| CPSF7 | ENSG00000149532 | 590,4006313 |
| MRFAP1 | ENSG00000179010 | 512,9569009 |
| MARK2 | ENSG00000072518 | 386,2495501 |
| GPX3 | ENSG00000211445 | 366,4673233 |
| ARCN1 | ENSG00000095139 | 554,5699337 |
| RBM3 | ENSG00000102317 | 604,1722436 |
| SSU72 | ENSG00000160075 | 483,0063855 |
| SLC38A1 | ENSG00000111371 | 421,17252 |
| HERPUD1 | ENSG00000051108 | 469,6902833 |

| | | |
|---|---|---|
| KHNYN | ENSG00000100441 | 548,9990067 |
| INPP4A | ENSG00000040933 | 631,2189165 |
| NORAD | ENSG00000260032 | 561,3738469 |
| STRN4 | ENSG00000090372 | 508,7723757 |
| IL17RA | ENSG00000177663 | 475,4031348 |
| SERTAD2 | ENSG00000179833 | 709,1408587 |
| ITCH | ENSG00000078747 | 658,0379473 |
| GALNT1 | ENSG00000141429 | 645,7427627 |
| UBE2L6 | ENSG00000156587 | 526,6421127 |
| CHMP5 | ENSG00000086065 | 649,3510301 |
| ZNF641 | ENSG00000167528 | 450,5593328 |
| TMED5 | ENSG00000117500 | 855,1451717 |
| HERC3 | ENSG00000138641 | 746,8227035 |
| UBALD2 | ENSG00000185262 | 591,1400554 |
| CAB39 | ENSG00000135932 | 694,9992551 |
| CYSTM1 | ENSG00000120306 | 650,2262435 |
| FBXO34 | ENSG00000178974 | 683,1829053 |
| E2F3 | ENSG00000112242 | 670,150904 |
| KMT2D | ENSG00000167548 | 527,2401021 |
| PCYT1A | ENSG00000161217 | 672,2267885 |
| SIRPB1 | ENSG00000101307 | 717,5333381 |
| RBM22 | ENSG00000086589 | 614,0544925 |
| PFDN5 | ENSG00000123349 | 633,6610677 |
| IPO7 | ENSG00000205339 | 633,1260148 |
| RGL2 ENSG00000237441 | ENSG00000237441 | 481,7015526 |
| GATAD2B ENSG00000143614 | ENSG00000143614 | 719,5425387 |
| CRK | ENSG00000167193 | 583,292919 |
| RNF168 | ENSG00000163961 | 594,7744865 |
| C1orf43 | ENSG00000143612 | 519,802118 |
| HNRNPH3 | ENSG00000096746 | 684,3521326 |
| RPL35A | ENSG00000182899 | 487,2147268 |
| NT5C3A | ENSG00000122643 | 421,9988045 |
| BRWD3 | ENSG00000165288 | 667,9152071 |
| MORF4L2 | ENSG00000123562 | 765,9533657 |
| YBX1 | ENSG00000065978 | 514,2729849 |
| ABI1 | ENSG00000136754 | 597,8485597 |
| SYF2 | ENSG00000117614 | 548,027298 |
| PDE7A | ENSG00000205268 | 549,7482911 |
| NR1H2 | ENSG00000131408 | 418,7396525 |
| OGFR | ENSG00000060491 | 501,8634924 |
| RIF1 | ENSG00000080345 | 651,2020779 |
| BTN2A1 | ENSG00000112763 | 572,6954503 |
| KDM5B | ENSG00000117139 | 671,8676917 |
| PPIG | ENSG00000138398 | 622,2708098 |
| RPS29 | ENSG00000213741 | 411,1939352 |
| GIMAP4 | ENSG00000133574 | 434,6466137 |
| ATP6AP1 | ENSG00000071553 | 544,6480203 |
| FYN | ENSG00000010810 | 359,4082823 |
| GTF2A1 | ENSG00000165417 | 531,3739028 |
| LPIN2 | ENSG00000101577 | 550,01438 |
| TNKS2 | ENSG00000107854 | 569,1224728 |
| HLA-DPA1 ENSG00000231389 | ENSG00000231389 | 443,1225107 |
| ZNF207 | ENSG00000010244 | 635,4314006 |
| TGFBR1 | ENSG00000106799 | 439,649776 |
| TFRC | ENSG00000072274 | 445,7723021 |

| | | |
|---|---|---|
| TRIM8 | ENSG00000171206 | 342,3528187 |
| NCOA1 | ENSG00000084676 | 759,3630033 |
| TMX4 | ENSG00000125827 | 626,4779062 |
| TMCC1 | ENSG00000172765 | 521,8863701 |
| SSR2 | ENSG00000163479 | 715,5385111 |
| PTGES3 | ENSG00000110958 | 528,160244 |
| IPMK | ENSG00000151151 | 662,4249953 |
| PRDM2 | ENSG00000116731 | 460,3455613 |
| TUBA4A | ENSG00000127824 | 617,4051207 |
| FBXO38 | ENSG00000145868 | 698,8286018 |
| PPP1R10 ENSG00000204569 | ENSG00000204569 | 621,9374882 |
| PROK2 | ENSG00000163421 | 593,1356449 |
| ATF4 | ENSG00000128272 | 592,0811558 |
| LAMTOR3 | ENSG00000109270 | 605,3508952 |
| CTDSP1 | ENSG00000144579 | 414,3111061 |
| NRBF2 | ENSG00000148572 | 553,2518274 |
| ENSG00000225889 | ENSG00000225889 | 633,7564547 |
| SSH1 | ENSG00000084112 | 515,0447054 |
| TERF2IP | ENSG00000166848 | 485,7888091 |
| TGIF1 | ENSG00000177426 | 490,8239978 |
| PPP4R3A | ENSG00000100796 | 516,2926663 |
| RPS13 | ENSG00000110700 | 487,1807461 |
| KIF13A | ENSG00000137177 | 482,973019 |
| XAF1 | ENSG00000132530 | 335,0103519 |
| ZFAND3 | ENSG00000156639 | 551,9437839 |
| CHURC1 | ENSG00000258289 | 463,1118938 |
| TRPC4AP | ENSG00000100991 | 554,2385389 |
| SH2B2 | ENSG00000160999 | 505,9072409 |
| PNRC2 | ENSG00000189266 | 588,7892309 |
| SH3GLB1 | ENSG00000097033 | 581,0126607 |
| TBC1D2B | ENSG00000167202 | 300,8280465 |
| GAK | ENSG00000178950 | 542,1684842 |
| N4BP2L2 | ENSG00000244754 | 610,0154476 |
| TM9SF2 | ENSG00000125304 | 574,5859919 |
| MEGF9 | ENSG00000106780 | 461,6429591 |
| TMCO3 | ENSG00000150403 | 620,2289261 |
| RPL18 | ENSG00000063177 | 408,7155465 |
| CRYBG1 | ENSG00000112297 | 431,3538542 |
| ATG3 | ENSG00000144848 | 604,8974026 |
| SREBF1 | ENSG00000072310 | 579,4384664 |
| MCOLN1 | ENSG00000090674 | 824,3047109 |
| ZFX | ENSG00000005889 | 533,7182178 |
| SAMD8 | ENSG00000156671 | 655,6293893 |
| XPO1 | ENSG00000082898 | 326,1355376 |
| SCYL2 | ENSG00000136021 | 650,8704276 |
| AK2 | ENSG00000004455 | 600,450049 |
| KPNA1 | ENSG00000114030 | 598,6278003 |
| RAB5C | ENSG00000108774 | 494,1086391 |
| ATP6V1E1 | ENSG00000131100 | 604,4038408 |
| CHMP2B | ENSG00000083937 | 627,5387108 |
| SMU1 | ENSG00000122692 | 611,8297332 |
| GPR160 | ENSG00000173890 | 677,3100856 |
| CCL3L3 ENSG00000276085 | ENSG00000276085 | 353,0375992 |
| TLE4 | ENSG00000106829 | 588,9104179 |
| PNPLA1 | ENSG00000180316 | 531,532583 |

| | | |
|---|---|---|
| NEK7 | ENSG00000151414 | 569,3566164 |
| UBL3 | ENSG00000122042 | 560,2470086 |
| HAL | ENSG00000084110 | 506,7879621 |
| SESN3 | ENSG00000149212 | 453,5108911 |
| ZNF592 | ENSG00000166716 | 476,4767793 |
| USF1 | ENSG00000158773 | 521,1615983 |
| SPOPL | ENSG00000144228 | 669,5677866 |
| PLEKHF2 | ENSG00000175895 | 517,9329517 |
| ABCF1 ENSG00000204574 | ENSG00000204574 | 492,7643987 |
| ZFYVE16 | ENSG00000039319 | 758,1983449 |
| NCL | ENSG00000115053 | 494,4631579 |
| LPGAT1 | ENSG00000123684 | 655,5218612 |
| HCAR3 | ENSG00000255398 | 618,5206375 |
| RPL14 | ENSG00000188846 | 446,5790172 |
| PHIP | ENSG00000146247 | 668,2549424 |
| IRF2 | ENSG00000168310 | 512,6339256 |
| PRDM1 | ENSG00000057657 | 349,1864564 |
| ARPC4 | ENSG00000241553 | 462,3262422 |
| ANKRD33B | ENSG00000164236 | 392,4251512 |
| PPP6C | ENSG00000119414 | 531,02106 |
| RPL10A | ENSG00000198755 | 416,1370836 |
| ATOSB | ENSG00000005238 | 649,2736184 |
| ARFIP1 | ENSG00000164144 | 590,292526 |
| EPB41 | ENSG00000159023 | 554,081052 |
| PLEKHM1 ENSG00000225190 | ENSG00000225190 | 492,4869421 |
| EIF5A | ENSG00000132507 | 417,2780507 |
| LTB ENSG00000227507 | ENSG00000227507 | 349,049752 |
| FMNL3 | ENSG00000161791 | 412,43233 |
| CAST | ENSG00000153113 | 742,9405582 |
| SEC24A | ENSG00000113615 | 737,7729508 |
| LENG8 ENSG00000167615 | ENSG00000167615 | 535,9063573 |
| CSF2RA | ENSG00000198223 | 589,4368955 |
| STARD7 | ENSG00000084090 | 356,4575155 |
| NOP10 | ENSG00000182117 | 585,6866538 |
| SMAD3 | ENSG00000166949 | 347,5388834 |
| DENND4B | ENSG00000198837 | 595,4970653 |
| BAG6 ENSG00000204463 | ENSG00000204463 | 483,2125247 |
| NACA | ENSG00000196531 | 562,9936007 |
| RBMS1 | ENSG00000153250 | 511,0762661 |
| RALGAPA1 | ENSG00000174373 | 508,1778573 |
| MAN1A1 | ENSG00000111885 | 538,2023245 |
| MAGT1 | ENSG00000102158 | 565,8315768 |
| NCF1B | ENSG00000182487 | 459,6975674 |
| ESCO1 | ENSG00000141446 | 531,8004591 |
| ENTPD4 | ENSG00000197217 | 621,6758583 |
| ZZEF1 | ENSG00000074755 | 549,2165325 |
| PHOSPHO1 | ENSG00000173868 | 317,2823944 |
| PSMB9 ENSG00000240065 | ENSG00000240065 | 391,1235171 |
| MVP | ENSG00000013364 | 484,4815975 |
| FBXO33 | ENSG00000165355 | 434,7234281 |
| ARFGAP3 | ENSG00000242247 | 633,6335071 |
| TOX4 | ENSG00000092203 | 566,2637827 |
| RAB3D | ENSG00000105514 | 479,1972691 |
| ENSG00000188897 | ENSG00000188897 | 497,3961397 |
| TOR1AIP2 | ENSG00000169905 | 655,3145216 |

| | | |
|---|---|---|
| PRCP | ENSG00000137509 | 530,8385098 |
| BROX | ENSG00000162819 | 513,1840366 |
| RPL37A | ENSG00000197756 | 401,3639129 |
| DIP2B | ENSG00000066084 | 601,7281643 |
| SH3BGRL | ENSG00000131171 | 735,0026077 |
| PNISR | ENSG00000132424 | 623,9677344 |
| TMEM259 | ENSG00000182087 | 477,0820737 |
| KDM5C | ENSG00000126012 | 462,3307723 |
| ZC3H13 | ENSG00000123200 | 625,5910348 |
| BCLAF1 | ENSG00000029363 | 597,8245371 |
| NQO2 | ENSG00000124588 | 340,1092016 |
| RFLNB ENSG00000183688 | ENSG00000183688 | 540,3769348 |
| EPM2AIP1 | ENSG00000178567 | 423,2875391 |
| IKZF1 | ENSG00000185811 | 461,0960306 |
| EHD4 | ENSG00000103966 | 468,9240779 |
| DCP2 | ENSG00000172795 | 519,8199008 |
| LONRF1 ENSG00000154359 | ENSG00000154359 | 527,8642597 |
| CTSD | ENSG00000117984 | 378,8920381 |
| TBXAS1 | ENSG00000059377 | 465,9193578 |
| PPP1R11 ENSG00000204619 | ENSG00000204619 | 532,1043598 |
| PPTC7 | ENSG00000196850 | 429,6820883 |
| PLEKHM2 | ENSG00000116786 | 410,0383471 |
| HNRNPL ENSG00000104824 | ENSG00000104824 | 508,4669373 |
| TUBB ENSG00000196230 | ENSG00000196230 | 400,1253826 |
| RPN1 | ENSG00000163902 | 547,3644096 |
| XRN2 | ENSG00000088930 | 475,4803729 |
| MACROH2A1 | ENSG00000113648 | 572,133255 |
| HIPK3 | ENSG00000110422 | 561,843198 |
| USF3 | ENSG00000176542 | 429,1604531 |
| GM2A | ENSG00000196743 | 243,5615321 |
| THBD | ENSG00000178726 | 470,2535595 |
| TUT4 | ENSG00000134744 | 631,0277405 |
| PDE3B | ENSG00000152270 | 463,4496789 |
| JAK3 | ENSG00000105639 | 445,4348409 |
| HNRNPF | ENSG00000169813 | 512,0273579 |
| USB1 | ENSG00000103005 | 585,1987389 |
| ANAPC16 | ENSG00000166295 | 487,6875546 |
| FRAT2 | ENSG00000181274 | 302,8912731 |
| FOXJ2 | ENSG00000065970 | 401,6336639 |
| CD300A | ENSG00000167851 | 428,6689409 |
| ITPRID2 | ENSG00000138434 | 591,1151793 |
| NOL4L | ENSG00000197183 | 493,7468318 |
| TCF20 ENSG00000100207 | ENSG00000100207 | 427,0252116 |
| PPP4R1L | ENSG00000124224 | 652,6359824 |
| FBXW7 | ENSG00000109670 | 635,3581538 |
| CHST7 | ENSG00000147119 | 273,8122388 |
| PPP1R2 | ENSG00000184203 | 610,4235028 |
| PNPLA6 | ENSG00000032444 | 716,0630881 |
| BIN2 | ENSG00000110934 | 636,0717888 |
| TCF25 | ENSG00000141002 | 493,1197824 |
| RIPK2 | ENSG00000104312 | 652,8446957 |
| FNDC3A | ENSG00000102531 | 535,082117 |
| UTRN | ENSG00000152818 | 549,5042866 |
| ATP5F1E | ENSG00000124172 | 568,412383 |
| VCPIP1 | ENSG00000175073 | 506,2060328 |

| | | |
|---|---|---|
| MAFF | ENSG00000185022 | 705,1944204 |
| IP6K1 | ENSG00000176095 | 342,6384319 |
| RASA3 ENSG00000185989 | ENSG00000185989 | 393,227531 |
| UPF1 | ENSG00000005007 | 404,6780341 |
| PRKCH | ENSG00000027075 | 611,4351966 |
| FHL3 | ENSG00000183386 | 575,0556026 |
| SEMA4A | ENSG00000196189 | 437,2781825 |
| SRRM1 | ENSG00000133226 | 502,3412231 |
| CTSC | ENSG00000109861 | 533,3709245 |
| HYCC2 | ENSG00000155744 | 568,7250085 |
| AGTRAP | ENSG00000177674 | 335,0961628 |
| MED28 | ENSG00000118579 | 525,5240385 |
| FGD5-AS1 | ENSG00000225733 | 464,4471402 |
| FAM120A | ENSG00000048828 | 509,6105797 |
| SURF4 ENSG00000148248 | ENSG00000148248 | 406,8330955 |
| UBE2J1 | ENSG00000198833 | 484,6196648 |
| TMOD3 | ENSG00000138594 | 566,7577526 |
| VAPA | ENSG00000101558 | 606,1173931 |
| ARAP3 | ENSG00000120318 | 442,3701292 |
| CAMK1D | ENSG00000183049 | 454,3738347 |
| TNRC6B | ENSG00000100354 | 492,147037 |
| BTF3 | ENSG00000145741 | 472,3011233 |
| PPP6R3 | ENSG00000110075 | 527,9920159 |
| IMPDH1 | ENSG00000106348 | 297,7531993 |
| BMF | ENSG00000104081 | 354,7489828 |
| RIOK3 | ENSG00000101782 | 593,8061798 |
| EEF1B2 ENSG00000114942 | ENSG00000114942 | 377,4795623 |
| TRAFD1 | ENSG00000135148 | 399,4391829 |
| KLF7 | ENSG00000118263 | 470,1138684 |
| ZC3H7A | ENSG00000122299 | 582,8219568 |
| XRCC6 | ENSG00000196419 | 492,1780762 |
| MAPK1IP1L | ENSG00000168175 | 550,960752 |
| PPIA | ENSG00000196262 | 432,1846456 |
| RPL29 | ENSG00000162244 | 368,6547974 |
| MTMR10 ENSG00000166912 | ENSG00000166912 | 464,90952 |
| CRKL | ENSG00000099942 | 443,0523561 |
| ATG7 | ENSG00000197548 | 336,6661689 |
| SPCS3 | ENSG00000129128 | 474,7073225 |
| WWP2 | ENSG00000198373 | 455,8655246 |
| ARHGAP27 ENSG00000159314 | ENSG00000159314 | 398,178197 |
| ERO1A | ENSG00000197930 | 505,0461732 |
| PCBP2 | ENSG00000197111 | 558,7946506 |
| FBXW2 | ENSG00000119402 | 414,7758205 |
| CCDC69 | ENSG00000198624 | 453,777379 |
| SLC38A2 | ENSG00000134294 | 486,0261712 |
| FKBP1A | ENSG00000088832 | 543,2465864 |
| C4orf3 | ENSG00000164096 | 529,5553023 |
| SUPT5H | ENSG00000196235 | 428,5486964 |
| LSM14A ENSG00000257103 | ENSG00000257103 | 461,9785725 |
| MAN2B1 | ENSG00000104774 | 433,2669765 |
| DHX40 | ENSG00000108406 | 398,1256986 |
| CBL | ENSG00000110395 | 563,383238 |
| FLII ENSG00000177731 | ENSG00000177731 | 427,8841072 |
| CLIP1 | ENSG00000130779 | 536,2799134 |
| PLXDC2 | ENSG00000120594 | 530,3343673 |

| | | |
|---|---|---|
| PRR33 | ENSG00000283787 | 455,940704 |
| SH3BP5 | ENSG00000131370 | 430,8075461 |
| VPS26A | ENSG00000122958 | 627,128275 |
| UBAP2L | ENSG00000143569 | 540,0329485 |
| BCL10 | ENSG00000142867 | 497,7984107 |
| MT-ND4L | ENSG00000212907 | 501,4931238 |
| ICAM3 | ENSG00000076662 | 427,19471 |
| SUMO1 | ENSG00000116030 | 629,8499592 |
| SRSF11 | ENSG00000116754 | 519,5183277 |
| CLK3 | ENSG00000179335 | 535,3443268 |
| PRR13 | ENSG00000205352 | 514,7830415 |
| FAM157A ENSG00000236438 | ENSG00000236438 | 525,9684676 |
| ZNF143 | ENSG00000166478 | 511,6903083 |
| CDC37 | ENSG00000105401 | 482,8312866 |
| GLRX | ENSG00000173221 | 324,6774911 |
| UHMK1 | ENSG00000152332 | 373,9695719 |
| TPI1 | ENSG00000111669 | 395,6320627 |
| RGS1 | ENSG00000090104 | 586,3617 |
| GPR132 | ENSG00000183484 | 417,2367098 |
| RNF19A | ENSG00000034677 | 467,9975073 |
| TMEM184B | ENSG00000198792 | 428,1057988 |
| DDX39B ENSG00000198563 | ENSG00000198563 | 587,3947195 |
| SFPQ | ENSG00000116560 | 457,6073182 |
| SSR1 | ENSG00000124783 | 438,5096485 |
| VPS18 | ENSG00000104142 | 320,1460497 |
| PPP4R3B | ENSG00000275052 | 515,9324571 |
| NSMAF | ENSG00000035681 | 461,2844725 |
| ENSA | ENSG00000143420 | 517,4759638 |
| SBNO1 | ENSG00000139697 | 507,2173701 |
| GDI2 | ENSG00000057608 | 481,1693679 |
| CIC | ENSG00000079432 | 450,8952234 |
| MR1 | ENSG00000153029 | 473,2417619 |
| TNPO1 | ENSG00000083312 | 409,5508617 |
| ERP44 | ENSG00000023318 | 627,7172108 |
| SETD2 | ENSG00000181555 | 468,0670399 |
| FOXN2 | ENSG00000170802 | 608,9492742 |
| RASA2 | ENSG00000155903 | 630,2421198 |
| ADD3 | ENSG00000148700 | 544,565964 |
| MRTFA | ENSG00000196588 | 524,1318209 |
| USP8 | ENSG00000138592 | 573,8597529 |
| BCAT1 | ENSG00000060982 | 656,5017776 |
| H2AZ2 | ENSG00000105968 | 425,7195544 |
| SPG11 | ENSG00000104133 | 527,2771873 |
| TUG1 | ENSG00000253352 | 484,6467191 |
| PSMB8 ENSG00000204264 | ENSG00000204264 | 415,7433255 |
| JUN | ENSG00000177606 | 431,9280099 |
| LGALS3 | ENSG00000131981 | 336,5135282 |
| SF3B2 | ENSG00000087365 | 431,2083303 |
| TSC22D2 | ENSG00000196428 | 465,0345617 |
| DOCK11 | ENSG00000147251 | 539,019323 |
| CCDC93 | ENSG00000125633 | 675,6442141 |
| DDX18 | ENSG00000088205 | 481,2029626 |
| KLHL24 | ENSG00000114796 | 429,9783528 |
| NONO | ENSG00000147140 | 430,3376021 |
| GNLY | ENSG00000115523 | 228,6827174 |

| | | |
|---|---|---|
| TBC1D30 | ENSG00000111490 | 473,8274582 |
| SAP30BP | ENSG00000161526 | 498,6589781 |
| MGRN1 | ENSG00000102858 | 445,8389719 |
| DR1 | ENSG00000117505 | 437,6096039 |
| SETD1B | ENSG00000139718 | 332,5805328 |
| ARHGDIA | ENSG00000141522 | 400,613729 |
| APOBEC3G | ENSG00000239713 | 397,3099186 |
| FMNL1-DT | ENSG00000267121 | 397,972337 |
| NFYA | ENSG00000001167 | 459,229961 |
| EMB | ENSG00000170571 | 452,5412478 |
| RPS6KA3 | ENSG00000177189 | 563,8475888 |
| ATP1B3 | ENSG00000069849 | 382,9381459 |
| CNOT2 | ENSG00000111596 | 574,493651 |
| RNASET2 | ENSG00000026297 | 546,1927361 |
| ATXN1 | ENSG00000124788 | 585,2431047 |
| SRA1 | ENSG00000213523 | 514,0023946 |
| MICAL1 | ENSG00000135596 | 586,3364371 |
| MAILR | ENSG00000253320 | 368,3376299 |
| MBD2 | ENSG00000134046 | 607,5861763 |
| PSMB4 | ENSG00000159377 | 450,4205997 |
| EXOC8 | ENSG00000116903 | 420,6979671 |
| MPZL3 | ENSG00000160588 | 369,6629764 |
| UNC93B1 | ENSG00000110057 | 391,7371046 |
| PYGL | ENSG00000100504 | 603,8278711 |
| CELF1 | ENSG00000149187 | 416,2224804 |
| NOP53 | ENSG00000105373 | 254,5633812 |
| TBL1X | ENSG00000101849 | 421,4107043 |
| HCG18 ENSG00000231074 | ENSG00000231074 | 446,1350684 |
| LMAN2 | ENSG00000169223 | 248,8077583 |
| USP47 | ENSG00000170242 | 563,7960373 |
| KDM4B | ENSG00000127663 | 401,5815703 |
| NUMA1 | ENSG00000137497 | 466,689754 |
| LTA4H | ENSG00000111144 | 403,8791408 |
| TASL | ENSG00000120280 | 398,2560987 |
| ATP5F1B | ENSG00000110955 | 398,2019571 |
| CEMIP2 | ENSG00000135048 | 369,2764687 |
| ZNF117 | ENSG00000152926 | 526,6158222 |
| PRNP | ENSG00000171867 | 471,6888491 |
| NCKAP1L | ENSG00000123338 | 533,4828535 |
| SRRT | ENSG00000087087 | 559,898624 |
| GLIPR1 | ENSG00000139278 | 436,8212083 |
| ITGAL | ENSG00000005844 | 314,6094838 |
| SLC25A3 | ENSG00000075415 | 430,8203945 |
| VAMP2 | ENSG00000220205 | 402,5322082 |
| CASP1 | ENSG00000137752 | 489,1562714 |
| OSBPL9 | ENSG00000117859 | 576,6004893 |
| COP1 | ENSG00000143207 | 509,8158364 |
| DGKD | ENSG00000077044 | 426,2366012 |
| CYP1B1 | ENSG00000138061 | 342,7131841 |
| BRK1 | ENSG00000254999 | 380,9099388 |
| TRIM21 | ENSG00000132109 | 409,9490434 |
| RPS5 | ENSG00000083845 | 279,2418078 |
| ARL8B | ENSG00000134108 | 483,8274277 |
| HERC1 | ENSG00000103657 | 453,557201 |
| SP140 | ENSG00000079263 | 477,485725 |

| | | |
|---|---|---|
| MINK1 | ENSG00000141503 | 471,3075564 |
| TRIM56 | ENSG00000169871 | 386,4846976 |
| RAB22A | ENSG00000124209 | 395,5723061 |
| LPCAT2 | ENSG00000087253 | 524,6997526 |
| TMEM248 | ENSG00000106609 | 482,234255 |
| CEBPD | ENSG00000221869 | 309,607744 |
| SYNE1 | ENSG00000131018 | 402,5436105 |
| LYSMD2 | ENSG00000140280 | 369,2645577 |
| NEK9 | ENSG00000119638 | 479,384963 |
| GSE1 | ENSG00000131149 | 352,5267669 |
| ENSG00000278600 | ENSG00000278600 | 614,3246496 |
| PHF3 | ENSG00000118482 | 484,4872603 |
| CAT | ENSG00000121691 | 477,6067993 |
| SATB1 | ENSG00000182568 | 538,4653472 |
| CEP350 | ENSG00000135837 | 412,1120011 |
| PNPT1 | ENSG00000138035 | 355,8545249 |
| AGAP3 | ENSG00000133612 | 277,1493148 |
| TMED2 | ENSG00000086598 | 419,1816476 |
| KCTD12 | ENSG00000178695 | 318,5463479 |
| SKIC2 ENSG00000204351 | ENSG00000204351 | 488,3384398 |
| PTPN1 | ENSG00000196396 | 397,7124005 |
| RBL2 | ENSG00000103479 | 409,4473069 |
| HDLBP | ENSG00000115677 | 392,1496494 |
| UBLCP1 | ENSG00000164332 | 439,4082493 |
| RAB14 | ENSG00000119396 | 412,5659215 |
| LILRA6 ENSG00000244482 | ENSG00000244482 | 597,1062892 |
| KREMEN1 | ENSG00000183762 | 294,7378091 |
| ILRUN | ENSG00000196821 | 411,7795351 |
| AGO1 | ENSG00000092847 | 545,4270471 |
| LILRA5 ENSG00000187116 | ENSG00000187116 | 554,0813284 |
| SECTM1 | ENSG00000141574 | 389,1139778 |
| PNP | ENSG00000198805 | 292,9338732 |
| ZBTB7B | ENSG00000160685 | 410,5428701 |
| HGSNAT | ENSG00000165102 | 475,8978617 |
| STK26 | ENSG00000134602 | 397,7873973 |
| ETV3 | ENSG00000117036 | 384,1961742 |
| PADI4 ENSG00000159339 | ENSG00000159339 | 476,0472331 |
| PPP1R16B | ENSG00000101445 | 282,6107347 |
| TBC1D15 | ENSG00000121749 | 499,4122839 |
| CCNT1 | ENSG00000129315 | 404,8472765 |
| SEC24B | ENSG00000138802 | 467,6424056 |
| RBBP6 | ENSG00000122257 | 439,4588937 |
| INPP5K | ENSG00000132376 | 362,7453207 |
| CDK11B | ENSG00000248333 | 483,1859051 |
| SEC23B | ENSG00000101310 | 416,243153 |
| SCLT1 | ENSG00000151466 | 481,7837479 |
| STAU1 | ENSG00000124214 | 467,7736179 |
| VPS8 | ENSG00000156931 | 590,5809877 |
| SYTL3 | ENSG00000164674 | 470,9970802 |
| ZEB1 | ENSG00000148516 | 513,6556894 |
| DHX58 | ENSG00000108771 | 261,400808 |
| ERI1 | ENSG00000104626 | 691,8968769 |
| ATP6V1G1 | ENSG00000136888 | 428,8182887 |
| MBD6 | ENSG00000166987 | 479,3394228 |
| REPS2 | ENSG00000169891 | 332,6346693 |

| | | |
|---|---|---|
| ELAPOR1 | ENSG00000116299 | 322,9437757 |
| NPL | ENSG00000135838 | 464,505848 |
| TINF2 ENSG00000092330 | ENSG00000092330 | 341,1732919 |
| ARID3A | ENSG00000116017 | 385,2041402 |
| SRPK2 | ENSG00000135250 | 497,9705752 |
| CBX3 | ENSG00000122565 | 432,8097236 |
| CCR7 | ENSG00000126353 | 368,3601447 |
| SLC20A1 | ENSG00000144136 | 390,4153296 |
| SPG21 | ENSG00000090487 | 388,1711568 |
| DAP | ENSG00000112977 | 379,7255133 |
| GLS | ENSG00000115419 | 525,0190107 |
| MED14 | ENSG00000180182 | 441,1061031 |
| TRIM33 | ENSG00000197323 | 467,6713753 |
| EFCAB14 | ENSG00000159658 | 368,9953704 |
| PISD | ENSG00000241878 | 297,8857918 |
| PSMD13 | ENSG00000185627 | 439,1818819 |
| AP1S2 | ENSG00000182287 | 358,1013898 |
| HS3ST3B1 | ENSG00000125430 | 365,913543 |
| GGA2 | ENSG00000103365 | 347,7166878 |
| DDX24 ENSG00000089737 | ENSG00000089737 | 380,6268102 |
| SLFN5 | ENSG00000166750 | 324,8397014 |
| GBA2 | ENSG00000070610 | 428,9460717 |
| CD274 | ENSG00000120217 | 297,2933756 |
| SELPLG | ENSG00000110876 | 345,9783555 |
| ACIN1 | ENSG00000100813 | 397,8419248 |
| CTSL | ENSG00000135047 | 363,0430432 |
| BTN3A1 | ENSG00000026950 | 369,0078008 |
| TMED10 | ENSG00000170348 | 460,3727513 |
| ATXN7L3 | ENSG00000087152 | 349,2162608 |
| GPAT3 | ENSG00000138678 | 352,9938798 |
| PLP2 | ENSG00000102007 | 414,9677527 |
| UBXN7 | ENSG00000163960 | 446,3455315 |
| ABHD17A | ENSG00000129968 | 321,411783 |
| CCNT2 | ENSG00000082258 | 383,1194228 |
| PKN2 | ENSG00000065243 | 547,5874795 |
| MYO1G | ENSG00000136286 | 380,459598 |
| COPB2 | ENSG00000184432 | 432,9792637 |
| PCBP1-AS1 | ENSG00000179818 | 450,6352578 |
| BTAF1 | ENSG00000095564 | 531,3286664 |
| RB1 | ENSG00000139687 | 524,746684 |
| VCAN | ENSG00000038427 | 371,7897297 |
| GOLM2 | ENSG00000166734 | 520,4436154 |
| IDS | ENSG00000010404 | 321,484008 |
| ITPKB | ENSG00000143772 | 304,2991262 |
| BANP | ENSG00000172530 | 277,4939787 |
| SRP14 | ENSG00000140319 | 410,7948996 |
| YWHAG | ENSG00000170027 | 310,483203 |
| SET | ENSG00000119335 | 404,3955329 |
| CUL3 | ENSG00000036257 | 439,1501259 |
| TNRC6C | ENSG00000078687 | 349,2339414 |
| EAF1 | ENSG00000144597 | 333,5041515 |
| SAFB | ENSG00000160633 | 409,9713818 |
| ABR ENSG00000159842 | ENSG00000159842 | 343,2729373 |
| PCMTD1 | ENSG00000168300 | 445,9069766 |
| FAR1 | ENSG00000197601 | 485,5513867 |

| | | |
|---|---|---|
| TNFRSF10C | ENSG00000173535 | 335,4123597 |
| BAG1 | ENSG00000107262 | 333,6479768 |
| CHD3 | ENSG00000170004 | 437,9816636 |
| PMAIP1 | ENSG00000141682 | 460,3979721 |
| ENSG00000237550 | ENSG00000237550 | 273,2761673 |
| TBC1D23 | ENSG00000036054 | 454,5099314 |
| PIK3CA | ENSG00000121879 | 417,3706999 |
| APOL2 | ENSG00000128335 | 400,8594699 |
| PSMA3-AS1 | ENSG00000257621 | 444,9750134 |
| CBFB | ENSG00000067955 | 339,6038589 |
| SH3GL1 | ENSG00000141985 | 344,8173489 |
| WBP1L | ENSG00000166272 | 272,9591164 |
| DCP1A | ENSG00000272886 | 423,9013749 |
| UBE2W | ENSG00000104343 | 508,1193861 |
| PHC3 | ENSG00000173889 | 392,6112262 |
| SKP1 | ENSG00000113558 | 413,1845027 |
| LYSMD3 | ENSG00000176018 | 419,6706851 |
| P2RX1 | ENSG00000108405 | 482,1152853 |
| CCDC186 | ENSG00000165813 | 436,5574074 |
| MCMBP | ENSG00000197771 | 316,7938812 |
| TOR1A | ENSG00000136827 | 459,6414923 |
| YY1AP1 | ENSG00000163374 | 399,3703701 |
| PRKCSH | ENSG00000130175 | 349,6923954 |
| YIPF3 | ENSG00000137207 | 340,3732033 |
| KIF21B | ENSG00000116852 | 436,6980813 |
| PRELID1 | ENSG00000169230 | 288,5477124 |
| LACTB | ENSG00000103642 | 411,6509655 |
| AGO3 | ENSG00000126070 | 395,0607065 |
| C11orf58 | ENSG00000110696 | 384,1872369 |
| COQ10B | ENSG00000115520 | 403,813479 |
| DDIT3 | ENSG00000175197 | 528,1354846 |
| UQCRC2 | ENSG00000140740 | 347,1632914 |
| YIPF4 | ENSG00000119820 | 392,0729921 |
| TRIM69 ENSG00000185880 | ENSG00000185880 | 392,9783769 |
| RPL36AL | ENSG00000165502 | 337,9294801 |
| HSPH1 | ENSG00000120694 | 253,9269307 |
| TMEM120A | ENSG00000189077 | 400,1684053 |
| PPP3CA | ENSG00000138814 | 491,0149788 |
| ARK2N | ENSG00000152242 | 378,3338863 |
| BLOC1S6 | ENSG00000104164 | 423,9712551 |
| GNA15 | ENSG00000060558 | 335,3694106 |
| LCOR | ENSG00000196233 | 482,9180226 |
| ADD1 | ENSG00000087274 | 376,445963 |
| DYNC1H1 | ENSG00000197102 | 328,8317661 |
| TAP2 ENSG00000204267 | ENSG00000204267 | 380,560538 |
| DYNLT3 | ENSG00000165169 | 475,7728249 |
| TSPYL1 | ENSG00000189241 | 347,3495352 |
| XIAP | ENSG00000101966 | 384,4212662 |
| SLC23A2 | ENSG00000089057 | 436,6995632 |
| EYA3 | ENSG00000158161 | 352,2718374 |
| CLEC2B | ENSG00000110852 | 509,3777302 |
| NDE1 ENSG00000072864 | ENSG00000072864 | 267,6126032 |
| GUCD1 | ENSG00000138867 | 331,940281 |
| TRAPPC10 | ENSG00000160218 | 376,6159496 |
| AFF1 | ENSG00000172493 | 470,3114086 |

| | | |
|---|---|---|
| HNRNPDL | ENSG00000152795 | 396,8343461 |
| CBX4 | ENSG00000141582 | 268,1044432 |
| IL18RAP | ENSG00000115607 | 449,0370015 |
| LINC00963 | ENSG00000204054 | 346,6125401 |
| NRBP1 | ENSG00000115216 | 336,697835 |
| RAB35 | ENSG00000111737 | 369,399687 |
| PLEKHM1P1 | ENSG00000214176 | 354,0980249 |
| GLG1 | ENSG00000090863 | 314,1914475 |
| HSPA13 | ENSG00000155304 | 481,3966738 |
| HP1BP3 | ENSG00000127483 | 413,8384425 |
| GPR107 | ENSG00000148358 | 315,4497906 |
| RNPEPL1 | ENSG00000142327 | 273,6062599 |
| TOPORS | ENSG00000197579 | 355,3551439 |
| FRMD8 | ENSG00000126391 | 235,4575793 |
| VNN1 | ENSG00000112299 | 233,5208238 |
| NUP50 | ENSG00000093000 | 409,6000802 |
| KLHL2 | ENSG00000109466 | 479,9848461 |
| BLOC1S2 | ENSG00000196072 | 361,4640162 |
| SLC9A1 | ENSG00000090020 | 329,8091051 |
| S1PR1 | ENSG00000170989 | 260,3905015 |
| OSTM1 | ENSG00000081087 | 414,9722548 |
| TMEM165 | ENSG00000134851 | 276,6220977 |
| OGDH | ENSG00000105953 | 260,196808 |
| ARGLU1 | ENSG00000134884 | 451,9603468 |
| LMBRD1 | ENSG00000168216 | 427,9179261 |
| DICER1 | ENSG00000100697 | 298,2654751 |
| PSME3 | ENSG00000131467 | 374,9473191 |
| SMAD7 | ENSG00000101665 | 306,4424991 |
| CANT1 | ENSG00000171302 | 283,6189849 |
| REC8 ENSG00000100918 | ENSG00000100918 | 420,2608343 |
| SBF1 | ENSG00000100241 | 276,8427229 |
| ASH1L | ENSG00000116539 | 381,0009906 |
| RRAGD | ENSG00000025039 | 284,4231007 |
| RPL38 | ENSG00000172809 | 324,6945474 |
| IL1A | ENSG00000115008 | 388,9329259 |
| NR2C2 | ENSG00000177463 | 370,2642355 |
| PGAM1 | ENSG00000171314 | 356,8624952 |
| TENT5A | ENSG00000112773 | 256,4583367 |
| ARID1A | ENSG00000117713 | 263,6331474 |
| S100A10 | ENSG00000197747 | 319,4705655 |
| STX7 | ENSG00000079950 | 376,5959774 |
| ILF3 | ENSG00000129351 | 342,4446046 |
| C9orf78 | ENSG00000136819 | 377,4532489 |
| KIF3B | ENSG00000101350 | 267,9511278 |
| HLA-DRB1 ENSG00000196126 | ENSG00000196126 | 315,226625 |
| RAD23B | ENSG00000119318 | 443,5001624 |
| TICAM1 | ENSG00000127666 | 273,1950609 |
| DHX9 | ENSG00000135829 | 378,0948645 |
| LAPTM4A | ENSG00000068697 | 413,320294 |
| SPTBN1 | ENSG00000115306 | 312,8656949 |
| RNF24 | ENSG00000101236 | 477,6106659 |
| SNX6 | ENSG00000129515 | 421,0827031 |
| MAP2K1 | ENSG00000169032 | 369,8821075 |
| DUSP5 | ENSG00000138166 | 285,5541774 |
| C6orf89 | ENSG00000198663 | 305,4352846 |

| | | |
|---|---|---|
| SNX27 | ENSG00000143376 | 363,7450799 |
| GOLGA4 | ENSG00000144674 | 419,6528073 |
| SBF2 | ENSG00000133812 | 445,5951865 |
| REST | ENSG00000084093 | 315,966002 |
| CCL5 ENSG00000271503 | ENSG00000271503 | 172,8378764 |
| HSPD1 | ENSG00000144381 | 272,0492388 |
| SERF2 | ENSG00000140264 | 309,9406701 |
| ZMAT2 | ENSG00000146007 | 350,6440733 |
| SRP54 | ENSG00000100883 | 418,9888044 |
| TNF ENSG00000232810 | ENSG00000232810 | 326,4688779 |
| NPM1 | ENSG00000181163 | 309,7269562 |
| RPL35 | ENSG00000136942 | 277,128187 |
| VPS13B | ENSG00000132549 | 427,7752978 |
| ZNF276 | ENSG00000158805 | 450,6708159 |
| NCOR1 | ENSG00000141027 | 386,3310329 |
| MIA3 | ENSG00000154305 | 399,4908018 |
| IRAG1 | ENSG00000072952 | 426,9121969 |
| NADSYN1 | ENSG00000172890 | 397,7512175 |
| ERF | ENSG00000105722 | 249,2523017 |
| CCNY | ENSG00000108100 | 318,5339952 |
| PTGER4 | ENSG00000171522 | 293,7653833 |
| PIGX | ENSG00000163964 | 301,2586142 |
| MAP2K4 | ENSG00000065559 | 362,5521769 |
| GSTO1 | ENSG00000148834 | 440,5487443 |
| LARP4B | ENSG00000107929 | 385,3215439 |
| FBXL3 | ENSG00000005812 | 326,4315598 |
| MRNIP ENSG00000161010 | ENSG00000161010 | 487,7586496 |
| TRAM1 | ENSG00000067167 | 386,9388461 |
| IKBKB | ENSG00000104365 | 379,1342956 |
| HNRNPM | ENSG00000099783 | 379,4697925 |
| HELB | ENSG00000127311 | 371,5926447 |
| MED15 | ENSG00000099917 | 323,4040532 |
| CCND2 | ENSG00000118971 | 357,7986903 |
| RAB8A | ENSG00000167461 | 330,7852698 |
| WASF2 | ENSG00000158195 | 311,89787 |
| EPS15L1 | ENSG00000127527 | 350,2228729 |
| CARS1 ENSG00000110619 | ENSG00000110619 | 307,843219 |
| GPR84 | ENSG00000139572 | 458,6517161 |
| PCF11 | ENSG00000165494 | 365,665649 |
| TAOK3 | ENSG00000135090 | 395,356121 |
| RAB20 | ENSG00000139832 | 304,2015208 |
| SPHK1 | ENSG00000176170 | 339,0444215 |
| IREB2 | ENSG00000136381 | 343,7888142 |
| CTC1 | ENSG00000178971 | 351,9489724 |
| PRDX5 | ENSG00000126432 | 273,836501 |
| DNTTIP1 | ENSG00000101457 | 327,1332796 |
| CHMP1A | ENSG00000131165 | 242,312596 |
| MKNK1 | ENSG00000079277 | 414,6427027 |
| NR4A2 | ENSG00000153234 | 395,4743016 |
| CD47 | ENSG00000196776 | 334,8190597 |
| TCP1 | ENSG00000120438 | 338,3586359 |
| USP25 | ENSG00000155313 | 439,1989221 |
| ADNP2 | ENSG00000101544 | 284,6045298 |
| NDFIP1 | ENSG00000131507 | 282,634506 |
| FYTTD1 | ENSG00000122068 | 352,1431869 |

| | | |
|---|---|---|
| NLRP12 | ENSG00000142405 | 291,4529573 |
| ANKLE2 | ENSG00000176915 | 333,0906636 |
| SH2D3A | ENSG00000125731 | 411,5797042 |
| CCSER2 | ENSG00000107771 | 328,2400454 |
| ITPK1 ENSG00000100605 | ENSG00000100605 | 331,9674719 |
| SMG7 | ENSG00000116698 | 277,1256827 |
| TGIF2 | ENSG00000118707 | 297,0511118 |
| HNRNPA0 | ENSG00000177733 | 260,2843482 |
| SOAT1 | ENSG00000057252 | 473,7790039 |
| LILRA2 ENSG00000239998 | ENSG00000239998 | 312,7713985 |
| P2RY13 | ENSG00000181631 | 348,1751865 |
| CERT1 | ENSG00000113163 | 318,65582 |
| ATXN1L | ENSG00000224470 | 337,3998644 |
| DCAF11 ENSG00000100897 | ENSG00000100897 | 401,1471594 |
| YY1 | ENSG00000100811 | 330,7924294 |
| AKIRIN2 | ENSG00000135334 | 446,4843092 |
| ZNF274 | ENSG00000171606 | 301,0177576 |
| CGGBP1 | ENSG00000163320 | 333,3680459 |
| ZNF281 | ENSG00000162702 | 448,3417606 |
| HNRNPD | ENSG00000138668 | 382,4133611 |
| GNAQ | ENSG00000156052 | 448,7326661 |
| BBC3 | ENSG00000105327 | 166,5872261 |
| TMEM43 | ENSG00000170876 | 287,8144925 |
| ALPK1 | ENSG00000073331 | 434,2777452 |
| EIF3E | ENSG00000104408 | 303,8387918 |
| OSBP | ENSG00000110048 | 302,0036646 |
| SEC31A | ENSG00000138674 | 396,7752936 |
| ZDHHC7 | ENSG00000153786 | 321,1933375 |
| RALBP1 | ENSG00000017797 | 308,3002488 |
| MAP3K13 | ENSG00000073803 | 351,3278567 |
| PPP1R12C | ENSG00000125503 | 301,3645786 |
| EGR3 | ENSG00000179388 | 423,1224952 |
| CSTB | ENSG00000160213 | 329,5916218 |
| CEP63 | ENSG00000182923 | 349,8244267 |
| XKR8 | ENSG00000158156 | 296,7103436 |
| RPS15 | ENSG00000115268 | 224,7326084 |
| EIF4G1 | ENSG00000114867 | 295,9755604 |
| TFEB | ENSG00000112561 | 250,0594054 |
| EXTL3 | ENSG00000012232 | 269,7560985 |
| RALY | ENSG00000125970 | 294,2376616 |
| CHD7 | ENSG00000171316 | 400,2411326 |
| USP36 | ENSG00000055483 | 287,9357789 |
| LINC01002 | ENSG00000282508 | 399,2294186 |
| FHIP2A | ENSG00000151553 | 315,685609 |
| ABCA7 | ENSG00000064687 | 291,5844445 |
| GTF2B | ENSG00000137947 | 363,6368924 |
| GPR183 | ENSG00000169508 | 362,2715628 |
| SIRT1 | ENSG00000096717 | 236,522236 |
| LSM10 | ENSG00000181817 | 283,4493862 |
| YOD1 | ENSG00000180667 | 501,7145841 |
| PSME3IP1 | ENSG00000172775 | 369,3444629 |
| TGFBI | ENSG00000120708 | 384,8881915 |
| MED12 | ENSG00000184634 | 383,4953392 |
| MARK3 | ENSG00000075413 | 305,2243974 |
| EIF3H | ENSG00000147677 | 346,4905575 |

| | | |
|---|---|---|
| MCTP1 | ENSG00000175471 | 296,7147602 |
| ADIPOR2 ENSG00000006831 | ENSG00000006831 | 359,0280362 |
| GPX1 | ENSG00000233276 | 279,0274096 |
| MYSM1 | ENSG00000162601 | 408,0593099 |
| TRAPPC8 | ENSG00000153339 | 331,2202969 |
| SMCR8 ENSG00000176994 | ENSG00000176994 | 245,9836311 |
| USP19 | ENSG00000172046 | 260,1541519 |
| MFSD1 | ENSG00000118855 | 412,2537048 |
| NPLOC4 | ENSG00000182446 | 269,8489602 |
| ARHGAP15 | ENSG00000075884 | 531,1670854 |
| USPL1 | ENSG00000132952 | 479,1763406 |
| CCDC71L | ENSG00000253276 | 361,7592679 |
| ZNF655 | ENSG00000197343 | 358,3654478 |
| HEXB | ENSG00000049860 | 331,5437285 |
| CMTR1 | ENSG00000137200 | 332,0233723 |
| RUNX1 | ENSG00000159216 | 302,4622204 |
| BAP1 | ENSG00000163930 | 265,8564396 |
| CREG1 | ENSG00000143162 | 407,1662556 |
| BCAP31 | ENSG00000185825 | 311,0647026 |
| BMP2K | ENSG00000138756 | 339,1820383 |
| TRABD | ENSG00000170638 | 403,7456574 |
| RNF103 | ENSG00000239305 | 338,9352145 |
| GRIPAP1 | ENSG00000068400 | 385,7321281 |
| HSPA1A ENSG00000204389 | ENSG00000204389 | 419,1786432 |
| SLC19A1 | ENSG00000173638 | 255,3074959 |
| USP20 | ENSG00000136878 | 326,1331682 |
| KCNE3 | ENSG00000175538 | 237,0066341 |
| RBM33 | ENSG00000184863 | 332,5935084 |
| CBX6 | ENSG00000183741 | 336,0783462 |
| STK38 | ENSG00000112079 | 392,2067479 |
| FKBP15 | ENSG00000119321 | 296,5529175 |
| VPS9D1 | ENSG00000075399 | 380,8245701 |
| RC3H2 | ENSG00000056586 | 302,4539497 |
| MMADHC | ENSG00000168288 | 308,3991244 |
| KANSL3 | ENSG00000114982 | 361,0801259 |
| YWHAQ | ENSG00000134308 | 299,1097671 |
| FEM1C | ENSG00000145780 | 448,6188818 |
| CNOT8 | ENSG00000155508 | 340,632572 |
| MKLN1 | ENSG00000128585 | 354,0461688 |
| YWHAE ENSG00000108953 | ENSG00000108953 | 334,6193101 |
| IL6ST | ENSG00000134352 | 360,259255 |
| MAP1LC3A | ENSG00000101460 | 257,5204875 |
| LIN54 | ENSG00000189308 | 350,7910435 |
| HELZ | ENSG00000198265 | 319,6601264 |
| PCGF5 | ENSG00000180628 | 361,1779289 |
| H1-2 | ENSG00000187837 | 196,5738148 |
| CARD16 | ENSG00000204397 | 390,9482502 |
| RPS21 | ENSG00000171858 | 220,6522346 |
| TRIM26 ENSG00000234127 | ENSG00000234127 | 301,1183403 |
| UBB | ENSG00000170315 | 311,7313942 |
| HSPA9 | ENSG00000113013 | 326,5908086 |
| TRIM5 | ENSG00000132256 | 290,1614182 |
| SUCO | ENSG00000094975 | 314,4459904 |
| ABCC1 ENSG00000103222 | ENSG00000103222 | 234,5939075 |
| SIRPB2 | ENSG00000196209 | 195,759361 |

| | | |
|---|---|---|
| NLRC4 | ENSG00000091106 | 402,5107118 |
| RGL4 | ENSG00000159496 | 312,1717728 |
| BST1 | ENSG00000109743 | 347,7781776 |
| ZMYM2 | ENSG00000121741 | 424,614981 |
| ASPH | ENSG00000198363 | 392,3406581 |
| ZNF746 | ENSG00000181220 | 253,0570775 |
| PCNP | ENSG00000081154 | 388,7028232 |
| SLC22A4 | ENSG00000197208 | 340,4074686 |
| ZNF333 | ENSG00000160961 | 364,9320597 |
| TM2D3 | ENSG00000184277 | 318,7713972 |
| CREB5 | ENSG00000146592 | 395,5560347 |
| GIT2 | ENSG00000139436 | 316,6455884 |
| OSBPL11 | ENSG00000144909 | 301,5035729 |
| IL2RB | ENSG00000100385 | 226,1469286 |
| PFKL | ENSG00000141959 | 235,1794726 |
| CDK12 | ENSG00000167258 | 292,5277234 |
| NECAP2 | ENSG00000157191 | 262,8453583 |
| PRPF40A | ENSG00000196504 | 363,0934797 |
| RHEB | ENSG00000106615 | 324,078904 |
| TMUB2 | ENSG00000168591 | 292,1885716 |
| AP5Z1 | ENSG00000242802 | 246,2885813 |
| KCMF1 | ENSG00000176407 | 302,1879888 |
| ENTPD1 | ENSG00000138185 | 400,0701323 |
| ARHGAP18 | ENSG00000146376 | 354,8783586 |
| BSDC1 | ENSG00000160058 | 297,6467105 |
| ENSG00000203644 | ENSG00000203644 | 379,6421242 |
| USP22 | ENSG00000124422 | 224,2542478 |
| NCOA3 | ENSG00000124151 | 348,3968307 |
| RRP12 | ENSG00000052749 | 326,5701863 |
| ARRDC4 | ENSG00000140450 | 244,4740318 |
| LINC00528 | ENSG00000269220 | 289,3078379 |
| MAX | ENSG00000125952 | 293,9187213 |
| LEPROTL1 | ENSG00000104660 | 291,2666312 |
| IGF1R | ENSG00000140443 | 326,9487638 |
| PACS1 | ENSG00000175115 | 299,1978001 |
| RSRC2 | ENSG00000111011 | 354,0794253 |
| HERC6 | ENSG00000138642 | 222,261607 |
| SMNDC1 | ENSG00000119953 | 320,6849054 |
| KTN1 | ENSG00000126777 | 368,2406239 |
| NPEPPS | ENSG00000141279 | 387,591566 |
| MON2 | ENSG00000061987 | 340,6961385 |
| FAM120AOS | ENSG00000188938 | 281,1685995 |
| CD96 | ENSG00000153283 | 205,6636799 |
| TCP11L2 | ENSG00000166046 | 293,1573145 |
| PSMA7 | ENSG00000101182 | 251,786159 |
| SEPTIN6 | ENSG00000125354 | 331,8711948 |
| HES4 | ENSG00000188290 | 268,0568766 |
| ATRX | ENSG00000085224 | 397,0676261 |
| ZKSCAN1 | ENSG00000106261 | 339,066477 |
| CTBS | ENSG00000117151 | 391,8205623 |
| SNW1 | ENSG00000100603 | 306,0999245 |
| ANKRD17 | ENSG00000132466 | 368,0734617 |
| DDAH2 ENSG00000213722 | ENSG00000213722 | 354,6282513 |
| SMARCA2 | ENSG00000080503 | 340,3074078 |
| PCNX4 | ENSG00000126773 | 314,2121639 |

| | | |
|---|---|---|
| MT-ND6 | ENSG00000198695 | 256,2211361 |
| GMPR2 ENSG00000100938 | ENSG00000100938 | 305,6283781 |
| EIF3A | ENSG00000107581 | 281,9285606 |
| STT3B | ENSG00000163527 | 347,5619201 |
| WDR45 | ENSG00000196998 | 260,4304385 |
| RPGR | ENSG00000156313 | 363,7057956 |
| DTX4 | ENSG00000110042 | 157,623329 |
| SLC25A6 | ENSG00000169100 | 203,1767955 |
| CLPTM1L ENSG00000049656 | ENSG00000049656 | 286,163215 |
| FXR1 | ENSG00000114416 | 300,289776 |
| RNF40 | ENSG00000103549 | 278,1361452 |
| FRY | ENSG00000073910 | 349,3534573 |
| INPP5A | ENSG00000068383 | 333,826565 |
| DVL3 | ENSG00000161202 | 373,5773841 |
| ANXA2 | ENSG00000182718 | 275,8005098 |
| ARF5 | ENSG00000004059 | 230,9407542 |
| GLYR1 | ENSG00000140632 | 290,6696421 |
| UVSSA | ENSG00000163945 | 231,9111973 |
| FGFR1OP2 | ENSG00000111790 | 373,8683845 |
| EREG | ENSG00000124882 | 239,1185567 |
| UBE2D2 | ENSG00000131508 | 299,223779 |
| LINC00877 | ENSG00000241163 | 266,2805869 |
| IL2RG | ENSG00000147168 | 313,9744867 |
| USP18 | ENSG00000184979 | 163,6121107 |
| LARP1 | ENSG00000155506 | 232,6253456 |
| EPS15 | ENSG00000085832 | 363,265074 |
| ZDHHC20 | ENSG00000180776 | 300,124668 |
| CRNKL1 | ENSG00000101343 | 250,6463691 |
| ANP32E | ENSG00000143401 | 289,5516847 |
| ZNF277 | ENSG00000198839 | 377,2644529 |
| RIPK1 | ENSG00000137275 | 295,3976137 |
| EIF1B | ENSG00000114784 | 390,7169522 |
| RTF1 | ENSG00000137815 | 316,9444097 |
| PITPNM1 | ENSG00000110697 | 246,670913 |
| ZBED1 | ENSG00000214717 | 240,910221 |
| RBM7 | ENSG00000076053 | 365,7806787 |
| PRR7 | ENSG00000131188 | 177,3718832 |
| UBE2M | ENSG00000130725 | 252,6791741 |
| HLA-DPB1 ENSG00000223865 | ENSG00000223865 | 206,0633027 |
| RETREG2 | ENSG00000144567 | 240,5919104 |
| NKTR | ENSG00000114857 | 384,434398 |
| PIK3CB | ENSG00000051382 | 328,4884462 |
| RNF44 | ENSG00000146083 | 269,3798316 |
| CDKN2D | ENSG00000129355 | 302,239693 |
| RPS28 | ENSG00000233927 | 206,4218342 |
| GGNBP2 ENSG00000278311 | ENSG00000278311 | 327,6222159 |
| IRF2BPL | ENSG00000119669 | 233,4981327 |
| PDCD6IP | ENSG00000170248 | 388,4760386 |
| PPP3R1 | ENSG00000221823 | 342,9732134 |
| MAPK8IP3 | ENSG00000138834 | 274,5088962 |
| PRKACA | ENSG00000072062 | 258,987586 |
| SYNRG ENSG00000275066 | ENSG00000275066 | 278,1677421 |
| ENSG00000230551 | ENSG00000230551 | 416,6417371 |
| RHBDD2 | ENSG00000005486 | 344,625599 |
| HOOK3 | ENSG00000168172 | 325,4702507 |

| | | |
|---|---|---|
| MLKL | ENSG00000168404 | 331,4543713 |
| KLHL5 | ENSG00000109790 | 311,6423833 |
| LPXN | ENSG00000110031 | 246,7667258 |
| DEK | ENSG00000124795 | 354,9864459 |
| ADSS2 | ENSG00000035687 | 242,9553713 |
| CIAO1 | ENSG00000144021 | 312,5741398 |
| FZR1 | ENSG00000105325 | 219,0060384 |
| TENT5C | ENSG00000183508 | 201,5857871 |
| SEMA7A | ENSG00000138623 | 422,2911067 |
| PHF21A | ENSG00000135365 | 370,296021 |
| PSMB1 ENSG00000008018 | ENSG00000008018 | 285,1213691 |
| C1GALT1 | ENSG00000106392 | 340,8880628 |
| ZNF316 | ENSG00000205903 | 183,6877531 |
| EPB41L3 | ENSG00000082397 | 255,1520447 |
| ABHD13 | ENSG00000139826 | 345,129541 |
| VPS39 | ENSG00000166887 | 264,0869255 |
| ADAMTSL4 | ENSG00000143382 | 325,574958 |
| ELMO1 | ENSG00000155849 | 317,852447 |
| TPRG1L | ENSG00000158109 | 198,144931 |
| RASGRP2 | ENSG00000068831 | 246,1913147 |
| DYNLT1 | ENSG00000146425 | 328,1455106 |
| ACADVL | ENSG00000072778 | 281,2737023 |
| CHCHD2 | ENSG00000106153 | 279,4353409 |
| UBE2Z | ENSG00000159202 | 208,4779454 |
| ABHD4 | ENSG00000100439 | 243,6345106 |
| IDI1 | ENSG00000067064 | 403,3498037 |
| PIKFYVE | ENSG00000115020 | 337,412099 |
| SNRNP70 | ENSG00000104852 | 297,0854484 |
| PAF1 | ENSG00000006712 | 318,078246 |
| TXN | ENSG00000136810 | 384,2303172 |
| CCL2 | ENSG00000108691 | 125,3753125 |
| ZNF394 | ENSG00000160908 | 280,8279304 |
| MAP3K1 | ENSG00000095015 | 281,4618733 |
| TAF13 | ENSG00000197780 | 354,0689739 |
| TNK2 | ENSG00000061938 | 263,9990858 |
| RSF1 | ENSG00000048649 | 316,50852 |
| RALGAPB | ENSG00000170471 | 286,2758746 |
| GSPT1 | ENSG00000103342 | 236,0093075 |
| SLC35B2 | ENSG00000157593 | 318,287368 |
| PI4KA | ENSG00000241973 | 306,3699509 |
| PSMA4 | ENSG00000041357 | 351,6709801 |
| RNF139 | ENSG00000170881 | 273,3678735 |
| ADAM9 ENSG00000168615 | ENSG00000168615 | 362,5161277 |
| AKAP9 | ENSG00000127914 | 325,6026053 |
| SEC13 | ENSG00000157020 | 245,754816 |
| ATP5PB | ENSG00000116459 | 256,9426484 |
| YBX3 | ENSG00000060138 | 278,398891 |
| FBXO11 | ENSG00000138081 | 349,167329 |
| DHX36 ENSG00000174953 | ENSG00000174953 | 298,4848313 |
| PPM1K | ENSG00000163644 | 200,8644929 |
| ZBTB48 | ENSG00000204859 | 242,8787214 |
| LMNA | ENSG00000160789 | 167,3621271 |
| SPTAN1 | ENSG00000197694 | 274,6675815 |
| FBRSL1 | ENSG00000112787 | 226,858092 |
| GOLPH3 | ENSG00000113384 | 262,8907325 |

| | | |
|---|---|---|
| RO60 | ENSG00000116747 | 308,8012061 |
| CCNL2 | ENSG00000221978 | 290,733526 |
| STIP1 | ENSG00000168439 | 270,3395291 |
| PITPNC1 | ENSG00000154217 | 258,9794578 |
| DHX8 | ENSG00000067596 | 239,0889199 |
| WDR48 | ENSG00000114742 | 300,4948834 |
| PRKAG1 | ENSG00000181929 | 285,4444833 |
| RPS26 | ENSG00000197728 | 206,1636914 |
| HIC1 | ENSG00000177374 | 217,4590909 |
| INKA2 | ENSG00000197852 | 212,3594427 |
| UGGT1 | ENSG00000136731 | 283,3265354 |
| ZNF646 | ENSG00000167395 | 241,7941158 |
| KAT2B | ENSG00000114166 | 373,035199 |
| METTL9 ENSG00000197006 | ENSG00000197006 | 308,8795245 |
| UBR1 | ENSG00000159459 | 407,2512977 |
| CLEC2D | ENSG00000069493 | 286,0399065 |
| PRDM8 | ENSG00000152784 | 282,1962492 |
| SYNGR2 | ENSG00000108639 | 230,2654328 |
| ZSWIM8 | ENSG00000214655 | 231,4679541 |
| IRGQ | ENSG00000167378 | 325,9543686 |
| NELFE ENSG00000204356 | ENSG00000204356 | 258,5274065 |
| VAMP3 | ENSG00000049245 | 318,1710342 |
| EFR3A | ENSG00000132294 | 303,3797412 |
| CHFR | ENSG00000072609 | 248,7287893 |
| TRA2A | ENSG00000164548 | 313,1361208 |
| CDC73 | ENSG00000134371 | 326,7590513 |
| SLC11A2 | ENSG00000110911 | 270,4553907 |
| EXOC3 | ENSG00000180104 | 259,0178966 |
| U2AF2 | ENSG00000063244 | 235,6454862 |
| HPS5 | ENSG00000110756 | 291,1768744 |
| EAPP | ENSG00000129518 | 237,4651463 |
| SAP18 | ENSG00000150459 | 276,8148068 |
| BRAT1 | ENSG00000106009 | 254,0483764 |
| MADD | ENSG00000110514 | 265,8931318 |
| ADNP | ENSG00000101126 | 257,6347135 |
| ICOSLG | ENSG00000160223 | 170,6231082 |
| E2F4 | ENSG00000205250 | 230,7818301 |
| ARAF | ENSG00000078061 | 319,3113124 |
| ZBTB34 | ENSG00000177125 | 302,4362439 |
| DENND1C | ENSG00000205744 | 238,053981 |
| MAU2 | ENSG00000129933 | 285,5073413 |
| YKT6 | ENSG00000106636 | 198,8114004 |
| PLAGL1 | ENSG00000118495 | 362,5103221 |
| SPART | ENSG00000133104 | 341,89016 |
| UBE2A | ENSG00000077721 | 358,3503111 |
| UBE4A | ENSG00000110344 | 301,936609 |
| FAM193B | ENSG00000146067 | 321,3223854 |
| CUL4B | ENSG00000158290 | 298,457648 |
| RPL23A | ENSG00000198242 | 208,640643 |
| PPP1CC | ENSG00000186298 | 195,6570901 |
| ATOSA | ENSG00000047346 | 294,3444816 |
| GOLGA2 | ENSG00000167110 | 277,9873089 |
| SELENOF | ENSG00000183291 | 326,009595 |
| UBL5 | ENSG00000198258 | 246,5664851 |
| TSEN34 ENSG00000170892 | ENSG00000170892 | 272,1584685 |

| | | |
|---|---|---|
| RUBCN | ENSG00000145016 | 249,4819886 |
| PPARD | ENSG00000112033 | 322,8237697 |
| ZNF687 | ENSG00000143373 | 290,432035 |
| PTPN22 | ENSG00000134242 | 259,7860195 |
| DNAJB2 | ENSG00000135924 | 251,0909994 |
| NLRP6 | ENSG00000174885 | 199,0376876 |
| AP2M1 | ENSG00000161203 | 270,2924998 |
| USF2 | ENSG00000105698 | 242,2577454 |
| BRWD1 | ENSG00000185658 | 314,2942264 |
| DCTN4 | ENSG00000132912 | 326,9838523 |
| ITPR2 | ENSG00000123104 | 298,4353685 |
| MEPCE | ENSG00000146834 | 206,9938387 |
| CISH | ENSG00000114737 | 386,1110779 |
| CLK4 | ENSG00000113240 | 262,7768133 |
| RIPOR1 | ENSG00000039523 | 251,7121791 |
| RAP1GAP2 | ENSG00000132359 | 273,7684347 |
| TAB3 | ENSG00000157625 | 276,9262562 |
| EIF3D | ENSG00000100353 | 227,9661286 |
| GORASP1 | ENSG00000114745 | 339,0668872 |
| AGFG1 | ENSG00000173744 | 335,2641938 |
| TLR8 | ENSG00000101916 | 286,6301124 |
| EIF1AX | ENSG00000173674 | 244,7745731 |
| CCDC9 | ENSG00000105321 | 272,3954542 |
| PSMD2 | ENSG00000175166 | 247,8030875 |
| DLST | ENSG00000119689 | 272,9179651 |
| UBE2E1 | ENSG00000170142 | 242,9362409 |
| ZFP91 | ENSG00000186660 | 293,7596669 |
| CFP | ENSG00000126759 | 224,6404143 |
| GMFB | ENSG00000197045 | 327,6108953 |
| VPS13D | ENSG00000048707 | 283,2055632 |
| PEF1 | ENSG00000162517 | 269,7088892 |
| ZFTRAF1 | ENSG00000187954 | 256,5503228 |
| PIP4K2A | ENSG00000150867 | 257,8589813 |
| ARIH2 | ENSG00000177479 | 270,6608049 |
| RBM12 | ENSG00000244462 | 254,5258514 |
| XIST | ENSG00000229807 | 216,2832666 |
| DHX15 | ENSG00000109606 | 331,2859855 |
| CHD8 | ENSG00000100888 | 247,6758351 |
| CHMP3 | ENSG00000115561 | 261,8627465 |
| STARD3 | ENSG00000131748 | 262,9584025 |
| CUX1 | ENSG00000257923 | 271,0211906 |
| ATP11A | ENSG00000068650 | 282,1380322 |
| TRIM39 ENSG00000204599 | ENSG00000204599 | 231,9382314 |
| ACVR1B | ENSG00000135503 | 247,5606169 |
| SDF4 | ENSG00000078808 | 196,134276 |
| GSAP | ENSG00000186088 | 325,7816432 |
| COA1 | ENSG00000106603 | 220,5938539 |
| WIPI2 | ENSG00000157954 | 209,2899913 |
| ZNF708 | ENSG00000182141 | 234,0257049 |
| ZNF200 | ENSG00000010539 | 299,0583936 |
| HINT1 | ENSG00000169567 | 196,7367553 |
| BRD1 | ENSG00000100425 | 219,3795648 |
| MOSPD2 | ENSG00000130150 | 308,8767282 |
| TMEM170B | ENSG00000205269 | 310,3613462 |
| DRAM1 | ENSG00000136048 | 209,3006779 |

| | | |
|---|---|---|
| CHUK | ENSG00000213341 | 278,8741788 |
| RNF41 | ENSG00000181852 | 242,1854095 |
| KLRD1 | ENSG00000134539 | 163,9182737 |
| PTPN7 | ENSG00000143851 | 348,3256585 |
| GLA | ENSG00000102393 | 319,5367679 |
| HMGCS1 | ENSG00000112972 | 269,4637596 |
| ZNF148 | ENSG00000163848 | 263,9655344 |
| EIF2D | ENSG00000143486 | 327,816308 |
| ACTR1A | ENSG00000138107 | 257,9273779 |
| NUCB1 | ENSG00000104805 | 297,140108 |
| SGTB | ENSG00000197860 | 230,924405 |
| PDE4D | ENSG00000113448 | 237,5469012 |
| FRYL | ENSG00000075539 | 279,5228971 |
| HECTD1 | ENSG00000092148 | 233,7165043 |
| COL18A1 | ENSG00000182871 | 253,9178877 |
| WDFY4 | ENSG00000128815 | 260,3089128 |
| TENT2 | ENSG00000164329 | 302,7865158 |
| APOL1 | ENSG00000100342 | 190,6509047 |
| IRF5 | ENSG00000128604 | 226,3024677 |
| RABGAP1L | ENSG00000152061 | 278,7458082 |
| FLI1 | ENSG00000151702 | 257,3252402 |
| LMTK2 | ENSG00000164715 | 278,4524712 |
| MAPK8 | ENSG00000107643 | 329,5939849 |
| MARS1 | ENSG00000166986 | 277,8011855 |
| SMURF1 | ENSG00000198742 | 279,4205163 |
| TREML2 | ENSG00000112195 | 379,0582712 |
| CFAP58-DT | ENSG00000231233 | 216,1591332 |
| GPX4 | ENSG00000167468 | 192,9751198 |
| MSL3 | ENSG00000005302 | 363,1628579 |
| PCNX3 | ENSG00000197136 | 247,2185115 |
| ZBTB21 | ENSG00000173276 | 250,0269515 |
| TRAF6 | ENSG00000175104 | 289,9830185 |
| RBCK1 | ENSG00000125826 | 250,1286369 |
| POGZ | ENSG00000143442 | 294,620374 |
| TP53BP1 | ENSG00000067369 | 333,3990973 |
| PPFIA1 | ENSG00000131626 | 256,8912709 |
| DNAJB12 | ENSG00000148719 | 265,3221438 |
| ZBTB43 | ENSG00000169155 | 366,4780562 |
| AHCYL1 | ENSG00000168710 | 299,7839587 |
| ENSG00000130723 | ENSG00000130723 | 195,2122465 |
| IRF9 ENSG00000213928 | ENSG00000213928 | 326,5315353 |
| METTL21A | ENSG00000144401 | 193,6850061 |
| WDTC1 | ENSG00000142784 | 209,9746047 |
| ENSG00000228327 | ENSG00000228327 | 312,6817339 |
| TUBA1A | ENSG00000167552 | 300,5142647 |
| ENSG00000250274 | ENSG00000250274 | 271,9976973 |
| CNP | ENSG00000173786 | 165,4057963 |
| ISCU | ENSG00000136003 | 290,2336692 |
| MACF1 | ENSG00000127603 | 304,1632016 |
| PPP2R3C | ENSG00000092020 | 290,9199075 |
| CHPF2 | ENSG00000033100 | 206,6218476 |
| SREBF2 | ENSG00000198911 | 192,2355755 |
| EMILIN2 | ENSG00000132205 | 228,3572715 |
| ZBTB37 | ENSG00000185278 | 285,0655713 |
| CD6 | ENSG00000013725 | 171,0667567 |

| | | |
|---|---|---|
| CTNNA1 | ENSG00000044115 | 305,0882691 |
| LMO2 | ENSG00000135363 | 210,7018698 |
| TRA2B | ENSG00000136527 | 287,3682841 |
| SPAST | ENSG00000021574 | 324,6818681 |
| CRLF3 | ENSG00000176390 | 260,1843898 |
| HNRNPR ENSG00000125944 | ENSG00000125944 | 226,5761354 |
| KRT23 ENSG00000108244 | ENSG00000108244 | 192,9484488 |
| BTN2A2 | ENSG00000124508 | 231,4854793 |
| RNMT | ENSG00000101654 | 223,1940814 |
| KMT2B | ENSG00000272333 | 201,7075136 |
| PRPF38A | ENSG00000134748 | 283,0977439 |
| STRAP | ENSG00000023734 | 289,9903767 |
| KANSL1 ENSG00000120071 | ENSG00000120071 | 228,5986003 |
| UBN2 | ENSG00000157741 | 223,6608078 |
| NAP1L4 ENSG00000205531 | ENSG00000205531 | 173,2577448 |
| LPAR2 | ENSG00000064547 | 255,0529672 |
| GNB4 | ENSG00000114450 | 206,6188723 |
| ASXL1 | ENSG00000171456 | 198,9808767 |
| NPC2 | ENSG00000119655 | 237,5609249 |
| PCM1 | ENSG00000078674 | 264,0398112 |
| STX5 | ENSG00000162236 | 194,448487 |
| WBP11 | ENSG00000084463 | 232,8928792 |
| GAS7 | ENSG00000007237 | 196,3685166 |
| ADAM19 | ENSG00000135074 | 295,2837356 |
| SF3A3 | ENSG00000183431 | 275,3630248 |
| CTSZ | ENSG00000101160 | 276,2168949 |
| EEF1D ENSG00000104529 | ENSG00000104529 | 238,1069662 |
| DEGS1 | ENSG00000143753 | 253,7619954 |
| SERPINB8 | ENSG00000166401 | 238,6174862 |
| MARCHF3 | ENSG00000173926 | 231,113714 |
| ESRRA | ENSG00000173153 | 191,308625 |
| ZNF12 | ENSG00000164631 | 185,7967685 |
| ENSG00000225886 | ENSG00000225886 | 144,4421748 |
| AKT1 | ENSG00000142208 | 239,2461624 |
| CSGALNACT1 | ENSG00000147408 | 186,0189236 |
| BTN3A2 | ENSG00000186470 | 177,612406 |
| IDH2 | ENSG00000182054 | 166,8780949 |
| GANAB | ENSG00000089597 | 202,2787161 |
| HGS | ENSG00000185359 | 203,9188761 |
| EEIG2 | ENSG00000162636 | 213,1446093 |
| TNFAIP8 | ENSG00000145779 | 220,2960997 |
| RPS23 | ENSG00000186468 | 200,5939623 |
| DAPK1 | ENSG00000196730 | 167,1204775 |
| RHOB | ENSG00000143878 | 242,8324613 |
| SCAF8 | ENSG00000213079 | 268,312708 |
| LNPEP | ENSG00000113441 | 248,8202181 |
| ZNF654 | ENSG00000175105 | 236,1650322 |
| CAPNS1 | ENSG00000126247 | 231,6594952 |
| STX16 | ENSG00000124222 | 326,3776765 |
| OTUD1 | ENSG00000165312 | 160,0501568 |
| RNF167 | ENSG00000108523 | 253,995464 |
| SEC61A1 | ENSG00000058262 | 203,5128742 |
| KIAA0040 | ENSG00000235750 | 294,7146181 |
| SENP6 | ENSG00000112701 | 320,4343997 |
| POLM | ENSG00000122678 | 239,355402 |

| | | |
|---|---|---|
| DGLUCY | ENSG00000133943 | 217,7985813 |
| RFTN1 | ENSG00000131378 | 181,445103 |
| POLDIP3 | ENSG00000100227 | 216,632966 |
| ENSG00000279159 | ENSG00000279159 | 276,6246616 |
| SLC36A4 | ENSG00000180773 | 275,7437956 |
| DNAJC8 | ENSG00000126698 | 265,3402811 |
| ARF3 | ENSG00000134287 | 229,8493891 |
| PPP2R5A | ENSG00000066027 | 208,8054131 |
| POLR2B | ENSG00000047315 | 245,273649 |
| CDK16 | ENSG00000102225 | 271,1547786 |
| RNF38 | ENSG00000137075 | 236,8620297 |
| PARVG | ENSG00000138964 | 259,3132873 |
| AVL9 | ENSG00000105778 | 222,3067713 |
| OTULIN | ENSG00000154124 | 219,4709193 |
| TAF1D | ENSG00000166012 | 298,9493974 |
| RREB1 | ENSG00000124782 | 201,6439447 |
| NEU1 ENSG00000204386 | ENSG00000204386 | 196,6091991 |
| SERBP1 | ENSG00000142864 | 206,4277583 |
| RGS18 | ENSG00000150681 | 280,331502 |
| NSD3 | ENSG00000147548 | 242,8140512 |
| SRSF4 | ENSG00000116350 | 241,242662 |
| LPP | ENSG00000145012 | 273,6074572 |
| SRSF6 | ENSG00000124193 | 249,2909275 |
| JDP2 | ENSG00000140044 | 230,7292243 |
| VPS37B | ENSG00000139722 | 222,247231 |
| PRKAA1 | ENSG00000132356 | 302,8302106 |
| PFKFB2 | ENSG00000123836 | 340,8976164 |
| KLHL36 | ENSG00000135686 | 195,0986614 |
| JMY | ENSG00000152409 | 206,8288732 |
| ARL8A | ENSG00000143862 | 305,5789697 |
| ATG13 | ENSG00000175224 | 176,8375951 |
| IKZF3 | ENSG00000161405 | 174,883051 |
| BET1L | ENSG00000177951 | 212,9007466 |
| TMEM167A | ENSG00000174695 | 260,5034916 |
| HM13 | ENSG00000101294 | 226,8275945 |
| TRIR | ENSG00000123144 | 194,4866173 |
| CLASP1 | ENSG00000074054 | 266,3033392 |
| C5orf58 | ENSG00000234511 | 341,943093 |
| H2BC4 | ENSG00000180596 | 222,9528144 |
| SENP2 | ENSG00000163904 | 236,9637848 |
| LGALS9 | ENSG00000168961 | 230,3896059 |
| SZRD1 | ENSG00000055070 | 207,3424411 |
| ENSG00000280138 | ENSG00000280138 | 285,2860132 |
| G6PD | ENSG00000160211 | 194,013282 |
| PIK3C2A | ENSG00000011405 | 251,8803969 |
| MANBA | ENSG00000109323 | 286,4476245 |
| TUBGCP2 | ENSG00000130640 | 254,3998494 |
| DTX2 ENSG00000091073 | ENSG00000091073 | 226,9335734 |
| P2RY10 | ENSG00000078589 | 220,1073722 |
| CAPN2 | ENSG00000162909 | 200,8845932 |
| AKT2 | ENSG00000105221 | 210,5659432 |
| PSMD4 | ENSG00000159352 | 255,0067362 |
| SLC49A4 | ENSG00000138463 | 250,0535183 |
| LATS2 | ENSG00000150457 | 259,5615188 |
| ZER1 | ENSG00000160445 | 144,5917836 |

| | | |
|---|---|---|
| NRIP1 | ENSG00000180530 | 256,2420324 |
| PELI2 | ENSG00000139946 | 341,4882834 |
| YTHDC2 | ENSG00000047188 | 327,589366 |
| MNT | ENSG00000070444 | 194,1440386 |
| ZNF33A | ENSG00000189180 | 205,4557262 |
| APBB3 | ENSG00000113108 | 257,51393 |
| KIDINS220 | ENSG00000134313 | 270,7185829 |
| PTBP1 | ENSG00000011304 | 187,1611188 |
| RPS15A | ENSG00000134419 | 199,9184527 |
| RORA | ENSG00000069667 | 159,8610268 |
| ATG12 | ENSG00000145782 | 241,2226495 |
| CD9 | ENSG00000010278 | 175,9370582 |
| TRIM28 | ENSG00000130726 | 166,2147219 |
| SENP5 | ENSG00000119231 | 233,2694065 |
| TADA2B | ENSG00000173011 | 147,8588708 |
| DEF6 | ENSG00000023892 | 247,0475702 |
| ACSL5 | ENSG00000197142 | 202,3258596 |
| MAP4K2 | ENSG00000168067 | 202,3286281 |
| SECISBP2L | ENSG00000138593 | 254,2173015 |
| NPAT | ENSG00000149308 | 230,6115582 |
| ERV3-1 | ENSG00000213462 | 244,313451 |
| PRPF38B | ENSG00000134186 | 250,9466237 |
| UBTF | ENSG00000108312 | 175,7642349 |
| LEMD3 | ENSG00000174106 | 235,422275 |
| SNX3 | ENSG00000112335 | 300,7835201 |
| BTBD19 | ENSG00000222009 | 293,6806775 |
| HLA-DQB1 ENSG00000179344 | ENSG00000179344 | 127,700629 |
| CA4 | ENSG00000167434 | 281,5836134 |
| LIG4 | ENSG00000174405 | 285,2240839 |
| HSPA4 | ENSG00000170606 | 204,0815834 |
| GALNT3 | ENSG00000115339 | 269,7270566 |
| GGA3 | ENSG00000125447 | 221,0590773 |
| CAPRIN1 | ENSG00000135387 | 215,2444981 |
| ZBTB11 | ENSG00000066422 | 247,322685 |
| TTC3 | ENSG00000182670 | 194,9393229 |
| PHLDA1 | ENSG00000139289 | 336,5807194 |
| POMP | ENSG00000132963 | 298,5509022 |
| UBAC2 | ENSG00000134882 | 167,2564326 |
| CAMTA2 | ENSG00000108509 | 244,7169251 |
| PKN1 | ENSG00000123143 | 192,9102211 |
| CALCOCO1 | ENSG00000012822 | 230,9207069 |
| STAT4 | ENSG00000138378 | 202,3242354 |
| CD59 | ENSG00000085063 | 246,821826 |
| EIF4A3 | ENSG00000141543 | 244,8139853 |
| HK3 | ENSG00000160883 | 214,6720301 |
| UQCRB | ENSG00000156467 | 259,6532799 |
| GLTP | ENSG00000139433 | 207,9988182 |
| DIDO1 | ENSG00000101191 | 187,7330019 |
| SMAD4 | ENSG00000141646 | 236,3161109 |
| AOAH | ENSG00000136250 | 290,7818452 |
| DGKZ | ENSG00000149091 | 179,5157751 |
| TP53I11 | ENSG00000175274 | 192,9522974 |
| BMAL1 | ENSG00000133794 | 292,193745 |
| PRKAG2 | ENSG00000106617 | 209,6346249 |
| SLAIN2 | ENSG00000109171 | 219,2344375 |

| | | |
|---|---|---|
| UBXN1 | ENSG00000162191 | 215,9140866 |
| CYBC1 | ENSG00000178927 | 203,7440296 |
| CCDC47 | ENSG00000108588 | 227,8527254 |
| ITK | ENSG00000113263 | 241,1711014 |
| PLCL2 ENSG00000154822 | ENSG00000154822 | 248,5932422 |
| GNL1 ENSG00000204590 | ENSG00000204590 | 229,9263389 |
| NCBP3 | ENSG00000074356 | 247,1203851 |
| ANKFY1 | ENSG00000185722 | 265,9686849 |
| CCNH | ENSG00000134480 | 348,6158295 |
| TMEM33 | ENSG00000109133 | 249,6137918 |
| BPTF | ENSG00000171634 | 233,0198417 |
| STX12 | ENSG00000117758 | 333,4328559 |
| RPL22 | ENSG00000116251 | 188,0338886 |
| CDC5L | ENSG00000096401 | 246,8317676 |
| SLBP | ENSG00000163950 | 231,675175 |
| CHTOP | ENSG00000160679 | 175,6800384 |
| AP3D1 | ENSG00000065000 | 186,2105438 |
| INO80D ENSG00000114933 | ENSG00000114933 | 255,9536894 |
| CBLL1 | ENSG00000105879 | 184,6170893 |
| PTPN11 | ENSG00000179295 | 220,5919654 |
| SMAD2 | ENSG00000175387 | 228,1450657 |
| EZH1 | ENSG00000108799 | 202,4714759 |
| ARAP2 | ENSG00000047365 | 237,9448405 |
| STXBP5 | ENSG00000164506 | 277,3143096 |
| PSMB3 ENSG00000277791 | ENSG00000277791 | 206,6133467 |
| TMEM170A | ENSG00000166822 | 208,8406655 |
| GBF1 | ENSG00000107862 | 209,0745329 |
| SLC43A3 | ENSG00000134802 | 218,1035144 |
| EIF4G3 | ENSG00000075151 | 281,8909553 |
| RAB3GAP1 | ENSG00000115839 | 287,858809 |
| ODC1 | ENSG00000115758 | 157,3333221 |
| EIF3L | ENSG00000100129 | 167,1124649 |
| DLD | ENSG00000091140 | 223,7734982 |
| PPP6R1 | ENSG00000105063 | 175,2286344 |
| U2SURP | ENSG00000163714 | 233,1856457 |
| ATXN2L | ENSG00000168488 | 253,5583934 |
| UBA6 | ENSG00000033178 | 313,0984036 |
| DOCK10 | ENSG00000135905 | 177,2048924 |
| TFG | ENSG00000114354 | 259,5399125 |
| ZNF429 | ENSG00000197013 | 235,8558641 |
| USP24 | ENSG00000162402 | 220,2281114 |
| TSPYL2 | ENSG00000184205 | 234,6020392 |
| CST7 | ENSG00000077984 | 222,0417834 |
| FAM86B3P ENSG00000173295 | ENSG00000173295 | 178,9418793 |
| MTDH | ENSG00000147649 | 183,1902489 |
| SUMO2 | ENSG00000188612 | 246,204879 |
| TMC6 | ENSG00000141524 | 217,5479002 |
| MYO5A | ENSG00000197535 | 241,978857 |
| PRKAR2A | ENSG00000114302 | 212,4557009 |
| ATM | ENSG00000149311 | 199,4934381 |
| GSK3A | ENSG00000105723 | 153,7062937 |
| SMARCC1 | ENSG00000173473 | 234,3993863 |
| PLIN3 | ENSG00000105355 | 200,5375413 |
| TRBC2 ENSG00000211772 | ENSG00000211772 | 148,2692992 |
| LBH | ENSG00000213626 | 126,1664496 |

| | | |
|---|---|---|
| DEF8 | ENSG00000140995 | 183,0233538 |
| BCL11B | ENSG00000127152 | 151,2180025 |
| LDB1 | ENSG00000198728 | 166,9934011 |
| RAB6A | ENSG00000175582 | 247,6977765 |
| NARF | ENSG00000141562 | 217,3630376 |
| SNX13 | ENSG00000071189 | 285,9214827 |
| TIFA | ENSG00000145365 | 175,434802 |
| RALGAPA2 | ENSG00000188559 | 283,0553177 |
| CTR9 | ENSG00000198730 | 214,3239899 |
| RNF4 | ENSG00000063978 | 159,4768914 |
| NFYC | ENSG00000066136 | 226,8078055 |
| RASGEF1B | ENSG00000138670 | 203,4562632 |
| CASP9 | ENSG00000132906 | 181,2747483 |
| CC2D1B | ENSG00000154222 | 240,1929775 |
| ZNF652 | ENSG00000198740 | 178,718848 |
| SRSF7 | ENSG00000115875 | 208,2637733 |
| SRC | ENSG00000197122 | 145,7463461 |
| BECN1 | ENSG00000126581 | 242,9403087 |
| ZC3H15 | ENSG00000065548 | 226,6231571 |
| SRF | ENSG00000112658 | 183,7937252 |
| CD48 | ENSG00000117091 | 193,6474343 |
| NSFL1C | ENSG00000088833 | 169,8098653 |
| TMEM164 | ENSG00000157600 | 175,6725925 |
| ZMYND15 | ENSG00000141497 | 173,1472946 |
| RPL36 | ENSG00000130255 | 148,975801 |
| CXorf38 | ENSG00000185753 | 214,9240321 |
| ARL6IP1 | ENSG00000170540 | 223,7904193 |
| CNTNAP3 | ENSG00000106714 | 329,9872161 |
| VPS13C | ENSG00000129003 | 205,0981756 |
| NPTN | ENSG00000156642 | 269,9234803 |
| MFSD12 | ENSG00000161091 | 213,2168771 |
| CCDC117 | ENSG00000159873 | 193,2415278 |
| ITPR1 | ENSG00000150995 | 180,9377758 |
| KMT5A | ENSG00000183955 | 249,3261495 |
| ZBTB44 | ENSG00000196323 | 201,543467 |
| STAM2 | ENSG00000115145 | 227,6069921 |
| XAB2 | ENSG00000076924 | 199,9670336 |
| PLIN4 | ENSG00000167676 | 252,9128198 |
| BATF2 | ENSG00000168062 | 103,9321141 |
| ARID2 | ENSG00000189079 | 208,6935765 |
| SLTM | ENSG00000137776 | 214,4488237 |
| VIM-AS1 | ENSG00000229124 | 163,7817323 |
| UBE2D1 | ENSG00000072401 | 315,2473229 |
| CERS2 | ENSG00000143418 | 153,9011629 |
| FAM168B | ENSG00000152102 | 211,6872518 |
| PRDX6 | ENSG00000117592 | 213,922993 |
| SLC35E1 | ENSG00000127526 | 150,343996 |
| SRSF10 | ENSG00000188529 | 230,8374851 |
| MYCBP2 | ENSG00000005810 | 242,8987608 |
| PRKX | ENSG00000183943 | 172,0931023 |
| BICRAL | ENSG00000112624 | 231,3207226 |
| DRAP1 | ENSG00000175550 | 189,3072973 |
| POLG | ENSG00000140521 | 158,864068 |
| ITGA4 | ENSG00000115232 | 193,5419003 |
| SECISBP2 | ENSG00000187742 | 237,5617992 |

| | | |
|---|---|---|
| FXYD5 | ENSG00000089327 | 165,8083544 |
| SUZ12 | ENSG00000178691 | 235,8245422 |
| HDAC5 | ENSG00000108840 | 132,8557047 |
| STX17 | ENSG00000136874 | 269,5495257 |
| EEPD1 | ENSG00000122547 | 165,5550331 |
| NCR3LG1 | ENSG00000188211 | 126,9973907 |
| INTS6 | ENSG00000102786 | 227,7880703 |
| ZNF106 | ENSG00000103994 | 214,7190154 |
| BCKDK | ENSG00000103507 | 204,5314208 |
| OSCAR ENSG00000170909 | ENSG00000170909 | 189,1224818 |
| TJAP1 | ENSG00000137221 | 178,6475782 |
| NR4A1 | ENSG00000123358 | 210,7136018 |
| HADHA | ENSG00000084754 | 168,5289718 |
| SIRT7 | ENSG00000187531 | 180,1890627 |
| XYLT1 ENSG00000103489 | ENSG00000103489 | 138,8348939 |
| SLC40A1 | ENSG00000138449 | 138,4443125 |
| APH1A | ENSG00000117362 | 196,2531085 |
| ADARB1 | ENSG00000197381 | 217,2258004 |
| G3BP1 | ENSG00000145907 | 174,8014326 |
| CPSF2 | ENSG00000165934 | 155,400237 |
| SYNCRIP | ENSG00000135316 | 198,1677541 |
| CLINT1 | ENSG00000113282 | 251,3282719 |
| KLHL6 | ENSG00000172578 | 241,0180801 |
| FOXJ3 | ENSG00000198815 | 232,3180102 |
| UGP2 | ENSG00000169764 | 230,924095 |
| LTBR | ENSG00000111321 | 237,7955595 |
| HMGCR | ENSG00000113161 | 203,0722132 |
| AMD1 | ENSG00000123505 | 193,0569751 |
| PPP1CA | ENSG00000172531 | 142,4806098 |
| CLEC5A | ENSG00000258227 | 189,1227553 |
| ZNF638 | ENSG00000075292 | 222,3759626 |
| TWF1 | ENSG00000151239 | 228,2548292 |
| FNBP4 ENSG00000109920 | ENSG00000109920 | 192,5156591 |
| POLR2E | ENSG00000099817 | 167,5804066 |
| RBMX | ENSG00000147274 | 201,1225551 |
| GAPVD1 | ENSG00000165219 | 231,0124256 |
| PIGA | ENSG00000165195 | 272,8734715 |
| ST13 | ENSG00000100380 | 188,7518454 |
| SMIM29 | ENSG00000186577 | 185,281333 |
| IFI35 | ENSG00000068079 | 150,078301 |
| PSTPIP1 | ENSG00000140368 | 177,0889237 |
| SGPL1 | ENSG00000166224 | 136,6230954 |
| EEIG1 | ENSG00000167106 | 127,5500783 |
| TGFBR3 | ENSG00000069702 | 101,3824961 |
| IKZF5 | ENSG00000095574 | 192,0964271 |
| DPEP2 | ENSG00000167261 | 177,2887308 |
| SLC35F5 | ENSG00000115084 | 232,4508507 |
| MXRA7 | ENSG00000182534 | 148,9950142 |
| ATP2B4 | ENSG00000058668 | 182,2557007 |
| CS | ENSG00000062485 | 235,3089769 |
| RHOH | ENSG00000168421 | 288,3916076 |
| NHERF1 | ENSG00000109062 | 172,181806 |
| CD247 | ENSG00000198821 | 129,8873753 |
| APPL2 | ENSG00000136044 | 223,1243187 |
| CLDND1 | ENSG00000080822 | 243,6354247 |

| | | |
|---|---|---|
| DMTF1 | ENSG00000135164 | 214,797357 |
| RIC8A | ENSG00000177963 | 178,880128 |
| SEC24D | ENSG00000150961 | 247,0994894 |
| PIK3R4 | ENSG00000196455 | 183,8279406 |
| TCF7 | ENSG00000081059 | 178,9196258 |
| RPA2 | ENSG00000117748 | 236,5206331 |
| MPHOSPH10P1 | ENSG00000260078 | 234,6085928 |
| CWC22 | ENSG00000163510 | 231,2674877 |
| FAM199X | ENSG00000123575 | 233,2510417 |
| CSF1 | ENSG00000184371 | 216,6187851 |
| C3 | ENSG00000125730 | 122,9017367 |
| PRR14L | ENSG00000183530 | 164,9085149 |
| LYPLA1 | ENSG00000120992 | 243,3046347 |
| ZFR | ENSG00000056097 | 193,7619965 |
| PPP2R2D | ENSG00000175470 | 189,10191 |
| TWF2 | ENSG00000247596 | 175,0385444 |
| PUM1 | ENSG00000134644 | 206,5070287 |
| POFUT2 | ENSG00000186866 | 221,5541052 |
| PPM1M | ENSG00000164088 | 258,6544417 |
| PPP2R2A | ENSG00000221914 | 267,0652143 |
| VPS37C | ENSG00000167987 | 160,3045329 |
| SIPA1L2 | ENSG00000116991 | 233,5396676 |
| ATP5F1A | ENSG00000152234 | 149,9786028 |
| LY96 | ENSG00000154589 | 226,9055088 |
| FUT4 | ENSG00000196371 | 156,2677645 |
| NCOA7 | ENSG00000111912 | 148,9708764 |
| MTATP6P1 | ENSG00000248527 | 243,8170613 |
| H3P6 | ENSG00000235655 | 187,328801 |
| NRIP3 | ENSG00000175352 | 257,7429462 |
| ISG20L2 | ENSG00000143319 | 194,0174367 |
| CD2BP2 | ENSG00000169217 | 171,7633662 |
| UBA3 | ENSG00000144744 | 198,0987078 |
| PRP4K | ENSG00000112739 | 206,4937275 |
| JPT1 | ENSG00000189159 | 242,9013097 |
| DGKA | ENSG00000065357 | 244,0934253 |
| GK-AS1 | ENSG00000243055 | 297,2361408 |
| OAT | ENSG00000065154 | 261,6892754 |
| LTN1 | ENSG00000198862 | 222,5733605 |
| WASH5P | ENSG00000282458 | 202,8571698 |
| GRK5 | ENSG00000198873 | 180,2617347 |
| PEA15 | ENSG00000162734 | 156,5232284 |
| BCAS2 | ENSG00000116752 | 204,3679273 |
| TNFSF8 | ENSG00000106952 | 180,3150053 |
| TOMM20 | ENSG00000173726 | 198,506217 |
| DYNC1LI1 | ENSG00000144635 | 172,6557443 |
| CDK9 | ENSG00000136807 | 188,5954696 |
| BLTP2 | ENSG00000007202 | 190,0052591 |
| SLC12A9 | ENSG00000146828 | 207,972472 |
| GCC1 | ENSG00000179562 | 154,9839477 |
| USP38 | ENSG00000170185 | 213,6704731 |
| FAM53B | ENSG00000189319 | 95,21280265 |
| RNASEL | ENSG00000135828 | 184,0320698 |
| CHIC2 | ENSG00000109220 | 206,1236167 |
| TLK2 | ENSG00000146872 | 179,8831054 |
| MICAL2 | ENSG00000133816 | 177,1035373 |

| | | |
|---|---|---|
| GID8 | ENSG00000101193 | 142,374216 |
| MBTPS1 | ENSG00000140943 | 197,2856115 |
| RSBN1L | ENSG00000187257 | 179,7518722 |
| F5 | ENSG00000198734 | 216,271493 |
| PSME2 ENSG00000100911 | ENSG00000100911 | 206,4846985 |
| TRAK1 | ENSG00000182606 | 233,6011004 |
| DDX27 | ENSG00000124228 | 210,2162765 |
| IGHM ENSG00000211899 | ENSG00000211899 | 163,7771249 |
| CYREN | ENSG00000122783 | 174,036268 |
| PIP4P1 | ENSG00000165782 | 178,2666349 |
| ASCC1 | ENSG00000138303 | 184,9832318 |
| GSK3B | ENSG00000082701 | 212,0487828 |
| RASAL3 | ENSG00000105122 | 190,7552373 |
| SLA2 | ENSG00000101082 | 132,7155459 |
| AAGAB | ENSG00000103591 | 114,7829287 |
| IL18BP | ENSG00000137496 | 156,0371437 |
| GK5 | ENSG00000175066 | 229,2664781 |
| MBD4 | ENSG00000129071 | 241,1555236 |
| DNAH17 | ENSG00000187775 | 268,9598022 |
| CSTA | ENSG00000121552 | 220,8519145 |
| EMC3 | ENSG00000125037 | 157,4942802 |
| TBC1D10C | ENSG00000175463 | 143,6233367 |
| SAR1B | ENSG00000152700 | 194,3979524 |
| ENSG00000241886 | ENSG00000241886 | 256,5461057 |
| TXNL1 | ENSG00000091164 | 217,5426772 |
| MGAT4A | ENSG00000071073 | 202,2905368 |
| TPP2 | ENSG00000134900 | 226,005114 |
| HDAC7 | ENSG00000061273 | 199,289055 |
| ECD | ENSG00000122882 | 223,1835433 |
| TLK1 | ENSG00000198586 | 187,097972 |
| DPP8 | ENSG00000074603 | 181,0522726 |
| MAP3K5 | ENSG00000197442 | 293,9499035 |
| PLEKHM3 | ENSG00000178385 | 164,9742918 |
| PCIF1 | ENSG00000100982 | 177,2544171 |
| EIF3F | ENSG00000175390 | 152,216694 |
| REV3L | ENSG00000009413 | 221,2721975 |
| EBLN3P | ENSG00000281649 | 194,902985 |
| GIGYF1 | ENSG00000146830 | 159,2284683 |
| MAGOH | ENSG00000162385 | 263,9743181 |
| EVL | ENSG00000196405 | 160,3240932 |
| ARSG | ENSG00000141337 | 141,7861831 |
| NFATC2 | ENSG00000101096 | 145,2297996 |
| NRAS | ENSG00000213281 | 177,597729 |
| ZBTB2 | ENSG00000181472 | 168,7855706 |
| MEF2C | ENSG00000081189 | 196,2100123 |
| CDK13 | ENSG00000065883 | 180,4667476 |
| DUSP4 | ENSG00000120875 | 133,8483677 |
| RABEP1 | ENSG00000029725 | 199,8801335 |
| WWC3 | ENSG00000047644 | 205,03541 |
| RCC2 ENSG00000179051 | ENSG00000179051 | 193,7596491 |
| RBM17 | ENSG00000134453 | 261,6141961 |
| PDE4A | ENSG00000065989 | 181,4280027 |
| LAMTOR1 | ENSG00000149357 | 150,7832205 |
| ZXDC | ENSG00000070476 | 188,3931034 |
| HAVCR2 | ENSG00000135077 | 192,9596272 |

| | | |
|---|---|---|
| MBD1 | ENSG00000141644 | 194,8503121 |
| PHF23 | ENSG00000040633 | 193,5989901 |
| SLC25A44 | ENSG00000160785 | 201,8184411 |
| BCL2L13 | ENSG00000099968 | 193,3983086 |
| FAM110A | ENSG00000125898 | 128,8501714 |
| SHFL | ENSG00000130813 | 157,7456577 |
| SCYL1 | ENSG00000142186 | 163,2696788 |
| ZBTB25 | ENSG00000089775 | 132,7093225 |
| SP140L | ENSG00000185404 | 206,7311201 |
| BIN3 | ENSG00000147439 | 220,2123021 |
| RGS3 | ENSG00000138835 | 140,3461083 |
| RBM8A | ENSG00000265241 | 199,3235307 |
| GMCL1 | ENSG00000087338 | 181,4586213 |
| PCTP | ENSG00000141179 | 185,4713242 |
| CARD8-AS1 | ENSG00000268001 | 159,6898646 |
| FAF2 | ENSG00000113194 | 195,3446093 |
| SPOP | ENSG00000121067 | 174,8072834 |
| CNEP1R1 | ENSG00000205423 | 185,3208849 |
| ARHGEF6 | ENSG00000129675 | 221,4735785 |
| KAT8 | ENSG00000103510 | 200,0257081 |
| MPHOSPH8 | ENSG00000196199 | 181,5171462 |
| NBPF14 | ENSG00000270629 | 164,411902 |
| NKG7 | ENSG00000105374 | 90,39145942 |
| COPG1 | ENSG00000181789 | 175,8936655 |
| CTCF | ENSG00000102974 | 123,0899561 |
| ABCA2 | ENSG00000107331 | 151,9693239 |
| B3GNTL1 ENSG00000175711 | ENSG00000175711 | 233,3092813 |
| GON4L | ENSG00000116580 | 200,3988239 |
| SNX1 | ENSG00000028528 | 177,3521742 |
| CST3 | ENSG00000101439 | 140,2508522 |
| HLA-DQA1 ENSG00000196735 | ENSG00000196735 | 110,891259 |
| CAMK2G | ENSG00000148660 | 148,9066008 |
| ZNF100 ENSG00000197020 | ENSG00000197020 | 177,509087 |
| P4HA1 | ENSG00000122884 | 165,937473 |
| TNFRSF9 | ENSG00000049249 | 122,097002 |
| UPB1 | ENSG00000100024 | 167,9968501 |
| NUDT4 | ENSG00000173598 | 170,5785693 |
| DDX50 | ENSG00000107625 | 176,420841 |
| SPN | ENSG00000197471 | 123,6897142 |
| FLCN | ENSG00000154803 | 141,3938867 |
| ZDHHC17 | ENSG00000186908 | 214,5647536 |
| MPC1 | ENSG00000060762 | 158,1544617 |
| CMPK1 | ENSG00000162368 | 184,7631331 |
| ARMC8 | ENSG00000114098 | 257,3975684 |
| FOXP1 | ENSG00000114861 | 188,1561675 |
| SMURF2 | ENSG00000108854 | 206,1811234 |
| HMGN4 | ENSG00000182952 | 141,5775444 |
| GNPTAB | ENSG00000111670 | 158,8670186 |
| ATF6 | ENSG00000118217 | 224,8246671 |
| PSMB8-AS1 ENSG00000204261 | ENSG00000204261 | 169,3398615 |
| ATF7IP | ENSG00000171681 | 139,0553567 |
| PDS5A | ENSG00000121892 | 191,2441059 |
| ELK4 | ENSG00000158711 | 205,5991787 |
| RAB24 | ENSG00000169228 | 156,1419024 |
| NUCKS1 | ENSG00000069275 | 167,2752344 |

| | | |
|---|---|---|
| LRWD1 | ENSG00000161036 | 133,579313 |
| GARS1-DT | ENSG00000196295 | 226,4444889 |
| GAB3 | ENSG00000160219 | 220,2334595 |
| CD5 | ENSG00000110448 | 107,7328226 |
| SUPT20H | ENSG00000102710 | 185,6785678 |
| FAM118A | ENSG00000100376 | 221,6898246 |
| RAB32 | ENSG00000118508 | 161,1975038 |
| TLE5 | ENSG00000104964 | 137,6138718 |
| SRP9 | ENSG00000143742 | 181,0033159 |
| MALT1 | ENSG00000172175 | 213,0382292 |
| BUD31 | ENSG00000106245 | 183,2569965 |
| BBX | ENSG00000114439 | 222,472762 |
| IP6K2 | ENSG00000068745 | 161,5111733 |
| GAA | ENSG00000171298 | 149,9283284 |
| CMTM3 | ENSG00000140931 | 184,093459 |
| ADGRE1 | ENSG00000174837 | 198,9742441 |
| YAF2 | ENSG00000015153 | 162,1526247 |
| RNF111 | ENSG00000157450 | 218,9366246 |
| COX7A2 | ENSG00000112695 | 165,2090419 |
| ATP6V0A1 | ENSG00000033627 | 203,3574614 |
| TRPM7 | ENSG00000092439 | 176,6609395 |
| CLCN3 | ENSG00000109572 | 195,9373905 |
| PSMD6 | ENSG00000163636 | 188,9627519 |
| RSL24D1 | ENSG00000137876 | 188,3688924 |
| XPR1 | ENSG00000143324 | 189,2774692 |
| EIF3I | ENSG00000084623 | 202,2477493 |
| RAB1B | ENSG00000174903 | 159,1940044 |
| ZKSCAN8 | ENSG00000198315 | 159,7973495 |
| NUP214 | ENSG00000126883 | 158,8549394 |
| KMT5B | ENSG00000110066 | 159,0123197 |
| IRAK4 | ENSG00000198001 | 202,1695954 |
| SSR3 | ENSG00000114850 | 158,1829615 |
| MAPRE2 | ENSG00000166974 | 156,9664216 |
| RER1 | ENSG00000157916 | 176,3961576 |
| PLIN5 | ENSG00000214456 | 212,7852123 |
| SLC7A7 | ENSG00000155465 | 141,2904956 |
| FBXO28 | ENSG00000143756 | 165,8491377 |
| RNF114 | ENSG00000124226 | 180,2569404 |
| JUP | ENSG00000173801 | 98,06741372 |
| TBC1D1 | ENSG00000065882 | 228,3144198 |
| SCAMP2 | ENSG00000140497 | 124,9721571 |
| GABPB1 | ENSG00000104064 | 219,8853761 |
| ENSG00000273837 | ENSG00000273837 | 184,3994369 |
| DUSP3 | ENSG00000108861 | 134,1627382 |
| VAMP7 | ENSG00000124333 | 192,5621364 |
| MAP3K3 | ENSG00000198909 | 199,4729428 |
| RBBP5 | ENSG00000117222 | 160,5608633 |
| IL18R1 | ENSG00000115604 | 264,2180186 |
| AP3B1 | ENSG00000132842 | 234,5366329 |
| HNRNPLL | ENSG00000143889 | 162,4132768 |
| TFIP11 | ENSG00000100109 | 151,0415353 |
| PAPOLG | ENSG00000115421 | 151,8876866 |
| VCL | ENSG00000035403 | 244,6134603 |
| ZNF791 | ENSG00000173875 | 186,7658789 |
| SLC39A13-AS1 | ENSG00000255197 | 245,1637436 |

| | | |
|---|---|---|
| TRBC1 ENSG00000211751 | ENSG00000211751 | 106,6517341 |
| FAM120B ENSG00000112584 | ENSG00000112584 | 141,0379521 |
| RELCH | ENSG00000134444 | 225,8133963 |
| DCUN1D1 | ENSG00000043093 | 209,6137386 |
| LUC7L3 | ENSG00000108848 | 198,2546678 |
| ABHD3 | ENSG00000158201 | 216,4258831 |
| PRR14 | ENSG00000156858 | 169,3795815 |
| MAFB | ENSG00000204103 | 131,3258975 |
| KLHL15 | ENSG00000174010 | 177,0415388 |
| RASGRP1 | ENSG00000172575 | 121,8965072 |
| H2BC18 | ENSG00000203814 | 155,5554259 |
| PTAR1 | ENSG00000188647 | 204,1635886 |
| MAFG | ENSG00000197063 | 226,523553 |
| TP53BP2 | ENSG00000143514 | 201,2993018 |
| SMARCD2 | ENSG00000108604 | 150,4738811 |
| RERE | ENSG00000142599 | 176,9314521 |
| CCND3 | ENSG00000112576 | 171,253182 |
| NBPF9 | ENSG00000269713 | 219,8408349 |
| PRPF3 | ENSG00000117360 | 207,5377729 |
| ARHGAP31 | ENSG00000031081 | 131,8477324 |
| SDF2 | ENSG00000132581 | 201,03194 |
| PI4KB | ENSG00000143393 | 193,4018746 |
| ZFYVE27 | ENSG00000155256 | 149,5252065 |
| KCNH4 | ENSG00000089558 | 143,8121197 |
| LAIR1 ENSG00000167613 | ENSG00000167613 | 164,9227572 |
| MON1B | ENSG00000103111 | 200,8911813 |
| RNPS1 | ENSG00000205937 | 154,317714 |
| ZNF319 | ENSG00000166188 | 146,5444243 |
| OCIAD1 | ENSG00000109180 | 198,5603352 |
| DDX42 | ENSG00000198231 | 155,2196332 |
| SWAP70 | ENSG00000133789 | 178,0027543 |
| UBQLN2 | ENSG00000188021 | 191,0661434 |
| THAP5 | ENSG00000177683 | 171,9949723 |
| MTMR3 | ENSG00000100330 | 178,4417195 |
| CCDC82 | ENSG00000149231 | 191,1379249 |
| TMED4 | ENSG00000158604 | 172,4543811 |
| PPM1F-AS1 | ENSG00000224086 | 111,666896 |
| CTDP1 ENSG00000060069 | ENSG00000060069 | 154,4092325 |
| UBE2I | ENSG00000103275 | 148,9675827 |
| MYO15B | ENSG00000266714 | 224,6865818 |
| UIMC1 | ENSG00000087206 | 186,8277533 |
| HSPA6 | ENSG00000173110 | 189,1599174 |
| DPY19L3 | ENSG00000178904 | 226,8624452 |
| MOB3C | ENSG00000142961 | 149,4934489 |
| SAFB2 | ENSG00000130254 | 154,3149371 |
| TTLL4 | ENSG00000135912 | 165,9874858 |
| ZNF397 | ENSG00000186812 | 210,8456405 |
| MAF1 | ENSG00000179632 | 142,9182327 |
| HELLPAR | ENSG00000281344 | 201,2940331 |
| DPYD | ENSG00000188641 | 199,9229114 |
| HEXIM1 | ENSG00000186834 | 146,2103691 |
| ENSG00000234290 | ENSG00000234290 | 175,682189 |
| ARID3B | ENSG00000179361 | 190,5442548 |
| BRD10 | ENSG00000183354 | 203,4384528 |
| AGAP2 | ENSG00000135439 | 150,4680589 |

| | | |
|---|---|---|
| CD86 | ENSG00000114013 | 187,027105 |
| RTN2 | ENSG00000125744 | 102,4765743 |
| B4GALT4 | ENSG00000121578 | 133,1114176 |
| C2CD3 | ENSG00000168014 | 182,2622508 |
| TAMALIN | ENSG00000161835 | 143,2641655 |
| S1PR4 | ENSG00000125910 | 106,1534945 |
| FAM13B | ENSG00000031003 | 214,2569812 |
| ZBTB17 | ENSG00000116809 | 140,8470248 |
| KPNA6 | ENSG00000025800 | 170,1511949 |
| LRPAP1 | ENSG00000163956 | 172,4063855 |
| RHOT2 | ENSG00000140983 | 145,8516889 |
| MOV10 | ENSG00000155363 | 145,7786064 |
| UAP1 | ENSG00000117143 | 200,0820834 |
| GNPDA1 | ENSG00000113552 | 121,8661998 |
| GNA12 | ENSG00000146535 | 163,3402099 |
| CARINH | ENSG00000197536 | 158,2199754 |
| COPE | ENSG00000105669 | 157,4422877 |
| CDC34 | ENSG00000099804 | 144,1109336 |
| SS18 | ENSG00000141380 | 131,3416502 |
| UFD1 | ENSG00000070010 | 155,3009509 |
| TRAC | ENSG00000277734 | 119,5034207 |
| RBM38 | ENSG00000132819 | 126,7338313 |
| CD300LB | ENSG00000178789 | 198,3505215 |
| XPC | ENSG00000154767 | 119,5128134 |
| ATP6V1D | ENSG00000100554 | 188,683948 |
| DPF2 | ENSG00000133884 | 133,6062804 |
| LRRK2-DT | ENSG00000225342 | 128,9773376 |
| NR1D2 | ENSG00000174738 | 144,8525427 |
| WDR45B | ENSG00000141580 | 159,8553557 |
| PTPN2 | ENSG00000175354 | 195,5760368 |
| SLC30A7 | ENSG00000162695 | 167,4191528 |
| CREBZF | ENSG00000137504 | 165,4136609 |
| CCDC57 | ENSG00000176155 | 188,492243 |
| ENSG00000237094 | ENSG00000237094 | 223,2255418 |
| UBE2G1 | ENSG00000132388 | 185,4142675 |
| LARP7 | ENSG00000174720 | 181,226935 |
| SF3B4 | ENSG00000143368 | 170,8997619 |
| MFNG | ENSG00000100060 | 122,6624091 |
| AP3S1 | ENSG00000177879 | 229,3478787 |
| KAT7 | ENSG00000136504 | 185,2155879 |
| HDGFL3 | ENSG00000166503 | 167,6157366 |
| SFMBT2 | ENSG00000198879 | 166,8243021 |
| HSBP1 | ENSG00000230989 | 175,1506024 |
| SMS | ENSG00000102172 | 194,4074502 |
| DERL2 | ENSG00000072849 | 164,7084302 |
| TMEM131 | ENSG00000075568 | 195,3929691 |
| EDEM3 | ENSG00000116406 | 208,7497363 |
| SIRT2 ENSG00000068903 | ENSG00000068903 | 161,2929274 |
| POLB | ENSG00000070501 | 191,7030445 |
| USP12 | ENSG00000152484 | 180,0406603 |
| PHTF1 | ENSG00000116793 | 242,6188544 |
| SART1 | ENSG00000175467 | 164,6038441 |
| ENSG00000276900 | ENSG00000276900 | 149,459834 |
| ANP32B | ENSG00000136938 | 162,6883947 |
| PSMA3 | ENSG00000100567 | 186,3137003 |

| | | |
|---|---|---|
| MBOAT2 | ENSG00000143797 | 247,3298379 |
| SLC66A2 | ENSG00000122490 | 165,3148565 |
| EPN1 | ENSG00000063245 | 114,8069317 |
| FOXO1 | ENSG00000150907 | 141,7431208 |
| CAPG | ENSG00000042493 | 166,8660471 |
| SNU13 | ENSG00000100138 | 148,6548936 |
| JAK2 | ENSG00000096968 | 205,7221494 |
| MFSD14B | ENSG00000148110 | 211,6547002 |
| MAFK | ENSG00000198517 | 133,2963584 |
| GSTK1 | ENSG00000197448 | 197,0126121 |
| PRPF6 | ENSG00000101161 | 140,1026622 |
| PDLIM5 | ENSG00000163110 | 168,9059134 |
| VAV3 | ENSG00000134215 | 199,01168 |
| RRAGC | ENSG00000116954 | 138,2168634 |
| RAB3GAP2 | ENSG00000118873 | 184,3097851 |
| NFKBIB ENSG00000104825 | ENSG00000104825 | 117,6272065 |
| ZC3H4 | ENSG00000130749 | 139,6494306 |
| PIK3C3 | ENSG00000078142 | 185,0596043 |
| DBN1 | ENSG00000113758 | 157,4924256 |
| USP16 | ENSG00000156256 | 166,4843999 |
| TTYH3 | ENSG00000136295 | 157,0238146 |
| MAN2A1 | ENSG00000112893 | 189,5808082 |
| SUPT16H | ENSG00000092201 | 129,1660123 |
| ZRANB1 | ENSG00000019995 | 171,1716315 |
| AUTS2 | ENSG00000158321 | 85,5905904 |
| ZDHHC3 | ENSG00000163812 | 186,6647982 |
| C10orf55 | ENSG00000222047 | 134,0336782 |
| EPC2 | ENSG00000135999 | 173,9597459 |
| NSD1 | ENSG00000165671 | 155,5274768 |
| DCAF5 | ENSG00000139990 | 190,7033919 |
| ATP2A2 | ENSG00000174437 | 179,1690964 |
| FUBP1 | ENSG00000162613 | 171,3648703 |
| AATF ENSG00000275700 | ENSG00000275700 | 161,8229838 |
| CASP3 | ENSG00000164305 | 154,3831959 |
| EPC1 | ENSG00000120616 | 168,5686347 |
| SFT2D2 | ENSG00000213064 | 157,5359184 |
| NIPA2 | ENSG00000140157 | 180,3373262 |
| YIPF5 | ENSG00000145817 | 146,7437824 |
| GUSB | ENSG00000169919 | 225,8384572 |
| PRKCI | ENSG00000163558 | 196,3591727 |
| QRICH1 | ENSG00000198218 | 176,1578815 |
| JADE1 | ENSG00000077684 | 126,4674924 |
| EXOSC4 | ENSG00000178896 | 178,3092401 |
| ZNF350 | ENSG00000256683 | 130,3247642 |
| HMGXB4 | ENSG00000100281 | 198,03172 |
| RNF138 | ENSG00000134758 | 206,2987393 |
| GBP3 | ENSG00000117226 | 138,4750101 |
| SLMAP | ENSG00000163681 | 190,3278408 |
| MED29 | ENSG00000063322 | 158,8695994 |
| THAP12 | ENSG00000137492 | 138,1842623 |
| COX6B1 | ENSG00000126267 | 149,5259925 |
| WDFY1 | ENSG00000085449 | 166,8257214 |
| PPIP5K2 | ENSG00000145725 | 225,8779322 |
| KDELR2 | ENSG00000136240 | 152,4203497 |
| TAF1 | ENSG00000147133 | 151,1331087 |

| | | |
|---|---|---|
| ZBTB4 ENSG00000174282 | ENSG00000174282 | 134,0123031 |
| EPG5 | ENSG00000152223 | 158,2524583 |
| ETNK1 | ENSG00000139163 | 182,624937 |
| GINM1 | ENSG00000055211 | 200,6360713 |
| LACC1 | ENSG00000179630 | 124,8289401 |
| C2CD5 | ENSG00000111731 | 220,1847688 |
| APAF1 | ENSG00000120868 | 129,4773247 |
| MARCHF8 ENSG00000165406 | ENSG00000165406 | 178,0302491 |
| DNAJB9 | ENSG00000128590 | 195,3939076 |
| RFX5 | ENSG00000143390 | 173,500356 |
| FAM168A | ENSG00000054965 | 194,6522514 |
| SDHC | ENSG00000143252 | 177,3359382 |
| TOMM7 | ENSG00000196683 | 112,4620709 |
| HEBP2 | ENSG00000051620 | 192,0925782 |
| SNHG29 | ENSG00000175061 | 171,7128607 |
| AURKAIP1 | ENSG00000175756 | 115,8359398 |
| FAM32A | ENSG00000105058 | 132,7476574 |
| POM121C | ENSG00000272391 | 144,5309674 |
| CDK7 ENSG00000134058 | ENSG00000134058 | 148,6954519 |
| PDK1 | ENSG00000152256 | 131,2637424 |
| ZNF44 | ENSG00000197857 | 175,4027303 |
| SP2 | ENSG00000167182 | 134,4441855 |
| TRMT112 | ENSG00000173113 | 129,8540209 |
| KIAA2013 | ENSG00000116685 | 115,0505082 |
| ARHGEF40 | ENSG00000165801 | 175,8372793 |
| TSPO | ENSG00000100300 | 116,2616267 |
| PDPK1 | ENSG00000140992 | 137,8115347 |
| FCHSD2 | ENSG00000137478 | 199,2120661 |
| SINHCAF ENSG00000139146 | ENSG00000139146 | 165,121827 |
| GAS5 | ENSG00000234741 | 134,430755 |
| PSTPIP2 | ENSG00000152229 | 149,9438393 |
| BCL9L | ENSG00000186174 | 122,8038607 |
| EIF4E3 | ENSG00000163412 | 216,343359 |
| RETREG3 | ENSG00000141699 | 128,3164609 |
| TMED7 | ENSG00000134970 | 190,2465624 |
| ASXL2 | ENSG00000143970 | 184,0410385 |
| SNX2 | ENSG00000205302 | 184,0733295 |
| PDLIM7-AS1 | ENSG00000248996 | 161,7788725 |
| FOXO4 | ENSG00000184481 | 123,0864619 |
| TBCC | ENSG00000124659 | 112,2267636 |
| PSMC4 ENSG00000013275 | ENSG00000013275 | 131,0898487 |
| PPP2R1A | ENSG00000105568 | 104,0566605 |
| PRF1 | ENSG00000180644 | 93,27691409 |
| SCARB2 | ENSG00000138760 | 162,6007472 |
| SDHD | ENSG00000204370 | 143,6170638 |
| MVD | ENSG00000167508 | 154,7481109 |
| ZNF266 | ENSG00000174652 | 182,5690165 |
| MMP25-AS1 | ENSG00000261971 | 157,3289475 |
| BMP6 | ENSG00000153162 | 178,4956193 |
| SPAG7 | ENSG00000091640 | 152,8468605 |
| TBCD ENSG00000141556 | ENSG00000141556 | 130,8922395 |
| LGALS1 | ENSG00000100097 | 127,9759732 |
| CCM2 | ENSG00000136280 | 110,0129475 |
| TMA7 | ENSG00000232112 | 173,3879014 |
| SNX9 | ENSG00000130340 | 172,8431418 |

| | | |
|---|---|---|
| WASHC2C | ENSG00000172661 | 183,6226001 |
| AAK1 | ENSG00000115977 | 168,0887843 |
| SERPINB2 | ENSG00000197632 | 126,0077857 |
| CCL3 ENSG00000277632 | ENSG00000277632 | 110,6657894 |
| MED4 | ENSG00000136146 | 163,7439969 |
| TTC7A | ENSG00000068724 | 99,52946305 |
| GPATCH8 | ENSG00000186566 | 137,5031456 |
| PDAP1 | ENSG00000106244 | 152,7826422 |
| KLHL28 | ENSG00000179454 | 142,2828316 |
| FAM157C | ENSG00000260528 | 169,7689825 |
| SRSF9 | ENSG00000111786 | 166,71496 |
| HDAC1 | ENSG00000116478 | 162,7843801 |
| SREK1 | ENSG00000153914 | 187,7812928 |
| HDAC4 | ENSG00000068024 | 157,893765 |
| ASB8 | ENSG00000177981 | 143,9026421 |
| CCPG1 | ENSG00000260916 | 213,3452181 |
| PHF2 | ENSG00000197724 | 133,4526618 |
| MIA2 | ENSG00000150527 | 186,3937874 |
| ASB6 | ENSG00000148331 | 137,5525515 |
| MGAT4B ENSG00000161013 | ENSG00000161013 | 130,4391107 |
| CCDC97 | ENSG00000142039 | 123,0458079 |
| TUBB4B | ENSG00000188229 | 121,1779868 |
| SDHB | ENSG00000117118 | 184,3069624 |
| RPN2 | ENSG00000118705 | 134,9695491 |
| SEC63 | ENSG00000025796 | 188,4019375 |
| PLD2 | ENSG00000129219 | 159,8606956 |
| C1orf162 | ENSG00000143110 | 218,5758841 |
| H2AZ1 | ENSG00000164032 | 169,323482 |
| TMC8 | ENSG00000167895 | 126,5244438 |
| CSAD | ENSG00000139631 | 189,763469 |
| POU2F2 | ENSG00000028277 | 112,2349411 |
| ENSG00000279838 | ENSG00000279838 | 199,7676291 |
| H1-10 | ENSG00000184897 | 77,78719045 |
| TATDN2 | ENSG00000157014 | 136,0263556 |
| LEPROT | ENSG00000213625 | 191,5359724 |
| FCF1 | ENSG00000119616 | 143,5600979 |
| HILPDA | ENSG00000135245 | 86,0605494 |
| GSDMD ENSG00000104518 | ENSG00000104518 | 171,1147799 |
| TMEM131L | ENSG00000121210 | 176,0667672 |
| CLC | ENSG00000105205 | 168,9945469 |
| ST6GAL1 | ENSG00000073849 | 123,9720547 |
| ALG1L13P ENSG00000253981 | ENSG00000253981 | 141,1362429 |
| FAM131A | ENSG00000175182 | 159,6909434 |
| SLPI | ENSG00000124107 | 91,04045052 |
| SNIP1 | ENSG00000163877 | 153,2272736 |
| PDZD8 | ENSG00000165650 | 149,7577591 |
| CTBP2 | ENSG00000175029 | 150,0624613 |
| TPRA1 | ENSG00000163870 | 86,6061987 |
| MGAT5 | ENSG00000152127 | 134,3895302 |
| XPNPEP3 | ENSG00000196236 | 53,21022697 |
| NFX1 | ENSG00000086102 | 173,3665126 |
| WDR55 | ENSG00000120314 | 177,5038554 |
| SFSWAP | ENSG00000061936 | 124,2661568 |
| LMBR1L | ENSG00000139636 | 193,8098536 |
| CTSA | ENSG00000064601 | 175,1266171 |

| | | |
|---|---|---|
| FBXO7 | ENSG00000100225 | 166,9760372 |
| LYRM1 | ENSG00000102897 | 197,0956352 |
| NCAPH2 | ENSG00000025770 | 127,5686551 |
| AP2A1 | ENSG00000196961 | 139,7104872 |
| DCAF7 | ENSG00000136485 | 115,263833 |
| EIF3K ENSG00000178982 | ENSG00000178982 | 137,8900115 |
| RPS2P5 | ENSG00000240342 | 112,9421399 |
| TNKS ENSG00000173273 | ENSG00000173273 | 195,8873457 |
| DNMBP | ENSG00000107554 | 253,7564783 |
| EMICERI | ENSG00000285103 | 229,2919446 |
| DIS3 | ENSG00000083520 | 127,3399476 |
| ENG | ENSG00000106991 | 150,3493306 |
| LGALS8 | ENSG00000116977 | 170,5329463 |
| NF1 | ENSG00000196712 | 167,5414863 |
| EXOC1 | ENSG00000090989 | 158,0489087 |
| PDS5B | ENSG00000083642 | 204,0167494 |
| ATF6B ENSG00000213676 | ENSG00000213676 | 128,0390289 |
| GADD45G | ENSG00000130222 | 73,1202753 |
| CYC1 | ENSG00000179091 | 145,9563707 |
| BBIP1 | ENSG00000214413 | 175,8943127 |
| SNHG5 | ENSG00000203875 | 126,1042442 |
| ARID1B | ENSG00000049618 | 141,4767998 |
| HIF1AN | ENSG00000166135 | 140,7512886 |
| TMEM183A | ENSG00000163444 | 147,6223027 |
| ALKBH5 | ENSG00000091542 | 104,6350333 |
| HMOX2 ENSG00000103415 | ENSG00000103415 | 127,3677844 |
| VPS41 | ENSG00000006715 | 165,3274476 |
| SRP72 | ENSG00000174780 | 146,2172044 |
| DIP2A | ENSG00000160305 | 133,9093708 |
| ME2 | ENSG00000082212 | 169,0480105 |
| C11orf68 | ENSG00000175573 | 104,2273628 |
| ATG2B | ENSG00000066739 | 152,3877937 |
| ELK3 | ENSG00000111145 | 150,1221516 |
| GOLGA5 | ENSG00000066455 | 144,5673389 |
| ORAI1 | ENSG00000276045 | 188,8956962 |
| CNPPD1 | ENSG00000115649 | 120,5277575 |
| PNPLA2 | ENSG00000177666 | 156,4224613 |
| UPF2 | ENSG00000151461 | 176,3346793 |
| REPIN1 | ENSG00000214022 | 82,80305003 |
| GPRIN3 | ENSG00000185477 | 170,9645722 |
| MFN1 | ENSG00000171109 | 198,2996185 |
| SMC3 | ENSG00000108055 | 133,7804737 |
| VDAC2 | ENSG00000165637 | 147,1294969 |
| ACSL3 | ENSG00000123983 | 253,7940526 |
| EIF4E | ENSG00000151247 | 203,981367 |
| HSDL2 | ENSG00000119471 | 207,4076112 |
| SH3D21 | ENSG00000214193 | 187,6395543 |
| HYOU1 ENSG00000149428 | ENSG00000149428 | 100,6807193 |
| BMT2 | ENSG00000164603 | 119,8182884 |
| SNRPB | ENSG00000125835 | 123,2371818 |
| TTL | ENSG00000114999 | 98,66337267 |
| USP1 | ENSG00000162607 | 118,7772225 |
| ERAP1 | ENSG00000164307 | 165,2442493 |
| TCEA1 | ENSG00000187735 | 145,9304908 |
| TESK2 | ENSG00000070759 | 100,0096439 |

| | | |
|---|---|---|
| ZC3H3 ENSG00000014164 | ENSG00000014164 | 185,6604632 |
| ANKRD10 | ENSG00000088448 | 188,2637985 |
| BAG5 | ENSG00000166170 | 126,4360226 |
| ZNF441 | ENSG00000197044 | 69,24588286 |
| MAP3K7 | ENSG00000135341 | 152,0436498 |
| DHX38 | ENSG00000140829 | 141,1192748 |
| DDB1 | ENSG00000167986 | 111,3085991 |
| SEC24C | ENSG00000176986 | 117,4071644 |
| ERLIN2 | ENSG00000147475 | 128,9293681 |
| CD300LF | ENSG00000186074 | 187,7217081 |
| CORO7 ENSG00000262246 | ENSG00000262246 | 145,1733005 |
| TCF7L2 | ENSG00000148737 | 148,5012296 |
| STIM1 | ENSG00000167323 | 128,5149948 |
| RNF185 | ENSG00000138942 | 157,7585839 |
| SPG7 | ENSG00000197912 | 146,8868245 |
| ACBD3 | ENSG00000182827 | 166,5414983 |
| MAN1A2 | ENSG00000198162 | 151,8589138 |
| CAMKK1 | ENSG00000004660 | 191,6599996 |
| ING3 | ENSG00000071243 | 123,2836048 |
| TMEM185B | ENSG00000226479 | 130,0366826 |
| CDS2 | ENSG00000101290 | 171,2391726 |
| BCL2L11 | ENSG00000153094 | 125,0813539 |
| STMP1 | ENSG00000243317 | 159,3438166 |
| SWT1 | ENSG00000116668 | 182,9684993 |
| PSPC1 | ENSG00000121390 | 126,9223522 |
| DYNLL2 | ENSG00000264364 | 131,5930027 |
| SLC35E2B | ENSG00000189339 | 96,73853565 |
| STRN | ENSG00000115808 | 163,3315356 |
| IWS1 | ENSG00000163166 | 139,7955137 |
| INPPL1 | ENSG00000165458 | 194,6952229 |
| LINC02035 | ENSG00000273033 | 156,8929658 |
| PPIL4 | ENSG00000131013 | 196,0262273 |
| TMEM230 | ENSG00000089063 | 145,6434914 |
| CCDC174 | ENSG00000154781 | 131,8487364 |
| HIPK2 | ENSG00000064393 | 129,0319298 |
| UBXN6 | ENSG00000167671 | 104,1297037 |
| AXIN1 | ENSG00000103126 | 122,0430791 |
| CPNE1 | ENSG00000214078 | 160,6477131 |
| CAND1 | ENSG00000111530 | 136,8196756 |
| GIMAP7 | ENSG00000179144 | 113,7173974 |
| DAPK2 | ENSG00000035664 | 142,3167278 |
| PRKACB | ENSG00000142875 | 89,16234215 |
| ATP8A1 | ENSG00000124406 | 163,3546058 |
| EIF3J | ENSG00000104131 | 173,2425579 |
| SACM1L | ENSG00000211456 | 150,6269456 |
| HTT | ENSG00000197386 | 156,2900605 |
| BICD2 | ENSG00000185963 | 94,85809946 |
| SMG9 | ENSG00000105771 | 131,3353689 |
| APC | ENSG00000134982 | 146,5216469 |
| CNOT7 | ENSG00000198791 | 137,2676528 |
| CEP57 | ENSG00000166037 | 142,4457709 |
| POM121 | ENSG00000196313 | 120,9506907 |
| AKAP10 | ENSG00000108599 | 131,4367421 |
| MFSD11 | ENSG00000092931 | 153,9407809 |
| ATP7A | ENSG00000165240 | 158,1109909 |

| | | |
|---|---|---|
| PPP1R3D | ENSG00000132825 | 96,97391813 |
| RAB18 | ENSG00000099246 | 173,6452026 |
| ARRB1 | ENSG00000137486 | 99,85376424 |
| LDLR | ENSG00000130164 | 121,1192869 |
| RAB29 | ENSG00000117280 | 126,774512 |
| PRELID3B | ENSG00000101166 | 162,3403556 |
| ITGAV | ENSG00000138448 | 150,9674068 |
| CYSLTR1 | ENSG00000173198 | 138,5732852 |
| SUDS3 | ENSG00000111707 | 178,3279569 |
| EMC7 | ENSG00000134153 | 138,3183551 |
| HLA-L ENSG00000243753 | ENSG00000243753 | 88,93518792 |
| CYB5R3 | ENSG00000100243 | 119,0722733 |
| KLHL12 | ENSG00000117153 | 118,4645447 |
| POU2F1 | ENSG00000143190 | 172,9658336 |
| LTB4R ENSG00000213903 | ENSG00000213903 | 156,9416929 |
| DAGLB | ENSG00000164535 | 124,2654215 |
| API5 | ENSG00000166181 | 122,2837277 |
| ASCC2 | ENSG00000100325 | 145,9303067 |
| CAPN7 | ENSG00000131375 | 169,2904725 |
| ELOB | ENSG00000103363 | 132,8111593 |
| FBXW11 | ENSG00000072803 | 157,940108 |
| MEF2A | ENSG00000068305 | 181,02552 |
| USP33 | ENSG00000077254 | 153,5828965 |
| ZNF160 | ENSG00000170949 | 103,1130657 |
| RAN | ENSG00000132341 | 128,6344888 |
| APOBEC3B | ENSG00000179750 | 134,6935951 |
| MT2A | ENSG00000125148 | 91,93293331 |
| LINC01506 | ENSG00000234506 | 162,6303436 |
| NEMF | ENSG00000165525 | 151,635529 |
| CSF1R | ENSG00000182578 | 110,4738844 |
| LEMD2 | ENSG00000161904 | 127,7312153 |
| CUTALP | ENSG00000226752 | 86,64161068 |
| COX7B | ENSG00000131174 | 152,6136663 |
| SART3 | ENSG00000075856 | 120,6550907 |
| ASTL | ENSG00000188886 | 184,7179784 |
| CCNK | ENSG00000090061 | 163,0018575 |
| SLC39A6 | ENSG00000141424 | 177,0360259 |
| ZNF185 | ENSG00000147394 | 145,0565363 |
| GTF2H1 | ENSG00000110768 | 163,7793581 |
| SNHG16 | ENSG00000163597 | 106,9724411 |
| PRKAB1 | ENSG00000111725 | 199,5531836 |
| ECPAS | ENSG00000136813 | 154,0307392 |
| CFD ENSG00000197766 | ENSG00000197766 | 105,4691572 |
| COPZ1 | ENSG00000111481 | 140,8840359 |
| NTNG2 | ENSG00000196358 | 164,2374301 |
| PCGF3 | ENSG00000185619 | 194,3491714 |
| MAPKAPK3 | ENSG00000114738 | 126,9548068 |
| HSDL1 | ENSG00000103160 | 174,7628408 |
| RBM18 | ENSG00000119446 | 139,05739 |
| GLUD1 | ENSG00000148672 | 141,5965427 |
| LINC02207 | ENSG00000258476 | 134,8519926 |
| STARD10 | ENSG00000214530 | 103,338109 |
| OTUD4 | ENSG00000164164 | 135,2036859 |
| PPP1R21 | ENSG00000162869 | 133,1674776 |
| SIGLEC5 ENSG00000105501 | ENSG00000105501 | 158,7645656 |

| | | |
|---|---|---|
| NIBAN2 | ENSG00000136830 | 110,9670457 |
| YIPF6 | ENSG00000181704 | 157,1701484 |
| RAPGEF6 | ENSG00000158987 | 173,8460495 |
| SCAF1 | ENSG00000126461 | 94,57870265 |
| PLIN2 | ENSG00000147872 | 164,4789384 |
| NUP62 | ENSG00000213024 | 106,9217234 |
| GZMB | ENSG00000100453 | 78,69172374 |
| MICB ENSG00000204516 | ENSG00000204516 | 123,2479008 |
| ARHGEF7 | ENSG00000102606 | 132,1937033 |
| MED1 | ENSG00000125686 | 103,4315174 |
| ERGIC2 | ENSG00000087502 | 164,1304328 |
| CCAR2 | ENSG00000158941 | 135,5375318 |
| SPTLC1 | ENSG00000090054 | 167,9929398 |
| IL1RL1 | ENSG00000115602 | 123,279764 |
| RGP1 | ENSG00000107185 | 150,3185138 |
| WASHC5 | ENSG00000164961 | 125,8137998 |
| LINC00641 | ENSG00000258441 | 195,5077938 |
| DOK1 | ENSG00000115325 | 83,11533672 |
| ZNF516 | ENSG00000101493 | 150,7618442 |
| NAA60 | ENSG00000122390 | 126,0585838 |
| REEP5 | ENSG00000129625 | 159,7964648 |
| CLSTN3 | ENSG00000139182 | 105,6475919 |
| NDUFS1 ENSG00000023228 | ENSG00000023228 | 99,31800633 |
| MAPK3 | ENSG00000102882 | 156,5422654 |
| ANKH | ENSG00000154122 | 99,14174196 |
| ST3GAL4 | ENSG00000110080 | 170,0860103 |
| MAT2A | ENSG00000168906 | 161,6082151 |
| PABPC4 | ENSG00000090621 | 133,8107368 |
| COMMD5 | ENSG00000170619 | 116,2735183 |
| SMG5 | ENSG00000198952 | 124,6772969 |
| FLNB | ENSG00000136068 | 135,5612131 |
| TNFRSF8 | ENSG00000120949 | 96,63213562 |
| DERL1 | ENSG00000136986 | 136,3416137 |
| C1D | ENSG00000197223 | 152,5092659 |
| RBM43 | ENSG00000184898 | 108,2176804 |
| NDUFA1 | ENSG00000125356 | 123,932522 |
| VPS28 ENSG00000160948 | ENSG00000160948 | 109,5164018 |
| EGLN1 | ENSG00000135766 | 136,6816069 |
| SEC11A | ENSG00000140612 | 138,8675914 |
| WDR91 | ENSG00000105875 | 150,1168706 |
| VIRMA | ENSG00000164944 | 123,3558671 |
| ACLY | ENSG00000131473 | 134,6388438 |
| LMF2 | ENSG00000100258 | 116,7059493 |
| GNL3L | ENSG00000130119 | 109,474736 |
| ZCWPW1 | ENSG00000078487 | 163,692893 |
| CREB3 | ENSG00000107175 | 143,5639671 |
| CD7 | ENSG00000173762 | 78,5891103 |
| CLTB | ENSG00000175416 | 127,0355037 |
| SLC25A5 | ENSG00000005022 | 123,3110124 |
| UBE2N | ENSG00000177889 | 142,4821834 |
| LIF | ENSG00000128342 | 192,1694225 |
| HSPBAP1 | ENSG00000169087 | 165,9557329 |
| EXOC5 | ENSG00000070367 | 123,341345 |
| ADAT1 | ENSG00000065457 | 106,41195 |
| NSRP1 | ENSG00000126653 | 162,0124789 |

| | | |
|---|---|---|
| UBE2J2 | ENSG00000160087 | 112,0166476 |
| SLC39A8 | ENSG00000138821 | 138,6942902 |
| GRK3 | ENSG00000100077 | 99,00911736 |
| STOM | ENSG00000148175 | 140,8649858 |
| VPS53 ENSG00000141252 | ENSG00000141252 | 117,8998961 |
| TMEM9B | ENSG00000175348 | 142,9526472 |
| VPS26C | ENSG00000157538 | 114,8455498 |
| PRKD3 | ENSG00000115825 | 138,01534 |
| CCT5 | ENSG00000150753 | 116,4316077 |
| WIPI1 | ENSG00000070540 | 132,3747856 |
| MED25 | ENSG00000104973 | 124,1626876 |
| CLASRP | ENSG00000104859 | 164,2073813 |
| SERPINB6 | ENSG00000124570 | 183,9202328 |
| LNCATV | ENSG00000238005 | 128,3912309 |
| ACTR10 | ENSG00000131966 | 143,5874749 |
| CIMAP1B | ENSG00000177989 | 106,2368457 |
| EGLN2 | ENSG00000269858 | 159,9633182 |
| INIP | ENSG00000148153 | 102,8635426 |
| RSL1D1 | ENSG00000171490 | 118,4273432 |
| ACO1 | ENSG00000122729 | 105,0047258 |
| ARFGEF2 | ENSG00000124198 | 130,4686302 |
| DPM1 | ENSG00000000419 | 140,687644 |
| ATF2 | ENSG00000115966 | 159,0717101 |
| YJU2B | ENSG00000104957 | 157,556984 |
| NOTCH2NLC | ENSG00000286219 | 99,47059156 |
| TTLL3 | ENSG00000214021 | 165,6551957 |
| SC5D | ENSG00000109929 | 174,5845492 |
| UBA7 | ENSG00000182179 | 150,1944467 |
| FNTA | ENSG00000168522 | 136,4282485 |
| NFRKB | ENSG00000170322 | 120,9698556 |
| PSMC6 | ENSG00000100519 | 141,871814 |
| ANKZF1 | ENSG00000163516 | 145,4687558 |
| SNAP23 | ENSG00000092531 | 152,1415827 |
| NFE2L1 | ENSG00000082641 | 90,11402697 |
| CENPC | ENSG00000145241 | 124,9299679 |
| AP1B1 | ENSG00000100280 | 92,92028651 |
| SNX19 | ENSG00000120451 | 125,5325074 |
| DENND6A | ENSG00000174839 | 185,1512556 |
| PPP1R12B | ENSG00000077157 | 124,3993112 |
| ARL11 | ENSG00000152213 | 125,6143019 |
| ZFAS1 | ENSG00000177410 | 153,4953164 |
| ZNF674 | ENSG00000251192 | 102,9316097 |
| STAG1 | ENSG00000118007 | 165,6263013 |
| ARFGAP2 | ENSG00000149182 | 97,37701953 |
| LAMTOR4 | ENSG00000188186 | 99,51953161 |
| HBB | ENSG00000244734 | 57,13074372 |
| ITPKC | ENSG00000086544 | 117,8935654 |
| LMBR1 | ENSG00000105983 | 156,2490277 |
| ENSG00000280614 | ENSG00000280614 | 21,2192273 |
| THUMPD1 | ENSG00000066654 | 142,9063521 |
| STK35 | ENSG00000125834 | 95,01142147 |
| H2BC20P | ENSG00000261716 | 108,2946398 |
| BDP1 ENSG00000145734 | ENSG00000145734 | 132,8005849 |
| PRPF4 | ENSG00000136875 | 139,5587861 |
| OPA1 | ENSG00000198836 | 147,1688835 |

| | | |
|---|---|---|
| BTNL8 | ENSG00000113303 | 143,1624598 |
| RBBP7 | ENSG00000102054 | 122,7624023 |
| RNF146 | ENSG00000118518 | 154,6491875 |
| MTMR4 | ENSG00000108389 | 121,2131448 |
| ABTB3 | ENSG00000151136 | 107,3517876 |
| LRRC8C | ENSG00000171488 | 130,1709384 |
| BFAR ENSG00000103429 | ENSG00000103429 | 147,9643333 |
| FCGR2C | ENSG00000244682 | 223,7105293 |
| TRIOBP | ENSG00000100106 | 100,3324906 |
| VRK3 | ENSG00000105053 | 111,7757312 |
| CHD4 | ENSG00000111642 | 122,3335149 |
| FKBP5 | ENSG00000096060 | 155,7296627 |
| CHMP7 | ENSG00000147457 | 97,20367537 |
| KHDC4 | ENSG00000132680 | 163,8207176 |
| ALCAM | ENSG00000170017 | 165,0037684 |
| LPAR6 | ENSG00000139679 | 92,91527423 |
| PUS10 | ENSG00000162927 | 139,1533369 |
| KARS1 | ENSG00000065427 | 109,9141256 |
| EXOSC9 | ENSG00000123737 | 122,1633468 |
| ASMTL | ENSG00000169093 | 121,2083863 |
| DDX23 | ENSG00000174243 | 121,7274773 |
| GTF2F1 | ENSG00000125651 | 114,773701 |
| DGCR2 | ENSG00000070413 | 118,0485774 |
| HMG20B | ENSG00000064961 | 126,6671967 |
| ZNF731P | ENSG00000227671 | 182,8342899 |
| IRF4 | ENSG00000137265 | 112,764885 |
| HK1 | ENSG00000156515 | 174,3780475 |
| GOSR1 | ENSG00000108587 | 139,4919765 |
| CTBP1 | ENSG00000159692 | 120,1726191 |
| PLEKHJ1 | ENSG00000104886 | 94,77299725 |
| CPNE3 | ENSG00000085719 | 159,5177494 |
| STX10 | ENSG00000104915 | 131,6319985 |
| ATG9A | ENSG00000198925 | 130,5951403 |
| UBE2Q1 | ENSG00000160714 | 118,7867294 |
| SLC38A10 | ENSG00000157637 | 107,5475693 |
| RNF216 | ENSG00000011275 | 141,7600593 |
| ACAP1 | ENSG00000072818 | 153,4430885 |
| SLC25A13 | ENSG00000004864 | 139,6822168 |
| ZNF644 | ENSG00000122482 | 151,6721291 |
| TARDBP | ENSG00000120948 | 123,0474941 |
| PBRM1 | ENSG00000163939 | 117,0862165 |
| ABCC5 | ENSG00000114770 | 183,4739565 |
| ANXA6 | ENSG00000197043 | 112,4986257 |
| SLC35C2 | ENSG00000080189 | 121,1218823 |
| FAM76B | ENSG00000077458 | 173,1979804 |
| MYNN | ENSG00000085274 | 142,1791448 |
| ZNF331 | ENSG00000130844 | 129,0716867 |
| GALNT7 | ENSG00000109586 | 136,3023128 |
| LHFPL5 | ENSG00000197753 | 164,1614819 |
| OXA1L | ENSG00000155463 | 115,9549019 |
| NARS1 | ENSG00000134440 | 145,436081 |
| GALNS | ENSG00000141012 | 129,0714552 |
| CTNNBL1 | ENSG00000132792 | 136,332693 |
| CREBL2 | ENSG00000111269 | 120,5893874 |
| ADGRG1 | ENSG00000205336 | 82,04896652 |

| | | |
|---|---|---|
| DPH3 | ENSG00000154813 | 144,7507864 |
| ZBTB7A | ENSG00000178951 | 121,8254529 |
| TIAL1 | ENSG00000151923 | 147,4345119 |
| ANKRD49 | ENSG00000168876 | 117,9305175 |
| AP3S2 | ENSG00000157823 | 119,6673269 |
| ZNF493 | ENSG00000196268 | 150,5138873 |
| RCOR3 | ENSG00000117625 | 138,0692377 |
| ELOVL1 | ENSG00000066322 | 91,93347617 |
| NKIRAS2 | ENSG00000168256 | 122,0591446 |
| MCM5 | ENSG00000100297 | 137,7427696 |
| EDF1 | ENSG00000107223 | 102,891291 |
| RBM27 | ENSG00000091009 | 132,5866224 |
| SGTA | ENSG00000104969 | 101,5085242 |
| THUMPD3-AS1 | ENSG00000206573 | 142,6376404 |
| JADE2 | ENSG00000043143 | 84,53604646 |
| SCPEP1 | ENSG00000121064 | 152,3997232 |
| GMEB1 | ENSG00000162419 | 129,6022244 |
| TBC1D20 | ENSG00000125875 | 122,2788547 |
| KMT2A | ENSG00000118058 | 96,88164997 |
| ESYT1 | ENSG00000139641 | 90,17124779 |
| PDXDC1 ENSG00000179889 | ENSG00000179889 | 112,2373468 |
| ZMIZ2 | ENSG00000122515 | 110,2320412 |
| GARRE1 ENSG00000166398 | ENSG00000166398 | 103,7865189 |
| IRF8 | ENSG00000140968 | 116,2185497 |
| DGAT1 ENSG00000185000 | ENSG00000185000 | 126,6637797 |
| PNKD | ENSG00000127838 | 94,51536497 |
| PANK4 ENSG00000157881 | ENSG00000157881 | 97,65893477 |
| HIP1R | ENSG00000130787 | 99,20274191 |
| USO1 | ENSG00000138768 | 148,0288729 |
| H6PD | ENSG00000049239 | 104,0136749 |
| AP1M1 | ENSG00000072958 | 86,41043023 |
| SENP7 | ENSG00000138468 | 185,1589982 |
| FBXO42 | ENSG00000037637 | 143,0666872 |
| ZNF131 | ENSG00000172262 | 142,5487269 |
| PLPP3 | ENSG00000162407 | 152,4693224 |
| DNAJB11 | ENSG00000090520 | 171,3606389 |
| CLCN7 | ENSG00000103249 | 137,0430064 |
| CTNND1 | ENSG00000198561 | 114,6819075 |
| ATAD2B | ENSG00000119778 | 141,3418227 |
| EIF2S2 | ENSG00000125977 | 125,9232453 |
| FAM217B | ENSG00000196227 | 75,13391692 |
| BACH2 | ENSG00000112182 | 106,2608672 |
| NBPF10 | ENSG00000271425 | 120,7382113 |
| SLC25A28 | ENSG00000155287 | 154,8485557 |
| DELE1 | ENSG00000081791 | 137,2768833 |
| HCG27 ENSG00000206344 | ENSG00000206344 | 130,6028237 |
| RBMXL1 | ENSG00000213516 | 144,7041228 |
| CXCL2 | ENSG00000081041 | 178,2991229 |
| TBC1D10B | ENSG00000169221 | 156,1759867 |
| RAB11B | ENSG00000185236 | 120,3842996 |
| NUP210 | ENSG00000132182 | 86,73476676 |
| CD79A | ENSG00000105369 | 90,016926 |
| HLA-DRB5 | ENSG00000198502 | 101,2826226 |
| PHF8 | ENSG00000172943 | 146,3885304 |
| OIP5-AS1 | ENSG00000247556 | 115,6950286 |

| | | |
|---|---|---|
| CXCL10 | ENSG00000169245 | 46,92763884 |
| LINC-PINT | ENSG00000231721 | 184,0688883 |
| ALDOA | ENSG00000149925 | 132,1456496 |
| DDX46 | ENSG00000145833 | 121,1152981 |
| TRMT1L | ENSG00000121486 | 122,8893859 |
| VILL | ENSG00000136059 | 77,60179969 |
| ZAP70 | ENSG00000115085 | 79,1804984 |
| ZFAND2B | ENSG00000158552 | 138,4589195 |
| ISCA1 | ENSG00000135070 | 175,4135003 |
| MFAP1 | ENSG00000140259 | 121,6232701 |
| CTDNEP1 | ENSG00000175826 | 113,3473568 |
| CASP10 | ENSG00000003400 | 114,7253522 |
| UVRAG | ENSG00000198382 | 131,1378419 |
| ANAPC5 | ENSG00000089053 | 105,1431753 |
| MANSC1 ENSG00000111261 | ENSG00000111261 | 158,4878253 |
| FRAT1 | ENSG00000165879 | 69,33474031 |
| UBR3 | ENSG00000144357 | 156,9133446 |
| SYPL1 | ENSG00000008282 | 126,6633012 |
| INTS1 | ENSG00000164880 | 89,65830877 |
| LYPD3 | ENSG00000124466 | 60,75338601 |
| RANBP9 | ENSG00000010017 | 132,5854249 |
| SERTAD3 | ENSG00000167565 | 162,9681933 |
| AMZ2 | ENSG00000196704 | 153,9017656 |
| CENPBD2P | ENSG00000213753 | 83,72051673 |
| FEM1B | ENSG00000169018 | 99,2604149 |
| STK11 | ENSG00000118046 | 92,62808828 |
| ZNF518B | ENSG00000178163 | 152,6326433 |
| CLEC4D | ENSG00000166527 | 170,0031969 |
| GALNT11 | ENSG00000178234 | 105,0124602 |
| UFM1 | ENSG00000120686 | 132,5045287 |
| FCHO2 | ENSG00000157107 | 178,6082895 |
| H1-0 | ENSG00000189060 | 111,5008285 |
| LRIF1 | ENSG00000121931 | 103,3835276 |
| ARMCX3 | ENSG00000102401 | 162,7738427 |
| TIA1 | ENSG00000116001 | 122,422856 |
| C21orf91 | ENSG00000154642 | 138,4610991 |
| DCAF8 | ENSG00000132716 | 133,3899364 |
| ABAT | ENSG00000183044 | 133,9194113 |
| TRIM44 | ENSG00000166326 | 114,326195 |
| FAM50A | ENSG00000071859 | 124,0703944 |
| DARS1 | ENSG00000115866 | 131,9631434 |
| C3orf62 | ENSG00000188315 | 127,273689 |
| DAXX ENSG00000204209 | ENSG00000204209 | 117,2732424 |
| TASOR2 | ENSG00000108021 | 111,3773786 |
| MAP1S | ENSG00000130479 | 93,1392824 |
| TNRC6A | ENSG00000090905 | 94,49816009 |
| IKBKG | ENSG00000269335 | 118,5961961 |
| SULT1A1 | ENSG00000196502 | 79,15722552 |
| TAF15 ENSG00000270647 | ENSG00000270647 | 121,324947 |
| YARS1 | ENSG00000134684 | 99,45028524 |
| CTSH | ENSG00000103811 | 102,8726788 |
| MPRIP | ENSG00000133030 | 83,78529784 |
| CWC15 | ENSG00000150316 | 98,63396481 |
| DDOST | ENSG00000244038 | 87,54060546 |
| AEBP2 | ENSG00000139154 | 136,0746747 |

| | | |
|---|---|---|
| KYAT3 | ENSG00000137944 | 152,8384945 |
| ZNF318 | ENSG00000171467 | 107,0205693 |
| ALYREF | ENSG00000183684 | 93,32477418 |
| UBP1 | ENSG00000153560 | 114,9525611 |
| FASN | ENSG00000169710 | 81,09574904 |
| ATP2C1 | ENSG00000017260 | 107,7227206 |
| MAD2L1BP | ENSG00000124688 | 118,9204412 |
| ST20 | ENSG00000180953 | 169,363264 |
| C19orf38 | ENSG00000214212 | 101,5074494 |
| HADHB | ENSG00000138029 | 140,1226658 |
| GMEB2 | ENSG00000101216 | 96,43426212 |
| BRAP | ENSG00000089234 | 103,7366612 |
| LRRC8B | ENSG00000197147 | 97,63151152 |
| GBE1 | ENSG00000114480 | 154,1056934 |
| TTN | ENSG00000155657 | 174,2317035 |
| PNN | ENSG00000100941 | 167,9737917 |
| TEP1 | ENSG00000129566 | 93,90584297 |
| GIGYF2 | ENSG00000204120 | 130,9281502 |
| RSPRY1 | ENSG00000159579 | 125,9462248 |
| RHOF | ENSG00000139725 | 109,3491485 |
| MRFAP1L1 | ENSG00000178988 | 94,30399767 |
| ZDHHC6 | ENSG00000023041 | 130,7332669 |
| ENY2 | ENSG00000120533 | 129,0043694 |
| CNST | ENSG00000162852 | 147,6707211 |
| TBC1D7 | ENSG00000145979 | 161,8150096 |
| TLNRD1 | ENSG00000140406 | 80,90645867 |
| SF3B6 | ENSG00000115128 | 121,6752925 |
| RNF34 | ENSG00000170633 | 112,4480933 |
| TBC1D5 | ENSG00000131374 | 128,871158 |
| NCOA6 | ENSG00000198646 | 131,4728648 |
| RBM10 | ENSG00000182872 | 103,7125934 |
| OTUB1 | ENSG00000167770 | 111,8158884 |
| PANK2 | ENSG00000125779 | 116,6854878 |
| LONP2 | ENSG00000102910 | 110,1929218 |
| COX7C | ENSG00000127184 | 98,78774066 |
| SLC36A1 | ENSG00000123643 | 107,6859441 |
| PLAA | ENSG00000137055 | 131,8201302 |
| KIFC3 | ENSG00000140859 | 106,481735 |
| SMC1A | ENSG00000072501 | 108,9567177 |
| PLEKHA3 | ENSG00000116095 | 121,793175 |
| BCL7B | ENSG00000106635 | 128,4632856 |
| DAD1 | ENSG00000129562 | 134,8189111 |
| CD3G | ENSG00000160654 | 68,39601882 |
| RSBN1 | ENSG00000081019 | 129,9798399 |
| SIAH1 | ENSG00000196470 | 120,9611975 |
| NPC1 | ENSG00000141458 | 120,5565236 |
| CTDSPL2 | ENSG00000137770 | 126,2915777 |
| PTPRN2 ENSG00000155093 | ENSG00000155093 | 99,88574645 |
| ITGB2-AS1 | ENSG00000227039 | 92,06684026 |
| CHD9 | ENSG00000177200 | 111,0102987 |
| PANK3 | ENSG00000120137 | 113,7135249 |
| SIAH2 | ENSG00000181788 | 93,01396894 |
| UBALD1 | ENSG00000153443 | 85,59383822 |
| SLC39A9 | ENSG00000029364 | 93,37731622 |
| NUDT21 | ENSG00000167005 | 97,26126907 |

| | | |
|---|---|---|
| ZNF672 | ENSG00000171161 | 110,9998456 |
| SLC30A9 | ENSG00000014824 | 125,0276535 |
| DYNC1LI2 | ENSG00000135720 | 132,8640075 |
| PITPNB | ENSG00000180957 | 121,3064541 |
| RIC1 | ENSG00000107036 | 125,838168 |
| NUS1 | ENSG00000153989 | 115,6924895 |
| GTF2IP12 | ENSG00000283050 | 137,8571902 |
| MOAP1 ENSG00000165943 | ENSG00000165943 | 100,4179219 |
| RBBP4 | ENSG00000162521 | 113,9674492 |
| SYVN1 | ENSG00000162298 | 129,5087458 |
| NEPRO | ENSG00000163608 | 128,9387257 |
| BST2 | ENSG00000130303 | 77,70961847 |
| DSC2 | ENSG00000134755 | 188,9760515 |
| STARD4 | ENSG00000164211 | 200,032994 |
| RTRAF | ENSG00000087302 | 140,9056515 |
| YWHAH | ENSG00000128245 | 96,86268904 |
| ZRANB2 | ENSG00000132485 | 115,7256857 |
| ACSS2 | ENSG00000131069 | 157,8831423 |
| EP400 | ENSG00000183495 | 99,20279643 |
| SF3B3 | ENSG00000189091 | 94,41266266 |
| SYNE3 | ENSG00000176438 | 119,2799493 |
| UBE4B | ENSG00000130939 | 135,2868063 |
| ILF2 | ENSG00000143621 | 104,4033842 |
| ATF3 | ENSG00000162772 | 97,61078629 |
| UBE2Q2 | ENSG00000140367 | 101,1764527 |
| TREML3P | ENSG00000184106 | 159,9788174 |
| DNMT3A | ENSG00000119772 | 120,726872 |
| IPO5 | ENSG00000065150 | 97,82103981 |
| GABPA | ENSG00000154727 | 122,0048126 |
| ZNF800 | ENSG00000048405 | 118,8114533 |
| RUFY1 ENSG00000176783 | ENSG00000176783 | 121,5207155 |
| DENND11 ENSG00000257093 | ENSG00000257093 | 109,5005391 |
| BAZ1B | ENSG00000009954 | 117,1170368 |
| BPNT2 | ENSG00000104331 | 101,5035852 |
| COX5B | ENSG00000135940 | 113,6457147 |
| SETD3 | ENSG00000183576 | 138,8418504 |
| MASTL | ENSG00000120539 | 138,1920771 |
| SLC35A4 | ENSG00000176087 | 113,9813733 |
| PRCC | ENSG00000143294 | 104,8495865 |
| GOLGA3 | ENSG00000090615 | 89,16841825 |
| GJB6 | ENSG00000121742 | 150,762903 |
| STARD3NL | ENSG00000010270 | 118,0946865 |
| PSMD11 | ENSG00000108671 | 103,8861398 |
| PABPN1 | ENSG00000100836 | 125,5074474 |
| PLB1 | ENSG00000163803 | 157,0786293 |
| GALNT10 | ENSG00000164574 | 67,40666656 |
| UBE2L3 | ENSG00000185651 | 97,01062659 |
| INO80 | ENSG00000128908 | 110,5666616 |
| CLPTM1 | ENSG00000104853 | 96,91620946 |
| HEATR5B | ENSG00000008869 | 94,47731733 |
| MPPE1 | ENSG00000154889 | 91,66419962 |
| B3GNT2 | ENSG00000170340 | 131,176024 |
| RNF125 | ENSG00000101695 | 75,21690348 |
| RCAN3 | ENSG00000117602 | 98,92596457 |
| SLC44A1 | ENSG00000070214 | 180,3361094 |

| | | |
|---|---|---|
| SMIM14 | ENSG00000163683 | 122,0260714 |
| THRA | ENSG00000126351 | 61,01460369 |
| RELL1 | ENSG00000181826 | 124,41677 |
| LRRFIP2 | ENSG00000093167 | 143,7994706 |
| TRIM23 | ENSG00000113595 | 97,58564742 |
| ATG16L1 ENSG0000085978 | ENSG00000085978 | 100,1062775 |
| APP | ENSG00000142192 | 122,9193215 |
| PDCD10 | ENSG00000114209 | 142,3361684 |
| SUMO3 | ENSG00000184900 | 109,6560886 |
| SRBD1 | ENSG00000068784 | 138,3074091 |
| ABT1 | ENSG00000146109 | 73,11355676 |
| MAP2K2 | ENSG00000126934 | 100,2202135 |
| FHIP1B | ENSG00000051009 | 136,2286777 |
| CLPX | ENSG00000166855 | 94,72124262 |
| MICU2 | ENSG00000165487 | 122,2601918 |
| HDAC2 | ENSG00000196591 | 110,97191 |
| TOLLIP | ENSG00000078902 | 79,88543433 |
| COX7A2L | ENSG00000115944 | 99,87777083 |
| LRRC59 | ENSG00000108829 | 88,57618093 |
| SH3BP5-AS1 | ENSG00000224660 | 164,0664076 |
| MAN2B2 | ENSG00000013288 | 102,9302566 |
| ZNF3 | ENSG00000166526 | 119,4723001 |
| RUNX2 | ENSG00000124813 | 109,6110803 |
| TBC1D2 | ENSG00000095383 | 96,66901274 |
| ARMT1 | ENSG00000146476 | 114,0267163 |
| TMEM273 | ENSG00000204161 | 101,4437293 |
| GARS1 | ENSG00000106105 | 139,6817761 |
| TRAPPC11 | ENSG00000168538 | 111,6369181 |
| TRAPPC1 | ENSG00000170043 | 101,4537021 |
| TUBGCP6 | ENSG00000128159 | 104,8829657 |
| SNAP29 | ENSG00000099940 | 102,0457574 |
| DNAH1 | ENSG00000114841 | 126,1032772 |
| PSMF1 | ENSG00000125818 | 85,48471223 |
| C11orf21 | ENSG00000110665 | 63,06924526 |
| BPGM | ENSG00000172331 | 97,78301635 |
| TMPO | ENSG00000120802 | 94,02798261 |
| CD109 | ENSG00000156535 | 107,5325789 |
| HIGD1A | ENSG00000181061 | 139,5537106 |
| KHSRP | ENSG00000088247 | 88,09747371 |
| MED18 ENSG00000130772 | ENSG00000130772 | 90,85826392 |
| RRBP1 | ENSG00000125844 | 137,2651131 |
| EIF3G | ENSG00000130811 | 84,07664433 |
| RANGAP1 | ENSG00000100401 | 76,20471339 |
| UBE2V2 | ENSG00000169139 | 121,5799447 |
| UBE3A | ENSG00000114062 | 99,18100578 |
| COPS2 | ENSG00000166200 | 121,578584 |
| CLTA | ENSG00000122705 | 88,11582884 |
| MGA | ENSG00000174197 | 99,29355024 |
| ELMO2 | ENSG00000062598 | 121,9088812 |
| IKBKE | ENSG00000263528 | 67,1314459 |
| SCN1B | ENSG00000105711 | 122,2978636 |
| CFLAR-AS1 | ENSG00000226312 | 106,856457 |
| UNKL | ENSG00000059145 | 97,24158828 |
| FFAR3 | ENSG00000185897 | 70,934867 |
| ZNF136 | ENSG00000196646 | 98,77182837 |

| | | |
|---|---|---|
| PSMD8 | ENSG00000099341 | 92,45134929 |
| TMCO1 | ENSG00000143183 | 89,18057522 |
| PLEKHG2 | ENSG00000090924 | 146,4024308 |
| SIGLEC9 | ENSG00000129450 | 84,28716922 |
| RPS6KB1 | ENSG00000108443 | 112,4513842 |
| PLXNB2 | ENSG00000196576 | 88,35210973 |
| LIMK1 | ENSG00000106683 | 98,976383 |
| LAMTOR5 | ENSG00000134248 | 166,6419044 |
| TRRAP | ENSG00000196367 | 106,2248361 |
| ERP29 | ENSG00000089248 | 109,6652788 |
| TGFA | ENSG00000163235 | 193,260824 |
| TPMT | ENSG00000137364 | 105,1573299 |
| TMED9 | ENSG00000184840 | 91,8252559 |
| BTN3A3 | ENSG00000111801 | 88,6051956 |
| C14orf119 | ENSG00000179933 | 99,88582049 |
| MLLT11 | ENSG00000213190 | 142,5477499 |
| PRMT2 | ENSG00000160310 | 98,35863118 |
| SMC5 | ENSG00000198887 | 116,8289335 |
| TTC17 | ENSG00000052841 | 131,2444972 |
| MTA2 | ENSG00000149480 | 106,7555218 |
| ENSG00000279861 | ENSG00000279861 | 123,471093 |
| EML3 | ENSG00000149499 | 98,58548686 |
| PHF10 | ENSG00000130024 | 127,8101372 |
| RAB9A | ENSG00000123595 | 103,4234527 |
| LYL1 | ENSG00000104903 | 75,13289143 |
| SGSM2 | ENSG00000141258 | 118,7472735 |
| KDM2B | ENSG00000089094 | 97,78464166 |
| CSTF1 | ENSG00000101138 | 84,86366953 |
| NGLY1 | ENSG00000151092 | 119,2594912 |
| SMC4 | ENSG00000113810 | 110,1061124 |
| RALA | ENSG00000006451 | 90,32157756 |
| ACP5 | ENSG00000102575 | 145,9601533 |
| ENSG00000260467 | ENSG00000260467 | 171,0049444 |
| TMOD2 | ENSG00000128872 | 92,70512462 |
| UQCRFS1 | ENSG00000169021 | 112,3445356 |
| LAMP3 | ENSG00000078081 | 105,8630858 |
| APPBP2 | ENSG00000062725 | 107,7662304 |
| CEACAM1 | ENSG00000079385 | 140,7828069 |
| PLCB3 | ENSG00000149782 | 137,9745572 |
| GTF3C1 | ENSG00000077235 | 83,71716012 |
| MUL1 | ENSG00000090432 | 68,6300539 |
| MT-ATP8 | ENSG00000228253 | 124,1473727 |
| RPRD1B | ENSG00000101413 | 96,47273638 |
| HARS2 | ENSG00000112855 | 120,052211 |
| OST4 | ENSG00000228474 | 103,8992008 |
| ZNF776 | ENSG00000152443 | 102,7445013 |
| CDK19 | ENSG00000155111 | 120,1165546 |
| IPCEF1 | ENSG00000074706 | 117,8279657 |
| FES | ENSG00000182511 | 124,1817599 |
| PIP5K1A | ENSG00000143398 | 134,0436559 |
| TRIM14 | ENSG00000106785 | 85,29086394 |
| MFSD2A | ENSG00000168389 | 93,00074857 |
| ATXN7L3B | ENSG00000253719 | 90,66175137 |
| WDR33 | ENSG00000136709 | 100,9204333 |
| CLASP2 | ENSG00000163539 | 102,5905425 |

| | | |
|---|---|---|
| DNAJC7 | ENSG00000168259 | 124,3245096 |
| FBXO30 | ENSG00000118496 | 104,5064442 |
| EML2 | ENSG00000125746 | 100,1284106 |
| DCAF12 | ENSG00000198876 | 99,69764504 |
| SESTD1 | ENSG00000187231 | 112,8632243 |
| LRCH4 | ENSG00000077454 | 116,3241471 |
| TAF2 | ENSG00000064313 | 118,5939445 |
| TENT4B | ENSG00000121274 | 117,9102108 |
| PTPRA | ENSG00000132670 | 106,0814416 |
| SEC23A | ENSG00000100934 | 120,8948468 |
| DGUOK | ENSG00000114956 | 106,779966 |
| DDX20 | ENSG00000064703 | 83,69714999 |
| C2orf68 | ENSG00000168887 | 75,4445272 |
| RBBP8 | ENSG00000101773 | 117,2998454 |
| DDI2 | ENSG00000197312 | 82,22275605 |
| PDCD4-AS1 | ENSG00000203497 | 100,4154801 |
| PIRAT1 | ENSG00000237803 | 94,95261572 |
| ZNF438 | ENSG00000183621 | 150,4979831 |
| ADA2 | ENSG00000093072 | 111,2406741 |
| RABGAP1 | ENSG00000011454 | 125,8092931 |
| DNAJB14 | ENSG00000164031 | 130,8446492 |
| MYBPC3 | ENSG00000134571 | 133,4119507 |
| SERINC5 | ENSG00000164300 | 106,1865686 |
| WDR47 | ENSG00000085433 | 132,9554264 |
| KANSL1L | ENSG00000144445 | 115,3928686 |
| MCM7 | ENSG00000166508 | 87,17128758 |
| SLC8B1 | ENSG00000089060 | 125,5047191 |
| HPS1 | ENSG00000107521 | 110,036386 |
| APOBEC3C | ENSG00000244509 | 91,3027446 |
| CBLB | ENSG00000114423 | 74,84580218 |
| SRSF1 | ENSG00000136450 | 130,8973942 |
| TAPT1 | ENSG00000169762 | 120,6707688 |
| EHMT1 | ENSG00000181090 | 96,90086264 |
| RGCC | ENSG00000102760 | 97,54190278 |
| MLXIP | ENSG00000175727 | 99,32666267 |
| ZNF101 | ENSG00000181896 | 84,15781138 |
| ARFGAP1 | ENSG00000101199 | 98,31399113 |
| MX1-AS1 | ENSG00000228318 | 71,4237331 |
| RBM6 | ENSG00000004534 | 148,5544924 |
| BCR | ENSG00000186716 | 92,68474148 |
| COQ8A | ENSG00000163050 | 64,40000775 |
| KCTD10 | ENSG00000110906 | 129,2464839 |
| ARNT | ENSG00000143437 | 120,3922412 |
| ZBTB6 | ENSG00000186130 | 77,80703293 |
| SPATA2L | ENSG00000158792 | 82,91696914 |
| STEAP4 | ENSG00000127954 | 119,0176979 |
| NUP155 | ENSG00000113569 | 132,1007392 |
| LINC01001 | ENSG00000230724 | 156,9385834 |
| SCAMP4 | ENSG00000227500 | 92,68337044 |
| CWF19L1 | ENSG00000095485 | 116,0562488 |
| DUSP10 | ENSG00000143507 | 59,74491702 |
| MEGF6 | ENSG00000162591 | 108,8868138 |
| PASK | ENSG00000115687 | 75,55348534 |
| AEN | ENSG00000181026 | 81,85478657 |
| TBRG1 | ENSG00000154144 | 117,168009 |

| | | |
|---|---|---|
| CCDC59 | ENSG00000133773 | 99,22990182 |
| DAZAP1 | ENSG00000071626 | 117,0565072 |
| ASPRV1 | ENSG00000244617 | 100,6284902 |
| TASOR | ENSG00000163946 | 110,8433083 |
| SENP1 | ENSG00000079387 | 96,95821365 |
| STRIP1 | ENSG00000143093 | 87,32782006 |
| BTK | ENSG00000010671 | 114,713797 |
| EIF2A | ENSG00000144895 | 122,0898327 |
| EPHB1 | ENSG00000154928 | 108,3804186 |
| ICE1 | ENSG00000164151 | 93,62938623 |
| TSNAX | ENSG00000116918 | 134,5756165 |
| CD2 | ENSG00000116824 | 83,39540189 |
| TMEM50B ENSG00000142188 | ENSG00000142188 | 112,3481637 |
| RPS6KA5 | ENSG00000100784 | 128,1080124 |
| EIF4E2 | ENSG00000135930 | 77,67994371 |
| ANXA3 | ENSG00000138772 | 180,8665495 |
| MAP2K7 | ENSG00000076984 | 95,56062519 |
| CYP27A1 | ENSG00000135929 | 85,33205193 |
| NOTCH2NLA | ENSG00000264343 | 76,65015525 |
| SLC22A18 ENSG00000110628 | ENSG00000110628 | 71,80324118 |
| FAM193A | ENSG00000125386 | 100,8913921 |
| PPM1B | ENSG00000138032 | 122,6556057 |
| CHERP | ENSG00000085872 | 79,34761263 |
| MAP3K20 | ENSG00000091436 | 97,91565882 |
| DCUN1D2 | ENSG00000150401 | 119,6638501 |
| ZNF701 | ENSG00000167562 | 61,71898249 |
| MXI1 | ENSG00000119950 | 111,5800569 |
| SFI1 | ENSG00000198089 | 96,86598749 |
| MAML3 | ENSG00000196782 | 109,34659 |
| KPNA3 | ENSG00000102753 | 111,7139722 |
| SP4 | ENSG00000105866 | 105,9161919 |
| RAB5IF | ENSG00000101084 | 104,15899 |
| RXRB ENSG00000204231 | ENSG00000204231 | 103,9679987 |
| SZT2 | ENSG00000198198 | 106,4962856 |
| DEDD | ENSG00000158796 | 126,5756793 |
| PUF60 | ENSG00000179950 | 83,39977818 |
| CNOT10 | ENSG00000182973 | 118,5057118 |
| DAP3 | ENSG00000132676 | 91,58450134 |
| ARID5B | ENSG00000150347 | 87,00664582 |
| BTBD1 | ENSG00000064726 | 101,7617213 |
| CHRAC1 | ENSG00000104472 | 103,9614994 |
| SPCS2 | ENSG00000118363 | 94,9504726 |
| RPL7L1 | ENSG00000146223 | 77,29479054 |
| KSR1 | ENSG00000141068 | 127,132468 |
| EIF3B | ENSG00000106263 | 82,52430724 |
| ERGIC3 | ENSG00000125991 | 85,59373026 |
| EID1 | ENSG00000255302 | 111,1779571 |
| VBP1 | ENSG00000155959 | 94,61559417 |
| CUL4A | ENSG00000139842 | 91,54922439 |
| SH3BP5L | ENSG00000175137 | 104,2092659 |
| UGCG | ENSG00000148154 | 145,7021763 |
| MAN2C1 | ENSG00000140400 | 102,8475803 |
| C2orf49 | ENSG00000135974 | 104,4433234 |
| TAF1C | ENSG00000103168 | 111,6308136 |
| TNPO3 | ENSG00000064419 | 107,0429251 |

| | | |
|---|---|---|
| RBM42 | ENSG00000126254 | 78,28181548 |
| R3HDM2 | ENSG00000179912 | 112,1692374 |
| TOB1 | ENSG00000141232 | 123,3520885 |
| EDC4 | ENSG00000038358 | 90,18456 |
| NAP1L4P1 | ENSG00000177173 | 136,842256 |
| CCNYL1 | ENSG00000163249 | 85,07465215 |
| LILRA1 ENSG00000104974 | ENSG00000104974 | 99,74922544 |
| PHRF1 ENSG00000070047 | ENSG00000070047 | 82,46829772 |
| PMPCB | ENSG00000105819 | 106,4180378 |
| DHX16 ENSG00000204560 | ENSG00000204560 | 93,95559746 |
| ANAPC13 | ENSG00000129055 | 118,609419 |
| CD99 | ENSG00000002586 | 70,64643056 |
| NINJ2 | ENSG00000171840 | 91,489906 |
| SLC7A11 | ENSG00000151012 | 94,1719072 |
| CCT7 | ENSG00000135624 | 78,49251932 |
| LSMEM1 | ENSG00000181016 | 159,8853658 |
| PLOD1 | ENSG00000083444 | 110,159933 |
| PSMD1 | ENSG00000173692 | 84,44965061 |
| R3HCC1L | ENSG00000166024 | 120,7276248 |
| EPOR | ENSG00000187266 | 131,309276 |
| CNTNAP3B | ENSG00000154529 | 145,3991392 |
| NUP160 | ENSG00000030066 | 98,67175594 |
| M6PR | ENSG00000003056 | 97,64301291 |
| NEK6 | ENSG00000119408 | 82,94856526 |
| MLLT1 | ENSG00000130382 | 107,0930571 |
| C5orf15 | ENSG00000113583 | 112,037728 |
| MPP7 | ENSG00000150054 | 120,9259332 |
| ZNF586 | ENSG00000083828 | 111,650676 |
| ZBED5 | ENSG00000236287 | 102,8655237 |
| VPS29 | ENSG00000111237 | 101,5088312 |
| INPP1 | ENSG00000151689 | 134,6380207 |
| TC2N ENSG00000165929 | ENSG00000165929 | 119,8588323 |
| RBM26 | ENSG00000139746 | 110,0894311 |
| PARK7 | ENSG00000116288 | 90,21280556 |
| AFG2B | ENSG00000171763 | 97,87820838 |
| UBA2 | ENSG00000126261 | 84,24769549 |
| RGPD2 | ENSG00000185304 | 115,0298681 |
| LIN7C | ENSG00000148943 | 84,37326927 |
| DCTN5 | ENSG00000166847 | 79,85760558 |
| TRMO | ENSG00000136932 | 121,5354814 |
| CYCS | ENSG00000172115 | 92,4413294 |
| RPS19BP1 | ENSG00000187051 | 77,56343363 |
| TADA3 | ENSG00000171148 | 81,18924683 |
| CCR3 | ENSG00000183625 | 142,8854399 |
| SCFD1 | ENSG00000092108 | 103,1594783 |
| GDAP2 | ENSG00000196505 | 130,4882197 |
| CCDC28A | ENSG00000024862 | 90,42987101 |
| RUNDC1 | ENSG00000198863 | 74,8334518 |
| SBDS | ENSG00000126524 | 125,541047 |
| PDIA6 | ENSG00000143870 | 106,551714 |
| CD3E | ENSG00000198851 | 64,86600547 |
| BRMS1 | ENSG00000174744 | 82,88787463 |
| DCTN2 | ENSG00000175203 | 110,8229483 |
| MAGED2 | ENSG00000102316 | 72,3397918 |
| REV1 | ENSG00000135945 | 122,4918286 |

| | | |
|---|---|---|
| PPM1D | ENSG00000170836 | 85,74193986 |
| MLX | ENSG00000108788 | 95,91408425 |
| IQCE | ENSG00000106012 | 78,1350653 |
| SHC1 | ENSG00000160691 | 86,36315145 |
| PSMB6 | ENSG00000142507 | 92,23094654 |
| HLA-H ENSG00000206341 | ENSG00000206341 | 128,5957866 |
| LRPPRC | ENSG00000138095 | 75,4701298 |
| MTHFR | ENSG00000177000 | 78,5315622 |
| ENSG00000237513 | ENSG00000237513 | 84,56861916 |
| B3GNT8 | ENSG00000177191 | 101,1362862 |
| MARCHF5 | ENSG00000198060 | 114,0010297 |
| GCN1 | ENSG00000089154 | 82,74402727 |
| ATG4B | ENSG00000168397 | 115,9790977 |
| VTI1A | ENSG00000151532 | 97,85105349 |
| AIDA | ENSG00000186063 | 106,1639492 |
| PTGR3 | ENSG00000180011 | 83,35013066 |
| BTBD7 ENSG00000011114 | ENSG00000011114 | 106,808643 |
| ARHGEF3 | ENSG00000163947 | 92,44174048 |
| FBXO46 | ENSG00000177051 | 90,54277458 |
| METTL22 | ENSG00000067365 | 120,8794365 |
| SLC25A36 | ENSG00000114120 | 111,5751438 |
| ITM2A | ENSG00000078596 | 78,58071507 |
| ADCY7 | ENSG00000121281 | 85,09882969 |
| TMX2 | ENSG00000213593 | 93,43277644 |
| PLPBP | ENSG00000147471 | 104,9984921 |
| UBE2K | ENSG00000078140 | 111,882805 |
| SPTLC2 | ENSG00000100596 | 87,03086216 |
| SGMS1 | ENSG00000198964 | 83,33058814 |
| MEX3C | ENSG00000176624 | 87,41867038 |
| LRRC75A | ENSG00000181350 | 82,61922371 |
| SLC30A1 | ENSG00000170385 | 62,59739978 |
| CGAS | ENSG00000164430 | 85,78694893 |
| FCHSD1 | ENSG00000197948 | 108,567132 |
| USP48 | ENSG00000090686 | 117,6514121 |
| CNOT9 | ENSG00000144580 | 88,7183712 |
| CDC27 | ENSG00000004897 | 106,0702248 |
| CAMK1G | ENSG00000008118 | 134,9645527 |
| LRCH3 | ENSG00000186001 | 88,92619097 |
| ABL1 | ENSG00000097007 | 62,47091541 |
| PLCD1 | ENSG00000187091 | 51,07976902 |
| CCNG1 | ENSG00000113328 | 93,51727192 |
| CNIH1 | ENSG00000100528 | 108,2266879 |
| TBC1D22A | ENSG00000054611 | 91,46079281 |
| PI4K2B | ENSG00000038210 | 95,40761917 |
| ARHGAP1 | ENSG00000175220 | 64,37828871 |
| CNOT3 ENSG00000088038 | ENSG00000088038 | 105,4804008 |
| RAB39B | ENSG00000155961 | 86,48413348 |
| PSIP1 | ENSG00000164985 | 107,7947811 |
| PRDX1 | ENSG00000117450 | 86,48309571 |
| CSNK2A1 | ENSG00000101266 | 89,31912933 |
| CD4 | ENSG00000010610 | 104,0980379 |
| ZNF451 | ENSG00000112200 | 104,9922884 |
| CYTOR | ENSG00000222041 | 105,6711623 |
| UBE3C | ENSG00000009335 | 73,37803502 |
| TNIK | ENSG00000154310 | 96,284604 |

| | | |
|---|---|---|
| RNF169 | ENSG00000166439 | 104,9922304 |
| METTL23 | ENSG00000181038 | 103,3218233 |
| EIF5B | ENSG00000158417 | 102,3344415 |
| ZMAT3 | ENSG00000172667 | 101,885525 |
| HSF1 ENSG00000185122 | ENSG00000185122 | 104,9194103 |
| KAT6B ENSG00000156650 | ENSG00000156650 | 99,91004615 |
| THOC2 | ENSG00000125676 | 113,147076 |
| AKTIP | ENSG00000166971 | 112,9884893 |
| SLAMF7 | ENSG00000026751 | 70,8240349 |
| UBR5-DT | ENSG00000246263 | 96,19346635 |
| MLH1 | ENSG00000076242 | 89,57925766 |
| WHAMM | ENSG00000156232 | 99,1240309 |
| SLC26A2 | ENSG00000155850 | 133,8857124 |
| TOMM40L | ENSG00000158882 | 143,9796874 |
| AUP1 | ENSG00000115307 | 79,61816617 |
| SIDT2 | ENSG00000149577 | 98,45537971 |
| FRMD3 | ENSG00000172159 | 71,04428735 |
| DDX41 | ENSG00000183258 | 111,6651549 |
| LRRK1 | ENSG00000154237 | 82,82726495 |
| LRFN1 | ENSG00000128011 | 91,65107496 |
| RANBP6 | ENSG00000137040 | 108,5043811 |
| MOB4 | ENSG00000115540 | 89,65230121 |
| CNOT6 | ENSG00000113300 | 108,7434343 |
| ZDHHC2 | ENSG00000104219 | 123,4692506 |
| BLZF1 | ENSG00000117475 | 111,1350162 |
| BAX | ENSG00000087088 | 93,46820201 |
| KIAA1143 ENSG00000163807 | ENSG00000163807 | 76,52720782 |
| ARSA | ENSG00000100299 | 103,3894624 |
| LSM14B | ENSG00000149657 | 71,61851579 |
| TRAF4 | ENSG00000076604 | 71,60589409 |
| HEG1 | ENSG00000173706 | 68,65655506 |
| NXT1 | ENSG00000132661 | 119,532197 |
| DESI1 | ENSG00000100418 | 97,89840025 |
| JAZF1 | ENSG00000153814 | 108,1223796 |
| ADORA2A-AS1 | ENSG00000178803 | 138,9780851 |
| ZNF335 | ENSG00000198026 | 86,3387245 |
| DNAJC13 | ENSG00000138246 | 98,26696174 |
| CCDC88C | ENSG00000015133 | 62,74031186 |
| DDX39A | ENSG00000123136 | 84,01983353 |
| TNS1 | ENSG00000079308 | 83,89144074 |
| MXD4 | ENSG00000123933 | 54,63653204 |
| ANKS1A | ENSG00000064999 | 91,83727742 |
| MIB2 | ENSG00000197530 | 75,79891812 |
| TMBIM4 | ENSG00000155957 | 104,8660826 |
| HMCES | ENSG00000183624 | 90,53524887 |
| STIM2 | ENSG00000109689 | 120,910919 |
| PPHLN1 | ENSG00000134283 | 119,1601772 |
| PDE12 | ENSG00000174840 | 97,41765631 |
| CUL1 | ENSG00000055130 | 96,41120757 |
| ETV7 | ENSG00000010030 | 78,56341092 |
| SYNJ2 | ENSG00000078269 | 104,6831121 |
| PSMD12 | ENSG00000197170 | 111,5249419 |
| RPRD1A | ENSG00000141425 | 121,7512029 |
| GPR157 | ENSG00000180758 | 64,09427428 |
| LY75 | ENSG00000054219 | 76,61154341 |

| | | |
|---|---|---|
| ABI3 | ENSG00000108798 | 64,74217122 |
| KATNIP | ENSG00000047578 | 96,07809687 |
| MTHFD2 | ENSG00000065911 | 108,8699051 |
| AMFR | ENSG00000159461 | 70,80770504 |
| SUGP2 | ENSG00000064607 | 90,53227448 |
| DYRK2 | ENSG00000127334 | 78,04965937 |
| HAX1 | ENSG00000143575 | 102,8782387 |
| GPD2 | ENSG00000115159 | 115,4357939 |
| PPP6R2 | ENSG00000100239 | 83,06934392 |
| PRPSAP1 | ENSG00000161542 | 86,46438344 |
| SMARCC2 | ENSG00000139613 | 86,7508022 |
| ZNF830 | ENSG00000198783 | 80,68521843 |
| FBN1 | ENSG00000166147 | 74,38438278 |
| TNIP2 | ENSG00000168884 | 58,48122518 |
| RENBP | ENSG00000102032 | 68,93997217 |
| SLC8A1 | ENSG00000183023 | 131,4597653 |
| PAPSS1 | ENSG00000138801 | 118,3582638 |
| WDR37 | ENSG00000047056 | 101,9838284 |
| AQR | ENSG00000021776 | 87,67941438 |
| OPLAH | ENSG00000178814 | 112,5951741 |
| FOXK1 | ENSG00000164916 | 73,31171296 |
| SLC10A3 | ENSG00000126903 | 72,94433219 |
| VDAC1 | ENSG00000213585 | 80,42868609 |
| SLC35E3 | ENSG00000175782 | 51,00606322 |
| SSBP3 | ENSG00000157216 | 83,58527957 |
| SH2D2A | ENSG00000027869 | 53,6139891 |
| AFDN | ENSG00000130396 | 70,34174099 |
| ABCF3 | ENSG00000161204 | 99,09380154 |
| GZMH | ENSG00000100450 | 36,67673671 |
| ALDH9A1 | ENSG00000143149 | 65,64041678 |
| CCSAP | ENSG00000154429 | 88,18068838 |
| GABPB1-IT1 | ENSG00000285410 | 91,37783278 |
| DSTN | ENSG00000125868 | 106,2288248 |
| PEX2 | ENSG00000164751 | 88,74660179 |
| MTREX | ENSG00000039123 | 75,19133299 |
| TM6SF1 | ENSG00000136404 | 95,60115579 |
| ZNF706 | ENSG00000120963 | 99,26757742 |
| USP11 | ENSG00000102226 | 87,04851469 |
| KCNK6 | ENSG00000099337 | 103,0911291 |
| UQCRH | ENSG00000173660 | 89,12459601 |
| TRAPPC3 | ENSG00000054116 | 94,59807821 |
| ZNF627 | ENSG00000198551 | 86,99945101 |
| PILRB | ENSG00000121716 | 106,5232252 |
| HINT3 | ENSG00000111911 | 81,41458767 |
| COPS5 | ENSG00000121022 | 96,80476858 |
| EIF1AD | ENSG00000175376 | 78,07757834 |
| VAT1 | ENSG00000108828 | 105,2916075 |
| ELK1 | ENSG00000126767 | 68,7701578 |
| HACD4 | ENSG00000188921 | 126,7507521 |
| EIF2AK3 | ENSG00000172071 | 113,0398693 |
| PLK2 | ENSG00000145632 | 68,15177683 |
| CCAR1 | ENSG00000060339 | 86,80262711 |
| MIR142HG | ENSG00000265206 | 92,99838829 |
| CD52 | ENSG00000169442 | 84,0822712 |
| CCT8 | ENSG00000156261 | 79,44475691 |

| | | |
|---|---|---|
| CLSTN1 | ENSG00000171603 | 68,83451977 |
| CYB5R1 | ENSG00000159348 | 114,150415 |
| SGSH | ENSG00000181523 | 100,3669204 |
| VCPKMT | ENSG00000100483 | 122,435989 |
| NINJ2-AS1 | ENSG00000177406 | 82,90065609 |
| EXOC7 | ENSG00000182473 | 109,0271897 |
| RPA1 | ENSG00000132383 | 75,4214287 |
| KEAP1 | ENSG00000079999 | 63,63496852 |
| NOLC1 | ENSG00000166197 | 80,05624316 |
| PSMA5 | ENSG00000143106 | 80,57365273 |
| RBSN | ENSG00000131381 | 93,0827152 |
| BCL2 | ENSG00000171791 | 93,18226817 |
| C1orf52 | ENSG00000162642 | 109,2210887 |
| SMIM15 | ENSG00000188725 | 90,53219996 |
| DLGAP4 | ENSG00000080845 | 101,947919 |
| SOD1 | ENSG00000142168 | 74,60152325 |
| PPIB | ENSG00000166794 | 83,29818183 |
| CCT6A | ENSG00000146731 | 89,60404905 |
| ANAPC2 | ENSG00000176248 | 75,11776528 |
| FAM111A | ENSG00000166801 | 107,1596573 |
| IL10RB | ENSG00000243646 | 86,85020765 |
| NLRC3 | ENSG00000167984 | 60,64658087 |
| PIGB | ENSG00000069943 | 112,7086581 |
| WLS | ENSG00000116729 | 98,79394724 |
| RAB37 | ENSG00000172794 | 95,1723257 |
| TOPBP1 | ENSG00000163781 | 95,83605808 |
| MCM3AP | ENSG00000160294 | 97,15571271 |
| TSC2 | ENSG00000103197 | 82,25787578 |
| GTDC1 | ENSG00000121964 | 114,4451287 |
| FBXO3 | ENSG00000110429 | 100,1724019 |
| RASSF1 | ENSG00000068028 | 64,22422824 |
| BTBD9 | ENSG00000183826 | 74,90772228 |
| CSNK2A2 | ENSG00000070770 | 74,72686112 |
| POFUT1 | ENSG00000101346 | 50,54417449 |
| ATMIN | ENSG00000166454 | 66,19334376 |
| EIF2AK1 | ENSG00000086232 | 75,59230103 |
| SYS1 | ENSG00000204070 | 79,36775349 |
| DHCR7 | ENSG00000172893 | 137,1152393 |
| SIKE1 | ENSG00000052723 | 96,08367055 |
| TSN | ENSG00000211460 | 78,26213597 |
| SEC23IP | ENSG00000107651 | 93,45707025 |
| USP42 | ENSG00000106346 | 91,22712686 |
| PRRG4 | ENSG00000135378 | 62,49573315 |
| POLR2C | ENSG00000102978 | 93,51666375 |
| BUB3 | ENSG00000154473 | 90,02148763 |
| IDH3B | ENSG00000101365 | 92,2170398 |
| TCFL5 | ENSG00000101190 | 60,1831079 |
| QARS1 | ENSG00000172053 | 72,39181994 |
| PPWD1 | ENSG00000113593 | 108,3847513 |
| EMSY | ENSG00000158636 | 98,67453918 |
| DOK2 | ENSG00000147443 | 51,90396392 |
| IGKC | ENSG00000211592 | 79,377439 |
| FXR2 | ENSG00000129245 | 87,35183139 |
| SERPING1 | ENSG00000149131 | 67,6559935 |
| NISCH | ENSG00000010322 | 86,99136238 |

| | | |
|---|---|---|
| LOXHD1 | ENSG00000167210 | 74,96715625 |
| SEMA4B | ENSG00000185033 | 71,01484799 |
| ATP2B1-AS1 | ENSG00000271614 | 100,7228662 |
| MEI1 | ENSG00000167077 | 83,53143696 |
| F13A1 | ENSG00000124491 | 77,99309269 |
| ZC3H12C | ENSG00000149289 | 50,42609385 |
| ZBTB33 | ENSG00000177485 | 92,42901355 |
| ZNF83 | ENSG00000167766 | 74,02885793 |
| SLC30A5 | ENSG00000145740 | 99,88776302 |
| PIGG | ENSG00000174227 | 83,29953392 |
| IGBP1 | ENSG00000089289 | 79,98333172 |
| BAZ1A-AS1 | ENSG00000258738 | 96,48105636 |
| C7orf50 | ENSG00000146540 | 43,02390754 |
| LCK | ENSG00000182866 | 64,35908122 |
| ZNF506 | ENSG00000081665 | 107,3904956 |
| CCT4 | ENSG00000115484 | 72,12828683 |
| SPPL3 | ENSG00000157837 | 96,0430665 |
| UHRF2 | ENSG00000147854 | 101,552181 |
| CEBPZ | ENSG00000115816 | 84,80711595 |
| COMMD6 | ENSG00000188243 | 77,39723087 |
| ALAS1 | ENSG00000023330 | 139,8220217 |
| GYG1 | ENSG00000163754 | 125,9090228 |
| CHD6 | ENSG00000124177 | 73,75694852 |
| CDC123 | ENSG00000151465 | 137,923919 |
| ZNF18 | ENSG00000154957 | 86,91469612 |
| COG3 | ENSG00000136152 | 91,41101392 |
| SLC35A5 | ENSG00000138459 | 86,72003906 |
| CNDP2 | ENSG00000133313 | 73,67512711 |
| KXD1 | ENSG00000105700 | 88,75270017 |
| GVINP1 | ENSG00000254838 | 73,08739157 |
| TIPRL | ENSG00000143155 | 100,0117612 |
| NDUFB1 | ENSG00000183648 | 89,13996896 |
| SEC61B | ENSG00000106803 | 77,56258003 |
| CAPN1 | ENSG00000014216 | 80,59044292 |
| CRY2 | ENSG00000121671 | 63,209071 |
| MED23 | ENSG00000112282 | 97,0348014 |
| GSKIP | ENSG00000100744 | 83,7219284 |
| ACP6 | ENSG00000162836 | 123,1001642 |
| HBEGF | ENSG00000113070 | 132,8846774 |
| RARA-AS1 | ENSG00000265666 | 111,4359196 |
| SLFN13 | ENSG00000154760 | 120,3867075 |
| UEVLD | ENSG00000151116 | 84,09205553 |
| IL3RA | ENSG00000185291 | 126,8111742 |
| SARS1 | ENSG00000031698 | 92,25950146 |
| HPS3 | ENSG00000163755 | 83,23358452 |
| ST3GAL5 | ENSG00000115525 | 66,96021141 |
| BRD7 | ENSG00000166164 | 97,40751228 |
| SELENOO | ENSG00000073169 | 104,0770459 |
| MAML2 | ENSG00000184384 | 93,97549906 |
| ADAM28 | ENSG00000042980 | 97,58284845 |
| VAMP1 | ENSG00000139190 | 88,22707133 |
| RNF20 | ENSG00000155827 | 81,25445453 |
| PDE4DIP | ENSG00000178104 | 52,53198578 |
| FRS2 | ENSG00000166225 | 78,06661898 |
| PPP2R5E | ENSG00000154001 | 94,31463454 |

| | | |
|---|---|---|
| CD3D | ENSG00000167286 | 56,52943768 |
| ZMYM5 | ENSG00000132950 | 95,55646332 |
| CMSS1 | ENSG00000184220 | 68,18623115 |
| CLK2 ENSG00000176444 | ENSG00000176444 | 82,72385638 |
| RNPEP | ENSG00000176393 | 76,68659953 |
| TNFAIP8L1 | ENSG00000185361 | 71,90131199 |
| PPP1R8 | ENSG00000117751 | 71,74123136 |
| AP5M1 | ENSG00000053770 | 81,64945898 |
| NPIPB5 ENSG00000243716 | ENSG00000243716 | 96,55217772 |
| ELP2 | ENSG00000134759 | 84,83474949 |
| CDC26 | ENSG00000176386 | 100,1038886 |
| CYB561A3 | ENSG00000162144 | 49,91532798 |
| ZCCHC8 | ENSG00000033030 | 77,03438811 |
| LTF | ENSG00000012223 | 79,64514537 |
| DCAF15 | ENSG00000132017 | 86,24175224 |
| LUC7L2 | ENSG00000146963 | 122,3532694 |
| CDC40 | ENSG00000168438 | 76,64959776 |
| CAMK4 | ENSG00000152495 | 92,97270416 |
| GCLM | ENSG00000023909 | 165,1500221 |
| CERS5 | ENSG00000139624 | 96,53546089 |
| RILP ENSG00000167705 | ENSG00000167705 | 95,57277392 |
| RPRD2 | ENSG00000163125 | 72,91188143 |
| SERTAD1 | ENSG00000197019 | 83,17402832 |
| RASA1 | ENSG00000145715 | 117,0834117 |
| LIPA | ENSG00000107798 | 84,77551538 |
| KLC1 | ENSG00000126214 | 88,96639739 |
| AP3M1 | ENSG00000185009 | 73,80406335 |
| ZNF385A | ENSG00000161642 | 91,39540371 |
| SRGAP2 | ENSG00000266028 | 77,03582629 |
| PAXBP1 ENSG00000159086 | ENSG00000159086 | 116,8683566 |
| ADAMTSL4-AS1 | ENSG00000203804 | 107,2216893 |
| TMEM245 | ENSG00000106771 | 70,27704997 |
| PSMD7 | ENSG00000103035 | 84,7193481 |
| PPM1G | ENSG00000115241 | 80,64653606 |
| SH2D1B | ENSG00000198574 | 63,55575135 |
| MILR1 | ENSG00000271605 | 98,09979215 |
| SDC2 | ENSG00000169439 | 120,3400024 |
| C4orf33 | ENSG00000151470 | 117,9108582 |
| RCHY1 | ENSG00000163743 | 96,18198781 |
| RNGTT | ENSG00000111880 | 112,2300182 |
| ENSG00000215022 | ENSG00000215022 | 140,6035366 |
| LINC02649 | ENSG00000215244 | 106,8532453 |
| AHR | ENSG00000106546 | 73,95572836 |
| HLA-DMA ENSG00000204257 | ENSG00000204257 | 78,21852742 |
| OFD1 | ENSG00000046651 | 87,45974813 |
| ALDH2 | ENSG00000111275 | 77,47504869 |
| VPS13A | ENSG00000197969 | 84,9862861 |
| CABLES2 | ENSG00000149679 | 87,20955509 |
| BLCAP | ENSG00000166619 | 68,88120898 |
| SLC5A3 | ENSG00000198743 | 93,87314154 |
| CITED2 | ENSG00000164442 | 73,22168627 |
| CCDC88A | ENSG00000115355 | 87,80885562 |
| DNM1L | ENSG00000087470 | 72,06761824 |
| HECTD4 | ENSG00000173064 | 98,39728329 |
| SPIN1 | ENSG00000106723 | 100,8083154 |

| | | |
|---|---|---|
| DENND1A | ENSG00000119522 | 91,65176976 |
| ENSG00000265975 | ENSG00000265975 | 100,210229 |
| NDUFA6 ENSG00000184983 | ENSG00000184983 | 89,94063234 |
| LEF1 | ENSG00000138795 | 92,16896431 |
| DCTN1 | ENSG00000204843 | 76,19394746 |
| COL6A2 | ENSG00000142173 | 33,49069439 |
| SNX17 | ENSG00000115234 | 63,25329977 |
| EIF3M | ENSG00000149100 | 85,86406993 |
| POLR2J | ENSG00000005075 | 62,05015467 |
| ZNF146 | ENSG00000167635 | 75,05771041 |
| SYTL1 | ENSG00000142765 | 86,19793293 |
| TMEM45B | ENSG00000151715 | 77,78425058 |
| C1RL | ENSG00000139178 | 102,3076253 |
| NSUN4 | ENSG00000117481 | 80,51281046 |
| WAC-AS1 | ENSG00000254635 | 62,18456281 |
| DMXL1 | ENSG00000172869 | 96,63436855 |
| TRPM2 | ENSG00000142185 | 106,6952321 |
| PATL2 ENSG00000229474 | ENSG00000229474 | 97,46402667 |
| TNNI2 | ENSG00000130598 | 98,49439427 |
| PSMD5 | ENSG00000095261 | 77,5221694 |
| PMEPA1 | ENSG00000124225 | 75,32495204 |
| POLR1F | ENSG00000105849 | 92,53630869 |
| TRAPPC4 ENSG00000196655 | ENSG00000196655 | 84,73766254 |
| MCM6 | ENSG00000076003 | 69,6014019 |
| PIGT | ENSG00000124155 | 84,34679131 |
| MTHFS | ENSG00000136371 | 93,32670116 |
| LRBA | ENSG00000198589 | 68,78558227 |
| SIK2 | ENSG00000170145 | 107,9106844 |
| LRRC8D | ENSG00000171492 | 51,82573117 |
| TBC1D14 | ENSG00000132405 | 99,44075576 |
| ZBTB38 | ENSG00000177311 | 61,55505981 |
| YTHDF1 | ENSG00000149658 | 69,35840487 |
| ERCC6 | ENSG00000225830 | 84,23060719 |
| LILRB1 ENSG00000104972 | ENSG00000104972 | 108,4263953 |
| CDKN2AIP | ENSG00000168564 | 53,03396058 |
| PSMB7 | ENSG00000136930 | 66,91817227 |
| SFXN3 | ENSG00000107819 | 89,83656057 |
| FAM157D ENSG00000282572 | ENSG00000282572 | 103,447509 |
| SND1 | ENSG00000197157 | 64,67777758 |
| POGK | ENSG00000143157 | 63,55282227 |
| CPVL | ENSG00000106066 | 85,3275668 |
| ZCCHC3 | ENSG00000247315 | 50,55955794 |
| FBXW5 | ENSG00000159069 | 56,76646807 |
| ZMPSTE24 | ENSG00000084073 | 91,24423866 |
| GALNT6 | ENSG00000139629 | 58,83347577 |
| TRIP11 | ENSG00000100815 | 86,75256382 |
| TMEM184C | ENSG00000164168 | 72,73926942 |
| ENSG00000255026 | ENSG00000255026 | 77,65333796 |
| SOS1 | ENSG00000115904 | 100,1333176 |
| CRTAP | ENSG00000170275 | 84,25450785 |
| PINK1 | ENSG00000158828 | 59,58276354 |
| RGS10 | ENSG00000148908 | 76,69196913 |
| SPCS1 | ENSG00000114902 | 89,88787581 |
| CEBPA | ENSG00000245848 | 45,90658577 |
| EPRS1 | ENSG00000136628 | 58,49160949 |

| | | |
|---|---|---|
| SULT1B1 | ENSG00000173597 | 126,1567719 |
| CAMLG | ENSG00000164615 | 79,54917737 |
| ZNF250 | ENSG00000196150 | 120,6320057 |
| MTOR | ENSG00000198793 | 89,29097004 |
| SPINT2 | ENSG00000167642 | 96,97654883 |
| CLMN | ENSG00000165959 | 67,10258949 |
| ZNF622 | ENSG00000173545 | 65,59698423 |
| GPI ENSG00000105220 | ENSG00000105220 | 86,80198383 |
| FBL ENSG00000105202 | ENSG00000105202 | 59,75629913 |
| AZI2 | ENSG00000163512 | 104,7788853 |
| TRIM27 ENSG00000204713 | ENSG00000204713 | 67,26879851 |
| XPOT | ENSG00000184575 | 80,33963746 |
| ZNF264 | ENSG00000083844 | 60,64049656 |
| TNPO2 | ENSG00000105576 | 69,86377623 |
| APBA3 | ENSG00000011132 | 76,35005006 |
| NACC1 | ENSG00000160877 | 73,2637063 |
| TCF4 | ENSG00000196628 | 82,26533372 |
| S1PR5 | ENSG00000180739 | 55,85373614 |
| AGPAT4 | ENSG00000026652 | 51,37234821 |
| ZYG11B | ENSG00000162378 | 87,95564299 |
| ST3GAL6 | ENSG00000064225 | 73,6498807 |
| TPST1 | ENSG00000169902 | 76,97138659 |
| LATS1 | ENSG00000131023 | 85,93561751 |
| BORCS5 ENSG00000165714 | ENSG00000165714 | 124,4287744 |
| TTC19 | ENSG00000011295 | 89,08828289 |
| SLC16A5 | ENSG00000170190 | 70,05722927 |
| STK16 | ENSG00000115661 | 92,54237759 |
| NOP58 | ENSG00000055044 | 63,13991111 |
| PHKA2 | ENSG00000044446 | 102,9425429 |
| ING1 | ENSG00000153487 | 65,28893076 |
| AKAP8L | ENSG00000011243 | 81,50061065 |
| YTHDF2 | ENSG00000198492 | 78,52541621 |
| OLIG1 | ENSG00000184221 | 54,47910961 |
| MTSS1 | ENSG00000170873 | 65,74811873 |
| PANX2 | ENSG00000073150 | 115,6034182 |
| QTRT2 | ENSG00000151576 | 78,02426153 |
| HTATIP2 | ENSG00000109854 | 74,95500475 |
| SLC2A1 | ENSG00000117394 | 90,34746489 |
| ENSG00000255823 | ENSG00000255823 | 99,14637573 |
| AKAP8 | ENSG00000105127 | 67,95163068 |
| TRERF1 | ENSG00000124496 | 98,02123405 |
| GALNT2 | ENSG00000143641 | 62,41954613 |
| EOMES | ENSG00000163508 | 44,77702987 |
| COQ2 | ENSG00000173085 | 100,3415282 |
| CNOT11 | ENSG00000158435 | 86,62618893 |
| GTF3A | ENSG00000122034 | 84,91622224 |
| TMEM39A | ENSG00000176142 | 59,84305475 |
| CARS2 | ENSG00000134905 | 89,7841356 |
| HS1BP3 | ENSG00000118960 | 61,18076348 |
| HCFC1 | ENSG00000172534 | 59,62681656 |
| CD81 | ENSG00000110651 | 76,05175567 |
| ZNF445 | ENSG00000185219 | 93,53397335 |
| GIMAP6 | ENSG00000133561 | 66,0908599 |
| WDR20 | ENSG00000140153 | 66,80602697 |
| SLC12A7 ENSG00000113504 | ENSG00000113504 | 46,64578141 |

| | | |
|---|---|---|
| HERC4 | ENSG00000148634 | 106,8653963 |
| RAB44 | ENSG00000255587 | 90,54951359 |
| INTS11 | ENSG00000127054 | 77,28987135 |
| ZNF518A | ENSG00000177853 | 91,079326 |
| SNX14 | ENSG00000135317 | 94,20430841 |
| CUL2 | ENSG00000108094 | 109,2246599 |
| SPACA6 | ENSG00000182310 | 63,94342351 |
| GORASP2 | ENSG00000115806 | 56,94718302 |
| GSR | ENSG00000104687 | 84,68805699 |
| CACYBP | ENSG00000116161 | 61,61736395 |
| RNPC3 | ENSG00000185946 | 111,5305232 |
| PDK3 | ENSG00000067992 | 87,84330044 |
| RNF6 | ENSG00000127870 | 76,96673922 |
| RFC1 | ENSG00000035928 | 73,74618028 |
| NEBL | ENSG00000078114 | 44,53167963 |
| FBXL20 | ENSG00000108306 | 79,06903545 |
| XPO7 | ENSG00000130227 | 72,41642538 |
| CEP19 | ENSG00000174007 | 77,64227313 |
| CIDECP1 | ENSG00000186162 | 72,64793194 |
| UQCRC1 | ENSG00000010256 | 73,88767327 |
| TTC14 | ENSG00000163728 | 87,66123846 |
| WDFY2 | ENSG00000139668 | 85,64655577 |
| ADGRE4P | ENSG00000268758 | 117,2916817 |
| CLN3 | ENSG00000188603 | 74,69361295 |
| CYTH2 | ENSG00000105443 | 78,67392034 |
| APOL3 | ENSG00000128284 | 70,06146219 |
| MRPL44 | ENSG00000135900 | 64,62011655 |
| NMT1 | ENSG00000136448 | 63,60014327 |
| PKNOX1 | ENSG00000160199 | 89,30121253 |
| NECAB2 | ENSG00000103154 | 61,09449545 |
| KIF1C | ENSG00000129250 | 69,85470889 |
| AGER ENSG00000204305 | ENSG00000204305 | 101,8243899 |
| TMEM120B | ENSG00000188735 | 87,55108582 |
| MITD1 | ENSG00000158411 | 77,59005941 |
| P2RY2 | ENSG00000175591 | 88,15733712 |
| CEP95 | ENSG00000258890 | 97,2487107 |
| AKR1A1 | ENSG00000117448 | 56,12840942 |
| CD2AP | ENSG00000198087 | 97,29155532 |
| NSUN2 | ENSG00000037474 | 79,71101175 |
| ATP5MC2 | ENSG00000135390 | 71,92761973 |
| TIMM17B | ENSG00000126768 | 92,65488119 |
| CRBN | ENSG00000113851 | 88,64636619 |
| ZNF766 | ENSG00000196214 | 71,32250505 |
| CLP1 | ENSG00000172409 | 71,24075717 |
| DNMT1 | ENSG00000130816 | 61,27222704 |
| CCR5 | ENSG00000160791 | 66,63847878 |
| LDLRAD4 | ENSG00000168675 | 77,0953981 |
| KIF2A | ENSG00000068796 | 83,22782939 |
| CABIN1 ENSG00000099991 | ENSG00000099991 | 81,50482297 |
| RETREG1 | ENSG00000154153 | 81,76134936 |
| GLCCI1 | ENSG00000106415 | 69,91266926 |
| REXO1 | ENSG00000079313 | 71,3362022 |
| ENSG00000286147 | ENSG00000286147 | 108,1599061 |
| DECR1 | ENSG00000104325 | 101,8291614 |
| GNG5 | ENSG00000174021 | 89,77255714 |

| | | |
|---|---|---|
| MFAP3 | ENSG00000037749 | 93,0247398 |
| ZFYVE1 | ENSG00000165861 | 68,94017021 |
| EEF1A1P5 | ENSG00000196205 | 69,96381544 |
| SNAI3 | ENSG00000185669 | 74,82433702 |
| UBXN11 | ENSG00000158062 | 61,24092731 |
| TMEM52B | ENSG00000165685 | 134,2884961 |
| IFI30 | ENSG00000216490 | 122,3602332 |
| LRRC4 | ENSG00000128594 | 80,93628454 |
| TAF12 | ENSG00000120656 | 89,76003887 |
| RAB33B | ENSG00000172007 | 63,67515564 |
| MTF2 | ENSG00000143033 | 85,3974543 |
| PEDS1 | ENSG00000240849 | 65,25933155 |
| SLC39A7 ENSG00000112473 | ENSG00000112473 | 69,0838054 |
| UBQLN4 | ENSG00000160803 | 99,25598493 |
| EDEM2 | ENSG00000088298 | 68,81470469 |
| RNF2 | ENSG00000121481 | 93,35529968 |
| NIPBL-DT | ENSG00000285967 | 84,83360716 |
| RIMOC1 | ENSG00000205765 | 94,24254878 |
| CRIPT | ENSG00000119878 | 79,37240963 |
| EXT1 | ENSG00000182197 | 61,29277618 |
| DHRSX | ENSG00000169084 | 84,74678534 |
| NAAA | ENSG00000138744 | 109,1513684 |
| MIB1 | ENSG00000101752 | 92,38202792 |
| RMC1 | ENSG00000141452 | 78,21166404 |
| RRAGA | ENSG00000155876 | 61,62068495 |
| SCAP | ENSG00000114650 | 71,58176377 |
| PARP1 | ENSG00000143799 | 60,52488245 |
| CDK14 | ENSG00000058091 | 79,67277981 |
| RNH1 ENSG00000023191 | ENSG00000023191 | 75,63421247 |
| PCMTD2 ENSG00000203880 | ENSG00000203880 | 61,39885648 |
| LRIG1 | ENSG00000144749 | 53,66662342 |
| LPIN1 | ENSG00000134324 | 81,706457 |
| KLHL9 | ENSG00000198642 | 70,97218651 |
| RCBTB2 | ENSG00000136161 | 75,88730806 |
| NDUFS2 | ENSG00000158864 | 79,02613013 |
| CD84 | ENSG00000066294 | 83,04403652 |
| LSM3 | ENSG00000170860 | 80,0251472 |
| NFE2 | ENSG00000123405 | 90,08486566 |
| NSA2 | ENSG00000164346 | 69,07397531 |
| POLR3E ENSG00000058600 | ENSG00000058600 | 74,58576944 |
| EFTUD2 | ENSG00000108883 | 83,06956409 |
| ACP3 | ENSG00000014257 | 75,51226318 |
| ADORA2A | ENSG00000128271 | 69,08154007 |
| POLDIP2 | ENSG00000004142 | 52,9192693 |
| SCAF4 | ENSG00000156304 | 88,06319282 |
| TMEM65 | ENSG00000164983 | 98,82830649 |
| CCT2 | ENSG00000166226 | 66,53131543 |
| GLE1 | ENSG00000119392 | 80,13524889 |
| MTR | ENSG00000116984 | 70,82668094 |
| HBS1L | ENSG00000112339 | 101,5087572 |
| MED30 | ENSG00000164758 | 81,57626624 |
| DENND2D | ENSG00000162777 | 77,88417838 |
| PDPR | ENSG00000090857 | 64,99962848 |
| MS4A7 | ENSG00000166927 | 70,81330374 |
| ZNF7 | ENSG00000147789 | 69,33018293 |

| | | |
|---|---|---|
| SIN3B | ENSG00000127511 | 78,64362428 |
| TBC1D8 | ENSG00000204634 | 57,49587685 |
| ASB2 ENSG00000100628 | ENSG00000100628 | 68,7634107 |
| COLGALT1 | ENSG00000130309 | 96,75595154 |
| KIAA0319L | ENSG00000142687 | 96,31495252 |
| RAB30 | ENSG00000137502 | 67,50697772 |
| AMZ2P1 | ENSG00000214174 | 119,5230707 |
| RTP4 | ENSG00000136514 | 40,78312973 |
| TUFM | ENSG00000178952 | 53,50172595 |
| CCNJL | ENSG00000135083 | 84,81958571 |
| AMMECR1L | ENSG00000144233 | 57,96508149 |
| TSC22D1 | ENSG00000102804 | 54,90354229 |
| VAPB | ENSG00000124164 | 74,47400015 |
| ECHDC1 | ENSG00000093144 | 79,97022063 |
| ARHGAP12 | ENSG00000165322 | 84,33016553 |
| LUZP1 | ENSG00000169641 | 71,83867445 |
| KAT5 | ENSG00000172977 | 94,25456302 |
| SLC30A4 | ENSG00000104154 | 64,96617914 |
| PRKRIP1 | ENSG00000128563 | 73,65820079 |
| HARS1 | ENSG00000170445 | 79,88120781 |
| POC1B | ENSG00000139323 | 114,5251652 |
| NDUFA10 | ENSG00000130414 | 69,12796645 |
| KCNQ1OT1 | ENSG00000269821 | 104,3315943 |
| PDLIM2 | ENSG00000120913 | 73,4790613 |
| DDX59 | ENSG00000118197 | 90,39072127 |
| CTSW | ENSG00000172543 | 45,00512076 |
| WASH6P | ENSG00000182484 | 70,58305728 |
| SEPHS2 | ENSG00000179918 | 62,09467901 |
| ENSG00000279500 | ENSG00000279500 | 50,45833319 |
| LSM8 | ENSG00000128534 | 90,37755993 |
| PHACTR2 | ENSG00000112419 | 89,56734856 |
| TRAF3IP2 | ENSG00000056972 | 74,52564213 |
| PTPN18 | ENSG00000072135 | 74,18416258 |
| CDH23 | ENSG00000107736 | 85,34969024 |
| LGMN | ENSG00000100600 | 60,21461995 |
| NMD3 | ENSG00000169251 | 114,3616833 |
| SRP68 | ENSG00000167881 | 60,70450471 |
| UFL1 | ENSG00000014123 | 78,75996706 |
| ERVK3-1 | ENSG00000142396 | 106,6607343 |
| FCHO1 | ENSG00000130475 | 67,58301617 |
| GNPTG | ENSG00000090581 | 75,36011302 |
| HTRA2 | ENSG00000115317 | 42,78928472 |
| PRDX3P1 | ENSG00000229598 | 131,3429683 |
| PTPN4 | ENSG00000088179 | 59,05336148 |
| NDUFS5 | ENSG00000168653 | 75,96066967 |
| TSEN54 | ENSG00000182173 | 48,4760431 |
| TBC1D9B | ENSG00000197226 | 50,39448092 |
| DDHD1 | ENSG00000100523 | 92,31664474 |
| RHOT1 | ENSG00000126858 | 103,3956771 |
| ADRM1 | ENSG00000130706 | 57,39968018 |
| MAP3K14 ENSG00000006062 | ENSG00000006062 | 58,70409336 |
| BTF3L4 | ENSG00000134717 | 81,5918827 |
| KLHDC2 | ENSG00000165516 | 59,33201717 |
| LILRB5 ENSG00000105609 | ENSG00000105609 | 101,9828045 |
| ATP11C | ENSG00000101974 | 88,37189071 |

| | | |
|---|---|---|
| ENSG00000272449 | ENSG00000272449 | 32,78281881 |
| CNTRL | ENSG00000119397 | 86,38812573 |
| TEPSIN | ENSG00000167302 | 72,12161407 |
| RPL26 | ENSG00000161970 | 59,87284249 |
| MIAT | ENSG00000225783 | 75,52751012 |
| SRI | ENSG00000075142 | 81,99052118 |
| CCDC159 | ENSG00000183401 | 77,92369195 |
| FGFBP2 | ENSG00000137441 | 37,96409476 |
| ATL2 | ENSG00000119787 | 113,4943777 |
| ZNF710-AS1 | ENSG00000259291 | 91,3522335 |
| PGPEP1 | ENSG00000130517 | 77,20102947 |
| KLHDC8B | ENSG00000185909 | 47,91446949 |
| NDUFA4 | ENSG00000189043 | 75,52054202 |
| DUSP22 | ENSG00000112679 | 64,87325278 |
| PSMC1 | ENSG00000100764 | 68,93934965 |
| TARS1 | ENSG00000113407 | 78,08981373 |
| ASNSD1 | ENSG00000138381 | 69,98443818 |
| ELOC | ENSG00000154582 | 74,92641903 |
| MAPKAP1 | ENSG00000119487 | 77,64286063 |
| THUMPD3 | ENSG00000134077 | 87,91424041 |
| OTULINL | ENSG00000145569 | 84,03081277 |
| PTP4A3 ENSG00000184489 | ENSG00000184489 | 74,99498391 |
| LHFPL2 | ENSG00000145685 | 65,81906091 |
| AGRN | ENSG00000188157 | 65,24597511 |
| SLC31A1 | ENSG00000136868 | 72,55881739 |
| F2R | ENSG00000181104 | 49,49474409 |
| ENSG00000272669 | ENSG00000272669 | 95,20156485 |
| MTURN | ENSG00000180354 | 67,19724262 |
| NUP133 | ENSG00000069248 | 64,06182646 |
| PHLPP1 | ENSG00000081913 | 72,88853423 |
| RANBP3 | ENSG00000031823 | 61,08919284 |
| SNHG1 | ENSG00000255717 | 61,34893892 |
| CRTC3 | ENSG00000140577 | 68,00541329 |
| ZBTB26 | ENSG00000171448 | 68,26618419 |
| PIK3R5-DT | ENSG00000266389 | 72,31282537 |
| TRPS1 | ENSG00000104447 | 78,62019035 |
| SLC25A11 | ENSG00000108528 | 62,5095063 |
| SSR4 | ENSG00000180879 | 57,42262617 |
| FDFT1 ENSG00000079459 | ENSG00000079459 | 85,0203476 |
| RBM15B | ENSG00000259956 | 49,61600429 |
| PARN ENSG00000140694 | ENSG00000140694 | 104,8203247 |
| PARP6 | ENSG00000137817 | 82,86800039 |
| WDR11 | ENSG00000120008 | 64,32260671 |
| NCKAP5L | ENSG00000167566 | 83,97524654 |
| MTG2 | ENSG00000101181 | 68,18911479 |
| COPS8 | ENSG00000198612 | 69,46653225 |
| ZSWIM4 | ENSG00000132003 | 66,00974293 |
| TECPR1 | ENSG00000205356 | 66,24840851 |
| GNL2 | ENSG00000134697 | 103,0001213 |
| AKAP12 | ENSG00000131016 | 76,52793703 |
| MACO1 | ENSG00000204178 | 74,10046225 |
| KRCC1 | ENSG00000172086 | 62,3206296 |
| NUTM2B-AS1 | ENSG00000225484 | 80,98864047 |
| SLC37A2 | ENSG00000134955 | 60,20766152 |
| LSG1 | ENSG00000041802 | 58,29192987 |

| | | |
|---|---|---|
| WSB2 | ENSG00000176871 | 79,56069586 |
| MED11 | ENSG00000161920 | 82,68301595 |
| PRDX3 | ENSG00000165672 | 94,10730544 |
| AKAP11 | ENSG00000023516 | 57,12133589 |
| DAPK3 | ENSG00000167657 | 66,91670913 |
| CNNM3 | ENSG00000168763 | 48,14217615 |
| PLXND1 | ENSG00000004399 | 33,68967533 |
| SLFN11 | ENSG00000172716 | 65,68774928 |
| RFX2 | ENSG00000087903 | 95,87953954 |
| SNRPB2 | ENSG00000125870 | 83,2705685 |
| LNPK | ENSG00000144320 | 101,1852398 |
| TRPV2 | ENSG00000187688 | 90,85964015 |
| KLHL18 | ENSG00000114648 | 82,33698436 |
| SNRPD2 | ENSG00000125743 | 63,76666386 |
| IL21R | ENSG00000103522 | 54,87690479 |
| SOCS4 | ENSG00000180008 | 91,8430427 |
| ANKDD1A | ENSG00000166839 | 116,2343176 |
| DENR | ENSG00000139726 | 72,69330363 |
| DENND4C | ENSG00000137145 | 65,36099209 |
| PRUNE1 | ENSG00000143363 | 67,12347435 |
| ZNF107 | ENSG00000196247 | 136,5369768 |
| ERLEC1 | ENSG00000068912 | 78,56020928 |
| EMP1 | ENSG00000134531 | 132,8675677 |
| OPA3 | ENSG00000125741 | 60,73533004 |
| ZNF317 | ENSG00000130803 | 57,68897052 |
| MRPL28 | ENSG00000086504 | 64,41811535 |
| CADM4 | ENSG00000105767 | 75,89830067 |
| C11orf54 | ENSG00000182919 | 59,77489655 |
| ACBD5 | ENSG00000107897 | 61,79954685 |
| IPO8 | ENSG00000133704 | 85,55973799 |
| ZNF326 | ENSG00000162664 | 74,32775651 |
| TMEM63A | ENSG00000196187 | 82,83123093 |
| DNAJC21 | ENSG00000168724 | 72,65160792 |
| DUSP11 | ENSG00000144048 | 69,16581821 |
| TGS1 | ENSG00000137574 | 68,07528199 |
| MED21 | ENSG00000152944 | 65,70164526 |
| ATXN3 | ENSG00000066427 | 70,42384 |
| FTH1P20 | ENSG00000226564 | 70,94639429 |
| ITM2C | ENSG00000135916 | 41,82060064 |
| CYTH3 | ENSG00000008256 | 78,16224346 |
| LILRB4 ENSG00000186818 | ENSG00000186818 | 92,73744512 |
| TOB2 | ENSG00000183864 | 68,90689373 |
| CFAP58 | ENSG00000120051 | 61,30218651 |
| ERH | ENSG00000100632 | 72,80645802 |
| KCTD5 | ENSG00000167977 | 56,82694016 |
| BICDL1 | ENSG00000135127 | 50,41711621 |
| ACOT13 | ENSG00000112304 | 74,64641554 |
| SORT1 | ENSG00000134243 | 70,52407419 |
| STK19 ENSG00000204344 | ENSG00000204344 | 69,47098532 |
| CDIP1 ENSG00000089486 | ENSG00000089486 | 67,38416911 |
| PCNT | ENSG00000160299 | 57,04089447 |
| C1orf21 | ENSG00000116667 | 40,82245771 |
| CSTF2T | ENSG00000177613 | 55,18630347 |
| ENSG00000255328 | ENSG00000255328 | 86,83071409 |
| NELL2 | ENSG00000184613 | 71,72568261 |

| | | |
|---|---|---|
| MDN1 | ENSG00000112159 | 55,40619705 |
| TREML4 | ENSG00000188056 | 95,10879486 |
| NOSIP | ENSG00000142546 | 59,61584229 |
| HAT1 | ENSG00000128708 | 71,93302538 |
| SNRPG | ENSG00000143977 | 65,07411057 |
| LRRC58 | ENSG00000163428 | 53,91120406 |
| SLC39A13 | ENSG00000165915 | 73,91738118 |
| BRPF1 | ENSG00000156983 | 53,85430297 |
| TOP3A ENSG00000177302 | ENSG00000177302 | 71,29155449 |
| FANCD2 | ENSG00000144554 | 86,45528332 |
| HSPA1B ENSG00000204388 | ENSG00000204388 | 109,9995129 |
| ATP2C2 | ENSG00000064270 | 106,6461632 |
| DESI2 | ENSG00000121644 | 66,02232226 |
| HNRNPUL2 | ENSG00000214753 | 51,04003174 |
| ATP6V1H | ENSG00000047249 | 79,2844608 |
| LSM12 | ENSG00000161654 | 78,36078413 |
| SNX11 | ENSG00000002919 | 53,39987279 |
| ACAT2 | ENSG00000120437 | 63,464997 |
| TANC2 | ENSG00000170921 | 105,17899 |
| PSMB2 | ENSG00000126067 | 64,98358464 |
| DPP9 | ENSG00000142002 | 66,76213571 |
| TAF9B | ENSG00000187325 | 76,20501531 |
| LUC7L | ENSG00000007392 | 91,34887437 |
| STAMBP | ENSG00000124356 | 70,2116181 |
| ZNF254 | ENSG00000213096 | 128,2027523 |
| CFAP45 | ENSG00000213085 | 119,3964093 |
| TMEM88 | ENSG00000167874 | 89,54891741 |
| ANKRA2 | ENSG00000164331 | 62,07699268 |
| RAC1P2 | ENSG00000249936 | 68,59056497 |
| MAN1B1 | ENSG00000177239 | 67,00548472 |
| DOP1B | ENSG00000142197 | 70,74566455 |
| UBE2G2 | ENSG00000184787 | 67,47178749 |
| CPSF6 | ENSG00000111605 | 95,35910542 |
| GATA2 | ENSG00000179348 | 50,50353452 |
| PLAC8 | ENSG00000145287 | 74,69409744 |
| NBPF26 | ENSG00000273136 | 59,49550703 |
| SAV1 | ENSG00000151748 | 56,56150044 |
| USP39 | ENSG00000168883 | 67,93791948 |
| SNX16 | ENSG00000104497 | 53,459689 |
| TNFRSF25 | ENSG00000215788 | 58,01973954 |
| KYNU | ENSG00000115919 | 79,12219955 |
| THAP6 | ENSG00000174796 | 70,46829277 |
| SNF8 | ENSG00000159210 | 56,94132316 |
| EPHA4 | ENSG00000116106 | 44,61867175 |
| WAKMAR2 | ENSG00000237499 | 69,67384059 |
| PPDPF | ENSG00000125534 | 46,21159334 |
| BRD8 | ENSG00000112983 | 55,52869451 |
| PRAM1 | ENSG00000133246 | 68,35156679 |
| MRPL33 | ENSG00000243147 | 87,78100393 |
| GPAA1 | ENSG00000197858 | 74,16787417 |
| UBE3B | ENSG00000151148 | 66,889446 |
| TBC1D17 | ENSG00000104946 | 63,45401026 |
| TAFAZZIN | ENSG00000102125 | 64,86253153 |
| LINC00869 | ENSG00000277147 | 44,74025718 |
| COPS4 | ENSG00000138663 | 66,65675559 |

| | | |
|---|---|---|
| TBL2 | ENSG00000106638 | 71,78783798 |
| NCK2 | ENSG00000071051 | 61,14197726 |
| SYTL2 | ENSG00000137501 | 43,37451591 |
| CNOT4 | ENSG00000080802 | 68,50578802 |
| STYX | ENSG00000198252 | 63,40429686 |
| BTBD10 | ENSG00000148925 | 81,6965207 |
| RABGEF1P1 | ENSG00000229180 | 63,99534298 |
| TAF10 | ENSG00000166337 | 85,55777809 |
| IMPA1 | ENSG00000133731 | 79,25074369 |
| NDUFB6 | ENSG00000165264 | 78,06176429 |
| ENSG00000260257 | ENSG00000260257 | 91,48411813 |
| FBXL13 | ENSG00000161040 | 77,19714542 |
| NAIP ENSG00000249437 | ENSG00000249437 | 114,7517981 |
| MBOAT1 | ENSG00000172197 | 101,7054543 |
| RNF144A | ENSG00000151692 | 66,62511165 |
| ITPR3 | ENSG00000096433 | 54,01203943 |
| MMP24OS | ENSG00000126005 | 63,77843049 |
| TUSC2 | ENSG00000114383 | 50,2482494 |
| BCL11A | ENSG00000119866 | 67,48320669 |
| SLC4A7 | ENSG00000033867 | 60,07775542 |
| TERF2 | ENSG00000132604 | 63,03418021 |
| ENSG00000272501 | ENSG00000272501 | 61,05032099 |
| COG5 ENSG00000164597 | ENSG00000164597 | 80,08396051 |
| KIN | ENSG00000151657 | 74,73161652 |
| TAPBPL | ENSG00000139192 | 60,5602072 |
| KLF9 | ENSG00000119138 | 68,74779656 |
| NFATC2IP | ENSG00000176953 | 74,84714198 |
| ARRDC1 | ENSG00000197070 | 63,63307253 |
| NASP | ENSG00000132780 | 54,6880894 |
| AGPAT3 | ENSG00000160216 | 68,85572461 |
| SPRYD3 | ENSG00000167778 | 43,83386943 |
| NUP205 | ENSG00000155561 | 62,46665587 |
| PHF19 | ENSG00000119403 | 56,16750229 |
| ZFAND2A | ENSG00000178381 | 77,98953976 |
| TRADD | ENSG00000102871 | 57,80335161 |
| GDE1 | ENSG00000006007 | 63,05635756 |
| FNIP2 | ENSG00000052795 | 71,18110105 |
| MXD3 | ENSG00000213347 | 50,87704879 |
| NKAP | ENSG00000101882 | 71,12810507 |
| SNAI1 | ENSG00000124216 | 64,26961907 |
| FAM200B | ENSG00000237765 | 84,97473104 |
| CCT3 | ENSG00000163468 | 54,14897559 |
| OR52K3P | ENSG00000225101 | 73,23867451 |
| LRRC8A | ENSG00000136802 | 57,63343094 |
| TMEM62 | ENSG00000137842 | 59,11725957 |
| MFSD10 | ENSG00000109736 | 49,93762074 |
| DLGAP1-AS1 | ENSG00000177337 | 64,03635918 |
| EXOC4 | ENSG00000131558 | 63,43736081 |
| PPP2CB | ENSG00000104695 | 61,97243817 |
| PPID | ENSG00000171497 | 53,13481154 |
| RNF181 | ENSG00000168894 | 56,09500595 |
| TAOK2 | ENSG00000149930 | 66,1733197 |
| PQBP1 | ENSG00000102103 | 72,63496202 |
| SPIDR | ENSG00000164808 | 69,62487603 |
| ATP5MJ | ENSG00000156411 | 73,19862928 |

| | | |
|---|---|---|
| C6orf120 | ENSG00000185127 | 56,22796797 |
| PYURF | ENSG00000145337 | 60,11400295 |
| CCL20 | ENSG00000115009 | 83,57366399 |
| UCP2 | ENSG00000175567 | 74,66097942 |
| LINC03108 | ENSG00000258082 | 53,80375154 |
| SMARCD1 | ENSG00000066117 | 47,51943411 |
| SETDB1 | ENSG00000143379 | 57,36272556 |
| UBE2O | ENSG00000175931 | 49,27856479 |
| NFATC1 | ENSG00000131196 | 61,34354236 |
| PIP4K2C | ENSG00000166908 | 59,87051658 |
| INTS15 | ENSG00000146576 | 51,88816596 |
| CDIPT | ENSG00000103502 | 55,14028125 |
| RING1 ENSG00000204227 | ENSG00000204227 | 69,92474002 |
| SLC7A1 | ENSG00000139514 | 44,95297465 |
| SAP130 | ENSG00000136715 | 75,25819069 |
| ANO10 | ENSG00000160746 | 70,32849275 |
| ZNF561 | ENSG00000171469 | 53,96588714 |
| YLPM1 | ENSG00000119596 | 58,96633604 |
| CARD11 | ENSG00000198286 | 53,31219454 |
| ACTR1B | ENSG00000115073 | 56,36619294 |
| IMMT | ENSG00000132305 | 61,72169488 |
| PNKP | ENSG00000039650 | 67,28274527 |
| RIN2 | ENSG00000132669 | 51,22792065 |
| ENSG00000225313 | ENSG00000225313 | 103,1022523 |
| DCAF16 | ENSG00000163257 | 69,41554475 |
| ZNF430 ENSG00000118620 | ENSG00000118620 | 71,08913379 |
| SF3B5 | ENSG00000169976 | 49,38012615 |
| PDIA4 | ENSG00000155660 | 56,5369074 |
| ZNF407 | ENSG00000215421 | 87,27377405 |
| COASY | ENSG00000068120 | 61,04185879 |
| RPAP3 | ENSG00000005175 | 65,03870955 |
| HELQ | ENSG00000163312 | 65,46355865 |
| GOLGA1 | ENSG00000136935 | 74,85152364 |
| MICU1 | ENSG00000107745 | 81,44226466 |
| STT3A | ENSG00000134910 | 55,37523686 |
| SLC35A2 | ENSG00000102100 | 74,96195286 |
| PRDM10 | ENSG00000170325 | 91,47721207 |
| ZNF818P | ENSG00000269001 | 45,0465491 |
| ACO2 | ENSG00000100412 | 45,1112517 |
| PRMT9 | ENSG00000164169 | 87,67393852 |
| SIRT6 | ENSG00000077463 | 46,36775039 |
| N4BP3 | ENSG00000145911 | 42,4290228 |
| DNAJC10 | ENSG00000077232 | 76,25921534 |
| CHRNE | ENSG00000108556 | 87,29888776 |
| ZNF431 | ENSG00000196705 | 72,73708378 |
| TTC9 | ENSG00000133985 | 33,465075 |
| MARK4 | ENSG00000007047 | 51,02958873 |
| CYLD-AS1 | ENSG00000261644 | 87,18720378 |
| SSH3 | ENSG00000172830 | 72,62386944 |
| ZNRD2 | ENSG00000173465 | 49,58426085 |
| MINDY1 | ENSG00000143409 | 54,46999784 |
| SNX29 | ENSG00000048471 | 55,75088144 |
| PEAK3 | ENSG00000188305 | 46,32246502 |
| SSB | ENSG00000138385 | 65,16027674 |
| RNF115 | ENSG00000265491 | 55,64119525 |

| | | |
|---|---|---|
| MCRIP1 | ENSG00000225663 | 40,63321834 |
| HSD17B12 | ENSG00000149084 | 70,12311709 |
| PSMD3 | ENSG00000108344 | 51,18807601 |
| PCMT1 | ENSG00000120265 | 64,44066991 |
| LINC01270 | ENSG00000203999 | 85,7253188 |
| ZNF76 | ENSG00000065029 | 77,69299126 |
| FCMR | ENSG00000162894 | 56,03095711 |
| SS18L1 | ENSG00000184402 | 69,55584953 |
| INTS6L | ENSG00000165359 | 90,19204087 |
| PHB2 | ENSG00000215021 | 64,58532245 |
| ORC4 | ENSG00000115947 | 86,90047447 |
| LIG3 | ENSG00000005156 | 55,91410161 |
| TRPM6 | ENSG00000119121 | 87,70032516 |
| UXS1 | ENSG00000115652 | 65,37388741 |
| TRAPPC9 | ENSG00000167632 | 79,35723992 |
| MCUB | ENSG00000005059 | 68,42418659 |
| XPO4 | ENSG00000132953 | 51,34063576 |
| UTP6 | ENSG00000108651 | 65,79076745 |
| COPS3 | ENSG00000141030 | 56,2844756 |
| PLEKHA1 | ENSG00000107679 | 57,07925367 |
| CINP | ENSG00000100865 | 74,6761676 |
| TVP23B | ENSG00000171928 | 70,43245992 |
| MS4A1 | ENSG00000156738 | 67,49397793 |
| ALDH3B1 | ENSG00000006534 | 58,7958538 |
| AIM2 | ENSG00000163568 | 81,55842906 |
| BTNL3 | ENSG00000168903 | 71,45528537 |
| PGRMC2 | ENSG00000164040 | 69,63691274 |
| CDK6 | ENSG00000105810 | 59,7792583 |
| MYO18A | ENSG00000196535 | 69,55818343 |
| TBC1D22A-DT | ENSG00000260708 | 51,07062678 |
| ZCCHC10 | ENSG00000155329 | 75,35727667 |
| GNG7 | ENSG00000176533 | 45,00965028 |
| NCK1 | ENSG00000158092 | 59,1520675 |
| HTATSF1 | ENSG00000102241 | 65,37988714 |
| MSI2 | ENSG00000153944 | 67,98680871 |
| ING4 | ENSG00000111653 | 53,16171024 |
| BIN1 | ENSG00000136717 | 57,96548425 |
| ELOA | ENSG00000011007 | 52,73597667 |
| INTS8 | ENSG00000164941 | 56,32973766 |
| CRY1 | ENSG00000008405 | 63,02126928 |
| PTDSS1 | ENSG00000156471 | 67,30322297 |
| MAVS | ENSG00000088888 | 55,14766857 |
| ZNF252P | ENSG00000196922 | 91,53557113 |
| ANKRD22 | ENSG00000152766 | 76,91487534 |
| GTF2I | ENSG00000263001 | 70,31159992 |
| PSMA6 | ENSG00000100902 | 82,76782938 |
| LINGO3 | ENSG00000220008 | 40,13441 |
| GK4P | ENSG00000178146 | 70,23163518 |
| GTF2F2 | ENSG00000188342 | 81,27331149 |
| WASHC2A | ENSG00000099290 | 65,1790836 |
| POLE3 | ENSG00000148229 | 65,12667322 |
| CD226 | ENSG00000150637 | 73,99429315 |
| ALG13 | ENSG00000101901 | 68,45349934 |
| PURA | ENSG00000185129 | 56,34795663 |
| KCTD2 | ENSG00000180901 | 48,69692295 |

| | | |
|---|---|---|
| SLC1A4 | ENSG00000115902 | 51,60519459 |
| ATP6V1F | ENSG00000128524 | 71,1597728 |
| VPS51 | ENSG00000149823 | 46,22487981 |
| CLCF1 | ENSG00000175505 | 55,63048682 |
| NDUFA5 | ENSG00000128609 | 79,86316936 |
| PFKFB4 | ENSG00000114268 | 71,64395173 |
| PELO | ENSG00000152684 | 51,08884989 |
| ARHGEF11 | ENSG00000132694 | 57,36375814 |
| AMDHD2 | ENSG00000162066 | 53,65880608 |
| HLA-DMB ENSG00000242574 | ENSG00000242574 | 56,09992431 |
| TPCN2 | ENSG00000162341 | 70,61741946 |
| DCAF10 | ENSG00000122741 | 88,56795496 |
| SLC25A51 | ENSG00000122696 | 88,22884367 |
| ADA | ENSG00000196839 | 47,22088073 |
| UQCR10 | ENSG00000184076 | 53,71294399 |
| METAP2 | ENSG00000111142 | 52,83367018 |
| FPR3 | ENSG00000187474 | 55,64069608 |
| TTF1 | ENSG00000125482 | 60,72470772 |
| MCRS1 | ENSG00000187778 | 48,40430925 |
| SNAPC1 | ENSG00000023608 | 94,90037611 |
| VPS37A | ENSG00000155975 | 81,25779785 |
| TSHZ1 | ENSG00000179981 | 44,9294178 |
| TSPAN2 | ENSG00000134198 | 89,6237234 |
| SNX5 | ENSG00000089006 | 71,59025541 |
| ENSG00000283839 | ENSG00000283839 | 50,20894986 |
| CXXC5 | ENSG00000171604 | 54,51474257 |
| VPS11 ENSG00000160695 | ENSG00000160695 | 49,50993221 |
| TP53 | ENSG00000141510 | 49,53857182 |
| SNRNP35 | ENSG00000184209 | 63,30138301 |
| CD38 | ENSG00000004468 | 70,46325904 |
| MDH2 | ENSG00000146701 | 52,38262531 |
| ZMYND11 | ENSG00000015171 | 46,81092238 |
| NBEAL1 | ENSG00000144426 | 81,10801018 |
| CDC25B | ENSG00000101224 | 41,18795901 |
| PDHA1 | ENSG00000131828 | 56,02372824 |
| PHF5A | ENSG00000100410 | 60,54912607 |
| ELAVL1 | ENSG00000066044 | 64,22710838 |
| COMMD2 | ENSG00000114744 | 61,24998412 |
| SERPINE1 | ENSG00000106366 | 64,49207073 |
| FUNDC2 | ENSG00000165775 | 76,24459248 |
| MDM1 | ENSG00000111554 | 44,96943329 |
| BAK1 | ENSG00000030110 | 55,66765948 |
| RABGGTB | ENSG00000137955 | 63,92985649 |
| DKC1 | ENSG00000130826 | 72,70729955 |
| CBX7 | ENSG00000100307 | 42,96696872 |
| CEP120 | ENSG00000168944 | 51,24723524 |
| ATP10D | ENSG00000145246 | 62,35733482 |
| CNR2 | ENSG00000188822 | 67,34938581 |
| NPIPB4 ENSG00000185864 | ENSG00000185864 | 73,83406753 |
| GRPEL1 | ENSG00000109519 | 75,90837536 |
| RASGRP3 | ENSG00000152689 | 89,86982605 |
| ENSG00000278918 | ENSG00000278918 | 74,22951415 |
| TRAK2 | ENSG00000115993 | 67,79619029 |
| SH2D1A | ENSG00000183918 | 47,54384103 |
| SLC37A1 | ENSG00000160190 | 63,51626702 |

| | | |
|---|---|---|
| HDC | ENSG00000140287 | 52,1545194 |
| TSC1 | ENSG00000165699 | 73,16341802 |
| LRRC41 | ENSG00000132128 | 65,1849688 |
| GUSBP11 ENSG00000228315 | ENSG00000228315 | 88,31587565 |
| IMPDH2 | ENSG00000178035 | 49,63329359 |
| DSTYK | ENSG00000133059 | 44,99427977 |
| ZNF417 | ENSG00000173480 | 55,15280303 |
| PWWP2A | ENSG00000170234 | 45,61106096 |
| CDC14A | ENSG00000079335 | 62,55784926 |
| FAM117A | ENSG00000121104 | 68,87180319 |
| TMEM175 | ENSG00000127419 | 59,25670469 |
| ZNF770 | ENSG00000198146 | 55,9164924 |
| FLVCR1 | ENSG00000162769 | 73,92399084 |
| PHACTR4 | ENSG00000204138 | 70,75216661 |
| FAM20B | ENSG00000116199 | 62,83549703 |
| HOMER1 | ENSG00000152413 | 67,60715464 |
| CHORDC1 ENSG00000110172 | ENSG00000110172 | 74,74286662 |
| ZFYVE26 | ENSG00000072121 | 67,25652132 |
| GSTP1 | ENSG00000084207 | 47,82299831 |
| PLCXD2 | ENSG00000240891 | 41,41104677 |
| CEP164 | ENSG00000110274 | 69,93911896 |
| MTND2P28 | ENSG00000225630 | 86,9511815 |
| VMA21 | ENSG00000160131 | 54,94185988 |
| HNRNPAB | ENSG00000197451 | 61,11480899 |
| MRPL49 | ENSG00000149792 | 47,83769445 |
| F2RL1 | ENSG00000164251 | 63,53042507 |
| USP14 | ENSG00000101557 | 59,48005834 |
| CEP192 | ENSG00000101639 | 52,19136524 |
| COL9A3 | ENSG00000092758 | 17,63339001 |
| PIEZO1 | ENSG00000103335 | 40,75709274 |
| TOR4A | ENSG00000198113 | 51,2401002 |
| CDK2AP1 | ENSG00000111328 | 54,24068855 |
| COX8A | ENSG00000176340 | 55,45743164 |
| HMGXB3 | ENSG00000113716 | 58,61105994 |
| PRDM4 | ENSG00000110851 | 50,80160898 |
| NAPG | ENSG00000134265 | 86,24232737 |
| KPNA5 | ENSG00000196911 | 60,62820352 |
| ATP5F1C | ENSG00000165629 | 59,03206689 |
| ZNF211 | ENSG00000121417 | 53,82626255 |
| SMARCA4 | ENSG00000127616 | 61,29756234 |
| CDK5RAP2 | ENSG00000136861 | 72,5093345 |
| CPSF1 ENSG00000071894 | ENSG00000071894 | 56,7143238 |
| SNX30 | ENSG00000148158 | 66,79034504 |
| MANF | ENSG00000145050 | 58,98078039 |
| MELTF | ENSG00000163975 | 62,66511999 |
| DNAAF11 | ENSG00000129295 | 92,68967416 |
| LIN7A | ENSG00000111052 | 82,93896404 |
| LINC00664 | ENSG00000268658 | 65,35922455 |
| SCNM1 | ENSG00000163156 | 58,23066708 |
| ATP5IF1 ENSG00000130770 | ENSG00000130770 | 67,63020434 |
| MOGS | ENSG00000115275 | 42,22594263 |
| ZNF230 | ENSG00000159882 | 73,76007495 |
| SLC9A7 | ENSG00000065923 | 54,02489285 |
| MARCHF2 | ENSG00000099785 | 62,54840337 |
| ATAD1 | ENSG00000138138 | 80,1960901 |

| | | |
|---|---|---|
| NSUN3 | ENSG00000178694 | 57,84678982 |
| TMEM106B | ENSG00000106460 | 107,7656202 |
| POLR1H ENSG00000066379 | ENSG00000066379 | 53,55653083 |
| FCGR2B | ENSG00000072694 | 71,57946598 |
| GRAMD1C | ENSG00000178075 | 83,13456075 |
| APPL1 | ENSG00000157500 | 69,32614533 |
| SAP30L | ENSG00000164576 | 56,28597789 |
| NCLN | ENSG00000125912 | 47,94451279 |
| ABCE1 | ENSG00000164163 | 45,8440148 |
| PCSK7 | ENSG00000160613 | 50,87526502 |
| LMAN1 | ENSG00000074695 | 61,89232909 |
| LINC01127 | ENSG00000281162 | 72,36068562 |
| SPATA2 | ENSG00000158480 | 60,71548928 |
| BAIAP3 | ENSG00000007516 | 55,15222669 |
| NUDC | ENSG00000090273 | 54,66437217 |
| STRN3 | ENSG00000196792 | 82,16641701 |
| RNF175 | ENSG00000145428 | 89,9632048 |
| TTC1 | ENSG00000113312 | 65,37555755 |
| ARHGAP21 | ENSG00000107863 | 54,70085883 |
| ENTPD7 | ENSG00000198018 | 92,30804117 |
| TDG | ENSG00000139372 | 61,31405012 |
| IKZF2 | ENSG00000030419 | 48,4374302 |
| ZNF384 | ENSG00000126746 | 59,05546562 |
| PGAP1 | ENSG00000197121 | 71,20604848 |
| BORCS8 | ENSG00000254901 | 48,77942839 |
| HDAC3 | ENSG00000171720 | 65,85727174 |
| ST20-AS1 | ENSG00000259642 | 63,94840207 |
| PGGT1B | ENSG00000164219 | 62,84424712 |
| STARD8 | ENSG00000130052 | 52,48918272 |
| SMPD2 | ENSG00000135587 | 52,87356055 |
| BTBD2 | ENSG00000133243 | 48,28732478 |
| QSOX2 | ENSG00000165661 | 78,93196569 |
| SYT11 | ENSG00000132718 | 49,66404099 |
| NELFCD | ENSG00000101158 | 63,12260499 |
| MEAF6 | ENSG00000163875 | 67,03982529 |
| PTGER2 | ENSG00000125384 | 40,81043282 |
| FBH1 | ENSG00000134452 | 57,87776399 |
| NFE2L3 | ENSG00000050344 | 83,36771259 |
| POLR1D | ENSG00000186184 | 55,88526229 |
| C17orf107 | ENSG00000205710 | 58,76954179 |
| CICP27 | ENSG00000233750 | 52,6072008 |
| HHEX | ENSG00000152804 | 54,64870319 |
| MAPK9 | ENSG00000050748 | 52,14512343 |
| TXNDC16 | ENSG00000087301 | 59,81447767 |
| AHSA2P | ENSG00000173209 | 71,95181848 |
| MPZ | ENSG00000158887 | 54,80622152 |
| IBTK ENSG00000005700 | ENSG00000005700 | 57,81017219 |
| RAVER1 | ENSG00000161847 | 57,72692243 |
| CAMSAP1 | ENSG00000130559 | 52,80160775 |
| NFATC3 | ENSG00000072736 | 59,96356355 |
| TAF4 ENSG00000130699 | ENSG00000130699 | 49,25246031 |
| VCF1 | ENSG00000133193 | 62,09899475 |
| ATL3 | ENSG00000184743 | 46,02769978 |
| PGAP2 | ENSG00000148985 | 55,7788331 |
| MMGT1 | ENSG00000169446 | 70,54235147 |

| | | |
|---|---|---|
| GRSF1 | ENSG00000132463 | 47,76584983 |
| DCTN3 | ENSG00000137100 | 53,28741546 |
| SEC22C | ENSG00000093183 | 53,86024989 |
| ISY1 | ENSG00000240682 | 53,55080543 |
| DENND10 | ENSG00000119979 | 76,14550258 |
| PRADX | ENSG00000235027 | 61,97118264 |
| FOXK2 | ENSG00000141568 | 60,94217739 |
| THADA | ENSG00000115970 | 60,66480456 |
| RFX1 | ENSG00000132005 | 59,47297642 |
| ACAP3 | ENSG00000131584 | 56,93148762 |
| SMIM12 | ENSG00000163866 | 55,12005255 |
| RALGPS2 | ENSG00000116191 | 87,54329729 |
| SELENOS | ENSG00000131871 | 59,50748817 |
| ATG5 | ENSG00000057663 | 57,51475825 |
| MKRN2 | ENSG00000075975 | 45,50317421 |
| BCL6-AS1 | ENSG00000285938 | 102,9423655 |
| MAF | ENSG00000178573 | 33,85053756 |
| ABCB1 | ENSG00000085563 | 47,44110834 |
| ZNF609 | ENSG00000180357 | 55,41444365 |
| VDAC3 | ENSG00000078668 | 68,18159534 |
| NIPSNAP2 | ENSG00000146729 | 51,55512471 |
| TFDP2 | ENSG00000114126 | 47,10251694 |
| NFE4 | ENSG00000230257 | 55,09465439 |
| PPP2R5B | ENSG00000068971 | 83,17765082 |
| RFWD3 | ENSG00000168411 | 49,67990335 |
| ENSG00000286067 | ENSG00000286067 | 76,66106276 |
| RSU1 | ENSG00000148484 | 69,67037442 |
| FIS1 | ENSG00000214253 | 41,98641805 |
| CBX5 | ENSG00000094916 | 60,74095255 |
| MBTD1 | ENSG00000011258 | 51,04002954 |
| ENSG00000267174 | ENSG00000267174 | 57,29133496 |
| LYPLA2 | ENSG00000011009 | 55,26975614 |
| SNX12 | ENSG00000147164 | 40,30034043 |
| MAP2K6 | ENSG00000108984 | 67,92770742 |
| EIF2B5 | ENSG00000145191 | 61,03037086 |
| CAPN10 | ENSG00000142330 | 49,95530078 |
| DPP7 | ENSG00000176978 | 48,05811225 |
| CD28 | ENSG00000178562 | 56,20815144 |
| CAMP | ENSG00000164047 | 46,22350983 |
| RPS27L | ENSG00000185088 | 66,33393029 |
| ZBTB10 | ENSG00000205189 | 61,26340614 |
| MLEC | ENSG00000110917 | 41,58367101 |
| KIAA0930 | ENSG00000100364 | 50,91968594 |
| ZNF213 | ENSG00000085644 | 42,28982462 |
| SERPINB9P1 | ENSG00000230438 | 57,13686107 |
| KATNB1 | ENSG00000140854 | 40,95588079 |
| RRP36 | ENSG00000124541 | 37,66441607 |
| SH3TC1 | ENSG00000125089 | 37,01391526 |
| PSMB10 | ENSG00000205220 | 44,46277109 |
| TSPYL4 | ENSG00000187189 | 47,17113917 |
| CTNNAL1 | ENSG00000119326 | 64,59013759 |
| FBXO6 | ENSG00000116663 | 47,36664976 |
| SLC22A15 | ENSG00000163393 | 83,25967091 |
| ERICH1 ENSG00000104714 | ENSG00000104714 | 64,5213741 |
| PTPN23 | ENSG00000076201 | 63,71255136 |

| | | |
|---|---|---|
| TXLNA | ENSG00000084652 | 46,73061052 |
| GNL3 | ENSG00000163938 | 47,98249748 |
| SUN1 | ENSG00000164828 | 52,69636905 |
| RGPD5 | ENSG00000015568 | 73,2861035 |
| ENSG00000268555 | ENSG00000268555 | 55,35771354 |
| CFAP92 | ENSG00000114656 | 50,6219778 |
| ZNF467 | ENSG00000181444 | 57,39992851 |
| DOCK9 | ENSG00000088387 | 56,19049516 |
| PPP3CB | ENSG00000107758 | 62,89421027 |
| ACER3 | ENSG00000078124 | 93,61744381 |
| TCERG1 | ENSG00000113649 | 55,69870297 |
| GPR174 | ENSG00000147138 | 43,81620271 |
| JAG1 | ENSG00000101384 | 65,55075077 |
| TSPAN3 | ENSG00000140391 | 63,88360079 |
| PYCARD | ENSG00000103490 | 56,54166194 |
| BCL2L1 | ENSG00000171552 | 74,25145165 |
| RRM1 | ENSG00000167325 | 48,89621526 |
| IARS1 | ENSG00000196305 | 41,79553734 |
| LINC02289 | ENSG00000258819 | 40,85086577 |
| AMBRA1 | ENSG00000110497 | 47,49551347 |
| TRIM41 | ENSG00000146063 | 42,33849318 |
| RBM14 | ENSG00000239306 | 54,33494222 |
| GPR35 | ENSG00000178623 | 54,02649875 |
| PRKCZ | ENSG00000067606 | 56,72974635 |
| RPS6KB2 | ENSG00000175634 | 49,06813571 |
| TMEM205 | ENSG00000105518 | 40,03068285 |
| FHIP1A | ENSG00000164142 | 28,21726125 |
| ARAP1-AS2 | ENSG00000245148 | 93,98065689 |
| NCS1 | ENSG00000107130 | 52,52664807 |
| C12orf76 | ENSG00000174456 | 49,59775343 |
| TRAT1 | ENSG00000163519 | 35,24241337 |
| TBPL1 | ENSG00000028839 | 47,89032149 |
| ENSG00000279821 | ENSG00000279821 | 69,75697979 |
| SSRP1 | ENSG00000149136 | 44,30623897 |
| NUP54 | ENSG00000138750 | 55,15097661 |
| AP3M2 | ENSG00000070718 | 51,30052027 |
| TMEM222 | ENSG00000186501 | 37,25872818 |
| WDR6 | ENSG00000178252 | 53,30512595 |
| DNAJB5 | ENSG00000137094 | 56,41526762 |
| AKT3 ENSG00000117020 | ENSG00000117020 | 44,73729456 |
| PRXL2B ENSG00000157870 | ENSG00000157870 | 33,00880467 |
| TTC39C | ENSG00000168234 | 39,88132102 |
| HLA-DOA ENSG00000204252 | ENSG00000204252 | 34,22977077 |
| RBX1 | ENSG00000100387 | 57,35304981 |
| ZNF140 | ENSG00000196387 | 55,92723724 |
| ATP9B ENSG00000166377 | ENSG00000166377 | 67,20769735 |
| MIOS | ENSG00000164654 | 76,88966174 |
| INAVA | ENSG00000163362 | 53,09053883 |
| DBI | ENSG00000155368 | 48,5158794 |
| TXN2 | ENSG00000100348 | 55,48394625 |
| C3orf38 | ENSG00000179021 | 52,14996935 |
| ZC3H6 | ENSG00000188177 | 61,28816343 |
| BMPR2 | ENSG00000204217 | 57,55726834 |
| NCBP2 | ENSG00000114503 | 49,81156232 |
| SLC25A46 | ENSG00000164209 | 55,67885461 |

| | | |
|---|---|---|
| GRAP2 | ENSG00000100351 | 38,60803158 |
| PKP4 | ENSG00000144283 | 62,51143254 |
| PCED1B-AS1 | ENSG00000247774 | 57,40225501 |
| GXYLT1 | ENSG00000151233 | 64,12376418 |
| ATP5PD | ENSG00000167863 | 55,52028995 |
| TFEC | ENSG00000105967 | 50,74574269 |
| ENSG00000281383 | ENSG00000281383 | 26,08853254 |
| ALG2 | ENSG00000119523 | 61,8274235 |
| UXT | ENSG00000126756 | 55,64618217 |
| UNK | ENSG00000132478 | 52,61104384 |
| EVA1B | ENSG00000142694 | 36,87251747 |
| NONOP2 | ENSG00000237522 | 65,20876568 |
| TTC21A | ENSG00000168026 | 66,47616477 |
| VPS54 | ENSG00000143952 | 72,66518817 |
| ENSG00000233461 | ENSG00000233461 | 73,31646797 |
| IL12RB1 | ENSG00000096996 | 41,0891322 |
| NOP56 | ENSG00000101361 | 54,42965802 |
| HACD2 | ENSG00000206527 | 48,27084131 |
| SMG6 | ENSG00000070366 | 53,94302275 |
| P2RX4 | ENSG00000135124 | 45,89829455 |
| OXCT1 | ENSG00000083720 | 46,88797093 |
| THAP9 | ENSG00000168152 | 70,30336406 |
| DGKQ | ENSG00000145214 | 25,72367605 |
| MTX1 | ENSG00000173171 | 73,83575008 |
| UBOX5 | ENSG00000185019 | 47,96309052 |
| ZNF721 | ENSG00000182903 | 54,89257825 |
| LINC02887 ENSG00000280279 | ENSG00000280279 | 70,93873598 |
| ZCCHC14 | ENSG00000140948 | 64,04102044 |
| SLCO4C1 | ENSG00000173930 | 74,81582176 |
| STRADA | ENSG00000266173 | 64,99112174 |
| CHAF1A | ENSG00000167670 | 34,01986923 |
| BMI1 | ENSG00000168283 | 43,43882159 |
| TRIM13 | ENSG00000204977 | 47,61149423 |
| DYM | ENSG00000141627 | 66,29185751 |
| FYCO1 | ENSG00000163820 | 52,06352492 |
| NTMT1 | ENSG00000148335 | 46,19938207 |
| MED8 | ENSG00000159479 | 53,83088099 |
| DHX29 | ENSG00000067248 | 45,75244227 |
| ZPR1 | ENSG00000109917 | 45,38871106 |
| DENND1B | ENSG00000213047 | 61,48744074 |
| NUCB2 | ENSG00000070081 | 93,51555214 |
| PSMC3 | ENSG00000165916 | 41,46237534 |
| SHISAL2A | ENSG00000182183 | 68,28171312 |
| KBTBD11 ENSG00000176595 | ENSG00000176595 | 65,1285387 |
| TRAF7 | ENSG00000131653 | 35,51663107 |
| POLR2K | ENSG00000147669 | 61,02394186 |
| TRIP4 | ENSG00000103671 | 45,00421982 |
| PPCS | ENSG00000127125 | 55,12229531 |
| STAM | ENSG00000136738 | 77,98086149 |
| NOL8 | ENSG00000198000 | 57,61624074 |
| PREPL | ENSG00000138078 | 37,86093602 |
| ICE2 | ENSG00000128915 | 47,2684435 |
| TMEM208 | ENSG00000168701 | 49,98424235 |
| OSER1-DT | ENSG00000223891 | 47,50869349 |
| TNNT3 | ENSG00000130595 | 80,82420353 |
